# Beyond motif recognition: Specificity of human transcription factors in yeast

**DOI:** 10.64898/2026.04.27.721015

**Authors:** Joshua Bugis, Dan Reuben Zlotnik-Weinberg, Wajd Manadre, Vladimir Mindel, Yunwei Lu, Juan Ignacio Fuxman Bass, Naama Barkai

## Abstract

Transcription factors (TFs) bind DNA through sequence-specific DNA-binding domains (DBDs), yet genome-wide analyses show that TFs occupy only a small fraction of their motif occurrences. This raises the question of how TFs distinguish specific targets from the many potential sites in the genome. To investigate determinants of binding specificity beyond the cognate motif and cofactor influences, we measured the binding of 60 human TFs across the budding yeast genome. Although human TFs robustly recognized their motifs, they displayed strong selectivity in site occupancy. Nucleosome abundance explained this selectivity only in part: among the 5-20% of motif sites that were bound, a substantial fraction remained nucleosome covered. Furthermore, TFs recognizing similar motif sequences independently localized to distinct subsets of sites within different promoters. Despite the absence of human-specific cofactors in yeast, both binding stability and genomic preferences depended on largely disordered non-DBD regions. These findings suggest intrinsically disordered regions (IDRs) may therefore direct genome binding TF target recognition across evolutionarily distant genomes.

**Graphical Abstract:** 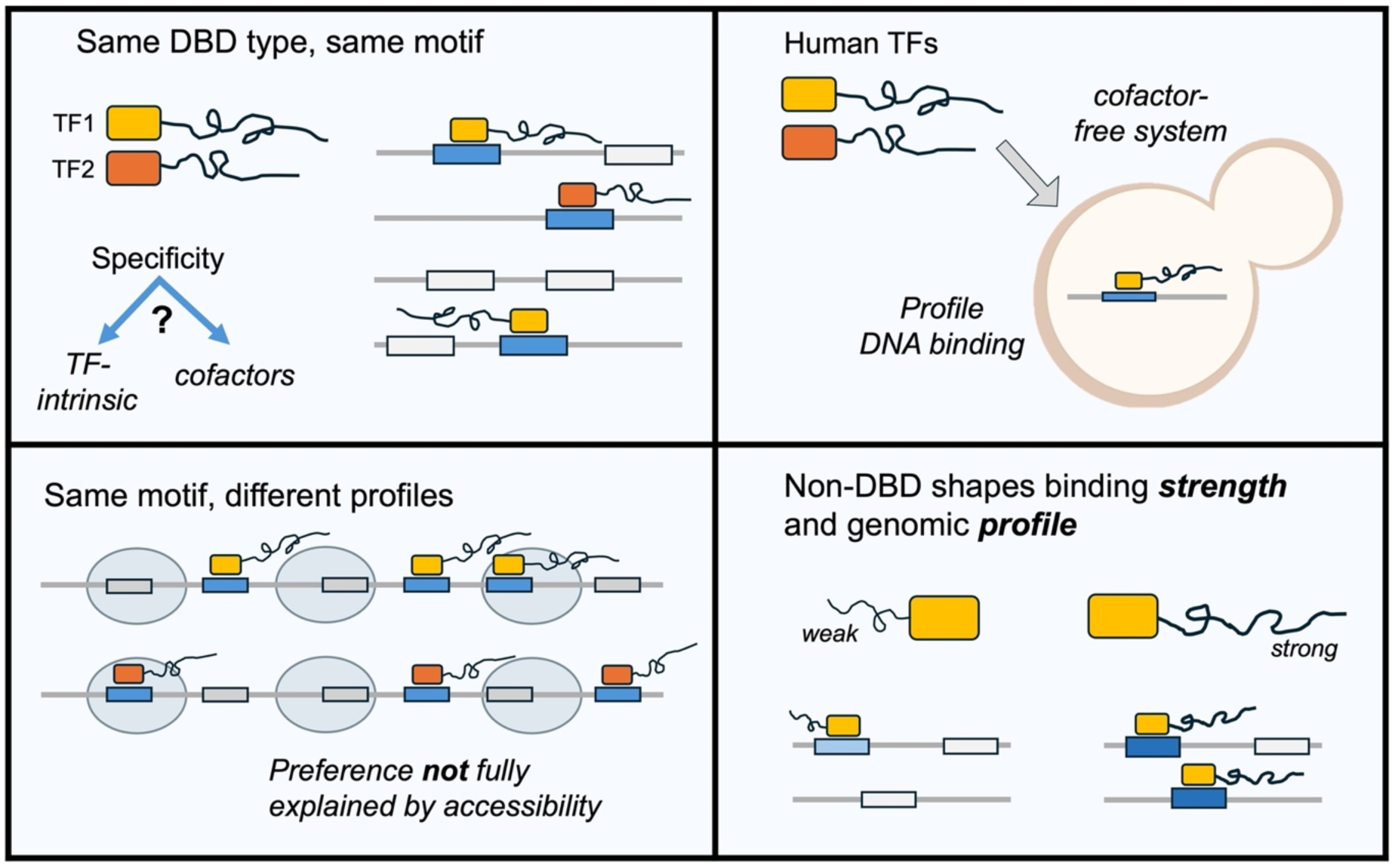

## Introduction

Transcription factors (TFs) regulate gene expression by binding to gene regulatory sequences scattered across genomes. The ability to rapidly and selectively recognize their binding targets is a hallmark of TFs, critical for their role in cellular adaptation, but how this recognition is achieved with high specificity and speed is only partially understood. TF binding sites contain short sequence motifs recognized by DNA-binding domains (DBDs) within TFs. Yet DBD-recognized motifs are highly abundant in genomes, with only a fraction of sites being bound in cells^1,2^. This pronounced disparity between motif abundance and occupancy raises the question of how TFs identify their specific genomic targets amid an overwhelming excess of similar sequences, and how this search process is executed rapidly enough to support timely gene regulation.

Beyond their DBDs, TFs are characterized by extensive intrinsically disordered regions (IDRs) that can span hundreds of residues^3–12^. IDRs remain understudied due to their rapid sequence divergence and the difficulty of experimentally handling structure-lacking domains. Notably, our recent studies in budding yeast point to IDRs as central effectors of TF target recognition^1,13–18^, consistent with related findings in mammalian cells^19–23^. We further noted that IDRs direct genome binding through multiple weak determinants that are dispersed throughout their sequence rather than a single localized region^13,14,16,17,24,25^. However, the molecular interactions through which IDRs direct genome binding remain unclear.

In prevailing models, the selective binding of TFs across their motif sites is explained by DNA accessibility and cooperative interactions^4,26^. DNA accessibility biases TF binding to motifs located within open regions, with chromatin acting as an initial filter that narrows the search space by limiting DNA access. Cooperative interactions complement this by biasing TF binding to sites of multiple motifs bound by respective interacting TFs. In this model, co-binding TFs stabilize each other, either through direct protein-protein contacts or indirectly by changing chromatin accessibility.

IDRs could direct genome binding by interacting with co-binding TFs, chromatin components, or perhaps DNA itself. Protein-protein interaction is best aligned with prevailing models of TF binding specificity; however, this explanation was challenged by systematic efforts to identify interactors of a model IDR-dependent yeast TF, Msn2 where the combined deletion of fourteen TFs that co-localize at Msn2 target promoters had minimal impact on the IDR-driven binding pattern^27^ and we noted a limited effect of motif cooperation^28^. Whether the IDR of Msn2 indeed acts independently of co-binding TFs, and whether this principle extends to other TFs, including in complex eukaryotes where cooperative interactions may play a broader role, remains unclear.

The human genome contains approximately 1600 TFs, compared to about 150 TFs in budding yeast^29,30^. Human TFs are organized into families sharing the same DBD type and recognizing the same consensus motif. For example, all TFs of the GATA family bind to the same, or highly similar motifs, as do all TFs of the forkhead family. This redundancy in motif usage includes TFs with different activities that regulate distinct sets of targets. While this divergence of binding preference could result from interactions with specific co-binding TFs it could also involve more direct recognition of DNA or chromatin features that are intrinsic to the individual TFs within a family and are not captured by the cognate motif.

Defining intrinsic binding preferences of TFs within their native cellular context is confounded by the abundance of potential co-binding interactors, many of which remain unknown. We therefore turned to budding yeast, as a tractable genetic system lacking human-specific proteins. We selected 60 human TFs from 10 DBD motif families and profiled their binding across the yeast genome. Our data revealed that both the binding strength and site preferences of human TFs across the yeast genome depend on their largely disordered non-DBD, and we provide evidence that these effects are independent of nucleosome occupancy or TF interactions. We discuss the implications of our results for understanding the molecular basis of IDR-directed binding and its conservation across eukaryotes.

## Results

### Yeast as a ‘cellular test tube’ for dissecting intrinsic binding preferences of human TFs

Budding yeast presents an experimentally accessible genetic system that enables studying intrinsic properties of human proteins in an environment lacking human-specific interactors^31^. Previously, yeast reporter assays were used to define motif preferences of human TFs, as well as their activation capacity and compare binding at selected human enhancers^32–37^. Here, we extended this approach to directly examine TF binding across the full yeast genome, focusing on binding preferences that are not explained by the TF cognate motifs. We integrated human TFs into the yeast genome, fusing them to yellow fluorescent protein (YFP) to monitor their abundance and nuclear localization, and to MNase to profile their genomic binding by ChEC-seq^38,39^ (Fig. S1A). In ChEC-seq, TF binding locations are defined at near base-pair resolution, by first triggering cleavage of TF-bound DNA using a short calcium pulse that activates the fused MNase and then collecting and sequencing the cleaved DNA fragments. We characterized binding of human TFs in the same conditions previously used to profile a compendium of 147 native yeast TFs and 39 of their DBD-only variants, enabling direct comparisons and the detection of any cross-species TF interactions, if present.

We began with the bZIP factor ATF1 and the ETS factor ERG, as these two well-characterized TFs represent DBD families with and without homology in yeast, respectively. Our ChEC-seq data revealed specific binding of both TFs across the yeast genome (Fig 1A), as indicated by high reproducibility (Fig 1B), and preferred association with their respective cognate motifs (Fig 1C). We further validated motif localization by visualizing the average MNase cleavage profiles around the top 200 bound motif sites, revealing a strong TF footprint (Fig. 1D-E).

**Figure 1:**
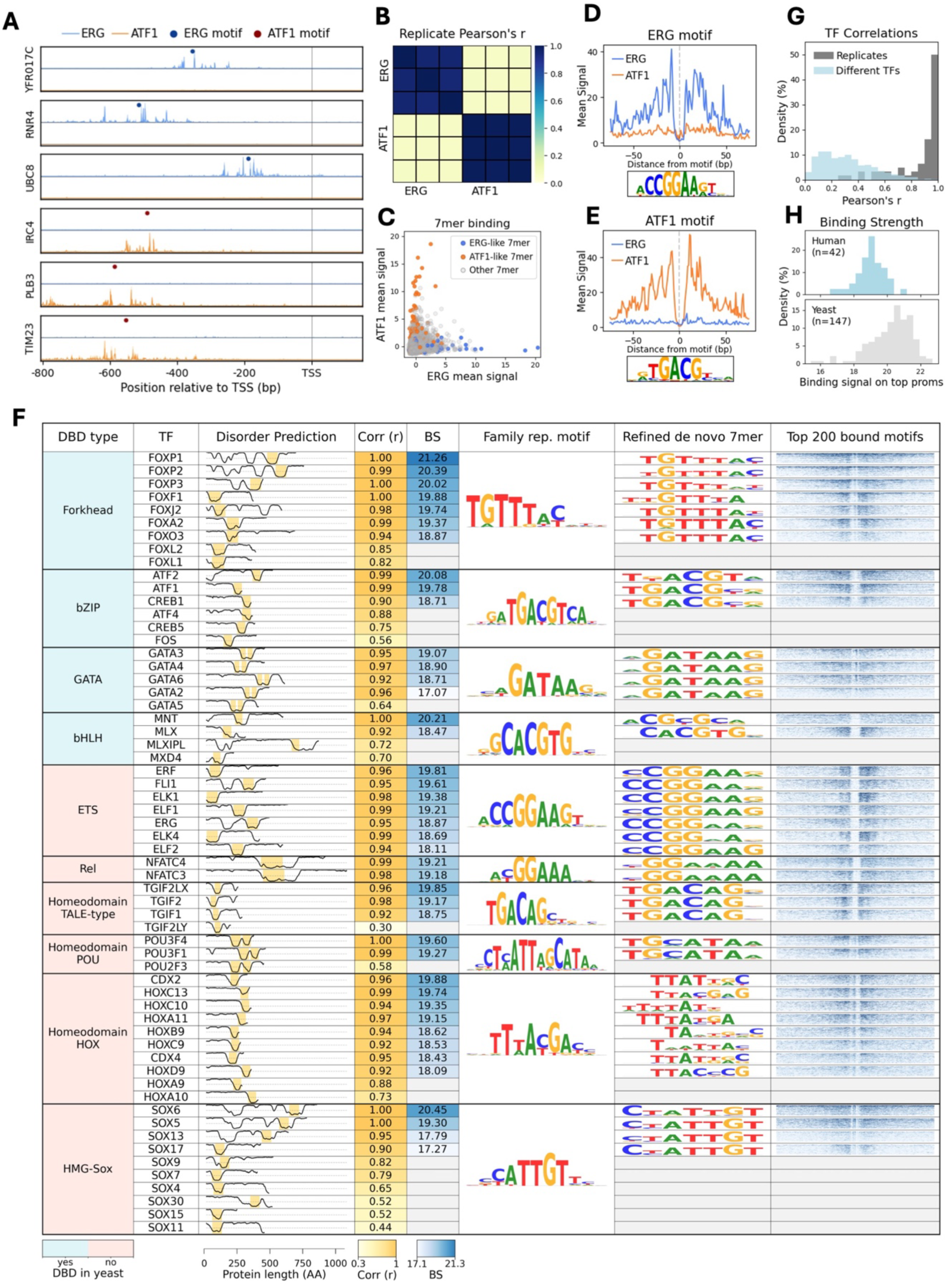
Mapping human TF binding in the budding yeast genome using ChEC-seq. *(1A) Human TFs bind specific promoters in yeast:* ChEC-seq cleavage tracks for the human TFs ERG and ATF1 at representative target promoters. Tracks represent sequencing read cut sites, and dots indicate the location of the HOCOMOCO-defined TF motif. *(1B) ChEC-seq measurements are reproducible across biological repeats:* Pearson correlations are shown between the summed, z-transformed promoter binding signals of three independent ChEC-seq replicates for ERG and ATF1 (see Methods). *(1C) Sequence-specific binding is captured by short k-mer analysis:* For each TF, the mean ChEC-seq signal across all promoter occurrences of every possible 7-mer is shown. 7-mers matching at least 6 out of 7 positions to the consensus ERG (CCGGAAR) or ATF1 (RTGACGY) motifs are highlighted. *(1D-E) Human TFs generate localized cleavage footprints at their cognate motifs:* Mean ChEC-seq signal is shown around the top 200 bound motif instances, aligned by motif center (see Methods). Corresponding sequence logos are shown below each footprint. *(1F) Summary of binding properties across human TFs expressed in yeast:* Shown are DNA-binding domain (DBD) types, predicted protein disorder, and annotated DBD locations (see Methods). The presence or absence of homologous DBDs in yeast is indicated. For each TF, reproducibility (“Corr”) denotes the maximum Pearson correlation between replicates based on summed and z-transformed promoter binding signals. Binding strength is defined as the log_2_ promoter binding signal of top-bound promoters (see Methods). For each DBD family, the representative HOCOMOCO motif is shown together with a refined 7-mer derived from weighted consensus-like matches. Heatmaps show the binding signal around the top 200 motif instances (±100 bp), with color saturation capped at the 92^nd^ percentile of all plotted scores per DBD family. *(1G) Human TFs show higher within-TF similarity than between-TF similarity:* Distributions of maximum replicate correlations for all human TFs (n=60) and between-TF correlations computed from repeat-averaged data for reproducible factors (Pearson’s r>0.9, n=42). *(1H) Human TFs display lower binding strengths compared to native yeast TFs:* Distributions of binding strengths for human and native yeast TFs, calculated as in *(F)*.

Building on this proof of principle, we extended the analysis to a total of 60 human TFs, covering ten DBD families, four of which have homology in yeast (Fig 1F). When selecting individual TFs to test, we prioritized factors with high motif similarity (defined by HOCOMOCO^40^), long IDRs, and minimal additional domains (Fig. 1F). All TFs were integrated into the yeast genome, and their binding was mapped using ChEC-seq with 2-6 replicates per TF. Expression of the YFP-tagged TF was verified by fluorescence microscopy in a representative subset of strains (n=38) (Fig. S1B). Overall, we obtained robust profiles for 42 of 60 TFs tested (70%), 41 of which associated with their cognate motif (Fig. 1F). Binding profiles were highly correlated between replicates of the same TF but differed widely between TFs (Fig. 1G). We conclude that the yeast genome landscape is complex enough to support distinct binding of human TFs.

We noted that TFs of the same motif families differed in their apparent total binding strengths (BS). For example, all seven ETS TFs tested localized to their known common motif, yet when averaged across bound promoters, the total binding strength was 3.2-fold higher (ΔBS (log₂) = 1.70) for ERF as compared to ELF2 (Fig. 1F). While our current data does not include a spike-in calibration, we previously noted that the average binding signal across the strongest bound promoters correlates well with absolute binding strength^17^. Overall, promoter binding signals of human TFs appeared weaker than native yeast TFs (Fig. 1H), suggesting that human TFs may bind the yeast genomes at affinities lower than native proteins.

### Success vs. failure of human TF binding to the yeast genome is explained by obligate dimerization and by stabilizing effects of the largely disordered non-DBD

A subset (18/60) of human TFs failed to bind detectably across the budding yeast genome. We reasoned that these cases could be informative for understanding binding requirements. The bHLH and bZIP DBD families represent exceptional cases, as they bind DNA as obligatory dimers. In these families, the absence of binding is consistent with the lack of a dimerization partner (Fig. 2A). Robust binding in our assay was limited to TFs that bind DNA as homodimers^41^, including ATF1, ATF2, or CREB1 from the bZIP family, or MLX and MNT from the bHLH family. By contrast, TFs that rely on heterodimeric partners^41^, such as the bZIP TFs FOS and ATF4, or the bHLH TFs MLXIPL and MXD4, failed to bind reproducibly across the yeast genome. CREB5 was an exception, as it was shown to bind DNA as homodimer yet failed to profile in our assay. Importantly, yeast contains both bZIP and bHLH TFs, but these did not rescue the binding of human TFs, suggesting that cross-species dimers in these families were not formed.

**Figure 2:**
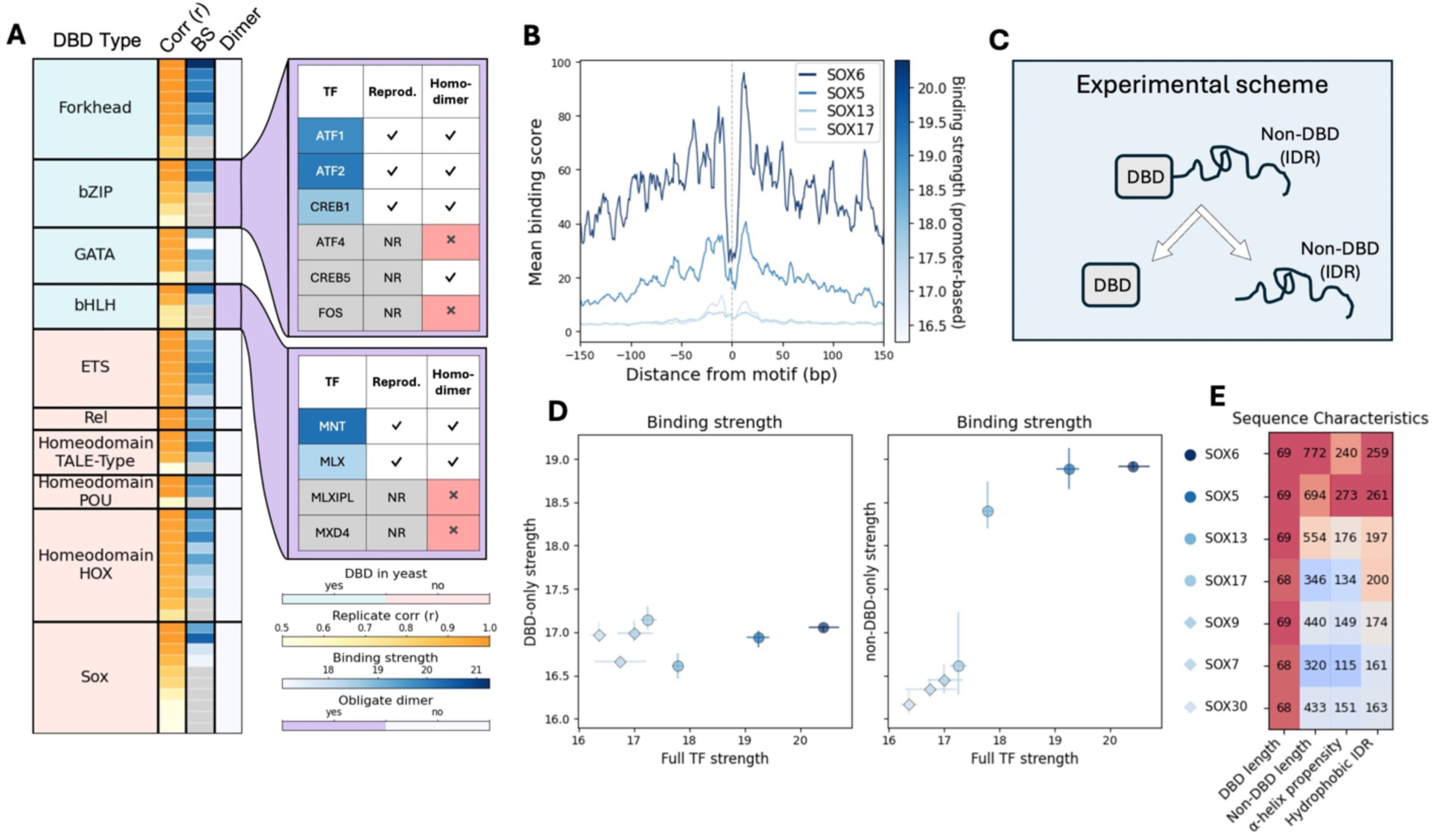
Binding success depends on dimerization status and stabilization by the non-DBD. *(2A) Failure to dimerize is associated with loss of detectable DNA binding:* Summary table of all human TF DNA-binding domain (DBD) families tested in yeast (n=60 TFs). For each TF, the maximum replicate correlation of summed, z-transformed promoter binding scores (as in Fig. 1) and the binding strength are shown, together with the known dimerization mode of the DBD^41^ (obligate dimer versus non-obligate). Zoomed-in tables highlight the two obligate-dimer DBD families. For individual TFs in these families, binding strength (blue shading), reproducibility (defined as Pearson’s r > 0.9), and the ability to bind DNA as a homodimer are indicated. All TFs shown are capable of heterodimerization, which is not explicitly displayed. *(2B) Members of the SOX DBD family span a wide range of binding strengths:* Shown are superimposed ChEC-seq footprints (see Methods) aligned by motif center to the HOCOMOCO-defined representative SOX motif, smoothed with a 5-bp sliding window mean. TF footprints are colored by promoter-based binding strength, computed per replicate and then averaged per TF. *(2C) Separation of DBD and non-DBD regions enables dissection of binding contributions:* Schematic portraying the creation of DBD-only and non-DBD-only strains for seven selected SOX TFs, representing strong binding TFs (n=2), weak binding TFs (n=2), and non-reproducible TFs (n=3). *(2D) Non-DBD binding strength predicts full-length TF binding:* Binding strengths of full-length TFs are compared with those of the corresponding DBD-only (left) or non-DBD-only (right) constructs. Binding strength was calculated as in *(B)*. Lines indicate the range from the minimum to maximum values across replicates. Circular and diamond markers represent reproducible and non-reproducible TFs, respectively. Markers and lines are colored as in *(B)*. *(2E) Strong binders are characterized by longer non-DBDs and specific sequence features:* Heatmaps summarize properties of the amino acid sequences of the SOX TFs, including DBD and non-DBD length (defined by InterProScan), the number of amino acids with predicted α-helix propensity (Chou–Fasman algorithm), and the number of hydrophobic residues within intrinsically disordered regions (IDRs). IDRs were defined using Metapredict (score>0.5), and hydrophobic residues were counted as G, A, V, P, L, I, M, W, and F. Values are color-scaled independently per column, with the lower bound set to 0.8 × the minimum value.

Failure to bind robustly across the yeast genome was also observed in families that bind DNA as monomers. In these cases, failure to bind could reflect weak intrinsic affinity of the DBD to its motif, which in the native human context is stabilized by co-binding TFs. Alternatively, regions outside the DBDs could inhibit or enhance association to the yeast genome. To distinguish between these possibilities, we focused on the SOX DBD family, where differences in binding were pronounced: Of the ten SOX TFs tested, two showed strong promoter and motif association, two showed weaker binding, and six did not show reproducible binding (Fig. 1F, 2B). Of note, the two strong binders (SOX5,6) and one intermediate binder (SOX13) are of the SoxD sub-family, sharing highly conserved DBDs. Within their non-DBD, both SOX5 and SOX6 contain coiled-coil domains, which enable dimerization and may stabilize binding at motif pairs, although we saw little evidence for such preference in our data (Fig. S2A). Most binding events occurred at isolated motifs (distance >30 bp), with only a minor enrichment in proximal motif pairs (≤30 bp) relative to the background motif distribution (Fig. S2B).

We selected seven SOX TFs and generated their isolated DBD and non-DBD variants (Fig. 2C). Notably, in our ChEC-seq profiles, all DBDs showed similar weak binding, including the DBDs of the strong-binding TFs SOX5 and SOX6 (Fig. 2D). By contrast, binding strengths of the isolated non-DBD correlated with those of the full-length TFs, with non-DBDs of SOX5, SOX6, and SOX13 showing strong binding despite the lack of a DBD. What distinguishes these three non-DBDs relative to those of the four other SOX factors tested remains unclear, although we note that non-DBD length and its propensity to form alpha-helices, as well as total IDR hydrophobic content largely correlate with binding strength of the full TF (Fig. 2E). Together, these data suggest that the DBDs of SOX TFs are of too weak affinities to support consistent binding across the yeast genome alone, but are stabilized by their non-DBDs, which dictate binding strength for the full protein.

### Human TF binding varies across motif occurrences

In their native context, TFs bind to only a fraction of their motif sites^1,2,42,43^. We asked whether human TFs show similar site selectivity when tested in yeast lacking their human-specific interactors. First, we focused on ERG and ATF1, collecting all motif occurrences within promoters and ordering them by the respective ChEC-seq signal. Binding signals of both TFs varied widely across motif sites, with strong binding signals limited to only a subset of occurrences (Fig. 3A). Repeating the analysis for the native yeast bZIP TF Cst6, which binds a highly similar motif to the human bZIP factor ATF1, revealed a comparable selectivity bias.

**Figure 3:**
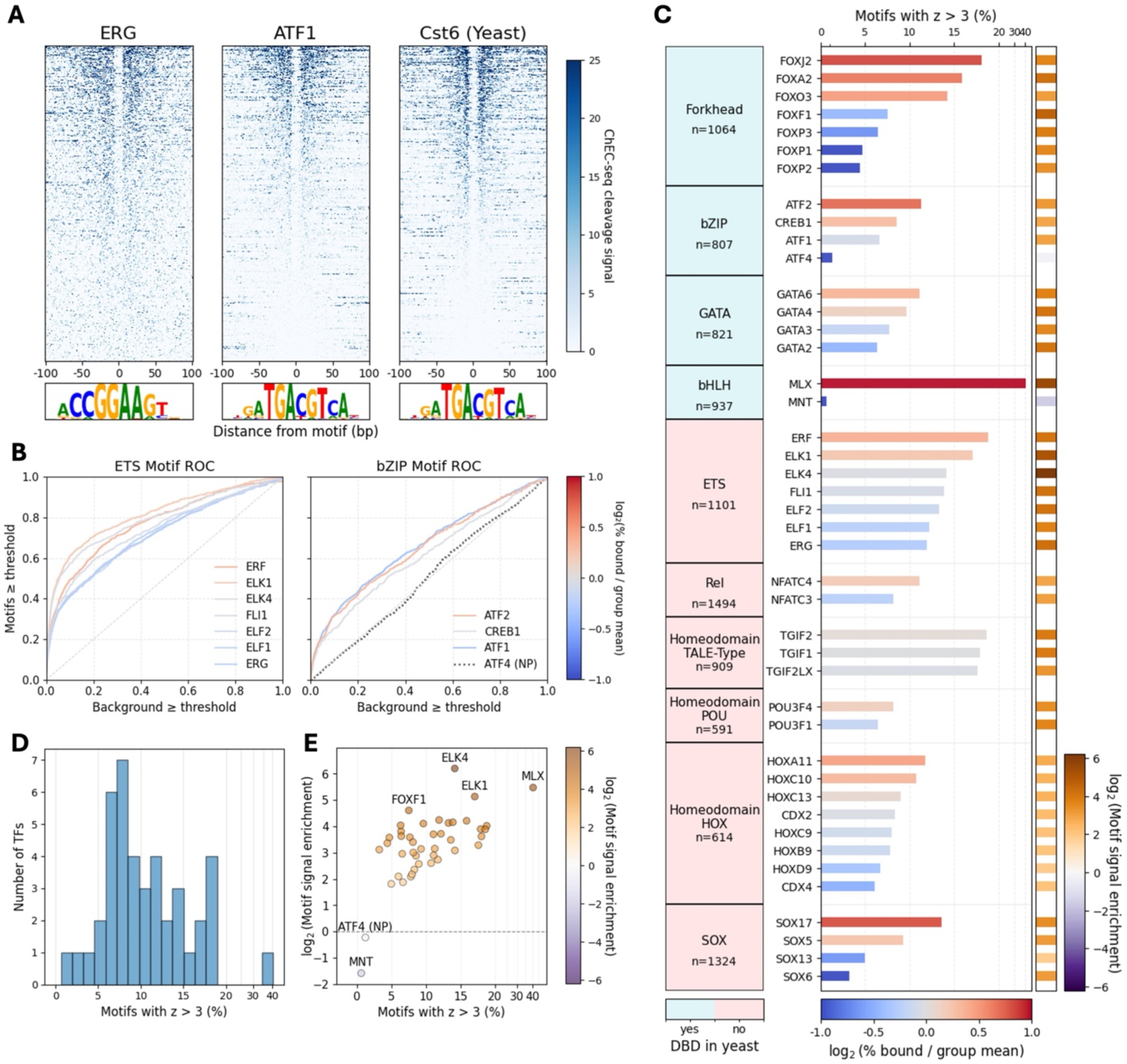
Human TF binding varies across motif occurrences. *(3A) Strong binding is restricted to a subset of motif sites:* Heatmaps of the ChEC-seq binding signal (±100 bp) around all occurrences of the family cognate motif for ERG (human ETS family), ATF1 (human bZIP family), and Cst6 (yeast bZIP family). Motif instances are ordered per TF by binding signal (see Methods). *(3B) Binding is enriched at motif sites, with family-specific variability:* ROC-like curves depict enrichment of binding at motif sites relative to background (see Methods). The y axis shows the fraction of motif instances exceeding a given z-score binding threshold, while the x axis shows the fraction of background windows exceeding the same threshold. Left, ETS family TFs; right, bZIP family TFs. ATF4, which could not be profiled (NP), is included as a dotted black line as a non-binding control. The dashed gray diagonal indicates the expected 1:1 relationship in the absence of motif-specific enrichment. Bars are colored by the log_2_-transformed fraction of bound motifs relative to the mean fraction bound across the entire corresponding motif family. *(3C) TFs recognizing the same motif differ in the fraction of motif sites they bind:* Bar plots show, for each TF, the fraction of motif occurrences with binding z-score >3 (see Methods). Bar colors correspond to the curve colors shown in *(B)*. The rightmost column shows a heatmap of motif binding signal enrichment, calculated as the log_2_ ratio between the fraction of motif sites and background sites exceeding z-score >3. *(3D) Most human TFs bind only a small fraction of their motif occurrences:* Histogram shows the number of TFs (y axis) as a function of the fraction of their cognate motifs bound with z-score >3 (x axis). *(3E) Motif enrichment is largely independent of the number of bound motifs*: Shown is motif-binding enrichment (calculated as in *(C)*) plotted against the percentage of motifs bound with z-score >3. Points are colored as in *(C)*. ATF4 (NP) is included as a control.

We summarized specific TF binding across motif sites using receiver operating characteristic (ROC)–like curves, plotting the fraction of motif sites above a given binding threshold against the fraction of background sites above the same threshold (Fig. 3B). These plots confirmed strong enrichment of motifs amongst the strongly bound sites but also emphasized that binding is limited to a subset of motif sites. Notably, using these ROC-like curves, we confirmed limited sensitivity to the stringency of motif site definition (FIMO^44^ thresholds) (Fig. S3A).

To quantify the fraction of bound sites across our full dataset, we measured the fraction of motifs bound with a z-score >3 (Fig. 3C). Notably, the percentage of bound motif sites varied among TFs within the same motif family. For example, in the forkhead family, FOXJ2 and FOXA2 bound greater than 15% of their motif occurrences, whereas FOXP1,2,3 localized to only 4-6% of their sites. The TF that bound the highest fraction of its motifs was MLX (41%), a bHLH TF. This relatively permissive binding was an exception, as most other human TFs bound at only 5-20% of their motif sites (Fig. 3D). For seven of the ten DBD families, the fraction of motifs bound differed by at least 1.5-fold between TFs. When comparing motif sites to background signal, clear enrichment was observed, generally increasing with the fraction of motifs bound (Fig. 3E). We conclude that human TFs preferentially bind to specific motif occurrences across the yeast genome at levels significantly higher than non-motif sites.

### Human TFs bind a higher fraction of motifs at low nucleosome coverage sites

Chromatin accessibility could explain the differential binding of human TFs across their motif sites^45^. In budding yeast, closed heterochromatin is limited to specific regions hence majority of promoters are located within open chromatin^46–48^. Still, promoters may differ in accessibility depending on their respective nucleosome arrangements. We asked whether preference for motifs in nucleosome-unoccupied DNA could explain the differential binding of human TFs across their motif sites. We reasoned that this could potentially explain differences in the fraction of bound sites between TFs of the same motif family, if TFs that bind a higher fraction of their sites are of pioneering-like activity, enabling their binding at nucleosome-covered motifs.

We visualized the nucleosome context of bound and unbound motifs using existing MNase-seq data^27^. In both ERG and ATF1, strongly bound motifs were located, on average, between nucleosome peaks whereas unbound motifs were more often nucleosome covered (Fig. 4A, S4A). To quantify the relationship between motif binding and nucleosome coverage, we separated motif sites into two groups based on their associated nucleosomes and compared the respective ROC-like curves of each group (Fig. 4B). We noted that the background signals at non-motif sites depend on nucleosome coverage (Fig. S4B), so we normalized the motif binding score accordingly. We expected the binding signal enrichment to differ substantially between high and low nucleosome motif classes, with bound motifs being enriched primarily in the low-nucleosome group. This was indeed the case for ERG, which strongly favored binding nucleosome-free motifs as compared to nucleosome-covered ones (Fig. 4B). However, in the case of ATF1, we noticed a much smaller difference in the area under the curve for high and low nucleosome motif occurrences, indicating a reduced relative preference for nucleosome-free motifs. This result motivated us to quantify nucleosome sensitivity across all human TFs in our dataset.

**Figure 4:**
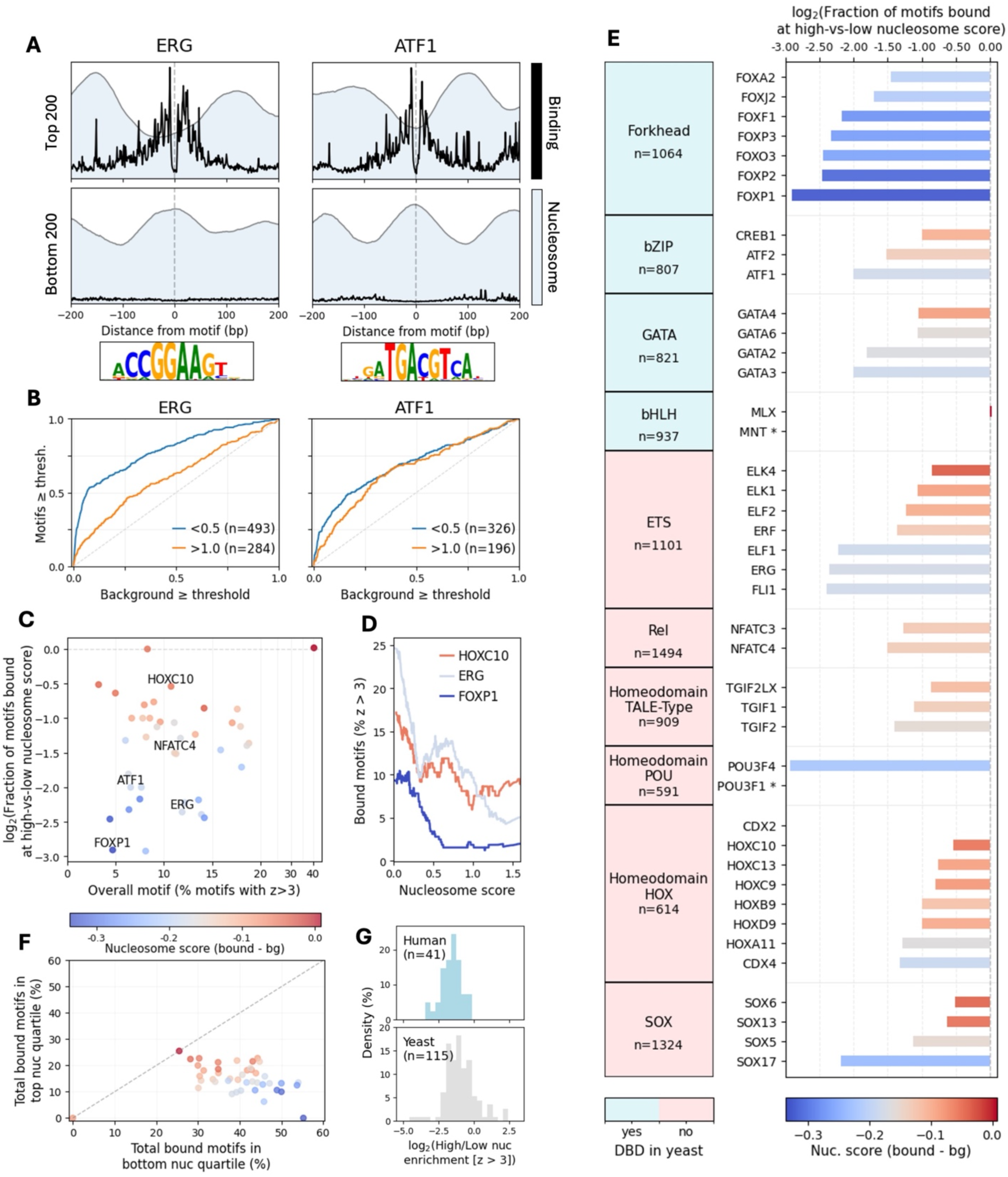
Human TFs bind a higher fraction of motifs at nucleosome-free sites. *(4A) Strongly bound sites are located between nucleosomes on average:* ChEC-seq signal footprints are shown for the strongest and weakest bound motif instances of ERG and ATF1 (see Methods). Mean nucleosome occupancy profiles derived from MNase-seq data^27^ are shown in the background for the same motif instances. Each TF’s motif logo is plotted below. *(4B) Binding is more strongly enriched at motifs located in low-nucleosome regions:* ROC-like curves compare enrichment of binding at motif sites relative to background for ERG and ATF1 (see Methods). Curves are shown separately for motifs classified as low- or high-nucleosome, based on the mean nucleosome score within ±7 bp of the motif center. Sample sizes for each nucleosome group are indicated in parentheses in the legend. *(4C-E) Nucleosome sensitivity varies across TFs and is independent of overall binding strength: (C)* Shown is a quantitative measure of nucleosome sensitivity defined as the ratio between the fraction of motifs bound at high nucleosome occupancy (95^th^ percentile of motif nucleosome scores) and the fraction bound at low nucleosome occupancy (5^th^ percentile). This ratio is plotted against the total fraction of motifs bound with z-score >3. *(D)* Fraction of bound motifs for a given nucleosome score is plotted for the nucleosome sensitive TFs FOXP1 and ERG, and nucleosome insensitive TF HOXC10 TFs. Binding fractions were calculated using sliding windows of ±10% of all motif occurrences ordered by nucleosome score. *(E)* Summary of high-versus-low nucleosome binding ratios across all TFs, with the total number of motifs reported under the DBD family name. Star next to TF name represents no binding at high nucleosome occupancy motifs. Markers in all three panels are colored by the mean nucleosome score at bound motifs compared to all motif sites. *(4F) Despite TF low-nucleosome preference, many bound motifs reside in nucleosome-occupied regions:* Scatterplot shows, for each TF, the fraction of all bound family representative HOCOMOCO motifs falling into the lowest quartile (x axis) and highest quartile (y axis) of motif nucleosome scores. Points are colored as in *(C–E)*. *(4G) Human and yeast TFs exhibit similar sensitivity to nucleosomes:* Histograms show the log_2_-transformed ratio between the fraction of bound individual HOCOMOCO motifs in the top versus bottom quartile of nucleosome scores. Distributions are shown separately for human and yeast TFs, with sample sizes indicated in parentheses.

For all human TFs, we quantified the fraction of motifs bound across the full spectrum of nucleosome occupancy (Fig. 4C,D). Nearly all TFs bound a higher fraction of motifs in low-nucleosome regions compared to nucleosome-covered regions (Fig. 4C,E), indicating a general preference for accessible DNA. Notably, this preference was independent of the overall fraction of bound motifs (Fig. 4C), arguing against a model in which differences in binding selectivity simply reflect an enhanced ability of some TFs to bind nucleosome-occupied DNA. Furthermore, for most TFs, a substantial fraction of bound motifs (∼10–22%) reside in nucleosome-covered regions (Fig. 4F). The high-to-low nucleosome binding ratios of human TFs is comparable to those of native yeast TFs (Fig. 4G). Together, these results indicate that while nucleosome occupancy strongly influences TF binding, it is not sufficient to explain selective motif occupancy across promoters.

Nucleosome-insensitive TFs were found across multiple families including HOX, ETS, and SOX. The bHLH TF MLX, which bound to the highest fraction of its motifs overall, showed total indifference to nucleosomes (Fig. 4E). We note, however, that only 16% of MLX motifs are classified here as high nucleosome-occupancy motifs, fewer than all other motifs, which range from 23-32% high nucleosome score motifs (Fig. S4C). Contrary to our expectations, members of the known pioneering family FOX showed more limited binding at nucleosome-covered motifs. While FOXA2, a canonical pioneering factor, showed the least nucleosome sensitivity within the FOX family, its nucleosome preference compared to all other TFs represents a relatively nucleosome-averse profile (Fig. 4E). Similarly, GATA family TFs did not exhibit enriched binding at high-nucleosome motifs compared with other, non-pioneering TF families (Fig. 4E). We conclude that nucleosome occupancy acts as a strong barrier to human TF binding but does not fully explain motif selectivity. The ability to bind nucleosome-covered motif sites in the yeast genome appears largely independent of canonical pioneering activity in human cells, potentially reflecting the generally accessible chromatin environment of yeast, or the absence of recruited pioneering partners present in the native context.

### Distinct genomic preferences of human TFs with a shared cognate motif

We initially expected that TFs of the same motif family would show the same binding preference across their motif sites, perhaps with different capacities for binding at nucleosome-covered motifs. However, this hypothesis was challenged by our results showing that almost all TFs could still bind high-nucleosome motifs, indicating a limited effect on motif selectivity. These findings motivated us to more directly compare the motif sites bound by members of the same DBD family. Indeed, examining binding of ATF1 and ATF2 across their common motif sites revealed common sites that were bound by both factors, as well as unique sites bound exclusively bound by either ATF1 or ATF2 (Fig. 5A-C). Similar differences in binding preference were seen when comparing binding across promoters, in which case the total ChEC-seq signal mapped to each promoter was compared (Fig. 5D).

**Figure 5:**
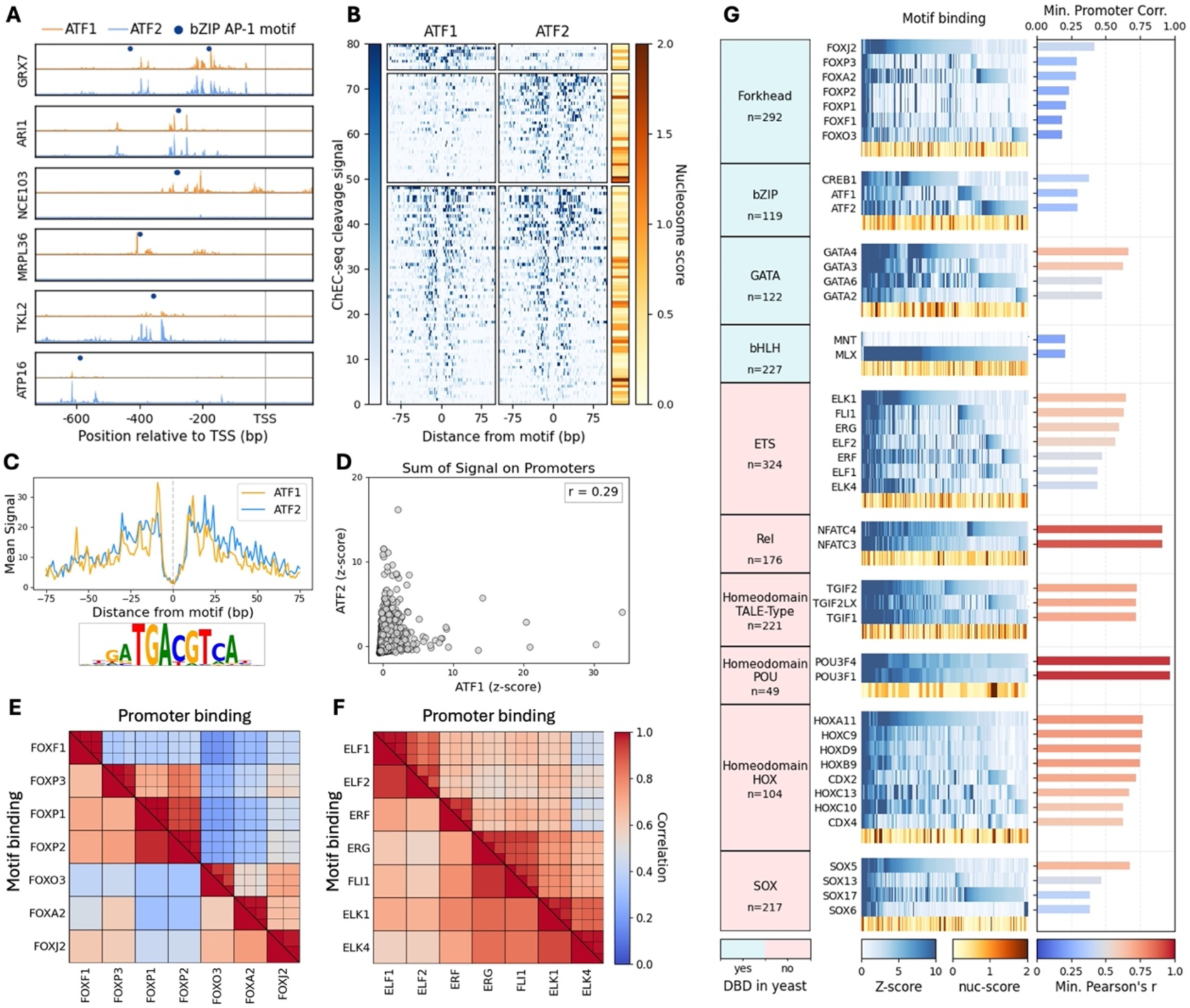
TFs binding the same cognate motif show distinct genomic preferences. *(5A-B) TFs from the same DBD family bind unique subsets of shared motif sites: (A)* ChEC-seq cleavage tracks for the bZIP TFs ATF1 and ATF2 at representative target promoters. Tracks represent MNase cleavage sites, and dots indicate occurrences of the representative bZIP AP-1 motif. *(B)* Heatmaps show the ChEC-seq binding signal (±100 bp around motif center) at motif occurrences classified as ATF1-unique (top; z-score >3 in ATF1 and <2 in ATF2; n=8, ATF2-unique (middle; z-score >3 in ATF2 and <2 in ATF1; n=35), or shared (bottom; z-score >1 in both TFs; n=73). Motif occurrences are ordered by binding strength, and the rightmost color bar indicates the nucleosome score associated with each motif site, calculated in a window of ±7 bp from motif center. *(5C) TFs from the same DBD family bind the same motif in yeast:* Mean ChEC-seq cleavage signal is shown around the top 200 bound motif instances for ATF1 and ATF2, aligned by motif center. The representative bZIP family motif logo is shown below the footprints. *(5D) TFs sharing a cognate motif bind distinct sets of promoters:* Scatterplot compares ATF1 and ATF2 promoter binding, where each point represents a promoter in the yeast genome and values correspond to the summed, z-transformed ChEC-seq signal across the promoter (see Methods). Pearson’s correlation coefficient (r) across all promoters is indicated. *(5E-F) Divergence in promoter and motif binding within DBD families:* Shown are Pearson correlation matrices for the forkhead *(E)* and ETS *(F)* DBD families. The upper right half of each matrix displays Pearson correlations computed between biological replicates based on promoter binding signals. The lower left half displays Pearson correlations of replicate-averaged binding scores across all motif sites. *(5G) Summary of motif- and promoter-level binding divergence across DBD families:* For each DBD family, the heatmap panel depicts binding at motif occurrences (columns), where motifs included are those bound by at least one TF in the family (bound motif sample size is reported per family). The nucleosome score associated with each motif site is shown as the bottom row of the heatmap. The bar plot panel reports, for each TF, the minimum Pearson correlation of promoter binding with any other TF in the same family.

We extended the analysis to compare the binding preferences of TFs from the same motif family by correlating binding signals across sites of their common motif and across promoters (5E-F). As expected, the correlation patterns between different TFs were largely consistent across these two metrics. For example, FOXP1 and FOXP2 bound similar promoters (r=0.93) and motif sites (r=0.96), while FOXP1 and FOXA2 showed distinct preferences for promoter (r=0.29) and motif (r=0.31) binding. Across human TFs from the same family, we observed remarkable differences in both promoter and motif site selectivity (Fig. 5G). The minimum promoter-binding correlation between each factor and all other members of its DBD family was less than 0.75 for 34/42 human TFs, and less than 0.50 for 20/42 factors. Additionally, within each family, an average of just 27% of bound motifs were shared across all family members. These differential preferences extended to most DBD families, except Rel and Homeodomain-POU, where only two TFs were successfully profiled. Collectively, our findings show that human TFs of the same motif family uniquely bind across their motif sites and promoters, indicating differential specificity independent of their cognate motif.

### Human TFs bind to OPN-type promoters enriched with yeast TFs

To better understand the differences in binding of TFs that share a common motif, we examined the promoters that are favorably bound by the tested human TFs. In yeast, native TFs strongly favor binding at the small subset of promoters displaying a fuzzy-nucleosome architecture^14,49^, termed occupied proximal nucleosome (OPN)^50^. We previously showed that TF preference for binding its motif at high-OPN promoters depends on the disordered non-DBD, as DBD-only mutants bind more uniformly across promoter types^14^. To examine whether human TFs show a similar bias for high-OPN promoters, we ordered promoters by their OPN score and plotted the binding signal of each TF at all promoters (Fig. 6A). Human TFs did show a bias for binding at high-OPN promoters, which was slightly lower on average than that of native yeast TFs, but higher than that of the yeast DBD mutants (Fig. S6A).

**Figure 6:**
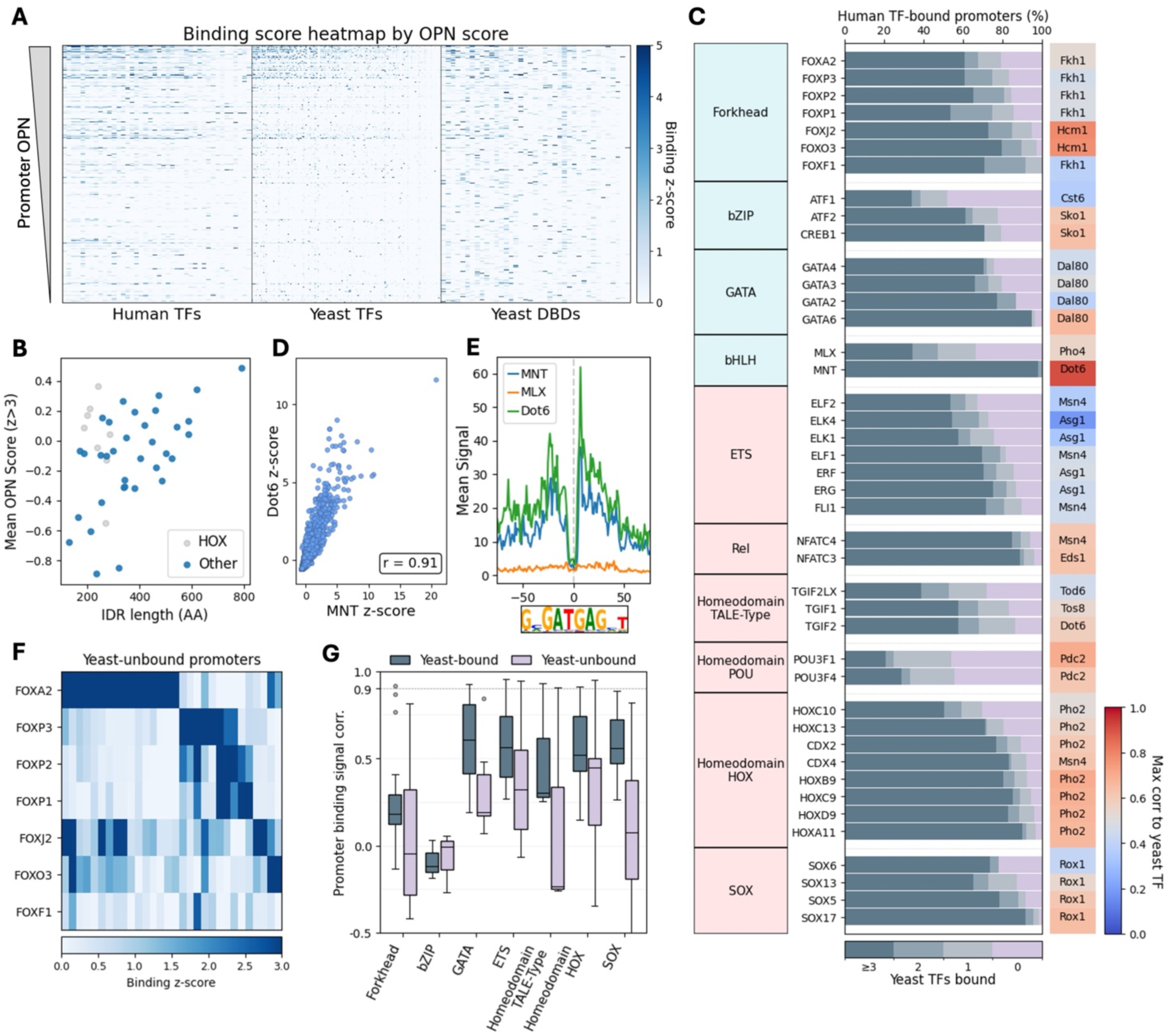
Human TFs bind promoters with fuzzy nucleosome architecture independently of yeast TFs. *(6A) Human and yeast TFs preferentially bind occupied proximal nucleosome (OPN) promoters:* Heatmap shows the summed, z-transformed promoter binding signal of human TFs (n=42), yeast TFs (n=147), and yeast DBD-only strains (n=39) across all budding yeast promoters, ordered by OPN score. Columns correspond to TFs and rows correspond to promoters. *(6B) TFs with longer intrinsically disordered regions (IDRs) bind more OPN-type promoters:* For each TF, mean OPN score of bound promoters (z-score>3) is plotted against IDR length (amino acids). IDRs were defined using Metapredict with a disorder score >0.5. *(6C) Human TFs bind promoters that are also targeted by yeast TFs:* Stacked bar plot shows, for each human TF, the fraction of bound promoters that are bound by 0, 1, 2, or ≥3 yeast TFs. The heatmap bar annotates, for each human TF, the yeast TF with the highest Pearson correlation across summed, z-transformed promoter binding scores, colored by the corresponding correlation value. *(6D-E) MNT colocalizes with the yeast TF Dot6 at shared target promoters:* Shown in *(D)* is a scatterplot comparing promoter binding of human MNT and yeast Dot6, where each point represents a promoter and values correspond to summed, z-transformed binding signal. Pearson’s correlation coefficient (r) is indicated. In *(E)*, mean footprints are shown for Dot6 (yeast Myb TF), and MNT and MLX (human bHLH TFs) at occurrences of the Dot6 PAC motif, aligned by motif center for the top 200 sites ranked by the number of cleavage events within ±25 bp of the motif center. *(6F) Novel promoter targets are not shared among TFs of the same family:* Heatmap shows promoter binding for forkhead family TFs at promoters unbound by any yeast TF, highlighting novel target promoters. Columns correspond to promoters. *(6G) Promoter binding correlation is further reduced at yeast-unbound promoters:* Boxplots summarize Pearson correlations of promoter binding between TFs within the same DNA-binding domain family, calculated separately for promoters bound by at least three yeast TFs (gray) and for promoters unbound by any yeast TF (purple). Only DBD families with at least three TFs were included.

Furthermore, this preference for high OPN promoters differed between human TFs in a manner that correlated with the lengths of their IDRs (Fig. 6B). Notably, the HOX DBD family deviated from this trend (Spearman’s *r*=0.30 for all TFs including HOX, Spearman’s *r*=0.53 excluding HOX) by binding high-OPN-type promoters despite relatively short IDRs, remaining an outlier among the tested TFs.

We observed a clear concordance of promoter binding targets between human and native yeast TFs. To measure this more directly, we calculated the number of yeast TFs bound to the target promoters of each of the human TFs in our dataset (Fig. 6C). Of the 5358 promoters analyzed in budding yeast, only 14% are bound by three or more of the 147 yeast TFs profiled under identical experimental conditions (Fig. S6B). In comparison, an average of 67% of human TF-bound promoters are bound by at least three yeast TFs. Overall, promoters bound by human TFs are significantly enriched with yeast TFs (P<0.05) for 41/42 profiled human TFs (Fig. S6B). We conclude that human TFs bind preferentially to high OPN promoters favored by their yeast counterparts, and this OPN preference correlates with the lengths of their IDRs.

### Human TFs bind independently of yeast TFs

The preference of human TFs for promoters enriched with yeast TFs raises the possibility that human TF binding is guided by emerging interactions with native yeast TFs. We reasoned that such interactions could explain differences in binding preferences between TFs of the same motif family if they undergo interactions with unique yeast TFs. To test this, we searched for human TFs whose binding profiles correlate well with those of individual yeast TFs tested under the same experimental conditions. Only one such case emerged: MNT, a human bHLH TF, localized to practically the same promoters as Dot6 (r=0.91), a yeast repressor of the Myb family (Fig. 6D). MNT’s profile was unique in our data, as it was the only reproducible human TF that showed no apparent preference for its cognate motif. It did, however, localize with a clear footprint to the Dot6 cognate motif (Fig. 6E). MLX, the other human bHLH TF profiled, showed no such interaction signatures. Together, our results suggest a single unique case of opportunistic interaction between MNT and Dot6, causing a convergence of target promoters.

Preferences of all other human TFs tested were distinct from those of individual yeast TFs. For motif families absent from yeast, such as ETS and POU, maximal correlations in promoter signal reached 18-70%. For motif families that do exist in yeast, such as bZIP and forkhead, maximum correlations ranged from 37-79%, but the respective yeast TFs were always of the same motif family, indicating a limited motif-based bias. Furthermore, these limited similarities do not explain the differences in binding preferences across same-family TFs, as differences between human and yeast TFs of the same motif family were larger than between the human TFs themselves (Fig. S6C-E). Therefore, binding of the human TFs across the yeast genome appears largely independent of interactions with yeast TFs.

Further supporting yeast TF-independent binding of human TFs, we observed a substantial fraction of promoter targets unbound by any yeast TFs (Fig. 6C). We explored the identity of these new sites to determine whether they are shared amongst TFs from the same family. For most TFs, we see a clear split of novel targets both in identity and quantity (Fig. 6F). Correlation in the binding signal of human TF-bound promoters that are unbound by any yeast TFs is, on average, even lower than the correlation of yeast-bound promoters (Fig. 6G). These results point to TF-specific binding preferences that are independent of native yeast TFs, consistent with the model of intrinsic sequence-based logic encoded in human TFs.

### Swapping the non-DBD shifts binding preferences of human TFs across the yeast genome

Having largely ruled out the formation of cross-species dimers, we hypothesized that the differential binding patterns between TFs from the same-motif family may arise from intrinsic DBD-based preferences. While consensus motifs derived from our data were consistent across motif families, differences amongst TFs emerged when more directly comparing binding signals across all 7-mers, potentially indicating preferences for specific bases flanking the consensus motif, or simply reflecting the identity of selected promoters. Supporting DBD-specific preferences, we saw that sequence similarities amongst DBDs from the same motif family were often consistent with similarities in binding profiles, while this trend was weaker for non-DBDs (Fig. 7A, S7A-C). Together, our results initially pointed to a DBD-driven TF binding pattern.

**Figure 7:**
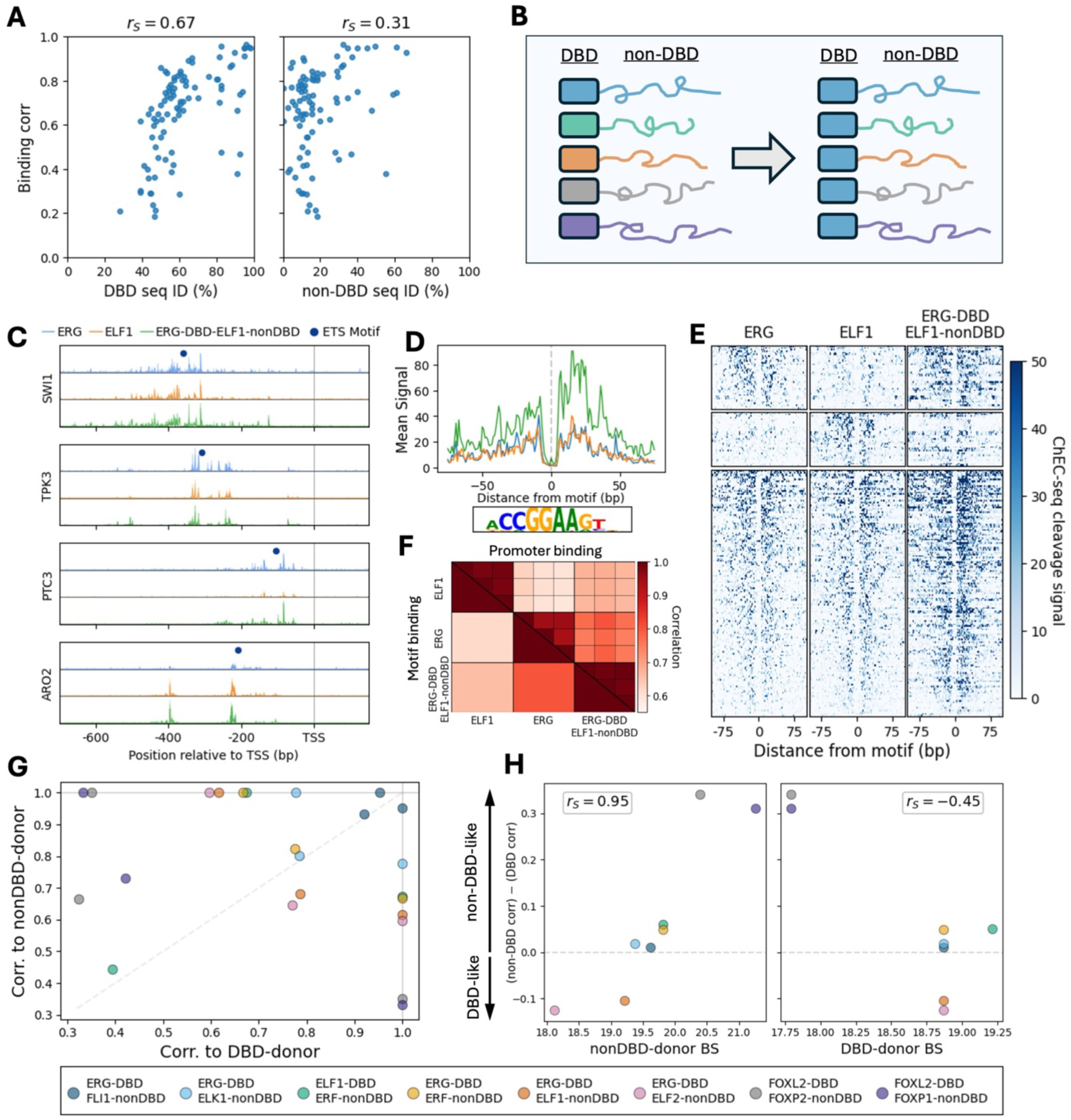
The largely disordered non-DBD plays a vital role in determining binding preferences. *(7A) DBD sequence similarity better predicts promoter binding similarity than non-DBD sequence similarity:* Scatterplots compare amino acid sequence alignment score of DBDs (left) or non-DBDs (right) with TF promoter binding correlation (see Methods). Spearman’s rank correlation coefficient (rₛ) between sequence similarity and binding similarity is indicated. *(7B) Experimental design for DBD swap strains:* Schematic illustrating the construction of DBD swap TFs, in which the DBDs of ERG, ELF1, and FOXL2 were exchanged into other members of their respective motif families to generate chimeric proteins. *(7C) DBD swap strains bind promoters targeted by both donor TFs:* Shown are ChEC-seq cleavage tracks for ERG, ELF1, and the ERG-DBD–ELF1 swap strain at representative target promoters. Tracks represent sequencing read cut sites, and dots indicate occurrences of the ETS family motif. *(7D) Swap strains bind the same cognate motif as their donor TFs:* Mean ChEC-seq signal is shown around the top 200 bound motif instances for ERG, ELF1, and ERG-DBD–ELF1, aligned by motif center. The representative ETS family motif logo is shown below the footprints. *(7E) Swap strains bind motif subsets associated with both the DBD and non-DBD donors:* Heatmaps show the ChEC-seq binding signal at motif occurrences (±100 bp) classified as ERG-unique (top; z-score >3 in ERG and <2 in ELF1; n=41), ELF1-unique (middle; z-score >3 in ELF1 and <2 in ERG; n=36), or shared (bottom; z-score >1 in both TFs; n=159). The binding of the ERG-DBD–ELF1 swap strain at the same motif sites is shown in the third column. *(7F) DBD swap strains exhibit intermediate promoter- and motif-level binding profiles:* Shown is a correlation matrix comparing ERG, ELF1, and ERG-DBD–ELF1. The upper right half displays Pearson correlations computed between biological replicates based on promoter binding signals, while the lower left half shows Pearson correlations of replicate-averaged binding scores across all motif sites. *(7G-H) non-DBD binding strength shapes swap strain binding profiles: (G)* Promoter binding similarity of each swap strain to its DBD (x axis) and non-DBD (y axis) donor is shown. The correlation of each donor to the other is shown along the x=1 and y=1 axes. *(H)* The difference between non-DBD and DBD promoter binding correlations to the swap strain is plotted as a function of non-DBD donor binding strength (left) or DBD donor binding strength (right) (see Methods). Spearman’s rank correlation coefficient (rₛ) is reported.

To experimentally test whether the DBDs can explain the differences in binding preferences between TFs of same-motif families, we considered the ETS and forkhead families, where TFs sharing the same cognate motif showed substantial differences in binding preferences (Fig. 5E-F). We generated 18 DBD-swapped yeast strains combining the DBDs of ERG, ELF1, or FOXL2 with non-DBD regions from other family members (Fig. 7B). Robust ChEC-seq profiles were obtained for 5/6 ERG DBD-swapped factors (Fig. S7D), with the relatively weak binding TF ELK4 failing to bind reproducibly when donating its non-DBD. In contrast, only 1/6 ELF1 DBD strains and 2/6 FOXL2 DBD strains were successfully profiled (Fig. S7D). The non-DBDs of the sole reproducible ELF1 and FOXL2 DBD strains came from the strongest binding wildtype TFs in each family, suggesting a joint DBD and non-DBD effect on binding success.

Consistent with their sequence donors, reproducible swap strains localized to specific promoters containing their cognate motif (Fig. 7C-D). We next compared the binding profile of the swaps to their DBD donor or their non-DBD donor. Analysis of motif selectivity revealed swap strain binding at both shared and donor-specific sites (Fig. 7E). Generally, an intermediate profile was produced, resulting in a higher correlation to each of the donors than between the donor strains themselves (Fig. 7F-G).

Finally, we examined how binding strength influences swap strain profiles across the eight successful swap strains from the ETS and forkhead families. Differences between correlations to the non-DBD donor versus the DBD donor were explained by the binding strength of the non-DBD donor (Fig. 7H). Specifically, non-DBDs derived from strongly binding TFs were more likely to shift swap strain profiles toward those of the non-DBD donor. In contrast, no clear relationship was observed with DBD donor binding strength. We conclude that TF binding profiles are shaped by both the DBD and non-DBD, but that the non-DBD is the primary determinant of binding behavior in strongly binding TFs.

## Discussion

Understanding how transcription factors (TFs), which recognize the same DNA motif, bind at distinct genomic sites and regulate different gene sets remains a central challenge in the field of gene regulation. The prevailing view is that motif-sharing TFs are recruited to different sites through cooperative interactions with co-binding partners. Our study demonstrates that motif-sharing human TFs consistently bind at unique genomic locations when tested across the evolutionarily distant yeast genome, which lacks human-specific cofactors. These findings reveal that intrinsic TF properties direct genome-binding preferences beyond simple cognate motif recognition and independent of evolved interactions with co-binding proteins.

Our data support the use of yeast as a system for studying intrinsic genomic binding preferences of human TFs. Of the 60 human TFs we tested, 42 (70%) bound the yeast genome robustly enough to be reproducibly captured by ChEC-seq. Robust binding was observed in all ten tested DBD families, regardless of whether homologous DBDs exist in yeast. This high success rate was unexpected, given the absence of human-specific cofactors in yeast that stabilize TF binding in their native cellular context.

Still, we noted that human TFs displayed weaker binding signals compared to their yeast counterparts, which may be explained by the lack of their evolved stabilizing factors in the yeast system.

The human TFs tested bound preferentially at their cognate motifs, as anticipated. Notably, however, binding signals varied substantially across motif sites spread across promoters, with only a limited fraction (5–20%) being strongly bound. Moreover, both the fraction and the identity of bound motif sites differed between TFs within the same motif family. In prevailing models, such differences in motif site selection are attributed to chromatin accessibility or recruitment through co-binding factors. Because yeast lacks evolved human-specific interactors, we initially hypothesized that differences in motif site selection result from differences in DNA accessibility and the capacity of human TFs to bind nucleosome-covered sites. We did indeed observe strong preferential binding at nucleosome-free motifs, but these effects were insufficient to explain both the spread of binding intensities across motif sites and the differences in motif site selection between motif-sharing TFs. We also noted that TFs capable of binding nucleosome-covered motifs neither localized to a higher fraction of their motif sites nor corresponded to known pioneering TFs. This discrepancy may reflect the generally accessible chromatin landscape of yeast promoters, which lack major repressive features of the human genome such as Polycomb- and HP-associated repression, or may point to the cofactor-dependent nature of pioneering activity in human cells.

Specificity of human TFs may also arise from *de novo* interactions with yeast TFs. We observed one apparent case of such emerging recruitment: the human TF MNT localized to a non-cognate PAC motif and co-occupied precisely the same genomic sites as the PAC-binding yeast TF Dot6. However, this example was unique, as all 41 other human TFs tested localized to their cognate motifs and showed low-moderate correlation with native yeast TFs. Moreover, although human TFs generally favored promoters enriched for yeast TF binding, they bound a substantial fraction of novel promoters lacking binding by any yeast TFs. Together, these results support a model in which differential human TF binding at cognate motif sites occurs independently of yeast TF recruitment.

Human TF specificity may also reflect DBD-specific preferences within same-motif families that favor, for example, different motif-flanking sequences. However, testing this hypothesis with domain-swapping experiments revealed that the DBD alone was insufficient to explain specificity. In ETS and forkhead TF families, binding preferences were driven by both the DBD and the non-DBD. However, swap strains containing a stronger-binding non-DBD donor produced a more non-DBD-like binding profile, whereas swap strains with weak non-DBD donors consistently failed to profile. In the SOX family, TF binding strength was conveyed by the non-DBD, whereas the DBD-only strains struggled to bind across all tested TFs. Together, these results indicate that while DBDs define motif recognition, non-DBD regions play a dominant role in shaping genome-wide binding preferences and binding strengths.

Notably, the non-DBD regions that dominate binding specificity in our experiments are composed largely of intrinsically disordered regions (IDRs). This extends our previous findings implicating TF IDRs in directing genome recognition, suggesting that IDR-localized specificity determinants remain functional across evolutionarily distant genomes lacking co-evolved interaction partners. Mechanistically, our results argue against nucleosome-mediated accessibility and emergent protein-protein recruitment as primary explanations for IDR-directed binding in this cross-species system. Whether IDRs engage in direct DNA recognition that complements the specificity encoded by canonical DNA-binding domains remains an intriguing possibility and calls for further investigation into the biophysical principles underlying IDR-mediated TF-DNA interactions.

## Materials and Methods

### Experimental design of strains

Yeast parent strain was generated by inserting a synthesized gBlock (IDT) fragment into the HO locus of *S. cerevisiae* BY4741 strain (genotype: MATa his3Δ1 leu2Δ0 met15Δ0 ura3Δ0 genotype) using CRISPR. All TFs were transformed into the gBlock to produce the following continuous structure: VMA5 promoter, yellow fluorescent protein (YFP), SV40 NLS, MNase, linker with FLAG-tag, TF, ADH1 terminator. TF sequences were sourced from the ORFeome^51^ or an arrayed collection of plasmids containing a TF and activation domain fusion (pDEST-AD)^34^.

### Budding yeast genetic manipulation

Genetic transformations were performed using the LiAc/SS DNA/PEG method^52^. Transformation success was confirmed with PCR and Oxford Nanopore Technologies sequencing (Plasmidsaurus). Subsequently, the pbRA89 plasmid (addgene #100950) was lost from transformed cells by growing them in liquid YPD (yeast extract peptone dextrose), followed by plate-based selection for colonies without bRA89-encoded Hygromycin resistance. For DBD-swap strains and IDR-only strains, ligation of the gene-specific guide-RNA into the bRA89 plasmid was performed as previously described^53^.

### Pre-experiment cell growth

Yeast strains were grown on YPD plates. Single colonies were picked and incubated at 30°C in liquid SD (synthetic complete with dextrose) medium overnight, reaching stationary phase OD_600_ ≈10, then diluted again into fresh SD medium for the experiment.

### Fluorescent microscopy imaging of yeast

Stationary phase cultures were diluted into 500 μL of low-fluorescence synthetic complete medium (Formedium LoFlo, riboflavin- and folate-free with adjusted inositol) and grown overnight to OD_600_ = 0.3. Cells were immobilized in ConA-coated Ibidi μ-Slide 18-well chambers (0.25 mg/mL ConA, 15 min), air-dried for 30 min, seeded with 100 μL cell suspension, allowed to settle for 15 min, and washed once with fresh medium to remove non-adherent cells. Imaging was performed on an Andor Dragonfly spinning-disk confocal microscope (Fusion v2.4.0.14; Andor Zyla4 sCMOS camera) using a 100×/1.47 NA oil-immersion objective with hardware drift stabilization. YFP fluorescence was acquired in CF40 confocal mode with 514 nm excitation (200 ms exposure) and matched brightfield images were acquired with 50 ms exposure.

### ChEC-seq experiments

The experiments were performed as described previously^38^, with some modifications. Stationary phase cultures were diluted into 5 mL fresh SD media and grown overnight to OD_600_ = 4. Cells were pelleted (1,500g, 2 min), resuspended in 0.5 mL Buffer A (15 mM Tris pH 7.5, 80 mM KCl, 0.1 mM EGTA, 0.2 mM spermine, 0.5 mM spermidine, 1× cOmplete EDTA-free protease inhibitors [Roche; one tablet per 50 mL], 1 mM PMSF), transferred to 2 mL 96-well plates (Thermo Scientific), and washed twice with 1 mL Buffer A. Cells were resuspended in 150 µL Buffer A + 0.1% digitonin, transferred to an Eppendorf 96-well plate, and permeabilized at 30°C for 5 min. CaCl₂ was added to 2 mM and incubated for 30 s to activate MNase, then stopped with an equal volume of stop buffer (400 mM NaCl, 20 mM EDTA, 4 mM EGTA, 1% SDS). Samples were treated with Proteinase K (0.5 mg/mL, 55°C, 30 min), extracted with equal volume phenol-chloroform pH 8 (Sigma-Aldrich), vortexed, and centrifuged (17,000g, 15 min).

DNA was precipitated with 3 volumes cold 96% EtOH, 45 mg Glycoblue, and 20 mM sodium acetate (–80°C, >1 hr), pelleted (17,000g, 4°C, 10 min), washed with 70% EtOH, dried, and resuspended in 30 µL RNase A solution (0.33 mg/mL in TE: 10 mM Tris, 1 mM EDTA) for 20 min at 37°C. Small fragments were enriched by SPRI cleanup (Ampure XP): 0.8× (24 µL) beads for reverse cleanup (5 min, RT), supernatant retained; then 1× (30 µL) beads plus 5.4× (162 µL) isopropanol (5 min, RT). Beads were washed twice with 85% EtOH and DNA eluted in 30 µL 0.1× TE.

### Chec-Seq next-generation sequencing library preparation

Library preparation was performed as described^54^, with modifications. Following RNase treatment and reverse SPRI cleanup, DNA fragments were used for end-repair and A-tailing (ERA). A 5.4 µL ERA mix (1× T4 DNA ligase buffer [NEB], 0.5 mM dNTPs, 0.25 mM ATP, 2.75% PEG 4000, 6 U T4 PNK [NEB], 0.5 U T4 DNA Polymerase [Thermo Scientific], 0.5 U Taq DNA polymerase [Bioline]) was added to 14.6 µL sample and incubated at 12°C for 20 min, 37°C for 15 min, and 58°C for 45 min. ERA products were subjected to reverse SPRI cleanup with 0.5× (10 µL) SPRI beads (Ampure XP, Beckman Coulter); supernatant was retained, and small fragments were purified with 1.3× (26 µL) SPRI beads plus 5.4× (108 µL) isopropanol. After two 85% EtOH washes, DNA was eluted in 17 µL 0.1× TE. For ligation, 16.4 µL eluate was added to a 40 µL reaction (1× Quick ligase buffer [NEB], 4000 U Quick ligase [NEB], 6.4 nM Y-shaped barcode adaptors with T-overhang) and incubated 15 min at 20°C. Ligation products were cleaned by double SPRI: first 1.2× (48 µL) SPRI, eluted in 30 µL 0.1× TE; then without bead separation, 1.3× (39 µL) HXN buffer (2.5 M NaCl, 20% PEG 8000) was added and DNA eluted in 24 µL 0.1× TE. Next, 23 µL eluate was amplified in a 50 µL PCR (1× Kappa HIFI [Roche], 0.32 µM barcoded forward and reverse primers) with 45 s at 98°C; 16 cycles of 15 s at 98°C and 15 s at 60°C; and 1 min at 72°C.

Final libraries were cleaned with 1.1× (55 µL) SPRI and eluted in 15 µL 0.1× TE. Concentration and size distribution were assessed by Qbit (Thermo Scientific) and TapeStation (Agilent). Libraries were pooled equimolarly, diluted to 2 nM, and sequenced on NovaSeq X (Illumina) with the following parameters: Read1, 61 nt; Index1, 8 nt; Index2, 8 nt; Read2, 61 nt.

### General computational analysis

NGS data was processed using Snakemake. Python (version 3.13.1) was used with Jupyter Notebooks for data analysis. Pandas and NumPy libraries were used for data structuring and calculations, while Matplotlib and Seaborn libraries were utilized for data visualization. SciPy was used for statistical analysis. Additional genomic data handling and manupilation was performed using the Biopython library. Finally, original python scripts were written to handle all data.

### ChEC-seq NGS data processing

After demultiplexing using bcl2fastq (Illumina), reads were filtered using cutadapt with the following parameters: ‘--pair-filter=any -m 15 --action=trim’. Alignment to the S. cerevisiae genome R64-1-1 was done using Bowtie2 (‘--very-sensitive --trim-to 30 --dovetail’), then genomeCoverageBed (‘-5 -fs 1 -d’) was used to achieve single bp resolution of MNase cut sites. The ribosomal locus (chr12: 450,000-490,001), CUP1 and CUP2 sites (chr8: 212,408-215,118), and two positions within GAL11 (chr15: 234,421; 234,548) were filtered out, and the remaining reads were normalized to 10e6.

### Promoter definition

Promoter regions were defined as in previous studies^17^, with a minimum length of 700 bp upstream of the transcription start site. The total ChEC-seq signal across the entire promoter was summed. Promoter binding scores were calculated by converting the raw summed signal into a z-score. Target promoters were defined as z-score >3. To measure promoter binding profile correlation between strains, the Pearson correlation coefficient is calculated across all promoter z-scores.

### TF binding strength

TF binding strength was calculated as the sum of the raw ChEC-seq score across all target promoters (z-score >3). This score was calculated once per human TF using the replicate-averaged promoter binding scores, unless explicitly stated that the binding score was calculated individually per replicate.

### Motif binding analysis

Human canonical motifs per TF and representative motifs per DBD family were defined by HOCOMOCO v12^40^. Yeast TF motifs were extracted from CISBP^29^, prioritizing high quality PBM and SELEX data. Motif sites within promoter regions were defined using MEME Suite’s Find Individual Motif Occurrences (FIMO)^44^ algorithm with yeast promoter nucleotide composition as background (A, T: 31%; G, C: 19%). Motif signal was defined as the average ChEC-seq signal ±25 bp from the motif center position (total window = 51 bp). The nucleosome score was defined using existing MNase-seq data^27^ and was calculated as the mean signal ±7 bp from the motif center (total window = 15 bp). Background motif windows and their associated nucleosome scores were calculated across all promoter bases (n=4,363,940). Motif scores were converted into z-scores using an array of background motif scores from the most similar 10% of nucleosome scores.

### 7-mer analysis

The average ChEC-seq signal at all 8192 reverse compliment-aware 7-mers was calculated and converted to a z-score. To create a refined 7-mer motif based on this experimental data, a novel position weight matrix was created using canonical motif-like 7mers. The HOCOMOCO^40^ canonical motif core was defined by identifying the first and last positions with information content (IC) ≥ 1.2 bits and designating the entire contiguous region between them as the core. If this region was shorter than five nucleotides, it was extended iteratively toward the adjacent position with the higher IC value until a minimum length of five bases was reached.

7-mers were classified as core-matching if they overlapped the canonical core with at least 4 of 5 matching bases in any 5-bp window, considering both orientations. For each sample, the top 10 highest Z-scoring core-matching 7-mers were aligned to the top-scoring sequence, and a position weight matrix was constructed by weighing each aligned nucleotide by its z-score and normalizing each position to sum to one.

### Footprint analysis

Footprint scores were calculated as the mean ChEC-seq cleavage signal per nucleotide across the top 200 bound motif instances (ranked by binding score, as defined above). Motif instances were aligned by their center and oriented according to strand prior to averaging.

### Receiver Operating Characteristic (ROC)-like curves

Motif binding scores were compared to background genomic windows as described above. To generate ROC-like curves, 1,000 z-score thresholds spanning the full range of observed values (from highest to lowest) were sampled. For each threshold, we calculated the fraction of motif sites with binding scores exceeding the threshold (y axis) and the corresponding fraction of background windows exceeding the same threshold (x axis). A dashed diagonal line (x = y) indicates the expected distribution in the absence of enrichment at motif sites.

### Sequence section definitions and dimerization status

DNA-binding domains (DBDs) were annotated using InterProScan^55^. All remaining residues outside the annotated DBD were classified as the non-DBD region. Sequence disorder was predicted using Metapredict^56^, and IDRs were defined as non-DBD residues with a disorder score >0.5. Dimerization status (homodimer, heterodimer, or no dimer) was retrieved from UniProtKB (release 2025_01)^57^.

## Data Availability

All sequencing data is available at NCBI’s Gene Expression Omnibus (GEO) database repository under the study ID GSE325768. Computational analysis code can be accessed at the following link: https://doi.org/10.5281/zenodo.19368402.

## Acknowledgements

This work is funded by Deutsche Forschungsgemeinschaft (DFG) – Project number: 550609939, European Research Council (ERC), Israel Science Foundation (ISF), and Minerva Foundation (NB) and by the National Institutes of Health (NIH) – Award number: R35GM128625.

We thank Dr. Gilad Yaakov for his expertise and assistance throughout the yeast strain creation process and ChEC-seq experiments. We also wish to thank Dr. Tamar Jana Lang for her help in acquiring and processing the pDEST-AD library, and Joseph Steinberger for his guidance with the microscopy imaging. We thank the whole Barkai laboratory for providing a supportive environment to everyone, including exciting scientific discussions and insightful comments that improved the manuscript. Lastly, we would like to thank our families. Thank you to Rachel, Tanyss, and Sam Bugis for your never-ending support.

## Supplementary Figures

**Figure S1:**
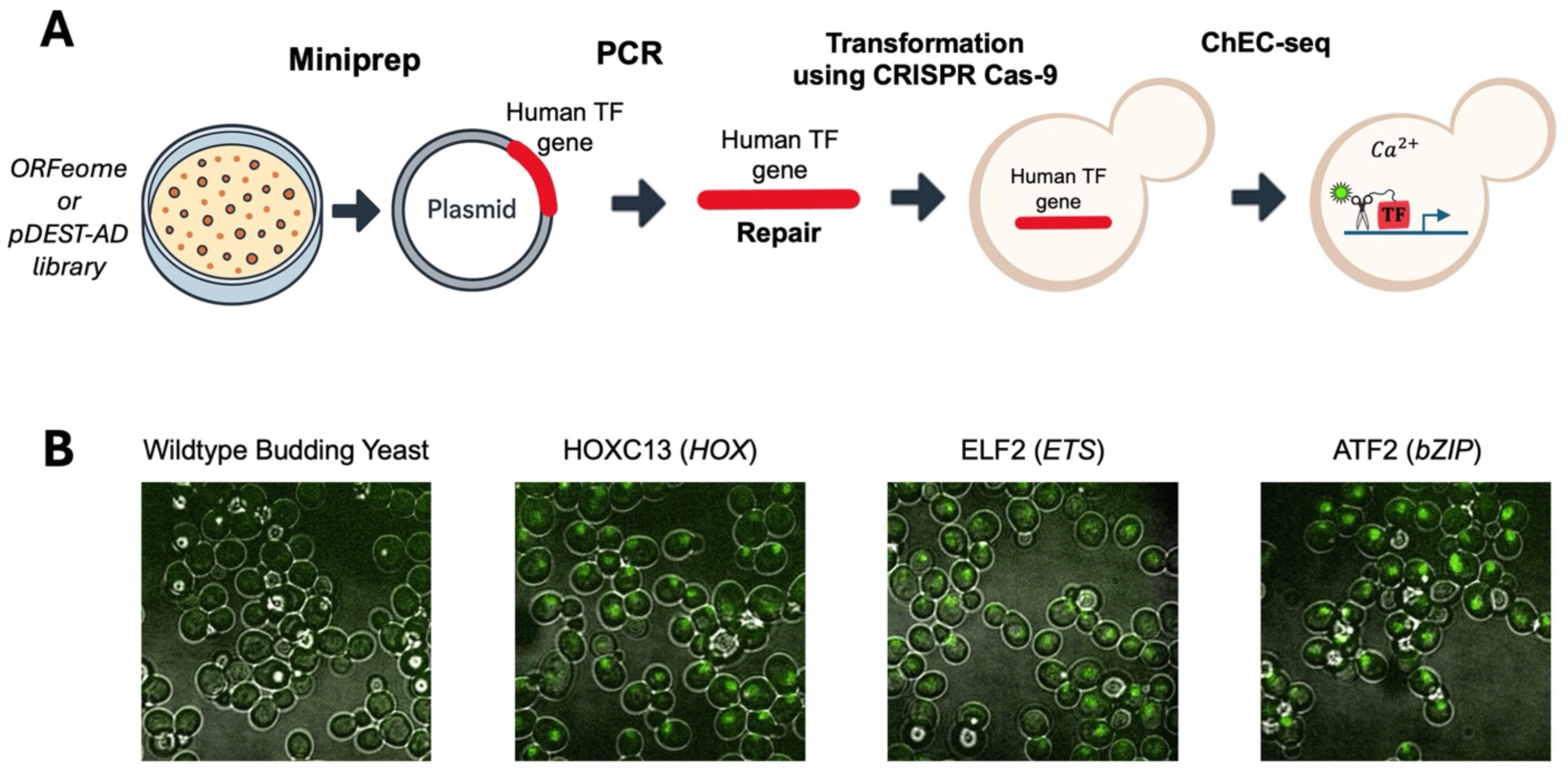
Overview of strain generation and ChEC-seq workflow. *(S1A) Schematic of yeast strain construction and ChEC-seq experimental workflow:* Diagram illustrates the genetic manipulation strategy used to generate yeast strains expressing MNase-tagged transcription factors (see Methods). Scissors and green star represent translated MNase and yellow fluorescent protein (YFP), respectively. *(S1B) Human transcription factors are expressed in the budding yeast system:* Fluorescent microscopy images (see Methods) of untransformed wildtype budding yeast, and three human TF-transformed budding yeast strains expressing YFP, showing nuclear-localized expression the human proteins.

**Figure S2:**
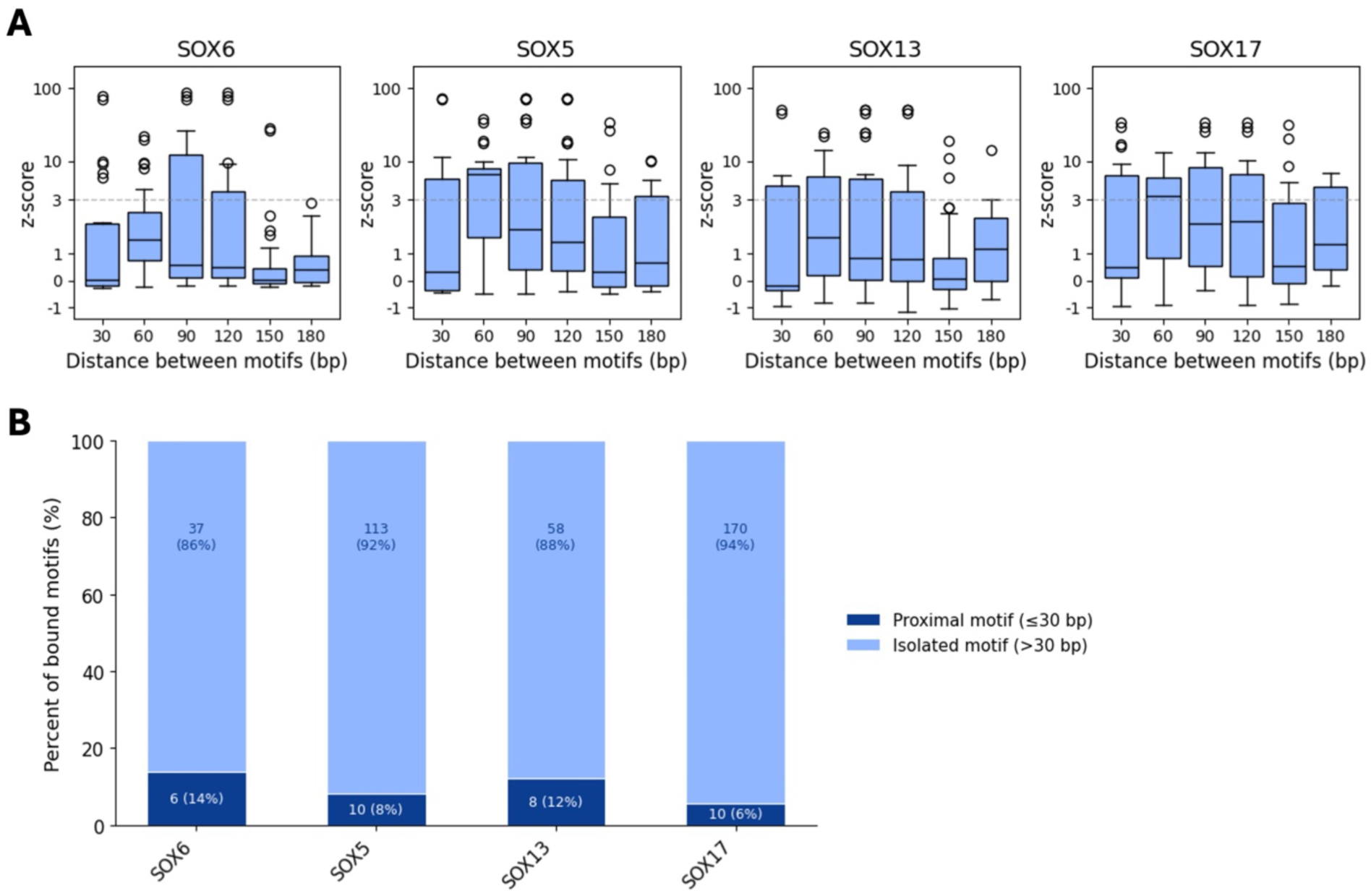
Limited evidence for cooperative binding at closely spaced SOX motifs. *(S2A) Little evidence for double-motif preference:* Boxplots show ChEC-seq z-scores for SOX motif occurrences on the same promoter, grouped by distance to the nearest cognate motif in 30 bp bins. Data are shown separately for each of the four reproducibly binding SOX TFs. *(S2B) Most binding occurs at isolated motifs:* Bar plots show, for each SOX TF, the number of bound motif occurrences classified as proximal (≤30 bp) or isolated (>30 bp). Binding is predominantly observed at isolated motifs.

**Figure S3:**
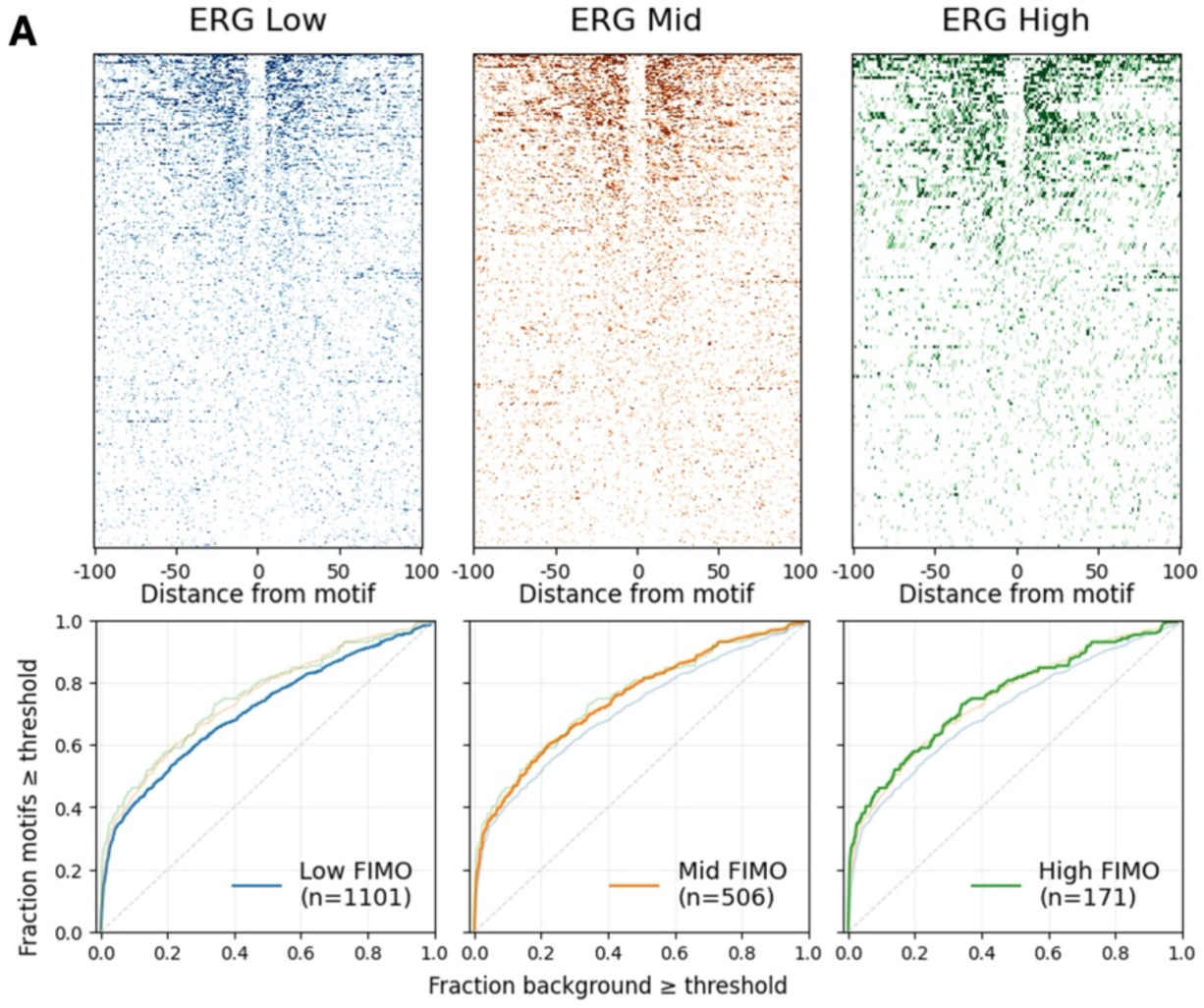
Motif binding is robust to motif score threshold. *(S3A) Similar fractions of motifs are bound across motif definitions:* Top, heatmaps show ERG ChEC-seq signal at all motif occurrences (±100 bp) defined using a default FIMO threshold (Low; left), the top two-thirds of FIMO scores (Mid; middle), or the top third of FIMO scores (High; right). Bottom, ROC-like curves depict enrichment of binding for each motif set, with the number of motif occurrences indicated.

**Figure S4:**
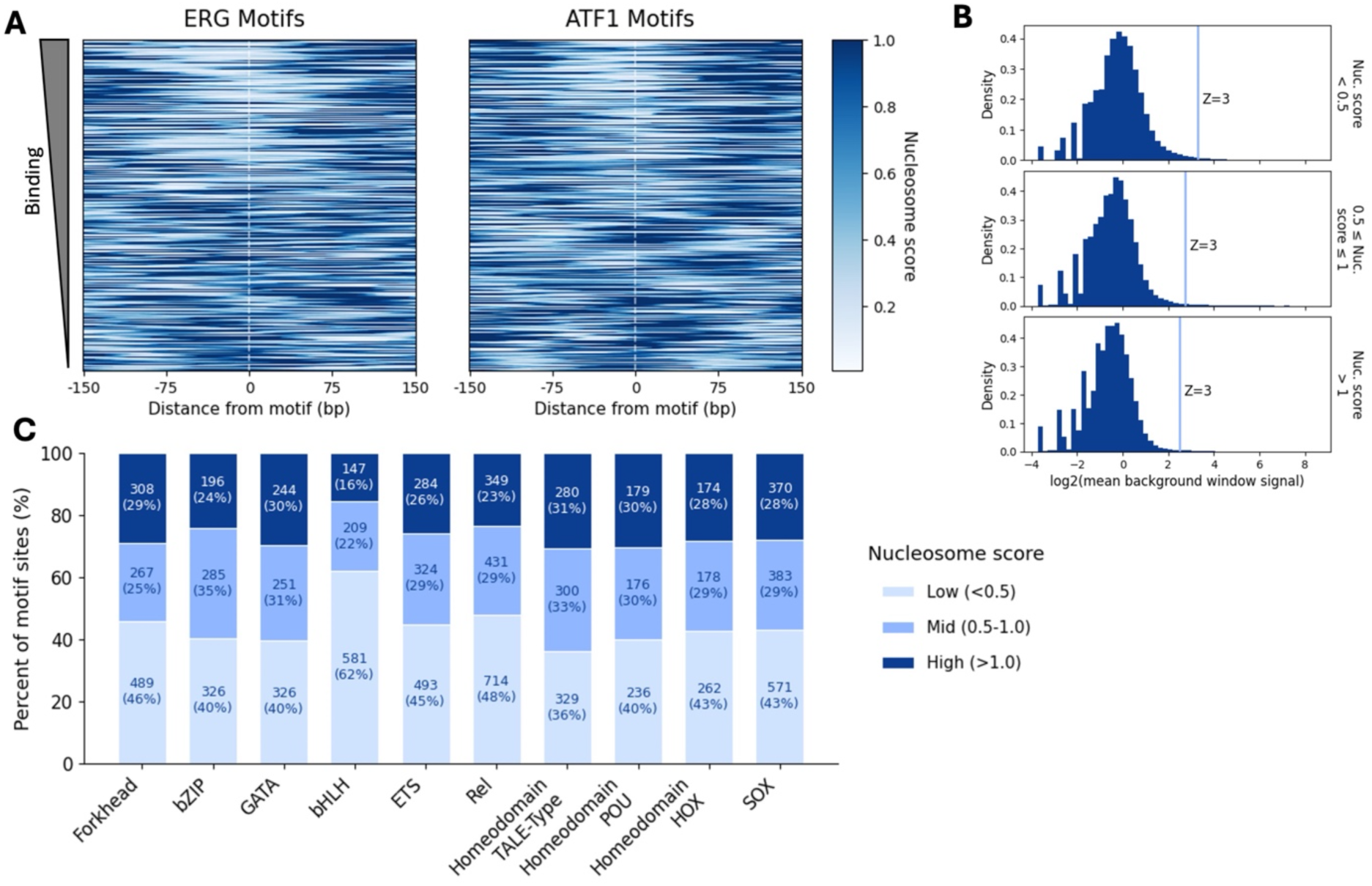
Additional analyses of nucleosome-dependent binding. *(S4A) Nucleosome occupancy across ERG and ATF1 motif instances:* Nucleosome score per base is shown for each motif occurrence of ERG (left) and ATF1 (right), ordered from strongest to weakest bound motif. Stronger binding is generally observed at lower nucleosome scores, although a substantial fraction of bound motifs resides in high-nucleosome regions. *(S4B) Background binding differs across nucleosome bins:* Histograms show the distribution of mean binding scores for all background genomic windows classified as low (<0.5), mid (0.5–1), or high (>1) nucleosome score (see Methods). The raw binding value corresponding to a z-score of 3 is higher in low-nucleosome windows than in high-nucleosome windows, reflecting differences in background signal across bins. *(S4C) Distribution of motif sites across nucleosome bins by family:* Stacked bar plots show the fraction and absolute number of all motif occurrences for each DBD family that fall into low, mid, or high nucleosome score bins.

**Figure S6:**
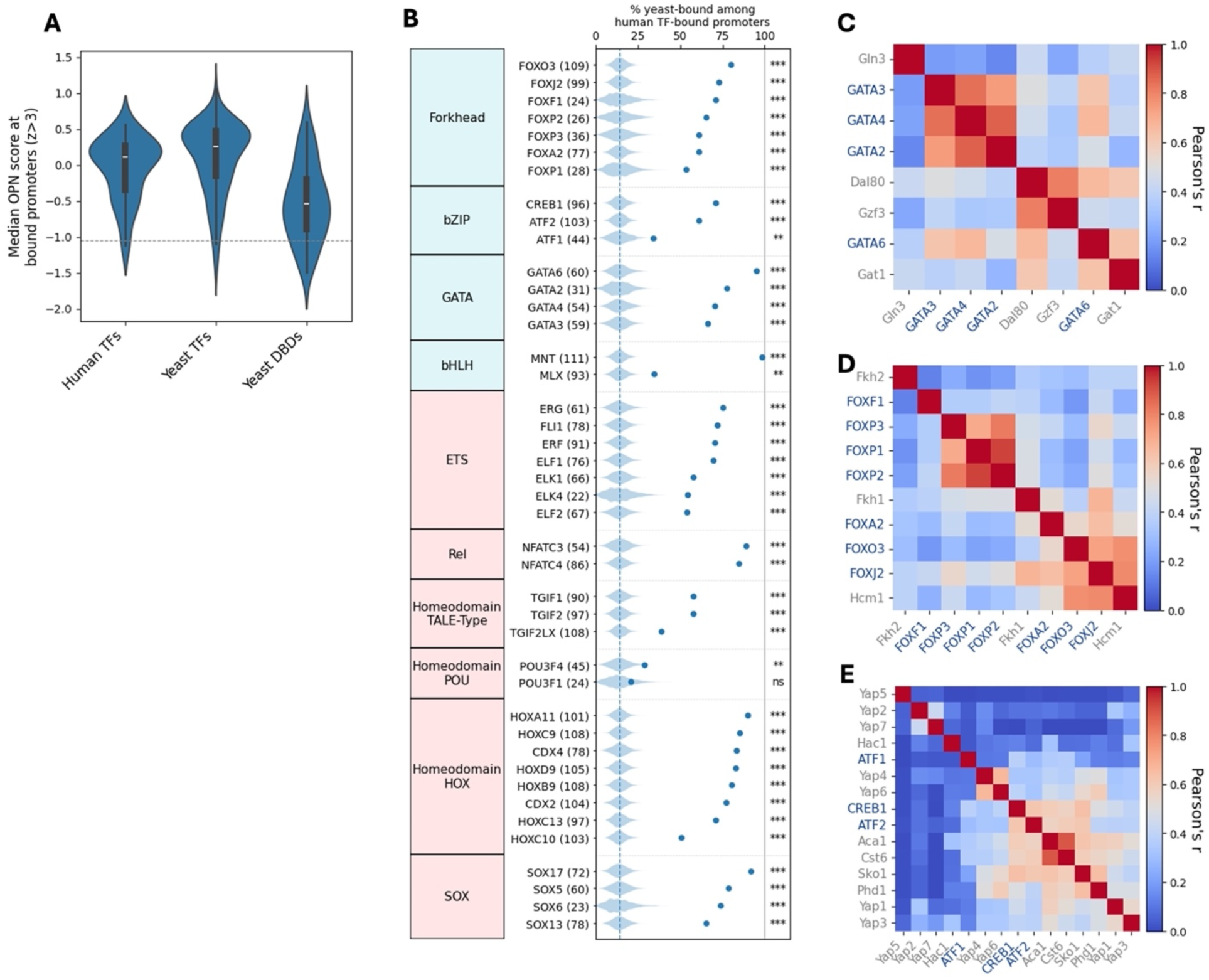
Additional analyses of OPN promoter binding and cross-species correlation. *(S6A) OPN preference across TF classes:* Violin plots show the distribution of the mean occupied proximal nucleosome (OPN) score of bound promoters (z-score >3) for human TFs, yeast TFs, and yeast DBD-only strains. Median (white line) and interquartile range are indicated. *(S6B) Enrichment of human TF binding at promoters co-bound by yeast TFs:* For each human TF, the blue dot indicates the fraction of bound promoters (z-score >3) that are also bound by at least three yeast TFs. The dashed line marks the true fraction of promoters in the entire yeast genome bound by at least three yeast TFs (14%). Distributions reflect 10,000 random samplings matched for the number of promoters bound by each human TF. Absolute numbers of human TF-bound promoters are shown in parentheses. One-tailed t-test p-values are reported in the rightmost column (*P<0.05; **P<0.01; ***P<0.001; ns, not significant). *(S6C–E) Promoter binding correlations between human and yeast TFs:* Heatmaps show Pearson correlations of summed, z-transformed promoter binding signals (see Methods) of GATA *(C)*, forkhead *(D)*, and bZIP *(E)* TFs. Human TFs are labeled in blue and yeast full-length TFs in gray.

**Figure S7:**
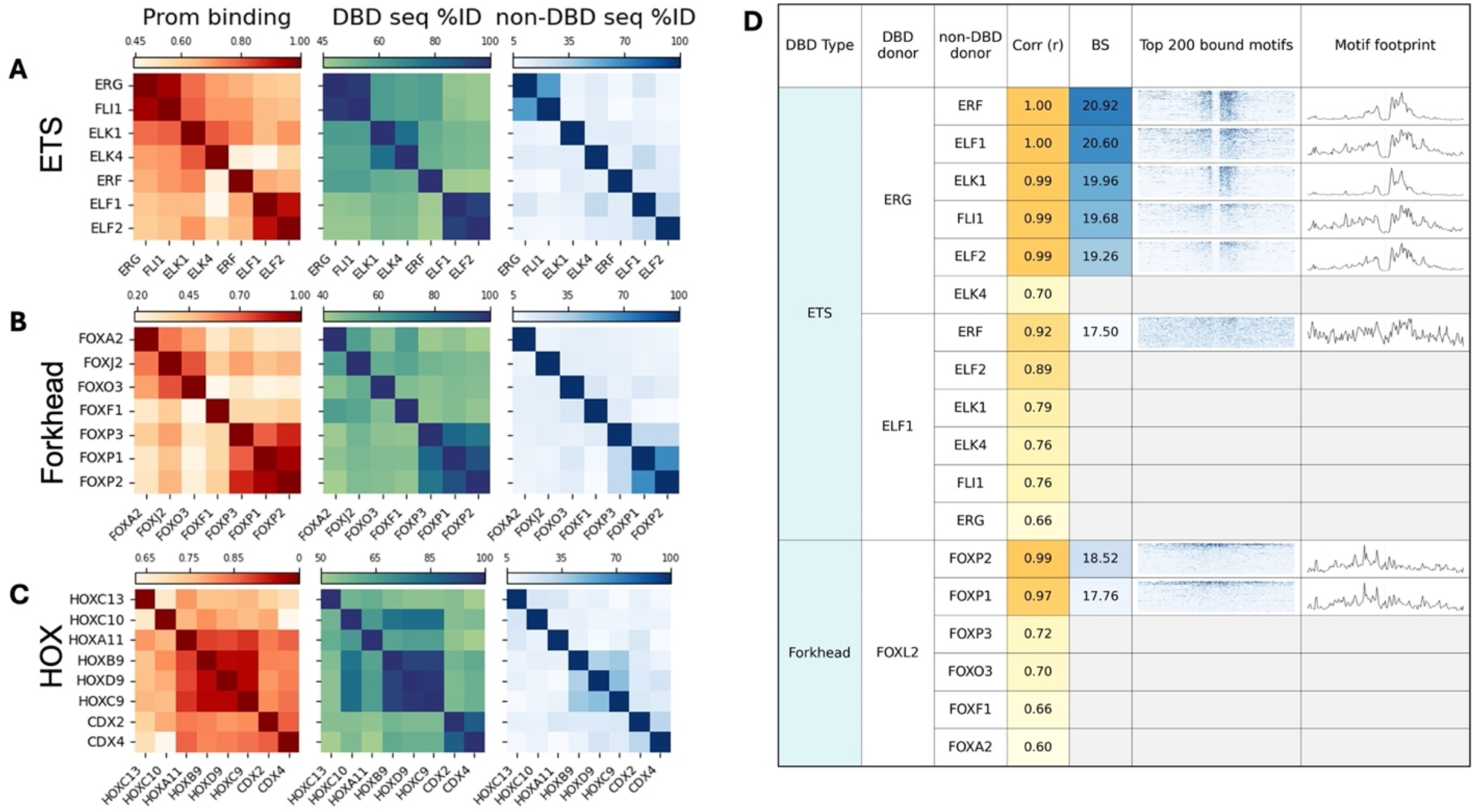
Sequence similarity and binding properties within and across TF families. *(S7A–C) Comparison of binding and sequence similarity across TF families:* For the ETS *(A)*, forkhead *(B)*, and HOX homeodomain *(C)* families, heatmaps show Pearson correlations of summed, z-transformed promoter binding signals (left), percent identity matrices for DBD multiple sequence alignments (center), and percent identity matrices for non-DBD regions (right). DBD and non-DBD boundaries were defined by InterProScan (see Methods). *(S7D) Summary of DBD swap strain binding properties:* Table lists DBD family, DBD donor TF, and non-DBD donor TF for each swap construct. Reproducibility (“Corr”) denotes the maximum Pearson correlation between biological replicates based on summed, z-transformed promoter binding signals (<0.9 considered irreproducible). Binding strength (BS) is defined in Methods. Heatmaps display ChEC-seq binding signal (±100 bp) at the top 200 bound motif instances for each swap strain, and the final panel shows the corresponding mean cleavage footprint (see Methods).

