## Supplemental Table for "Beyond motif recognition: Specificity of human transcription factors in yeast"

| TF Name | Source of repair | Primer F | Primer R | Anneal (°C) | gRNA | Strain seq |
|---|---|---|---|---|---|---|
| ATF4 | ORFeome | JBP007_ATF4_F | JBP044_attL2_pENTR223 | 58 | ura | ATGACCGAAATGAGCTTCCTGAGCAGCGAGGTGTTGGTGGGGGACTTGATGTCCCCCTTCGACCAGTCGGGTTTGGGGGCTGAAGAAAGCCTAGGTCTCTTAGATGATTACCTGGAGGTGGCCAAGCACTTCAAACCTCATGGGTTCTCCAGCGACAAGGCTAAGGCGGGCTCCTCCGAATGGCTGGCTGTGGATGGGTTGGTCAGTCCCTCCAACAACAGCAAGGAGGATGCCTTCTCCGGGACAGATTGGATGTTGGAGAAAATGGATTTGAAGGAGTTCGACTTGGATGCCCTGTTGGGTATAGATGACCTGGAAACCATGCCAGATGACCTTCTGACCACGTTGGATGACACTTGTGATCTCTTTGCCCCCCTAGTCCAGGAGACTAATAAGCAGCCCCCCCAGACGGTGAACCCAATTGGCCATCTCCCAGAAAGTTTAACAAAACCCGACCAGGTTGCCCCCTTCACCTTCTTACAACCTCTTCCCCTTTCCCCAGGGGTCCTGTCCTCCACTCCAGATCATTCCTTTAGTTTAGAGCTGGGCAGTGAAGTGGATATCACTGAAGGAGATAGGAAGCCAGACTACACTGCTTACGTTGCCATGATCCCTCAGTGCATAAAGGAGGAAGACACCCCTTCAGATAATGATAGTGGCATCTGTATGAGCCCAGAGTCCTATCTGGGGTCTCCTCAGCACAGCCCCTCTACCAGGGGCTCTCCAAATAGGAGCCTCCCATCTCCAGGTGTTCTCTGTGGGTCTGCCCGTCCCAAACCTTACGATCCTCCTGGAGAGAAGATGGTAGCAGCAAAAGTAAAGGGTGAGAAACTGGATAAGAAGCTGAAAAAAATGGAGCAAAACAAGACAGCAGCCACTAGGTACCGCCAGAAGAAGAGGGCGGAGCAGGAGGCTCTTACTGGTGAGTGCAAAGAGCTGGAAAAGAAGAACGAGGCTCTAAAAGAGAGGGCGGATTCCCTGGCCAAGGAGATCCAGTACCTGAAAGATTTGATAGAAGAGGTCCGCAAGGCAAGGGGGAAGAAAAGGGTCCCC |
| FOS | ORFeome | JBP008_FOS_F | JBP045_attL2_pDONR221_201 | 58 | ura | ATGATGTTCTCGGGCTTCAACGCAGACTACGAGGCGTCATCCTCCCGCTGCAGCAGCGCGTCCCCGGCCGGGGATAGCCTCTCTTACTACCACTCACCCGCAGACTCCTTCTCCAGCATGGGCTCGCCTGTCAACGCGCAGGACTTCTGCACGGACCTGGCCGTCTCCAGTGCCAACTTCATTCCCACGGTCACTGCCATCTCGACCAGTCCGGACCTGCAGTGGCTGGTGCAGCCCGCCCTCGTCTCCTCCGTGGCCCCATCGCAGACCAGAGCCCCTCACCCTTTCGGAGTCCCCGCCCCCTCCGCTGGGGCTTACTCCAGGGCTGGCGTTGTGAAGACCATGACAGGAGGCCGAGCGCAGAGCATTGGCAGGAGGGGCAAGGTGGAACAGTTATCTCCAGAAGAAGAAGAGAAAAGGAGAATCCGAAGGGAAAGGAATAAGATGGCTGCAGCCAAATGCCGCAACCGGAGGAGGGAGCTGACTGATACACTCCAAGCGGAGACAGACCAACTAGAAGATGAGAAGTCTGCTTTGCAGACCGAGATTGCCAACCTGCTGAAGGAGAAGGAAAAACTAGAGTTCATCCTGGCAGCTCACCGACCTGCCTGCAAGATCCCTGATGACCTGGGCTTCCCAGAAGAGATGTCTGTGGCTTCCCTTGATCTGACTGGGGGCCTGCCAGAGGTTGCCACCCCGGAGTCTGAGGAGGCCTTCACCCTGCCTCTCCTCAATGACCCTGAGCCCAAGCCCTCAGTGGAACCTGTCAAGAGCATCAGCAGCATGGAGCTGAAGACCGAGCCCTTTGATGACTTCCTGTTCCCAGCATCATCCAGGCCCAGTGGCTCTGAGACAGCCCGCTCCGTGCCAGACATGGACCTATCTGGGTCCTTCTATGCAGCAGACTGGGAGCCTCTGCACAGTGGCTCCCTGGGGATGGGGCCCATGGCCACAGAGCTGGAGCCCCTGTGCACTCCGGTGGTCACCTGTACTCCCAGCTGCACTGCTTACACGTCTTCCTTCGTCTTCACCTACCCCGAGGCTGACTCCTTCCCCAGCTGTGCAGCTGCCCACCGCAAGGGCAGCAGCAGCAATGAGCCTTCCTCTGACTCGCTCAGCTCACCCACGCTGCTGGCCCTGGAC |
| CREB5 | pDEST-AD | JBP005 | JBP006 | 58 | ura | ATGTTCTGCACCTCAGGAGGGAATTCAGCCTCAGTGATGTCCATGAGGCCTGTCCCAGGCTCTCTATCTTCTCTGCTACATCTCCACAACAGACAGAGACAGCCCATGCCAGCCTCCATGCCTGGGACCCTGCCCAACCCTACAATGCCAGGATCTTCCGCCGTCTTGATGCCAATGGAGCGACAAATGTCAGTGAACTCCAGCATCATGGGGATGCAAGGTCCAAATCTCAGCAACCCCTGTGCTTCTCCCCAGGTCCAGCCAATGCATTCAGAAGCCAAAATGAGGTTGAAGGCTGCATTGACTCACCACCCTGCTGCCATGTCAAATGGGAACATGAACACCATGGGACACATGATGGAGATGATGGGCTCCCGGCAGGACCAGACGCCACACCATCACATGCACTCGCACCCGCATCAGCACCAGACACTGCCACCCCATCACCCTTACCCACACCAGCACCAGCACCCAGCACACCATCCTCACCCTCAACCCCATCACCAGCAGAACCATCCACATCACCACTCCCATTCCCACCTTCATGCACACCCAGCACATCACCAGACCTCGCCACATCCGCCCCTGCACACCGGCAACCAAGCACAGGTTTCACCAGCAACACAACAGATGCAGCCAACCCAGACAATACAGCCACCCCAGCCCACAGGGGGGCGCCGGCGAAGGGTGGTAGACGAGGATCCGGACGAGAGGCGGCGGAAATTTCTGGAACGGAACCGGGCAGCTGCCACCCGCTGCAGACAGAAGAGGAAGGTCTGGGTGATGTCATTGGAAAAGAAAGCAGAAGAACTCACCCAGACAAACATGCAGCTTCAGAATGAAGTGTCTATGTTGAAAAATGAGGTGGCCCAGCTGAAACAGTTGTTGTTAACACATAAAGACTGCCCAATAACAGCCATGCAGAAAGAATCACAAGGATATCTAAGTCCAGAGAGTAGCCCTCCTGCTAGTCCTGTCCCAGCTTGCTCCCAGCAACAAGTCATCCAGCATAATACCATCACTACTTCCTCATCGGTCAGCGAGGTGGTAGGAAGCTCCACCCTCAGCCAGCTCACCACTCACAGAACAGACCTGAATCCGATTCTTTAG |
| ATF1 | pDEST-AD | JBP005 | JBP006 | 58 | ura | ATGGAAGATTCCCACAAGAGTACCACGTCAGAGACAGCACCTCAACCTGGTTCAGCAGTTCAGGGAGCTCACATTTCTCATATTGCTCAACAGGTATCATCTTTATCAGAAAGTGAGGAGTCCCAGGACTCATCCGACAGCATAGGCTCCTCACAGAAAGCCCACGGGATCCTAGCACGGCGCCCATCTTACAGAAAAATTTTGAAAGACTTATCTTCTGAAGATACACGGGGCAGAAAAGGAGACGGAGAAAATTCTGGAGTTTCTGCTGCTGTCACTTCTATGTCTGTTCCAACTCCCATCTATCAGACTAGCAGCGGACAGTATATTGCCATTGCCCCAAATGGAGCCTTACAGTTGGCAAGTCCAGGCACAGATGGAGTACAGGGACTTCAGACATTAACCATGACAAATTCAGGCAGTACTCAGCAAGGTACAACTATTCTTCAGTATGCACAGACCTCTGATGGACAGCAGATACTTGTGCCCAGCAATCAGGTGGTCGTACAAACTGCATCAGGAGATATGCAAACATATCAGATCCGAACTACACCTTCAGCTACTTCTCTGCCACAAACTGTGGTGATGACATCTCCTGTGACTCTCACCTCTCAGACAACTAAGACAGATGACCCCCAATTGAAAAGAGAAATAAGGTTAATGAAAAACAGAGAAGCTGCTCGAGAATGTCGCAGAAAGAAGAAAGAATATGTGAAATGCCTGGAAAACCGAGTTGCAGTCCTGGAAAATCAAAATAAAACTCTAATAGAAGAGTTAAAAACTTTGAAGGATCTTTATTCCAATAAAAGTGTTTAG |
| CREB1 | pDEST-AD | JBP005 | JBP006 | 58 | ura | ATGACCATGGAATCTGGAGCCGAGAACCAGCAGAGTGGAGATGCAGCTGTAACAGAAGCTGAAAACCAACAAATGACAGTTCAAGCCCAGCCACAGATTGCCACATTAGCCCAGGTATCTATGCCAGCAGCTCATGCAACATCATCTGCTCCCACCGTAACTCTAGTACAGCTGCCCAATGGGCAGACAGTTCAAGTCCATGGAGTCATTCAGGCGGCCCAGCCATCAGTTATTCAGTCTCCACAAGTCCAAACAGTTCAGTCTTCCTGTAAGGACTTAAAAAGACTTTTCTCCGGAACACAGATTTCAACTATTGCAGAAAGTGAAGATTCACAGGAGTCAGTGGATAGTGTAACTGATTCCCAAAAGCGAAGGGAAATTCTTTCAAGGAGGCCTTCCTACAGGAAAATTTTGAATGACTTATCTTCTGATGCACCAGGAGTGCCAAGGATTGAAGAAGAGAAGTCTGAAGAGGAGACTTCAGCACCTGCCATCACCACTGTAACGGTGCCAACTCCAATTTACCAAACTAGCAGTGGACAGTATATTGCCATTACCCAGGGAGGAGCAATACAGCTGGCTAACAATGGTACCGATGGGGTACAGGGCCTGCAAACATTAACCATGACCAATGCAGCAGCCACTCAGCCGGGTACTACCATTCTACAGTATGCACAGACCACTGATGGACAGCAGATCTTAGTGCCCAGCAACCAAGTTGTTGTTCAAGCTGCCTCTGGAGACGTACAAACATACCAGATTCGCACAGCACCCACTAGCACTATTGCCCCTGGAGTTGTTATGGCATCCTCCCCAGCACTTCCTACACAGCCTGCTGAAGAAGCAGCACGAAAGAGAGAGGTCCGTCTAATGAAGAACAGGGAAGCAGCTCGAGAGTGTCGTAGAAAGAAGAAAGAATATGTGAAATGTTTAGAAAACAGAGTGGCAGTGCTTGAAAATCAAAACAAGACATTGATTGAGGAGCTAAAAGCACTTAAGGACCTTTACTGCCACAAATCAGAT |
| ATF2 | pDEST-AD | JBP005 | JBP006 | 58 | ura | ATGAAATTCAAGTTACATGTGAATTCTGCCAGGCAATACAAGGACCTGTGGAATATGAGTGATGACAAACCCTTTCTATGTACTGCGCCTGGATGTGGCCAGCGTTTTACCAACGAGGATCATTTGGCTGTCCATAAACATAAACATGAGATGACACTGAAATTTGGTCCAGCACGTAATGACAGTGTCATTGTGGCTGATCAGACCCCAACACCAACAAGATTCTTGAAAAACTGTGAAGAAGTGGGTTTGTTTAATGAGTTGGCGAGTCCATTTGAGAATGAATTCAAGAAAGCTTCAGAAGATGACATTAAAAAAATGCCTCTAGATTTATCCCCTCTTGCAACACCTATCATAAGAAGCAAAATTGAGGAGCCTTCTGTTGTAGAAACAACTCACCAGGATAGTCCTTTACCTCACCCAGAGTCTACTACCAGTGATGAGAAGGAAGTACCATTGGCACAAACTGCACAGCCCACATCAGCTATTGTTCGTCCAGCATCATTACAGGTTCCCAATGTGCTGCTTACAAGTTCTGACTCAAGTGTAATTATTCAGCAGGCAGTACCTTCACCAACCTCAAGTACTGTAATCACCCAGGCACCATCCTCTAACAGGCCAATTGTCCCTGTACCAGGCCCATTTCCTCTTCTGTTACATCTTCCTAATGGACAAACCATGCCTGTTGCTATTCCTGCATCAATTACAAGTTCTAATGTGCATGTTCCAGCTGCAGTCCCACTCGTTCGACCAGTCACCATGGTGCCTAGTGTTCCAGGAATCCCAGGTCCTTCCTCTCCCCAACCAGTACAGTCAGAAGCAAAAATGAGATTAAAAGCTGCTTTGACCCAGCAACATCCTCCAGTTACCAATGGTGATACTGTCAAAGGTCATGGTAGCGGATTGGTTAGGACTCAGTCAGAGGAATCTCGACCGCAGTCATTACAACAGCCAGCCACATCCACTACAGAAACTCCGGCTTCTCCAGCTCACACAACTCCACAGACCCAAAGTACAAGTGGTCGTCGGAGAAGAGCAGCTAACGAAGATCCTGATGAAAAAAGGAGAAAGTTTTTAGAGCGAAATAGAGCAGCAGCTTCAAGATGCCGACAAAAAAGGAAAGTCTGGGTTCAGTCTTTAGAGAAGAAAGCTGAAGACTTGAGTTCATTAAATGGTCAGCTGCAGAGTGAAGTCACCCTGCTGAGAAATGAAGTGGCACAGCTGAAACAGCTTCTTCTGGCTCATAAAGATTGCCCTGTAACCGCCATGCAGAAGAAATCTGGCTATCATACTGCTGATAAAGATGATAGTTCAGAAGACATTTCAGTGCCGAGTAGTCCACATACAGAAGCTATACAGCATAGTTCGGTCAGCACATCCAATGGAGTCAGTTCAACCTCCAAGGCAGAAGCTGTAGCCACTTCAGTCCTCACCCAGATGGCGGACCAGAGTACAGAGCCTGCTCTTTCACAGATCGTTATGGCTCCTTCCTCCCAGTCACAGCCCTCAGGAAGT |
| FOXP3 | ORFeome | JBP009_FOXP3_F | JBP043_attL2_pENTR223.1 | 58 | ura | ATGCCCAACCCCAGGCCTGGCAAGCCCTCGGCCCCTTCCTTGGCCCTTGGCCCATCCCCAGGAGCCTCGCCCAGCTGGAGGGCTGCACCCAAAGCCTCAGACCTGCTGGGGGCCCGGGGCCCAGGGGGAACCTTCCAGGGCCGAGATCTTCGAGGCGGGGCCCATGCCTCCTCTTCTTCCTTGAACCCCATGCCACCATCGCAGCTGCAGCTGCCCACACTGCCCCTAGTCATGGTGGCACCCTCCGGGGCACGGCTGGGCCCCTTGCCCCACTTACAGGCACTCCTCCAGGACAGGCCACATTTCATGCACCAGCTCTCAACGGTGGATGCCCACGCCCGGACCCCTGTGCTGCAGGTGCACCCCCTGGAGAGCCCAGCCATGATCAGCCTCACACCACCCACCACCGCCACTGGGGTCTTCTCCCTCAAGGCCCGGCCTGGCCTCCCACCTGGGATCAACGTGGCCAGCCTGGAATGGGTGTCCAGGGAGCCGGCACTGCTCTGCACCTTCCCAAATCCCAGTGCACCCAGGAAGGACAGCACCCTTTCGGCTGTGCCCCAGAGCTCCTACCCACTGCTGGCAAATGGTGTCTGCAAGTGGCCCGGATGTGAGAAGGTCTTCGAAGAGCCAGAGGACTTCCTCAAGCACTGCCAGGCGGACCATCTTCTGGATGAGAAGGGCAGGGCACAATGTCTCCTCCAGAGAGAGATGGTACAGTCTCTGGAGCAGCAGCTGGTGCTGGAGAAGGAGAAGCTGAGTGCCATGCAGGCCCACCTGGCTGGGAAAATGGCACTGACCAAGGCTTCATCTGTGGCATCATCCGACAAGGGCTCCTGCTGCATCGTAGCTGCTGGCAGCCAAGGCCCTGTCGTCCCAGCCTGGTCTGGCCCCCGGGAGGCCCCTGACAGCCTGTTTGCTGTCCGGAGGCACCTGTGGGGTAGCCATGGAAACAGCACATTCCCAGAGTTCCTCCACAACATGGACTACTTCAAGTTCCACAACATGCGACCCCCTTTCACCTACGCCACGCTCATCCGCTGGGCCATCCTGGAGGCTCCAGAGAAGCAGCGGACACTCAATGAGATCTACCACTGGTTCACACGCATGTTTGCCTTCTTCAGAAACCATCCTGCCACCTGGAAGAACGCCATCCGCCACAACCTGAGTCTGCACAAGTGCTTTGTGCGGGTGGAGAGCGAGAAGGGGGCTGTGTGGACCGTGGATGAGCTGGAGTTCCGCAAGAAACGGAGCCAGAGGCCCAGCAGGTGTTCCAACCCTACACCTGGCCCC |
| FOXA2 | ORFeome | JBP010_FOXA2_F | JBP045_attL2_pDONR221_201 | 58 | ura | ATGCTGGGAGCGGTGAAGATGGAAGGGCACGAGCCGTCCGACTGGAGCAGCTACTATGCAGAGCCCGAGGGCTACTCCTCCGTGAGCAACATGAACGCCGGCCTGGGGATGAACGGCATGAACACGTACATGAGCATGTCGGCGGCCGCCATGGGCAGCGGCTCGGGCAACATGAGCGCGGGCTCCATGAACATGTCGTCGTACGTGGGCGCTGGCATGAGCCCGTCCCTGGCGGGGATGTCCCCCGGCGCGGGCGCCATGGCGGGCATGGGCGGCTCGGCCGGGGCGGCTGGCGTGGCGGGCATGGGGCCGCACTTGAGTCCCAGCCTGAGCCCGCTCGGGGGGCAGGCGGCCGGGGCCATGGGCGGCCTGGCCCCCTACGCCAACATGAACTCCATGAGCCCCATGTACGGGCAGGCGGGCCTGAGCCGCGCCCGCGACCCCAAGACCTACAGGCGCAGCTACACGCACGCAAAGCCGCCCTACTCGTACATCTCGCTCATCACCATGGCCATCCAGCAGAGCCCCAACAAGATGCTGACGCTGAGCGAGATCTACCAGTGGATCATGGACCTCTTCCCCTTCTACCGGCAGAACCAGCAGCGCTGGCAGAACTCCATCCGCCACTCGCTCTCCTTCAACGACTGTTTCCTGAAGGTGCCCCGCTCGCCCGACAAGCCCGGCAAGGGCTCCTTCTGGACCCTGCACCCTGACTCGGGCAACATGTTCGAGAACGGCTGCTACCTGCGCCGCCAGAAGCGCTTCAAGTGCGAGAAGCAGCTGGCGCTGAAGGAGGCCGCAGGCGCCGCCGGCAGCGGCAAGAAGGCGGCCGCCGGGGCCCAGGCCTCACAGGCTCAACTCGGGGAGGCCGCCGGGCCGGCCTCCGAGACTCCGGCGGGCACCGAGTCGCCTCACTCGAGCGCCTCCCCGTGCCAGGAGCACAAGCGAGGGGGCCTGGGAGAGCTGAAGGGGACGCCGGCTGCGGCGCTGAGCCCCCCAGAGCCGGCGCCCTCTCCCGGGCAGCAGCAGCAGGCCGCGGCCCACCTGCTGGGCCCGCCCCACCACCCGGGCCTGCCGCCTGAGGCCCACCTGAAGCCGGAACACCACTACGCCTTCAACCACCCGTTCTCCATCAACAACCTCATGTCCTCGGAGCAGCAGCACCACCACAGCCACCACCACCACCAGCCCCACAAAATGGACCTCAAGGCCTACGAACAGGTGATGCACTACCCCGGCTACGGTTCCCCCATGCCTGGCAGCTTGGCCATGGGCCCGGTCACGAACAAAACGGGCCTGGACGCCTCGCCCCTGGCCGCAGATACCTCCTACTACCAGGGGGTGTACTCCCGGCCCATTATGAACTCCTCTTTG |
| FOXF1 | ORFeome | JBP025_FOXF1_F | JBP045_attL2_pDONR221_201 | 58 | ura | ATGGACCCCGCGTCGTCCGGCCCGTCCAAGGCCAAGAAGACCAACGCCGGCATCCGGCGCCCGGAGAAGCCGCCCTATTCCTACATCGCGCTCATCGTCATGGCCATCCAGAGTTCACCCACCAAGCGCCTGACGCTGAGCGAGATCTACCAGTTCCTGCAGAGCCGCTTCCCCTTCTTCCGGGGCTCCTACCAGGGCTGGAAGAACTCCGTGCGCCACAACCTCTCGCTCAACGAGTGCTTCATCAAGCTACCCAAGGGCCTTGGGCGGCCCGGCAAGGGCCACTACTGGACCATCGACCCGGCCAGCGAGTTCATGTTCGAGGAGGGCTCCTTTCGGCGGCGGCCGCGCGGCTTCCGAAGGAAATGCCAGGCGCTCAAGCCCATGTACAGCATGATGAACGGGCTCGGCTTCAACCACCTCCCGGACACCTACGGCTTCCAGGGCTCGGCCGGCGGCCTCTCGTGCCCGCCCAACAGCCTGGCGCTGGAGGGCGGCCTGGGCATGATGAACGGCCACTTGCCGGGCAACGTGGACGGCATGGCCCTGCCCAGCCACTCGGTGCCCCACCTGCCTTCCAACGGCGGCCACTCGTACATGGGCGGCTGCGGCGGCGCGGCGGCCGGCGAGTACCCGCACCACGACAGCTCGGTGCCCGCCTCCCCGCTGCTGCCCACCGGCGCCGGTGGGGTCATGGAGCCGCACGCCGTCTACTCGGGCTCGGCGGCGGCCTGGCCGCCCTCGGCGTCCGCGGCGCTCAACAGCGGCGCCTCTTATATCAAGCAGCAGCCCCTGTCCCCCTGTAACCCCGCGGCCAACCCCCTGTCCGGCAGCCTCTCCACGCACTCCCTGGAGCAGCCGTATCTGCACCAGAACAGCCACAACGCCCCAGCCGAGCTGCAAGGCATCCCGCGGTATCACTCGCAGTCGCCCAGCATGTGTGACCGAAAGGAGTTTGTCTTCTCTTTCAACGCCATGGCGTCCTCTTCCATGCACTCGGCCGGCGGGGGCTCCTACTACCACCAGCAGGTCACCTACCAAGACATCAAGCCTTGCGTGATGTGGATCCAC |

| | | | | | | |
|---|---|---|---|---|---|---|
| FOXL1 | ORFeome | JBP028_FOXL1_F | JBP045_attL2_pDONR221_201 | 56 | ura | ATGAGTCACCTCTTCGATCCCCGGCTGCCTGCCCTGGCCGCCTCGCCCATGCTGTATCTGTACGGTCCCGAGAGACCCGG<br>CCTCCCTCTGGCCTTCGCCCCCGCGGCTGCTCTAGCTGCCTCGGGCCGGGCCGAGACCCCGCAGAAGCCTCCCTACAGCT<br>ACATCGCGCTCATCGCCATGGCGATCCAGGACGCGCCCGAGCAGAGGGTCACGCTCAACGGCATCTACCAGTTCATCATG<br>GACCGCTTCCCCTTCTACCACGACAACCGGCAGGGCTGGCAGAACAGCATCCGCCACAACCTCTCGCTCAACGACTGCTT<br>CGTCAAGGTGCCCCGCGAGAAAGGGCGGCCGGGCAAGGGCAGCTACTGGACGCTGGACCCCCGCTGCCTGGACATGTTTG<br>AGAACGGCAACTACCGGCGCCGGAAGAGGAAGCCCAAGCCGGGCCCCGGGGCCCCGGAGGCCAAGAGGCCCCGCGCCGAG<br>ACGCACCAGCGCAGCGCGGAGGCGCAGCCGGAGGCGGGGAGCGGGGCAGGGGGCTCGGGCCCCGCAATCTCCCGCCTGCA<br>GGCAGCGCCCGCGGGCCCCTCGCCCCTCCTGGACGGCCCCTCTCCGCCGGCGCCCCTCCACTGGCCGGGGACCGCGTCCC<br>CGAACGAGGACGCTGGTGACGCTGCCCAGGGCGCAGCGGCCGTGGCGGTCGGCCAGGCAGCGCGCACAGGGGACGGCCCG<br>GGGTCCCCTCTGCGCCCCGCCTCCCGCAGCTCTCCGAAGAGCTCCGACAAGTCCAAGAGCTTCAGCATAGACAGCATCCT<br>GGCGGGAAAGCAGGGCCAGAAGCCGCCTTCAGGGGACGAACTCCTAGGGGGTGCCAAGCCTGGGCCCGGCGGCCGTCTGG<br>GTGCCTCGCTCCTGGCCGCCTCCTCCAGCCTCCGTCCGCCTTTCAACGCTTCCCTGATGCTCGACCCGCATGTCCAGGGC<br>GGCTTTTACCAGCTCGGGATCCCCTTCCTCTCTTATTTCCCCCTGCAGGTTCCCGACACGGTACTCCACTTCCAG |
| FOXL2 | ORFeome | JBP029_FOXL2_F | JBP045_attL2_pDONR221_201 | 56 | ura | ATGATGGCCAGCTACCCCGAGCCCGAGGACGCGGCGGGGGCCCTGCTGGCCCCAGAGACCGGTCGCACAGTCAAGGAGCC<br>AGAAGGGCCGCCGCCGAGCCCAGGCAAGGGCGGTGGGGGTGGCGGCGGGACAGCCCCGGAGAAGCCGGACCCGGCGCAGA<br>AGCCCCCGTACTCGTACGTGGCGCTCATCGCCATGGCGATCCGCGAGAGCGCGGAGAAGAGGCTCACGCTGTCCGGCATC<br>TACCAGTACATCATCGCGAAGTTCCCGTTCTACGAGAAGAATAAGAAGGGCTGGCAAAATAGCATCCGCCACAACCTCAG<br>CCTCAACGAGTGCTTCATCAAGGTGCCGCGCGAGGGCGGCGGCGAGCGCAAGGGCAACTACTGGACGCTGGACCCGGCCT<br>GCGAAGACATGTTCGAGAAGGGCAACTACCGGCGCCGCCGCCGCATGAAGAGGCCCTTCCGGCCGCCGCCCGCGCACTTC<br>CAGCCCGGCAAGGGGCTCTTCGGGGCCGGAGGCGCCGCAGGCGGGTGCGGCGTGGCGGGCGCCGGGGCCGACGGCTACGG<br>CTACCTGGCGCCCCCCAAGTACCTGCAGTCTGGCTTCCTCAACAACTCGTGGCCGCTACCGCAGCCTCCCTCACCCATGC<br>CCTATGCCTCCTGCCAGATGGCGGCAGCCGCAGCGGCTGCAGCAGCTGCGGCTGCAGCCGCGGGCCCCGGTAGCCCTGGC<br>GCGGCCGCTGTGGTCAAGGGGCTGGCGGGCCCGGCCGCCTCGTACGGGCCGTACACACGCGTGCAGAGCATGGCGCTGCC<br>CCCCGGCGTAGTGAACTCGTACAATGGCCTGGGAGGCCCGCCGGCCGCACCCCCGCCTCCGCCGCACCCCCACCCGCATC<br>CGCACGCACACCATCTGCACGCGGCCGCCGCACCGCCGCCTGCCCCACCGCACCACGGGGCCGCCGCGCCGCCGCCGGGC<br>CAGCTCAGCCCTGCCAGCCCAGCCACCGCCGCGCCCCCGGCGCCCGCGCCCACCAGTGCGCCGGGCCTGCAGTTCGCTTG<br>TGCCCGGCAGCCCGAGCTCGCCATGATGCATTGCTCTTACTGGGACCACGACAGCAAGACCGGCGCGCTGCATTCGCGCC<br>TCGATCTC |
| FOXJ2 | pDEST-AD | JBP005 | JBP006 | 58 | ura | ATGGCTTCTGACCTAGAGAGTAGCCTCACCTCCATAGACTGGCTCCCCCAGCTGACCCTCCGAGCTACCATTGAGAAGCT<br>TGGAAGTGCCTCCCAGGCTGGGCCTCCCGGGAGCAGCCGCAAGTGTTCACCAGGGTCACCCACAGATCCTAATGCCACCC<br>TGAGCAAAGACGAGGCAGCAGTGCACCAGGACGGCAAGCCACGATACAGCTATGCCACTCTCATCACCTATGCCATCAAC<br>TCCTCTCCAGCCAAGAAGATGACCCTCAGCGAGATTTACCGCTGGATCTGTGATAACTTCCCCTATTACAAGAATGCTGG<br>CATTGGTTGGAAGAATTCAATACGGCACAACCTTTCTCTCAACAAGTGTTTCCGGAAGGTGCCCAGACCTCGGGATGACC<br>CTGGGAAGGGTTCCTATTGGACAATTGACACCTGCCCTGACATTTCCCGAAAGAGAAGACACCCTCCAGATGATGATCTG<br>TCCCAAGACTCACCAGAACAGGAGGCAAGCAAGAGCCCACGGGGAGGCGTTGCAGGGAGTGGAGAAGCCTCACTGCCTCC<br>TGAGGGGAATCCGCAGATGTCACTTCAGAGCCCCACATCTATAGCCAGCTACAGCCAGGGCACAGGATCTGTGGATGGTG<br>GAGCAGTGGCAGCAGGGGCTTCAGGCCGAGAAAGTGCTGAGGGTCCCCCTCCCCTCTATAACACCAACCATGACTTTAAA<br>TTCTCCTACTCAGAGATCAACTTTCAGGATCTAAGCTGGTCCTTCCGCAACCTCTATAAGTCCATGCTGGAGAAGTCCTC<br>TTCCTCCTCTCAGCACGGCTTTTCTTCTCTCCTGGGGGACATCCCACCCTCGAACAACTACTACATGTATCAGCAGCAGG<br>CACCGCCACCTCAACAGCAGCAGCAGCAGCAGCAGCCGCCACAGCCACCTCCCCAGCAGTCCCAGCCACAGCAGCAGCAG<br>GCACCTGCCCAGGGCCCCTCAGCTGTAGGGGGTGCTCCTCCACTGCACACCCCAAGCACAGATGGTTGTACCCCACCAGG<br>GGGAAAGCAAGCTGGGGCGGAAGGCTATGGGCCTCCCCCTGTAATGGCCATGCATCCACCCCCGCTGCAGCATGGAGGCT<br>ACCACCCTCATCAGCACCATCCCCACTCCCACCCTGCCCAGCAGCCACCACCTCCACAGCCACAGGCACAAGGCCAGGCT<br>CCCATCAACAACACTGGCTTTGCCTTTCCTTCTGACTGGTGCTCTAATATTGACTCTTTAAAGGAAAGCTTCAAGATGGT<br>GAATCGGCTCAATTGGTCCAGCATTGAGCAGTCACAATTCTCAGAACTGATGGAGAGTCTACGACAGGCAGAGCAGAAGA<br>ACTGGACCCTCGACCAGCATCACATTGCCAATCTGTGTGACTCCCTCAACCACTTCCTTACTCAGACTGGTCACGTGCCC<br>CCTCAAGGGGGTACCCACCGCCCACCAGCCCCTGCCCGTATTGCTGACTCCTGTGCCCTCACCAGTGGCAAACAGGAGTC<br>AGCCATGAGCCAAGTGAACTCTTATGGGCACCCACAAGCTCCCCACCTCTACCCTGGCCCATCACCAATGTACCCAATCC<br>CCACCCAGGACTCAGCAGGATACAATCGCCCAGCACACCATATGGTCCCTCGGCCATCAGTGCCACCTCCTGGTGCCAAT<br>GAGGAGATCCCTGATGACTTCGACTGGGACTTGATCACT |
| FOXO3 | pDEST-AD | JBP005 | JBP006 | 58 | ura | ATGGCAGAGGCACCGGCTTCCCCGGCCCCGCTCTCTCCGCTCGAAGTGGAGCTGGACCCGGAGTTCGAGCCCCAGAGCCG<br>TCCGCGATCCTGTACGTGGCCCCTGCAAAGGCCGGAGCTCCAAGCGAGCCCTGCCAAGCCCTCGGGGGAGACGGCCGCCG<br>ACTCCATGATCCCCGAGGAGGAGGACGATGAAGACGACGAGGACGGCGGGGGACGGGCCGGCTCGGCCATGGCGATCGGC<br>GGCGGCGGCGGGAGCGGCACGCTGGGCTCCGGGCTGCTCCTTGAGGACTCGGCCCGGGTGCTGGCACCCGGAGGGCAAGA<br>CCCCGGGTCTGGGCCAGCCACCGCGGCGGGCGGGCTGAGCGGGGGTACACAGGCGCTGCTGCAGCCTCAGCAACCGCTGC<br>CACCACCGCAGCCGGGGGCGGCTGGGGGCTCCGGGCAGCCGAGGAAATGTTCGTCGCGGCGGAACGCCTGGGGAAACCTG<br>TCCTACGCGGACCTGATCACCCGCGCCATCGAGAGCTCCCCGGACAAACGGCTCACTCTGTCCCAGATCTACGAGTGGAT<br>GGTGCGTTGCGTGCCCTACTTCAAGGATAAGGGCGACAGCAACAGCTCTGCCGGCTGGAAGAACTCCATCCGGCACAACC<br>TGTCACTGCATAGTCGATTCATGCGGGTCCAGAATGAGGGAACTGGCAAGAGCTCTTGGTGGATCATCAACCCTGATGGG<br>GGGAAGAGCGGAAAAGCCCCCCGGCGGCGGGCTGTCTCCATGGACAATAGCAACAAGTATACCAAGAGCCGTGGCCGCGC<br>AGCCAAGAAGAAGGCAGCCCTGCAGACAGCCCCCGAATCAGCTGACGACAGTCCCTCCCAGCTCTCCAAGTGGCCTGGCA<br>GCCCCACGTCACGCAGCAGTGATGAGCTGGATGCGTGGACGGACTTCCGTTCACGCACCAATTCTAACGCCAGCACAGTC<br>AGTGGCCGCCTGTCGCCCATCATGGCAAGCACAGAGTTGGATGAAGTCCAGGACGATGATGCGCCTCTCTCGCCCATGCT<br>CTACAGCAGCTCAGCCAGCCTGTCACCTTCAGTAAGCAAGCCGTGCACGGTGGAACTGCCACGGCTGACTGATATGGCAG<br>GCACCATGAATCTGAATGATGGGCTGACTGAAAACCTCATGGACGACCTGCTGGATAACATCACGCTCCCGCCATCCCAG<br>CCATCGCCCACTGGGGGACTCATGCAGCGGAGCTCTAGCTTCCCGTATACCACCAAGGGCTCGGGCCTGGGCTCCCCAAC<br>CAGCTCCTTTAACAGCACGGTGTTCGGACCTTCATCTCTGAACTCCCTACGCCAGTCTCCCATGCAGACCATCCAAGAGA<br>ACAAGCCAGCTACCTTCTCTTCCATGTCACACTATGGTAACCAGACACTCCAGGACCTGCTCACTTCGGACTCACTTAGC<br>CACAGCGATGTCATGATGACACAGTCGGACCCCTTGATGTCTCAGGCCAGCACCGCTGTGTCTGCCCAGAATTCCCGCCG<br>GAACGTGATGCTTCGCAATGATCCGATGATGTCCTTTGCTGCCCAGCCTAACCAGGGAAGTTTGGTCAATCAGAACTTGC<br>TCCACCACCAGCACCAAACCCAGGGCGCTCTTGGTGGCAGCCGTGCCTTGTCGAATTCTGTCAGCAACATGGGCTTGAGT<br>GAGTCCAGCAGCCTTGGGTCAGCCAAACACCAGCAGCAGTCTCCTGTCAGCCAGTCTATGCAAACCCTCTCGGACTCTCT<br>CTCAGGCTCCTCCTTGTACTCAACTAGTGCAAACCTGCCCGTCATGGGCCATGAGAAGTTCCCCAGCGACTTGGACCTGG<br>ACATGTTCAATGGGAGCTTGGAATGTGACATGGAGTCCATTATCCGTAGTGAACTCATGGATGCTGATGGGTTGGATTTT<br>AACTTTGATTCCCTCATCTCCACACAGAATGTTGTTGGTTTGAACGTGGGGAACTTCACTGGTGCTAAGCAGGCCTCATC<br>TCAGAGCTGGGTGCCAGGC |
| FOXP1 | ORFeome | JBP012_FOXP1_F | JBP045_attL2_pDONR221_201 | 58 | ura | ATGATGCAAGAATCTGGGACTGAGACAAAAAGTAACGGTTCAGCCATCCAGAATGGGTCGGGCGGCAGCAACCACTTACT<br>AGAGTGCGGCGGTCTTCGGGAGGGGCGGTCCAACGGAGAGACGCCGGCCGTGGACATCGGGGCAGCTGACCTCGCCCACG<br>CCCAGCAGCAGCAGCAACAGGCACTTCAGGTGGCAAGACAGCTCCTTCTTCAGCAGCAACAGCAGCAGCAAGTTAGTGGA<br>TTAAAATCTCCCAAGAGGAATGACAAACAACCAGCTCTTCAGGTTCCCGTGTCAGTGGCTATGATGACACCTCAAGTTAT<br>CACTCCCCAGCAAATGCAGCAGATCCTCCAGCAACAAGTGCTGAGCCCTCAGCAGCTCCAGGTTCTCCTCCAGCAGCAGC<br>AGGCCCTCATGCTTCAACAGCAGCAGCTTCAAGAGTTTTATAAAAAACAACAGGAACAGTTGCAGCTTCAACTTTTACAA<br>CAACAACATGCTGGAAAACAGCCTAAAGAGCAACAGCAGGTGGCTACCCAGCAGTTGGCTTTTCAGCAGCAGCTTTTACA<br>GATGCAGCAGTTACAGCAGCAGCACCTCCTGTCTTTGCAGCGCCAAGGCCTTCTGACAATTCAGCCCGGGCAGCCTGCCC<br>TTCCCCTTCAACCTCTTGCTCAAGGCATGATTCCAACAGAACTGCAGCAGCTCTGGAAAGAAGTGACAAGTGCTCATACT<br>GCAGAAGAAACCACAGGCAACAATCACAGCAGTTTGGATCTGACCACGACATGTGTCTCCTCCTCTGCACCTTCCAAGAC<br>CTCCTTAATAATGAACCCACATGCCTCTACCAATGGACAGCTCTCAGTCCACACTCCCAAAAGGGAAAGTTTGTCCCATG<br>AGGAGCACCCCCATAGCCATCCTCTCTATGGACATGGTGTATGCAAGTGGCCAGGCTGTGAAGCAGTGTGCGAAGATTTC<br>CAATCATTTCTAAAACATCTCAACAGTGAGCATGCGCTGGACGATAGAAGTACAGCCCAATGTAGAGTACAAATGCAGGT<br>TGTACAGCAGTTAGAGCTACAGCTTGCAAAAGACAAAGAACGCCTGCAAGCCATGATGACCCACCTGCATGTGAAGTCTA<br>CAGAACCCAAAGCCGCCCCTCAGCCCTTGAATCTGGTATCAAGTGTCACTCTCTCCAAGTCCGCATCGGAGGCTTCTCCA<br>CAGAGCTTACCTCATACTCCAACGACCCCAACCGCCCCCCTGACTCCCGTCACCCAAGGCCCCTCTGTCATCACAACCAC<br>CAGCATGCACACGGTGGGACCCATCCGCAGGCGGTACTCAGACAAATACAACGTGCCCATTTCGTCAGCAGATATTGCGC<br>AGAACCAAGAATTTTATAAGAACGCAGAAGTTAGACCACCATTTACATATGCATCTTTAATTAGGCAGGCCATTCTCGAA<br>TCTCCAGAAAAGCAGCTAACACTAAATGAGATCTATAACTGGTTCACACGAATGTTTGCTTACTTCCGACGCAACGCGGC<br>CACGTGGAAGAATGCAGTGCGTCATAATCTTAGTCTTCACAAGTGTTTTGTGCGAGTAGAAAACGTTAAAGGGGCAGTAT<br>GGACAGTGGATGAAGTAGAATTCCAAAAACGAAGGCCACAAAAGATCAGTGGTAACCCTTCCCTTATTAAAAACATGCAG<br>AGCAGCCACGCCTACTGCACACCTCTCAATGCAGCTTTACAGGCTTCAATGGCTGAGAATAGTATACCTCTATACACTAC<br>CGCTTCCATGGGAAATCCCACTCTGGGCAACTTAGCCAGCGCAATACGGGAAGAGCTGAACGGGGCAATGGAGCATACCA<br>ACAGCAACGAGAGTGACAGCAGTCCAGGCAGATCTCCTATGCAAGCCGTGCATCCTGTACACGTCAAAGAAGAGCCCCTC<br>GATCCAGAGGAAGCTGAAGGGCCCCTGTCCTTAGTGACAACAGCCAACCACAGTCCAGATTTTGACCATGACAGAGATTA<br>CGAAGATGAACCAGTAAACGAGGACATGGAGTCAGGCCTCATGGGC |
| FOXP2 | ORFeome | JBP030_FOXP2_F | JBP045_attL2_pDONR221_201 | 58 | ura | ATGATGCAGGAATCTGCGACAGAGACAATAAGCAACAGTTCAATGAATCAAAATGGAATGAGCACTCTAAGCAGCCAATT<br>AGATGCTGGCAGCAGAGATGGAAGATCAAGTGGTGACACCAGCTCTGAAGTAAGCACAGTAGAACTGCTGCATCTGCAAC<br>AACAGCAGGCTCTCCAGGCAGCAAGACAACTTCTTTTACAGCAGCAAACAAGTGGATTGAAATCTCCTAAGAGCAGTGAT<br>AAACAGAGACCACTGCAGGAATTGCTTCCAGAAACAAAATTATGTATCTGTGGCCACTCTTCTGGTGATGGGCATCCTCA<br>CAACACATTTGCAGTGCCTGTGTCAGTGGCCATGATGACTCCCCAGGTGATCACCCCTCAGCAAATGCAGCAGATCCTTC<br>AGCAACAAGTCCTGTCTCCTCAGCAGCTACAAGCCCTTCTCCAACAACAGCAGGCTGTCATGCTGCAGCAGCAACAACTA<br>CAAGAGTTTTACAAGAAACAGCAAGAGCAGTTACATCTTCAGCTTTTGCAGCAGCAGCAGCAACAGCAGCAGCAGCAACA<br>ACAGCAGCAACAACAGCAGCAGCAACAACAACAACAACAGCAGCAACAACAGCAGCAGCAGCAGCAACAGCAGCAGCAGC<br>AGCAACAGCATCCTGGAAAGCAAGCGAAAGAGCAGCAGCAGCAGCAGCAGCAGCAACAGCAATTGGCAGCCCAGCAGCTT<br>GTCTTCCAGCAGCAGCTTCTCCAGATGCAACAACTCCAGCAGCAGCAGCATCTGCTCAGCCTTCAGCGTCAGGGACTCAT<br>CTCCATTCCACCTGGCCAGGCAGCACTTCCTGTCCAATCGCTGCCTCAAGCTGGCTTAAGTCCTGCTGAGATTCAGCAGT<br>TATGGAAAGAAGTGACTGGAGTTCACAGTATGGAAGACAATGGCATTAAACATGGAGGGCTAGACCTCACTACTAACAAT<br>TCCTCCTCGACTACCTCCTCCAACACTTCCAAAGCATCACCACCAATAACTCATCATTCCATAGTGAATGGACAGTCTTC<br>AGTTCTAAGTGCAAGACGAGACAGCTCGTCACATGAGGAGACTGGGGCCTCTCACACTCTCTATGGCCATGGAGTTTGCA<br>AATGGCCAGGCTGTGAAAGCATTTGTGAAGATTTTGGACAGTTTTTAAAGCACCTTAACAATGAACACGCATTGGATGAC<br>CGAAGCACTGCTCAGTGTCGAGTGCAAATGCAGGTGGTGCAACAGTTAGAAATACAGCTTTCTAAAGAACGCGAACGTCT<br>TCAAGCAATGATGACCCACTTGCACATGCGACCCTCAGAGCCCAAACCATCTCCCAAACCTCTAAATCTGGTGTCTAGTG<br>TCACCATGTCGAAGAATATGTTGGAGACATCCCCACAGAGCTTACCTCAAACCCCTACCACACCAACGGCCCCAGTCACC<br>CCGATTACCCAGGGACCCTCAGTAATCACCCCAGCCAGTGTGCCCAATGTGGGAGCCATACGAAGGCGACATTCAGACAA<br>ATACAACATTCCCATGTCATCAGAAATTGCCCCAAACTATGAATTTTATAAAAATGCAGATGTCAGACCTCCATTTACTT<br>ATGCAACTCTCATAAGGCAGGCTATCATGGAGTCATCTGACAGGCAGTTAACACTTAATGAAATTTACAGCTGGTTTACA<br>CGGACATTTGCTTACTTCAGGCGTAATGCAGCAACTTGGAAGAATGCAGTACGTCATAATCTTAGCCTGCACAAGTGTTT<br>TGTTCGAGTAGAAAATGTTAAAGGAGCAGTATGGACTGTGGATGAAGTAGAATACCAGAAGCGAAGGTCACAAAAGATAA<br>CAGGAAGTCCAACCTTAGTAAAAAATATACCTACCAGTTTAGGCTATGGAGCAGCTCTTAATGCCAGTTTGCAGGCTGCC<br>TTGGCAGAGAGCAGTTTACCTTTGCTAAGTAATCCTGGACTGATAAATAATGCATCCAGTGGCCTACTGCAGGCCGTCCA<br>CGAAGACCTCAATGGTTCTCTGGATCACATTGACAGCAATGGAAACAGTAGTCCGGGCTGCTCACCTCAGCCGCACATAC<br>ATTCAATCCACGTCAAGGAAGAGCCAGTGATTGCAGAGGATGAAGACTGCCCAATGTCCTTAGTGACAACAGCTAATCAC<br>AGTCCAGAATTAGAAGACGACAGAGAGATTGAAGAAGAGCCTTTATCTGAAGATCTGGAA |

| | | | | | | |
|---|---|---|---|---|---|---|
| ELF1 | pDEST-AD | JBP005 | JBP006 | 58 | ura | ATGGCTGCTGTTGTCCAACAGAACGACCTAGTATTTGAATTTGCTAGTAACGTCATGGAGGATGAACGACAGCTTGGTGA<br>TCCAGCTATTTTTCCTGCCGTAATTGTGGAACATGTTCCTGGTGCTGATATTCTCAATAGTTATGCCGGTCTAGCCTGTG<br>TGGAAGAGCCCAATGACATGATTACTGAGAGTTCACTGGATGTTGCTGAAGAAGAAATCATAGACGATGATGATGATGAC<br>ATCACCCTTACAGTTGAAGCTTCTTGTCATGACGGGGATGAAACAATTGAAACTATTGAGGCTGCTGAGGCACTCCTCAA<br>TATGGATTCCCCTGGCCCTATGCTGGATGAAAAACGAATAAATAATAATATATTTAGTTCACCTGAAGATGACATGGTTG<br>TTGCCCCAGTCACCCATGTGTCCGTCACATTAGATGGGATTCCTGAAGTGATGGAAACACAGCAGGTGCAAGAAAAATAT<br>GCAGACTCACCGGGAGCCTCATCACCAGAACAGCCTAAGAGGAAAAAAGGAAGAAAAACTAAACCACCACGACCAGATTC<br>CCCAGCCACTACGCCAAATATATCTGTGAAGAAGAAAAACAAAGATGGAAAGGGAAACACAATTTATCTTTGGGAGTTTT<br>TACTGGCACTGCTCCAGGACAAGGCTACTTGTCCTAAATACATCAAGTGGACCCAGCGAGAGAAAGGCATTTTTAAATTG<br>GTGGATTCTAAAGCAGTGTCCAGGTTGTGGGGGAAGCACAAAAACAAACCTGATATGAATTATGAGACCATGGGAAGAGC<br>ACTCAGGTACTATTACCAAAGGGGTATTCTGGCAAAAGTGGAAGGTCAGCGCTTGGTGTATCAGTTTAAAGAAATGCCAA<br>AAGATCTTATATATATAAATGATGAGGATCCAAGTTCCAGCATAGAGTCTTCAGATCCATCACTATCTTCATCAGCCACT<br>TCAAATAGGAATCAAACCAGCCGGTCGAGAGTATCTTCAAGTCCAGGGGTAAAAGGAGGAGCCACTACAGTTCTAAAACC<br>AGGGAATTCTAAAGCTGCAAAACCCAAAGATCCTGTGGAAGTTGCACAACCATCAGAAGTTTTGAGGACAGTGCAGCCCA<br>CGCAGTCTCCATATCCTACCCAGCTCTTCCGGACTGTTCATGTAGTACAGCCAGTACAGGCTGTCCCAGAGGGAGAAGCA<br>GCTAGAACCAGTACCATGCAGGATGAAACATTAAATTCTTCCGTTCAGAGTATTAGGACTATACAGGCTCCAACCCAAGT<br>TCCAGTGGTTGTGTCTCCTAGGAATCAGCAGTTGCATACAGTAACACTCCAAACAGTGCCACTCACAACAGTTATAGCCA<br>GCACAGATCCATCAGCAGGTACTGGATCTCAGAAGTTTATTTTACAAGCCATTCCATCATCACAGCCCATGACAGTACTG<br>AAAGAAAATGTCATGCTGCAGTCACAAAAGGCGGGCTCTCCTCCTTCAATTGTCTTGGGCCCTGCCCAGGTTCAGCAGGT<br>CCTTACTAGCAATGTTCAGACCATTTGCAATGGAACCGTCAGTGTGGCTTCCTCTCCATCCTTCAGTGCTACTGCACCTG<br>TGGTGACCTTTTCTCCTCGCAGTTCACAGCTGGTTGCTCACCCACCTGGCACTGTAATCACTTCAGTTATCAAAACTCAA<br>GAAACAAAAACTCTTACACAGGAAGTAGAGAAAAAGGAATCTGAAGATCATTTGAAAGAGAACACTGAGAAAACGGAGCA<br>GCAGCCACAGCCTTATGTGATGGTAGTGTCCAGTTCCAATGGATTTACTTCTCAGGTAGCTATGAAACAAAACGAACTGC<br>TGGAACCCAACTCTTTTTAG |
| ELF2 | ORFeome | JBP031_ELF2_F | JBP045_attL2_pDONR221_201 | 58 | ura | ATGACATCAGCAGTGGTTGACAGTGGAGGTACTATTTTGGAGCTTTCCAGCAATGGAGTAGAAAATCAAGAGGAAAGTGA<br>AAAGGTTTCTGAATATCCAGCAGTGATTGTGGAGCCAGTTCCAAGTGCCAGATTAGAGCAGGGCTATGCAGCCCAGGTTC<br>TGGTTTATGATGATGAGACTTATATGATGCAAGATGTGGCAGAAGAACAAGAAGTTGAGACAGAGAATGTGGAAACAGTG<br>GAAGCATCAGTTCACAGCAGTAATGCACACTGTACAGATAAGACAATTGAAGCTGCTGAAGCCCTGCTTCATATGGAATC<br>TCCTACCTGCTTGAGGGATTCAAGAAGTCCTGAATTCATCCATGCTGCTATGAGGCCAGATGTCATTACAGAAACTGTAG<br>TGGAGGTGTCAACTGAAGAGTCTGAACCCATGGATACCTCTCCTATTCCAACATCACCAGATAGCCATGAACCAATGAAA<br>AAGAAAAAAGTTGGCCGTAAACCAAAGACCCAGCAATCACCAATTTCCAATGGCTCTCCTGAGTTAGGTATAAAGAAGAA<br>ACCAAGAGAAGGAAAAGGAAACACAACCTATTTGTGGGAGTTTCTTTTAGATCTACTTCAAGATAAAAATACTTGTCCCA<br>GGTATATTAAATGGACTCAGAGAGAAAAAGGCATATTCAAGCTGGTGGATTCAAAGGCTGTCTCTAAGCTTTGGGGAAAG<br>CATAAGAACAAACCAGACATGAACTATGAAACCATGGGACGAGCTTTGAGATACTACTACCAAAGGGGAATTCTTGCAAA<br>GGTTGAAGGACAGAGGCTTGTATATCAGTTCAAGGATATGCCGAAAAACATAGTGGTCATAGATGATGACAAAAGTGAAA<br>CCTGTAATGAAGATTTAGCAGGAACTACTGATGAAAAATCATTAGAACGAGTGTCACTGTCTGCAGAAAGTCTCCTGAAA<br>GCAGCATCCTCTGTTCGCAGTGGAAAAAATTCATCCCCTATAAACTGCTCCAGAGCAGAGAAGGGTGTAGCTAGAGTTGT<br>GAATATCACTTCCCCTGGGCACGATGCTTCATCCAGATCTCCTACTACCACTGCATCTGTGTCAGCAACAGCAGCTCCAA<br>GGACAGTTCGTGTGGCAATGCAGGTACCTGTTGTAATGACATCATTGGGTCAGAAAATTTCAACTGTGGCAGTTCAGTCA<br>GTTAATGCAGGCGCACCATTAATAACCAGCACTAGTCCAACAACAGCGACCTCTCCAAAGGTAGTCATTCAGACAATCCC<br>TACTGTGATGCCAGCTTCTACTGAAAATGGAGACAAAATCACCATGCAGCCTGCCAAAATTATTACCATCCCAGCTACAC<br>AGCTTGCACAGTGTCAACTGCAGACAAAGTCAAATCTGACTGGATCAGGAAGCATTAACATTGTTGGAACCCCATTGGCT<br>GTGAGAGCACTTACCCCTGTTTCAATAGCCCATGGTACACCTGTAATGAGACTATCAATGCCTACTCAGCAGGCATCTGG<br>CCAGACTCCTCCTCGAGTTATCAGTGCAGTCATAAAGGGGCCAGAGGTTAAATCGGAAGCAGTGGCAAAAAAGCAAGAAC<br>ATGATGTGAAAACTTTGCAGCTAGTAGAAGAAAAACCAGCAGATGGAAATAAGACAGTGACCCACGTAGTGGTTGTCAGT<br>GCGCCTTCAGCTATTGCCCTTCCTGTAACTATGAAAACAGAAGGACTAGTGACATGTGAGAAA |
| ERF | pDEST-AD | JBP005 | JBP006 | 58 | ura | ATGAAGACCCCGGCGGACACAGGGTTTGCCTTCCCGGATTGGGCCTACAAGCCAGAGTCGTCCCCTGGCTCAAGGCAGAT<br>CCAGCTGTGGCACTTTATCCTGGAGCTGCTGCGGAAGGAGGAGTACCAGGGCGTCATTGCCTGGCAGGGGGACTACGGGG<br>AATTCGTCATCAAAGACCCTGATGAGGTGGCCCGGCTGTGGGGCGTTCGCAAGTGCAAGCCCCAGATGAATTACGACAAG<br>CTGAGCCGGGCCCTGCGCTATTACTATAACAAGCGCATTCTGCACAAGACCAAGGGGAAACGGTTCACCTACAAGTTCAA<br>TTTCAACAAACTGGTGCTGGTCAATTACCCATTCATTGATGTGGGGTTGGCTGGGGGTGCAGTGCCCCAGAGTGCCCCGC<br>CAGTGCCGTCGGGTGGTAGCCACTTCCGCTTCCCTCCCTCAACGCCCTCCGAGGTGCTGTCCCCCACCGAGGACCCCCGC<br>TCACCACCAGCCTGCTCTTCATCTTCATCTTCCCTCTTCTCGGCTGTGGTGGCCCGCCGCCTGGGCCGAGGCTCAGTCAG<br>TGACTGTAGTGATGGCACGTCAGAGCTGGAGGAACCGCTGGGAGAGGATCCCCGCGCCCGACCACCCGGCCCTCCGGATC<br>TGGGTGCCTTCCGAGGGCCCCCGCTGGCCCGCCTGCCCCATGACCCTGGTGTCTTCCGAGTCTATCCCCGGCCTCGGGGT<br>GGCCCTGAACCCCTCAGCCCCTTCCCTGTGTCGCCTCTGGCCGGTCCTGGATCCCTGCTGCCCCCTCAGCTCTCCCCGGC<br>TCTGCCCATGACGCCCACCCACCTGGCCTACACTCCCTCGCCCACGCTGAGCCCGATGTACCCCAGTGGTGGCGGGGGGC<br>CCAGCGGCTCAGGGGGAGGCTCCCACTTCTCCTTCAGCCCTGAGGACATGAAACGGTACCTGCAGGCCCACACCCAAAGC<br>GTCTACAACTACCACCTCAGCCCCCGCGCCTTCCTGCACTACCCTGGGCTGGTGGTGCCCCAGCCCCAGCGCCCTGACAA<br>GTGCCCGCTGCCGCCCATGGCACCCGAGACCCCACCGGTCCCCTCCTCGGCCTCGTCATCCTCTTCTTCTTCTTCCTCCC<br>CATTCAAGTTTAAGCTCCAGCCGCCCCCACTCGGACGCCGGCAGCGGGCAGCTGGGGAGAAGGCCGTAGCCGGTGCTGAC<br>AAGAGCGGTGGCAGTGCAGGCGGGCTGGCTGAGGGGGCAGGGGCGCTAGCCCCACCGCCCCCGCCACCACAGATCAAGGT<br>GGAGCCCATCTCGGAAGGCGAGTCGGAGGAGGTAGAGGTGACTGACATCAGTGATGAGGATGAGGAAGACGGGGAGGTGT<br>TCAAGACGCCCCGTGCCCCACCTGCACCCCCTAAGCCTGAGCCCGGCGAGGCACCCGGGGCATCCCAGTGCATGCCCCTC<br>AAGCTACGCTTTAAGCGGCGCTGGAGTGAAGACTGTCGCCTCGAAGGGGGTGGGGGCCCCGCTGGGGGCTTTGAGGATGA<br>GGGTGAGGACAAGAAGGTGCGTGGGGAGGGGCCTGGGGAGGCTGGGGGGCCCCTCACCCCAAGGCGGGTGAGCTCTGACC<br>TCCAGCATGCCACGGCCCAGCTCTCCCTGGAGCACCGAGACTCCTAG |
| ELK1 | pDEST-AD | JBP005 | JBP006 | 58 | ura | ATGGACCCATCTGTGACGCTGTGGCAGTTTCTGCTGCAGCTGCTGAGAGAGCAAGGCAATGGCCACATCATCTCCTGGAC<br>TTCACGGGATGGTGGTGAATTCAAGCTGGTGGATGCAGAGGAGGTGGCCCGGCTGTGGGGGCTACGCAAGAACAAGACCA<br>ACATGAATTACGACAAGCTCAGCCGGGCCTTGCGGTACTACTATGACAAGAACATCATCCGCAAGGTGAGCGGCCAGAAG<br>TTCGTCTACAAGTTTGTGTCCTACCCTGAGGTCGCAGGGTGCTCCACTGAGGACTGCCCGCCCCAGCCAGAGGTGTCTGT<br>TACCTCCACCATGCCAAATGTGGCCCCTGCTGCTATACATGCCGCCCCAGGGGACACTGTCTCTGGAAAGCCAGGCACAC<br>CCAAGGGTGCAGGAATGGCAGGCCCAGGCGGTTTGGCACGCAGCAGCCGGAACGAGTACATGCGCTCGGGCCTCTATTCC<br>ACCTTCACCATCCAGTCTCTGCAGCCGCAGCCACCCCCTCATCCTCGGCCTGCTGTGGTGCTCCCCAGTGCAGCTCCTGC<br>AGGGGCAGCAGCGCCCCCCTCGGGGAGCAGGAGCACCAGTCCAAGCCCCTTGGAGGCCTGTCTGGAGGCTGAAGAGGCCG<br>GCTTGCCTCTGCAGGTCATCCTGACCCCGCCCGAGGCCCCAAACCTGAAATCGGAAGAGCTTAATGTGGAGCCGGGTTTG<br>GGCCGGGCTTTGCCCCCAGAAGTGAAAGTAGAAGGGCCCAAGGAAGAGTTGGAAGTTGCGGGGGAGAGAGGGTTTGTGCC<br>AGAAACCACCAAGGCCGAGCCAGAAGTCCCTCCACAGGAGGGCGTGCCAGCCCGGCTGCCCGCGGTTGTTATGGACACCG<br>CAGGGCAGGCGGGCGGCCATGCGGCTTCCAGCCCTGAGATCTCCCAGCCGCAGAAGGGCCGGAAGCCCCGGGACCTAGAG<br>CTTCCACTCAGCCCGAGCCTGCTAGGTGGGCCGGGACCCGAACGGACCCCAGGATCGGGAAGTGGCTCCGGCCTCCAGGC<br>TCCGGGGCCGGCGCTGACCCCATCCCTGCTTCCTACGCATACATTGACCCCGGTGCTGCTGACACCCAGCTCGCTGCCTC<br>CTAGCATTCACTTCTGGAGCACCCTGAGTCCCATTGCGCCCCGTAGCCCGGCCAAGCTCTCCTTCCAGTTTCCATCCAGT<br>GGCAGCGCCCAGGTGCACATCCCTTCTATCAGCGTGGATGGCCTCTCGACCCCCGTGGTGCTCTCCCCAGGGCCCCAGAA<br>GCCATAG |
| ELK4 | pDEST-AD | JBP005 | JBP006 | 58 | ura | ATGGACAGTGCTATCACCCTGTGGCAGTTCCTTCTTCAGCTCCTGCAGAAGCCTCAGAACAAGCACATGATCTGTTGGAC<br>CTCTAATGATGGGCAGTTTAAGCTTTTGCAGGCAGAAGAGGTGGCTCGTCTCTGGGGGATTCGCAAGAACAAGCCTAACA<br>TGAATTATGACAAACTCAGCCGAGCCCTCAGATACTATTATGTAAAGAATATCATCAAAAAAGTGAATGGTCAGAAGTTT<br>GTGTACAAGTTTGTCTCTTATCCAGAGATTTTGAACATGGATCCAATGACAGTGGGCAGGATTGAGGGTGACTGTGAAAG<br>TTTAAACTTCAGTGAAGTCAGCAGCAGTTCCAAAGATGTGGAGAATGGAGGGAAAGATAAACCACCTCAGCCTGGTGCCA<br>AGACCTCTAGCCGCAATGACTACATACACTCTGGCTTATATTCTTCATTTACTCTCAACTCTTTGAACTCCTCCAATGTA<br>AAGCTTTTCAAATTGATAAAGACTGAGAATCCAGCCGAGAAACTGGCAGAGAAAAAATCTCCTCAGGAGCCCACACCATC<br>TGTCATCAAATTTGTCACGACACCTTCCAAAAAGCCACCGGTTGAACCTGTTGCTGCCACCATTTCAATTGGCCCAAGTA<br>TTTCTCCATCTTCAGAAGAAACTATCCAAGCTTTGGAGACATTGGTTTCCCCAAAACTGCCTTCCCTGGAAGCCCCAACC<br>TCTGCCTCTAACGTAATGACTGCTTTTGCCACCACACCACCCATTTCGTCCATACCCCCTTTGCAGGAACCTCCCAGAAC<br>ACCTTCACCACCACTGAGTTCTCACCCAGACATCGACACAGACATTGATTCAGTGGCTTCTCAGCCAATGGAACTTCCAG<br>AGAATTTGTCACTGGAGCCTAAAGACCAGGATTCAGTCTTGCTAGAAAAGGACAAAGTAAATAATTCATCAAGATCCAAG<br>AAACCCAAAGGGTTAGAACTGGCACCCACCCTTGTGATCACGAGCAGTGATCCAAGCCCACTGGGAATACTGAGCCCATC<br>TCTCCCTACAGCTTCTCTTACACCAGCATTTTTTTCACAGGTAGCTTGCTCGCTCTTTATGGTGTCACCATTGCTTTCAT<br>TTATTTGCCCTTTTAAGCAAATCCAGAATTTATACACTCAAGTTTGCTTTCTGTTACTTAGGTTTGTCTTAGAAAGGTTA<br>TGTGTGACTGTCATGTAG |
| ERG | pDEST-AD | JBP005 | JBP006 | 58 | ura | ATGGTCAGCACTATTAAGGAAGCCTTATCAGTTGTGAGTGAGGACCAGTCGTTGTTTGAGTGTGCCTACGGAACGCCACA<br>CCTGGCTAAGACAGAGATGACCGCGTCCTCCTCCAGCGACTATGGACAGACTTCCAAGATGAGCCCACGCGTCCCTCAGC<br>AGGATTGGCTGTCTCAACCCCCAGCCAGGGTCACCATCAAAATGGAATGTAACCCTAGCCAGGTGAATGGCTCAAGGAAC<br>TCTCCTGATGAATGCAGTGTGGCCAAAGGCGGGAAGATGGTGGGCAGCCCAGACACCGTTGGGATGAACTACGGCAGCTA<br>CATGGAGGAGAAGCACATGCCACCCCCAAACATGACCACGAACGAGCGCAGAGTTATCGTGCCAGCAGATCCTACGCTAT<br>GGAGTACAGACCATGTGCGGCAGTGGCTGGAGTGGGCGGTGAAAGAATATGGCCTTCCAGACGTCAACATCTTGTTATTC<br>CAGAACATCGATGGGAAGGAACTGTGCAAGATGACCAAGGACGACTTCCAGAGGCTCACCCCCAGCTACAATGCCGACAT<br>CCTTCTCTCACATCTCCACTACCTCAGAGAGACTCCTCTTCCACATTTGACTTCAGATGATGTTGATAAAGCCTTACAAA<br>ACTCTCCACGGTTAATGCATGCTAGAAACACAGGGGGTGCAGCTTTTATTTTCCCAAATACTTCAGTATATCCTGAAGCT<br>ACGCAAAGAATTACAACTAGGCCAGATTTACCATATGAGCCCCCCAGGAGATCAGCCTGGACCGGTCACGGCCACCCCAC<br>GCCCCAGTCGAAAGCTGCTCAACCATCTCCTTCCACAGTGCCCAAAACTGAAGACCAGCGTCCTCAGTTAGATCCTTATC<br>AGATTCTTGGACCAACAAGTAGCCGCCTTGCAAATCCAGGCAGTGGCCAGATCCAGCTTTGGCAGTTCCTCCTGGAGCTC<br>CTGTCGGACAGCTCCAACTCCAGCTGCATCACCTGGGAAGGCACCAACGGGGAGTTCAAGATGACGGATCCCGACGAGGT<br>GGCCCGGCGCTGGGGAGAGCGGAAGAGCAAACCCAACATGAACTACGATAAGCTCAGCCGCGCCCTCCGTTACTACTATG<br>ACAAGAACATCATGACCAAGGTCCATGGGAAGCGCTACGCCTACAAGTTCGACTTCCACGGGATCGCCCAGGCCCTCCAG<br>CCCCACCCCCCGGAGTCATCTCTGTACAAGTACCCCTCAGACCTCCCGTACATGGGCTCCTATCACGCCCACCCACAGAA<br>GATGAACTTTGTGGCGCCCCACCCTCCAGCCCTCCCCGTGACATCTTCCAGTTTTTTGCTGCCCCAAACCCATACTGGA<br>ATTCACCAACTGGGGGTATATACCCCAACACTAGGCTCCCCACCAGCCATATGCCTTCTCATCTGGGCACTTACTAC |
| FLI1 | ORFeome | JBP011_FLI1_F | JBP045_attL2_pDONR221_201 | 58 | ura | ATGGACGGGACTATTAAGGAGGCTCTGTCGGTGGTGAGCGACGACCAGTCCCTCTTTGACTCAGCGTACGGAGCGGCAGC<br>CCATCTCCCCAAGGCCGACATGACTGCCTCGGGGAGTCCTGACTACGGGCAGCCCCACAAGATCAACCCCCTCCCACCAC<br>AGCAGGAGTGGATCAATCAGCCAGTGAGGGTCAACGTCAAGCGGGAGTATGACCACATGAATGGATCCAGGGAGTCTCCG<br>GTGGACTGCAGCGTTAGCAAATGCAGCAAGCTGGTGGGCGGAGGCGAGTCCAACCCCATGAACTACAACAGCTATATGGA<br>CGAGAAGAATGGCCCCCCTCCTCCCAACATGACCACCAACGAGAGGAGAGTCATCGTCCCCGCAGACCCCACACTGTGGA<br>CACAGGAGCATGTGAGGCAATGGCTGGAGTGGGCCATAAAGGAGTACAGCTTGATGGAGATCGACACATCCTTTTTCCAG<br>AACATGGATGGCAAGGAACTGTGTAAAATGAACAAGGAGGACTTCCTCCGCGCCACCACCCTCTACAACACGGAAGTGCT<br>GTTGTCACACCTCAGTTACCTCAGGGAAAGTTCACTGCTGGCCTATAATACAACCTCCCACACCGACCAATCCTCACGAT<br>TGAGTGTCAAAGAAGACCCTTCTTATGACTCAGTCAGAAGAGGAGCTTGGGGCAATAACATGAATTCTGGCCTCAACAAA<br>AGTCCTCCCCTTGGAGGGGCACAAACGATCAGTAAGAATACAGAGCAACGGCCCCAGCCAGATCCGTATCAGATCCTGGG<br>CCCGACCAGCAGTCGCCTAGCCAACCCTGGAAGCGGGCAGATCCAGCTGTGGCAATTCCTCCTGGAGCTGCTCTCCGACA<br>GCGCCAACGCCAGCTGTATCACCTGGGAGGGGACCAACGGGGAGTTCAAAATGACGGACCCCGATGAGGTGGCCAGGCGC<br>TGGGGCGAGCGGAAAAGCAAGCCCAACATGAATTACGACAAGCTGAGCCGGGCCCTCCGTTATTACTATGATAAAAACAT<br>TATGACCAAAGTGCACGGCAAAAGATATGCTTACAAATTTGACTTCCACGGCATTGCCCAGGCTCTGCAGCCACATCCGA<br>CCGAGTCGTCCATGTACAAGTACCCTTCTGACATCTCCTACATGCCTTCCTACCATGCCCACCAGCAGAAGGTGAACTTT<br>GTCCCTCCCCATCCATCCTCCATGCCTGTCACTTCCTCCAGCTTCTTTGGAGCCGCATCACAATACTGGACCTCCCCCAC<br>GGGGGGAATCTACCCCAACCCCAACGTCCCCCGCCATCCTAACACCCACGTGCCTTCACACTTAGGCAGCTACTACTTG |

| | | | | | | |
|---|---|---|---|---|---|---|
| MNT | ORFeome | JBP032_MNT_F | JBP045_attL2_pDONR221_201 | 58 | ura | ATGAGCATAGAGACGCTACTGGAGGCGGCCCGCTTCCTGGAATGGCAAGCGCAGCAACAACAGAGAGCACGTGAGGAGCA<br>GGAGCGGCTTCGCTTGGAGCAGGAGCGAGAGCAGGAACAGAAGAAGGCCAATAGCCTGGCCAGGCTGGCACATACCCTTC<br>CTGTGGAGGAACCCCGCATGGAGGCGCCACCCCTGCCTCTGTCTCCACCGGCTCCCCCGCCGGCACCCCCACCACCACTT<br>GCCACCCCTGCCCCACTGACTGTCATCCCTATCCCTGTGGTGACCAACTCCCCTCAGCCTCTACCCCCACCCCCACCCTT<br>GCCCGCGGCAGCCCAGCCTCTGCCCCTGGCGCCTCGTCAGCCGGCCCTGGTTGGCGCCCCCGGACTCAGCATTAAGGAGC<br>CTGCCCCCCTGCCCAGCAGGCCGCAGGTGCCCACCCCTGCTCCCCTACTGCCGGACTCGAAGGCCACCATTCCACCCAAT<br>GGCAGCCCCAAGCCTTTGCAGCCCCTCCCCACGCCTGTCCTGACCATAGCGCCACACCCTGGAGTCCAGCCTCAGCTGGC<br>CCCCCAGCAGCCGCCCCCACCCACGCTGGGGACCCTGAAGTTGGCACCAGCTGAAGAAGTCAAATCCAGTGAACAGAAGA<br>AGAGGCCCGGGGGGATCGGAACCAGAGAAGTCCACAACAAATTGGAGAAGAACAGGAGGGCCCATCTGAAAGAGTGCTTT<br>GAGACCCTGAAGCGGAACATCCCCAACGTGGATGACAAGAAGACGTCCAATCTGAGCGTGCTGCGGACGGCGCTGCGGTA<br>CATCCAGTCCCTGAAGAGGAAGGAGAAGGAATATGAGCATGAAATGGAGCGGCTGGCACGTGAGAAGATTGCCACGCAGC<br>AGCGGCTGGCAGAGCTCAAGCACGAGCTGAGCCAGTGGATGGACGTACTGGAGATTGACCGCGTGCTGCGGCAGACGGGC<br>CAGCCCGAGGATGACCAGGCCTCCACCTCCACCGCCTCTGAGGGTGAGGACAACATAGACGAGGATATGGAGGAGGACCG<br>GGCGGGCCTGGGCCCACCTAAGCTGAGCCATCGTCCCCAGCCGGAGCTGCTGAAGTCCACCCTGCCACCCCCCAGCACCA<br>CCCCTGCGCCTCTGCCTCCACACCCACACCCTCACCCCCACTCCGTGGCCCTACCTCCTGCCCACCTCCCCGTGCAGCAG<br>CAGCAGCCACAGCAGAAGACCCCTCTGCCAGCCCCTCCTCCCCCACCGGCTGCCCCTGCCCAGACACTGGTGCCAGCTCC<br>AGCCCATCTGGTGGCGACGGCTGGGGGTGGCTCCACGGTCATCGCCCACACAGCCACCACTCACGCTTCAGTCATCCAGA<br>CTGTGAACCACGTTCTGCAGGGGCCAGGCGGCAAGCACATCGCCCACATCGCCCCCTCGGCCCCCAGCCCTGCGGTGCAA<br>CTGGCGCCTGCCACACCCCCCATTGGGCACATCACTGTGCACCCTGCCACCCTCAACCATGTGGCCCACCTGGGCTCCCA<br>GCTGCCCTTGTACCCGCAGCCCGTGGCAGTGAGCCACATCGCCCACACCCTCTCGCACCAGCAAGTCAACGGCACGGCCG<br>GCCTGGGGCCCCCGGCTACTGTCATGGCAAAGCCGGCCGTGGGGGCTCAGGTGGTGCACCACCCCCAGCTGGTGGGCCAG<br>ACCGTGCTCAACCCTGTGACCATGGTCACCATGCCCTCCTTCCCAGTCAGCACACTCAAGCTGGCTTTG |
| MLXIPL | ORFeome | JBP033_MLXIPL_F | JBP045_attL2_pDONR221_201 | 58 | ura | ATGGCTGGCGCGCTGGCAGGCCTGGCCGCTGGCTTACAGGTCCCGAGGGTCGCGCCCAGCCCAGATTCTGATTCGGACAC<br>AGACTCTGAGGACCCGAGTCTCAGGCGCAGCGCGGGCGGCTTGCTCCGCTCGCAGGTCATTCACAGCGGTCACTTCATGG<br>TGTCTTCTCCGCACAGCGACTCTCTGCCCAGGCGGCGCGACCAGGAGGGGTCCGTGGGGCCCTCTGATTTCGGGCCTCGC<br>AGTATCGACCCCACACTCACACGCCTTTTCGAGTGCTTGAGCCTGGCCTACAGTGGCAAGCTGGTGTCACCCAAGTGGAA<br>GAATTTCAAAGGCCTCAAGCTCCTCTGCAGGGACAAGATCCGCCTGAACAACGCCATCTGGAGGGCCTGGTATATCCAGT<br>ATGTGAAGAGGAGGAAGAGCCCCGTGTGTGGCTTCGTGACCCCCCTGCAGGGGCCTGAGGCTGATGCGCACAGGAAGCCG<br>GAGGCCGTGGTCCTCGAAGGGAACTACTGGAAGCGGCGCATCGAGGTGGTGATGCGGGAATACCACAAGTGGCGCATCTA<br>CTACAAGAAGCGGCTTCGTAAGCCCAGTAGAGAAGATGACCTGCTGGCCCCTAAGCAGGCGGAAGGCAGATGGCCGCCGC<br>CGGAGCAATGGTGCAAACAGCTGTTCTCCAGTGTTGTCCCCGTGCTGCTGGGGGACCCAGAGGAAGAGCCGGGTGGGCGG<br>CAGCTCCTGGACCTCAATTGCTTTTTGTCCGACATCTCAGACACTCTCTTCACCATGACTCAGTCCGGCCCCTCGCCCCT<br>GCAGCTGCCGCCTGAGGATGCCTACGTCGGCAATGCTGACATGATCCAGCCGGACCTGACGCCACTGCAGCCAAGCCTGG<br>ATGACTTCATGGACATCTCAGATTTCTTTACCAACTCCCGCCTCCCACAGCCGCCCATGCCTTCCAACTTCCCAGAGCCT<br>CCCAGCTTCTCCCCTGTGGTTGACTCCCTCTTCAGCAGCGGGACCCTGGGCCCAGAGGTGCCCCCGGCTTCCTCCGCCAT<br>GACCCACCTCTCTGGACACAGCAGACTTCAGGCTCGGAACAGCTGCCCTGGACCCTTGGACTCCAGCGCCTTCCTGAGCT<br>CTGATTTCCTCCTTCCTGAAGACCCCAAACCTCGGCTCCCACCTCCTCCTGTACCCCCACCTCTGCTGCATTACCCTCCC<br>CCTGCCAAGGTGCCAGGACTGGAGCCCTGCCCCCCACCTCCCTTCCCTCCCATGGCACCACCTACTGCTTTGCTGCAGGA<br>AGAGCCTCTCTTCTCTCCCAGGTTTCCTTTCCCCACCGTCCCTCCTGCCCCAGGAGTGTCTCCTCTGCCTGCTCCTGCAG<br>CCTTCCCACCCACCCCACAGTCTGTCCCTAGCCCCGCCCCTACCCCTTTCCCTATAGAGCTTCTACCCTTGGGGTATTCG<br>GAGCCTGCCTTTGGCCCTTGCTTCTCCATGCCCAGAGGCAAGCCCCCCGCCCCATCCCCTAGGGGACAGAAAGCCAGCCC<br>TCCTACATTAGCCCCTGCCACTGCCAGTCCTCCTACCACCGCGGGGAGCAACAACCCCTGCCTCACACAGCTCCTCACAG<br>CAGCTAAGCCGGAGCAAGCCCTGGAGCCACCACTTGTATCCAGTACCCTCCTCCGGTCACCAGGGTCCCCGCAGGAGACA<br>GTCCCTGAATTCCCCTGCACATTCCTTCCCCCGACCCCTGCCCCTACACCGCCTCGGCCACCTCCAGGACCTGCTACATT<br>GGCCCCTTCAAGACCCCTGCTTGTCCCCAAAGCGGAGAGGCTCTCACCCCCAGCGCCCAGCGGGAGTGAACGGCGGCTGT<br>CAGGGGACCTTAGTTCCATGCCAGGCCCTGGGACTCTGAGTGTCCGTGTCTCTCCCCCGCAACCCATCCTCAGCCGGGGC<br>CGTCCAGACAGCAACAAGACCGAGAACCGGCGTATCACACACATCTCCGCGGAGCAGAAGCGGCGCTTCAACATCAAGCT<br>GGGGTTTGACACCCTTCATGGGCTCGTGAGCACACTCAGCGCCCAGCCCAGCCTCAAGGTGAGCAAAGCTACCACGCTGC<br>AGAAGACAGCTGAGTACATCCTTATGCTACAGCAGGAGCGTGCGGGCCTGCAGGAGGAGGCCCAGCAGCTGCGGGATGAG<br>ATTGAGGAGCTCAATGCCGCCATTAACCTGTGCCAGCAGCAGCTGCCCGCCACAGGGGTGCCCATCACACACCAGCGTTT |
| MLX | ORFeome | JBP034_MLX_F | JBP045_attL2_pDONR221_201 | 58 | ura | ATGACGGAGCCTGGAGCTTCTCCTGAGGATCCTTGGGTTAAAGCAAGCCCTGTGGGAGCTCATGCTGGAGAAGGAAGAGC<br>TGGTAGGGCTCGTGCACGTAGAGGGGCTGGAAGACGAGGAGCTTCTCTTCTTTCCCCAAAGTCTCCCACACTTTCCGTGC<br>CTAGGGGCTGCAGAGAAGACAGCTCTCATCCCGCGTGTGCTAAGGTGGAGTATGCCTATAGTGATAACAGCCTGGACCCC<br>GGGCTTTTTGTAGAAAGCACCCGCAAGGGGAGTGTAGTGTCCAGAGCTAATAGCATCGGTTCCACCAGTGCCTCTTCTGT<br>CCCCAACACAGATGATGAGGACAGTGATTACCACCAGGAGGCCTACAAGGAGTCCTACAAAGACCGGCGGCGGCGAGCAC<br>ACACTCAGGCTGAGCAGAAAAGGAGAGACGCCATCAAGAGAGGCTATGATGACCTTCAGACCATCGTCCCCACTTGTCAG<br>CAGCAGGACTTCTCCATTGGCTCCCAAAAGCTCAGCAAAGCCATCGTTCTACAAAAGACCATTGACTACATTCAGTTTTT<br>GCACAAGGAGAAGAAAAAGCAGGAGGAGGAGGTGTCCACGTTACGCAAGGATGTCACCGCCCTAAAGATCATGAAAGTGA<br>ACTATGAGCAGATTGTGAAGGCACACCAGGACAACCCCCATGAAGGGGAGGACCAGGTGTCTGACCAGGTCAAGTTCAAC<br>GTGTTTCAAGGCATCATGGATTCCCTGTTCCAGTCCTTCAATGCCTCCATCTCAGTGGCCAGCTTCCAGGAGCTGTCAGC<br>GTGTGTCTTCAGCTGGATCGAGGAGCACTGTAAGCCTCAGACCCTGCGGGAGATTGTGATTGGAGTCCTGCACCAATTGA<br>AAAACCAGCTTTAC |
| MXD4 | ORFeome | JBP035_MXD4_F | JBP045_attL2_pDONR221_201 | 58 | ura | ATGGAGCTGAACTCCCTGCTGATCCTGCTGGAGGCGGCCGAGTACCTGGAGCGCAGGGATCGAGAGGCCGAGCACGGCTA<br>CGCCTCGGTGCTGCCCTTCGACGGCGACTTCGCCAGGGAGAAAACAAAGGCGGCCGGCCTGGTGCGCAAGGCCCCGAACA<br>ACAGGTCTTCACACAACGAGCTAGAAAAGCACAGACGAGCCAAACTCAGGCTGTACCTTGAGCAGCTCAAGCAACTGGTG<br>CCCCTGGGCCCCGACAGCACCCGCCACACCACGCTGAGCCTCCTGAAGCGGGCCAAGGTGCACATCAAGAAACTGGAGGA<br>GCAGGACCGCCGGGCACTGAGCATCAAGGAGCAGCTGCAGCAGGAGCATCGTTTCCTGAAGCGGCGCCTGGAGCAGCTGT<br>CGGTGCAGAGCGTGGAGCGCGTGCGCACAGATAGCACGGGCTCTGCTGTCTCCACGGACGACTCAGAGCAAGAAGTGGAC<br>ATAGAGGGCATGGAGTTTGGCCCTGGTGAGCTGGACAGTGTTGGCAGCAGCAGTGACGCGGACGACCACTACAGCCTGCA<br>GAGTGGCACCGGCGGCGACAGTGGCTTCGGGCCCCACTGCCGGCGGCTGGGCCGCCCCGCCCTCTCG |
| CDX2 | pDEST-AD | JBP005 | JBP006 | 62 | ura | ATGTACGTGAGCTACCTCCTGGACAAGGACGTGAGCATGTACCCTAGCTCCGTGCGCCACTCTGGCGGCCTCAACCTGGC<br>GCCGCAGAACTTCGTCAGCCCCCCGCAGTACCCGGACTACGGCGGTTACCACGTGGCGGCCGCAGCTGCAGCGGCAGCGA<br>ACTTGGACAGCGCGCAGTCCCCGGGGCCATCCTGGCCGGCAGCGTATGGCGCCCCACTCCGGGAGGACTGGAATGGCTAC<br>GCGCCCGGAGGCGCCGCGGCCGCCGCCAACGCCGTGGCTCACGGCCTCAACGGTGGCTCCCCGGCCGCAGCCATGGGCTA<br>CAGCAGCCCCGCAGACTACCATCCGCACCACCACCCGCATCACCACCCGCACCACCCGGCCGCCGCGCCTTCCTGCGCTT<br>CTGGGCTGCTGCAAACGCTCAACCCCGGCCCTCCTGGGCCCGCCGCCACCGCTGCCGCCGAGCAGCTGTCTCCCGGCGGC<br>CAGCGGCGGAACCTGTGCGAGTGGATGCGGAAGCCGGCGCAGCAGTCCCTCGGCAGCCAAGTGAAAACCAGGACGAAAGA<br>CAAATATCGAGTGGTGTACACGGACCACCAGCGGCTGGAGCTGGAGAAGGAGTTTCACTACAGTCGCTACATCACCATCC<br>GGAGGAAAGCCGAGCTAGCCGCCACGCTGGGGCTCTCTGAGAGGCAGGTTAAAATCTGGTTTCAGAACCGCAGAGCAAAG<br>GAGAGGAAAATCAACAAGAAGAAGTTGCAGCAGCAACAGCAGCAGCAGCCACCACAGCCGCCTCCGCCGCCACCACAGCC<br>TCCCCAGCCTCAGCCAGGTCCTCTGAGAAGTGTCCCAGAGCCCTTGAGTCCGGTGTCTTCCCTGCAAGCCTCAGTGTCTG<br>GCTCTGTCCCTGGGGTTCTGGGGCCAACTGGGGGGGTGCTAAACCCCACCGTCACCCAGTGA |
| HOXA11 | ORFeome | JBP013_HOXA11_F | JBP045_attL2_pDONR221_201 | 58 | ura | ATGGATTTTGATGAGCGTGGTCCCTGCTCCTCTAACATGTATTTGCCAAGTTGTACTTACTACGTCTCGGGTCCAGATTT<br>CTCCAGCCTCCCTTCTTTTCTGCCCCAGACCCCGTCTTCGCGCCCAATGACATACTCCTACTCCTCCAACCTGCCCCAGG<br>TCCAACCCGTGCGCGAAGTGACCTTCAGAGAGTACGCCATTGAGCCCGCCACTAAATGGCACCCCCGCGGCAATCTGGCC<br>CACTGCTACTCCGCGGAGGAGCTCGTGCACAGAGACTGCCTGCAGGCGCCCAGCGCGGCCGGCGTGCCTGGCGACGTGCT<br>GGCCAAGAGCTCGGCCAACGTCTACCACCACCCCACCCCCGCAGTCTCGTCCAATTTCTATAGCACCGTGGGCAGGAACG<br>GCGTCCTGCCACAGGCTTTCGACCAGTTTTTCGAGACAGCCTACGGCACCCCGGAAAACCTCGCCTCCTCCGACTACCCC<br>GGGGACAAGAGCGCCGAGAAGGGGCCCCCGGCGGCCACGGCGACCTCCGCGGCGGCGGCGGCTGCAACGGGCGCGCC<br>GGCAACTTCAAGTTCGGACAGCGGCGGCGGCGGCTGCCGGGAGATGGCGGCGGCAGCAGAGGAGAAAGAGCGGCGGC<br>GGCGCCCCGAGAGCAGCAGCAGCCCCGAGTCGTCTTCCGGCCACACTGAGGACAAGGCCGGCGGCTCCAGTGGCCAACGC<br>ACCCGCAAAAAGCGCTGCCCCTATACCAAGTACCAGATCCGAGAGCTGGAACGGGAGTTCTTCTTCAGCGTCTACATTAA<br>CAAAGAGAAGCGCCTGCAACTGTCCCGCATGCTCAACCTCACTGATCGTCAAGTCAAAATCTGGTTTCAGAACAGGAGAA<br>TGAAGGAAAAAAAATTAACAGAGACCGTTTACAGTACTACTCAGCAAATCCACTCCTCTTG |
| HOXC10 | pDEST-AD | JBP005 | JBP006 | 58 | ura | ATGACATGCCCTCGCAATGTAACTCCGAACTCGTACGCGGAGCCCTTGGCTGCGCCCGGCGGAGGAGAGCGCTATAGCCG<br>GAGCGCAGGCATGTATATGCAGTCTGGGAGTGACTTCAATTGCGGGGTGATGAGGGGCTGCGGGCTCGCGCCCTCGCTCT<br>CCAAGAGGGACGAGGGCAGCAGCCCCAGCCTCGCCCTCAACACCTATCCGTCCTACCTCTCGCAGCTGGACTCCTGGGGC<br>GACCCCAAAGCCGCCTATCGCCTGGAACAACCTGTTGGCAGGCCGCTGTCCTCCTGCTCCTACCCACCTAGTGTCAAGGA<br>GGAGAATGTCTGCTGCATGTACAGCGCAGAGAAGCGGGCGAAAAGTGGCCCCGAGGCAGCTCTCTACTCCCACCCCTTGC<br>CGGAGTCCTGCCTTGGGGAGCACGAGGTACCCGTGCCCAGCTACTACCGCGCCAGCCCGAGCTACTCCGCGCTGGACAAG<br>ACGCCCCACTGTTCTGGGGCCAACGACTTCGAAGCCCCTTTCGAGCAGCGGGCCAGTCTCAACCCGCGCGCCGAACATCT<br>GGAATCGCCTCAGCTGGGGGGCAAAGTGAGTTTCCCTGAGACCCCCAAGTCCGACAGCCAGACCCCCAGCCCCAATGAAA<br>TCAAGACGGAGCAGAGCCTGGCGGGCCCTAAAGGGAGCCCCTCGGAGAGCGAAAAGGAGAGGGCCAAAGCTGCCGACTCC<br>AGCCCAGACACCTCGGATAACGAAGCGAAAGAGGAGATAAAGGCAGAAAACACCACAGGAAATTGGCTGACAGCAAAGAG<br>CGGAAGGAAGAAGAGGTGCCCCTATACTAAACACCAGACGCTGGAATTGGAGAAAGAATTTCTGTTCAATATGTATTTGA<br>CGCGAGAGCGCCGCCTGGAGATTAGCAAGACCATTAACCTTACAGACAGACAAGTCAAAATCTGGTTTCAAAATCGCAGA<br>ATGAAACTCAAGAAAATGAACCGAGAGAATCGGATCCGGGAACTGACCTCCAATTTTAATTTCACCTGA |
| HOXD9 | pDEST-AD | JBP005 | JBP006 | 58 | ura | ATGTCTTCCAGTGGCACCCTCAGCAACTACTACGTGGACTCGCTTATAGGCCATGAGGGCGACGAGGTGTTCGCGGCGCG<br>CTTCGGGCCGCCGGGGCCAGGCGCGCAGGGCCGGCCTGCAGGTGTGGCTGATGGCCCGGCCGCCACCGCCGCCGAGTTCG<br>CCTCGTGTAGTTTTGCCCCCAGATCGGCCGTGTTCTCTGCCTCGTGGTCCGCGGTGCCCTCCCAGCCCCCGGCAGCGGCG<br>GCGATGAGCGGCCTCTACCACCCGTACGTTCCCCCGCCGCCCCTGGCCGCCTCTGCCTCCGAGCCCGGCCGCTACGTGCG<br>CTCCTGGATGGAGCCGCTGCCCGGCTTCCCGGGCGGTGCGGGCGGTGGCGGTGGTGGTGGAGGCGGCGGTCCGGGCCGCG<br>GTCCCAGCCCTGGCCCCAGCGGCCCAGCCAACGGGCGCCACTACGGGATTAAGCCTGAAACCCGAGCGGCCCCGGCCCCC<br>GCCACGGCCGCCTCCACCACCTCCTCCTCCTCCACTTCCTTATCCTCCTCCTCCAAACGGACTGAGTGCTCCGTGGCCCG<br>GGAGTCCCAGGGGAGCAGCGGCCCCGAGTTCTCGTGCAACTCGTTCCTGCAGGAGAAGGCGGCAGCGGCGACGGGGGGAA<br>CCGGGCCTGGGGCAGGGATCGGGGCCGCGACTGGGACGGGCGGCTCGTCGGAGCCCTCAGCTTGCAGCGACCACCCGATC<br>CCAGGCTGTTCGCTGAAGGAGGAGGAGAAGCAGCATTCGCAGCCGCAGCAGCAGCAACTTGACCCAAACAACCCCGCCGC<br>GAACTGGATCCACGCTCGCTCCACCCGGAAAAAGCGCTGTCCCTACACCAAATACCAGACGCTTGAGCTGGAGAAAGAAT<br>TCCTCTTCAACATGTACCTCACCCGGGACCGGCGCTACGAGGTGGCCAGGATTCTCAACCTAACAGAGAGACAGGTCAAA<br>ATCTGGTTTCAGAACCGTAGGATGAAAATGAAAAAGATGAGCAAGGAGAAATGCCCCAAAGGAGAC |
| HOXA10 | ORFeome | JBP014_HOXA10_F | JBP045_attL2_pDONR221_201 | 58 | ura | ATGTCATGCTCGGAGAGCCCCGCCGCGAACTCTTTTTTGGTCGACTCGCTCATCAGCTCGGGCAGAGGCGAGGCAGGCGG<br>CGGTGGTGGTGGCGCGGGGGGCGGCGGCGGTGGCGGTTACTACGCCCACGGCGGGGTCTACCTGCCGCCCGCCGCCGACC<br>TGCCCTACGGGCTGCAGAGCTGCGGGCTCTTCCCCACGCTGGGCGGCAAGCGCAATGAGGCAGCGTCGCCGGGCAGCGGT<br>GGCGGTGGCGGGGGTCTAGGTCCCGGGGCGCACGGCTACGGGCCCTCGCCCATAGACCTGTGGCTAGACGCGCCCCGGTC<br>TTGCCGGATGGAGCCGCCTGACGGGCCGCCGCCGCCGCCCCAGCAGCAGCCGCCGCCCCCGCCGCAACCACCCCAGCCAG<br>CGCCGCAGGCCACCTCGTGCTCTTTCGCGCAGAACATCAAAGAAGAGAGCTCCTACTGCCTCTACGACTCGGCGGACAAA<br>TGCCCCAAAGTCTCGGCCACCGCCGCCGAACTGGCTCCCTTCCCGCGGGGCCCGCCGCCCGACGGCTGCGCCCTGGGCAC<br>CTCCAGCGGGGTGCCAGTGCCTGGCTACTTCCGCCTTTCTCAGGCCTACGGCACCGCCAAGGGCTATGGCAGCGGCGGCG<br>GCGGCGCGCAGCAACTCGGGGCTGGCCCGTTCCCCGCGCAGCCCCCGGGGCGCGGTTTCGATCTCCCGCCCGCGCTAGCC<br>TCCGGCTCGGCCGATGCGGCCCGGAAGGAGCGAGCCCTCGATTCGCCGCCGCCCCCCACGCTGGCTTGCGGCAGCGGCGG<br>GGGCTCGCAGGGCGACGAGGAGGCGCACGCGTCGTCCTCGGCCGCGGAGGAGCTCTCCCCGGCCCCTTCCGAGAGCAGCA<br>AAGCCTCGCCGGAGAAGGATTCCCTGGGCAATTCCAAAGGTGAAAACGCAGCCAACTGGCTCACGGCAAAGAGTGGTCGG<br>AAGAAGCGCTGCCCCTACACGAAGCACCAGACACTGGAGCTGGAGAAGGAGTTTCTGTTCAATATGTACCTTACTCGAGA<br>GCGGCGCCTAGAGATTAGCCGCAGCGTCCACCTCACGGACAGACAAGTGAAAATCTGGTTTCAGAACCGCAGGATGAAAC<br>TGAAGAAAATGAATCGAGAAAACCGGATCCGGGAGCTCACAGCCAACTTTAATTTTTCCTTG |

| | | | | | | |
|---|---|---|---|---|---|---|
| HOXC9 | ORFeome | JBP015_HOXC9_F | JBP044_attL2_pENTR 223 | 58 | ura | ATGTCGGCGACGGGGCCCATCAGTAACTATTACGTGGACTCGCTCATCTCTCACGACAATGAAGACCTCCTAGCGTCCAG<br>GTTTCCGGCCACCGGGGCTCATCCCGCCGCCGCCAGACCCAGCGGTTTGGTGCCGGACTGTAGCGATTTTCCGTCCTGTA<br>GCTTCGCGCCCAAGCCGGCAGTGTTCAGCACGTCGTGGGCGCCCGTGCCCTCTCAGTCGTCCGTGGTATATCACCCGTAC<br>GGCCCCCAGCCCCACCTCGGCGCCGACACGCGCTACATGCGGACTTGGCTCGAGCCGCTGTCCGGCGCCGTCTCCTTCCC<br>CAGCTTCCCGGCCGGGGGCCGTCACTACGCCCTCAAGCCGGACGCCTACCCCGGGCGCCGCGCGGACTGCGGCCCAGGGG<br>AGGGCCGCAGCTACCCGGACTACATGTACGGCTCGCCCGGGGAGCTGCGCGACCGCGCCCCGCAGACACTGCCCTCGCCC<br>GAGGCGGACGCGCTCGCCGGCAGCAAGCACAAAGAGGAGAAGGCTGACCTGGACCCCAGCAACCCCGTGGCCAACTGGAT<br>TCACGCCCGCTCCACGAGGAAGAAGCGCTGCCCCTACACCAAGTACCAGACGCTGGAACTGGAGAAGGAGTTTCTCTTCA<br>ATATGTATTTAACCAGGGACCGTCGGTATGAGGTGGCCCGGGTTCTCAATCTCACCGAGCGGCAGGTCAAAATCTGGTTT<br>CAGAATCGAAGGATGAAGATGAAAAAGATGAATAAAGAGAAAACCGACAAGGAGCAGTCC |
| HOXA9 | pDEST-AD | JBP005 | JBP006 | 58 | ura | ATGGCCACCACTGGGGCCCTGGGCAACTACTACGTGGACTCGTTCCTGCTGGGCGCCGACGCCGCGGATGAGCTGAGCGT<br>TGGCCGCTATGCGCCGGGGACCCTGGGCCAGCCTCCCCGGCAGGCGGCGACGCTGGCCGAGCACCCCGACTTCAGCCCGT<br>GCAGCTTCCAGTCCAAGGCGACGGTGTTTGGCGCCTCGTGGAACCCAGTGCACGCGGCGGGCGCCAACGCTGTACCCGCT<br>GCGGTGTACCACCACCATCACCACCACCCCTACGTGCACCCCCAGGCGCCCGTGGCGGCGGCGGCGCCGGACGGCAGGTA<br>CATGCGCTCCTGGCTGGAGCCCACGCCCGGTGCGCTCTCCTTCGCGGGCTTGCCCTCCAGCCGGCCTTATGGCATTAAAC<br>CTGAACCGCTGTCGGCCAGAAGGGGTGACTGTCCCACGCTTGACACTCACACTTTGTCCCTGACTGACTATGCTTGTGGT<br>TCTCCTCCAGTTGATAGAGAAAAACAACCCAGCGAAGGCGCCTTCTCTGAAAACAATGCTGAGAATGAGAGCGGCGGAGA<br>CAAGCCCCCCATCGATCCCAATAACCCAGCAGCCAACTGGCTTCATGCGCGCTCCACTCGGAAAAAGCGGTGCCCCTATA<br>CAAAACACCAGACCCTGGAACTGGAGAAAGAGTTTCTGTTCAACATGTACCTCACCAGGGACCGCAGGTACGAGGTGGCT<br>CGACTGCTCAACCTCACCGAGAGGCAGGTCAAGATCTGGTTCCAGAACCGCAGGATGAAAATGAAGAAAATCAACAAAGA<br>CCGAGCAAAAGACGAG |
| HOXB9 | ORFeome | JBP036_HOXB9_F | JBP045_attL2_pDONR 221_201 | 58 | ura | ATGTCCATTTCTGGGACGCTTAGCAGCTATTATGTCGACTCGATCATAAGTCACGAGAGTGAGGACGCGCCTCCAGCCAA<br>GTTTCCTTCTGGCCAGTACGCGAGCTCGCGGCAGCCGGGCCACGCGGAGCACCTGGAGTTCCCCTCGTGCAGCTTCCAGC<br>CCAAAGCGCCGGTGTTCGGCGCCTCCTGGGCGCCGCTGAGCCCGCACGCGTCCGGGAGCCTGCCGTCCGTCTACCACCCT<br>TACATCCAGCCCCAGGGCGTCCCGCCGGCCGAGAGCAGGTACCTCCGCACCTGGCTGGAGCCGGCGCCGCGCGGCGAAGC<br>GGCCCCGGGGCAGGGCCAGGCGGCGGTGAAGGCGGAGCCGCTGCTGGGCGCGCCTGGGGAGCTGCTCAAACAGGGCACGC<br>CCGAGTACAGTTTGGAAACTTCGGCGGGCAGGGAGGCCGTGCTGTCTAATCAAAGACCCGGCTACGGGGACAATAAAATT<br>TGCGAAGGAAGCGAGGACAAAGAGAGGCCGGATCAAACCAACCCCTCCGCCAACTGGCTGCACGCTCGCTCTTCCCGGAA<br>AAAGCGCTGTCCCTACACCAAATACCAGACGCTGGAGCTAGAGAAGGAGTTTCTGTTCAATATGTACCTCACCAGGGACC<br>GTAGGCACGAAGTGGCCAGACTCCTCAATCTGAGTGAGAGACAAGTCAAAATCTGGTTTCAGAACCGGCGGATGAAAATG<br>AAGAAAATGAATAAGGAGCAGGGCAAAGAG |
| CDX4 | pDEST-AD | JBP005 | JBP006 | 58 | ura | ATGTACGGAAGCTGTCTTTTGGAGAAAGAAGCAGGCATGTACCCGGGCACTCTCATGAGCCCTGGGGGCGACGGCACAGC<br>TGGGACAGGCGGCACAGGGGGCGGTGGGAGTCCGATGCCAGCCTCCAATTTCGCTGCGGCACCGGCTTTCTCGCACTATA<br>TGGGGTATCCTCATATGCCCAGCATGGATCCTCACTGGCCGTCTCTGGGAGTCTGGGGCTCACCCTACAGTCCCCCGCGA<br>GAAGACTGGAGCGTGTATCCTGGGCCGTCTAGTACAATGGGCACAGTGCCGGTGAACGACGTGACCTCTAGCCCCGCCGC<br>TTTCTGCTCGACCGACTACAGCAACTTGGGCCCTGTGGGCGGTGGAACTAGCGGCAGCAGCCTACCAGGCCAGGCTGGCG<br>GGTCGCTTGTCCCGACGGACGCAGGCGCCGCCAAGGCCAGTTCCCCCAGCAGGAGCCGCCACAGCCCCTATGCATGGATG<br>CGCAAGACGGTGCAGGTGACGGGGAAAACCAGGACAAAAGAAAAGTATCGTGTAGTTTACACTGATCATCAAAGATTGGA<br>GCTGGAAAAGGAATTCCATTGCAATAGATATATCACCATCCAGAGAAAATCAGAGCTGGCAGTTAACCTGGGCCTTTCCG<br>AGAGACAGGTGAAAATCTGGTTTCAGAATCGCAGAGCCAAGGAGAGAAAGATGATCAAAAAGAAAATCTCCCAGTTTGAG<br>AATAGTGGAGGCTCGGTGCAAAGTGACTCTGACTCCATCAGCCCTGGGGAACTACCTAACACTTTTTTCACCACACCATC<br>TGCTGTTCGTGGATTTCAACCTATTGAGATACAGCAGGTTATAGTCTCCGAA |
| HOXC13 | ORFeome | JBP037_HOXC13_F | JBP045_attL2_pDONR 221_201 | 58 | ura | ATGACGACTTCGCTGCTCCTGCATCCACGCTGGCCGGAGAGCCTTATGTACGTCTATGAGGACAGCGCGGCGGAGAGCGG<br>CATCGGCGGCGGCGGCGGAGGAGGAGGCGGCGGCACGGGCGGAGCGGGGGGTGGCTGCAGCGGAGCGAGCCCCGGCAAAG<br>CCCCGAGCATGGATGGTCTGGGCAGCAGCTGCCCGGCCAGCCACTGCCGCGACCTGCTTCCGCACCCCGTGCTGGGCCGC<br>CCGCCGGCTCCCCTGGGCGCCCCTCAGGGCGCCGTCTATACGGACATCCCGGCCCCGGAGGCGGCGCGCCAGTGTGCCCC<br>GCCGCCCGCACCCCCCACCTCGTCCAGCGCCACCCTGGGCTACGGCTACCCCTTCGGGGGCAGCTACTACGGCTGCCGCC<br>TGTCGCACAACGTGAACCTGCAGCAGAAGCCTTGCGCCTACCACCCGGGCGATAAATACCCGGAGCCGTCGGGCGCCCTG<br>CCCGGTGACGACCTGTCCTCTAGGGCCAAGGAGTTCGCCTTCTACCCCAGCTTCGCCAGCTCCTACCAGGCGATGCCCGG<br>CTACCTGGACGTGTCGGTGGTGCCCGGGATCAGCGGGCACCCGGAGCCGCGTCACGACGCCCTCATCCCCGTCGAAGGCT<br>ACCAGCACTGGGCTCTCTCCAATGGCTGGGACAGTCAGGTGTACTGCTCCAAGGAGCAGTCGCAGTCCGCCCACCTCTGG<br>AAGTCTCCCTTCCCAGACGTGGTTCCCCTGCAGCCCGAGGTGAGCAGCTACCGGCGCGGGCGCAAGAAACGCGTGCCCTA<br>CACTAAGGTGCAGCTGAAGGAGCTAGAGAAGGAATACGCGGCTAGCAAGTTCATCACCAAAGAGAAGCGCCGGCGCATCT<br>CCGCCACCACGAACCTCTCTGAGCGCCAGGTAACCATCTGGTTCCAGAACCGGCGGGTCAAAGAGAAGAAGGTGGTCAGC<br>AAATCGAAAGCGCCTCATCTCCACTCCACC |
| NFATC4 | pDEST-AD | JBP005 | JBP006 | 58 | ura | ATGGGGGCGGCCAGCTGCGAGGATGAGGAGCTGGAATTTAAGCTGGTGTTCGGGGAGGAAAAGGAGGCCCCCCCGCTGGG<br>CGCGGGGGGATTGGGGGAAGAACTGGACTCAGAGGATGCCCCGCCATGCTGCCGTCTGGCCTTGGGAGAGCCCCCTCCCT<br>ATGGCGCTGCACCTATCGGTATTCCCCGACCTCCACCCCCTCGGCCTGGCATGCATTCGCCACCGCCGCGACCAGCCCCC<br>TCACCTGGCACCTGGGAGAGCCAGCCCGCCAGGTCGGTGAGGCTGGGAGGACCAGGAGGGGGTGCTGGGGGTGCTGGGGG<br>TGGCCGTGTTCTCGAGTGTCCCAGCATCCGCATCACCTCCATCTCTCCCACGCCGGAGCCGCCAGCAGCGCTGGAGGACA<br>ACCCTGATGCCTGGGGGGACGGCTCTCCTAGAGATTACCCCCCACCAGAAGGCTTTGGGGGCTACAGAGAAGCAGGGGGC<br>CAGGGTGGGGGGGCCTTCTTCAGCCCAAGCCCTGGCAGCAGCAGCCTGTCCTCGTGGAGCTTCTTCTCCGATGCCTCTGA<br>TGAGGCAGCCCTGTATGCAGCCTGCGACGAGGTGGAGTCTGAGCTAAATGAGGCGGCCTCCCGCTTTGGCCTGGGCTCCC<br>CGCTGCCCTCGCCCCGGGCCTCCCCTCGGCCATGGACCCCCGAAGATCCCTGGAGCCTGTATGGTCCAAGCCCCGGAGGC<br>CGAGGGCCAGAGGATAGCTGGCTACTCCTCAGTGCTCCTGGGCCCACCCCAGCCTCCCCGCGGCCTGCCTCTCCATGTGG<br>CAAGCGGCGCTATTCCAGCTCGGGAACCCCATCTTCAGCCTCCCCAGCTCTGTCCCGCCGTGGCAGCCTGGGGGAAGAGG<br>GGTCTGAGCCACCTCCACCACCCCATTGCCTCTGGCCCGGGACCCGGGCTCCCCTGGTCCCTTTGACTATGTGGGGGCC<br>CCACCAGCTGAGAGCATCCCTCAGAAGACACGGCGGACTTCCAGCGAGCAGGCAGTGGCTCTGCCTCGGTCTGAGGAGCC<br>TGCCTCATGCAATGGGAAGCTGCCCTTGGGAGCAGAGGAGTCTGTGGCTCCTCCAGGAGGTTCCCGGAAGGAGGTGGCTG<br>GCATGGACTACCTGGCAGTGCCCTCCCCACTCGCTTGGTCCAAGGCCCGGATTGGGGGACACAGCCCTATCTTCAGGACC<br>TCTGCCCTACCCCCACTGGACTGGCCTCTGCCCAGCCAATATGAGCAGCTGGAGCTGAGGATCGAGGTACAGCCTAGAGC<br>CCACCACCGGGCCCACTATGAGACAGAAGGCAGCCGTGGAGCTGTCAAAGCTGCCCCTGGCGGTCACCCCGTAGTCAAGC<br>TCCTAGGCTACAGTGAGAAGCCACTGACCCTACAGATGTTCATCGGCACTGCAGATGAAAGGAACCTGCGGCCTCATGCC<br>TTCTATCAGGTGCACCGTATCACAGGCAAGATGGTGGCCACGGCCAGCTATGAAGCCGTAGTCAGTGGCACCAAGGTGTT<br>GGAGATGACTCTGCTGCCTGAGAACAACATGGCGGCCAACATTGACTGCGCGGGAATCCTGAAGCTTCGGAATTCAGACA<br>TTGAGCTTCGGAAGGGTGAGACGGACATCGGGCGCAAAAACACACGTGTACGGCTGGTGTTCCGGGTACACGTGCCCCAG<br>GGCGGCGGGAAGGTCGTCTCAGTACAGGCAGCATCGGTGCCCATCGAGTGCTCCCAGCGCTCAGCCCAGGAGCTGCCCCA<br>GGTGGAGGCCTACAGCCCCAGTGCCTGCTCTGTGAGAGGAGGCGAGGAACTGGTACTGACTGGCTCCAACTTCCTGCCAG<br>ACTCCAAGGTGGTGTTCATTGAGAGGGGTCCTGATGGGAAGCTGCAATGGGAGGAGGAGGCCACAGTGAACCGACTGCAG<br>AGCAACGAGGTGACGCTGACCCTGACTGTCCCCGAGTACAGCAACAAGAGGGTTTCCCGGCCAGTCCAGGTCTACTTTTA<br>TGTCTCCAATGGGCGGAGGAAACGCAGTCCTACCCAGAGTTTCAGGTTTCTGCCTGTGATCTGCAAAGAGGAGCCCCTAC<br>CGGACTCATCTCTGCGGGGTTTCCCTTCAGCATCGGCAACCCCCTTTGGCACTGACATGGACTTCTCACCACCCAGGCCC<br>CCCTACCCCTCCTATCGCCATGAAGACCCTGCTTGCGAAACTCCTTACCTATCAGAAGGCTTCGGCTATGGCATGCCCCC |
| NFATC3 | pDEST-AD | JBP005 | JBP006 | 58 | ura | ATGACTACTGCAAACTGTGGCGCCCACGACGAGCTCGACTTCAAACTCGTCTTTGGCGAGGACGGGGCGCCGGCGCCGCC<br>GCCCCCGGGCTCGCGGCCTGCAGATCTTGAGCCAGATGATTGTGCATCCATTTACATCTTTAATGTAGATCCACCTCCAT<br>CTACTTTAACCACACCACTTTGCTTACCACATCATGGATTACCGTCTCACTCTTCTGTTTTGTCACCATCGTTTCAGCTC<br>CAAAGTCACAAAAACTATGAAGGAACTTGTGAGATTCCTGAATCTAAATATAGCCCATTAGGTGGTCCCAAACCCTTTGA<br>GTGCCCAAGTATTCAAATTACATCTATCTCTCCTAACTGTCATCAAGAATTAGATGCACATGAAGATGACCTACAGATAA<br>ATGACCCAGAACGGGAATTTTTGGAAAGGCCTTCTAGAGATCATCTCTATCTTCCTCTTGAGCCATCCTACCGGGAGTCT<br>TCTCTTAGTCCTAGTCCTGCCAGCAGCATCTCTTCTAGGAGTTGGTTCTCTGATGCATCTTCTTGTGAATCGCTTTCACA<br>TATTTATGATGATGTGGACTCAGAGTTGAATGAAGCTGCAGCCCGATTTACCCTTGGATCCCCTCTGACTTCTCCTGGTG<br>GCTCTCCAGGGGGCTGCCCTGGAGAAGAAACTTGGCATCAACAGTATGGACTTGGACACTCATTATCACCCAGGCAATCT<br>CCTTGCCACTCTCCTAGATCCAGTGTCACTGATGAGAATTGGCTGAGCCCCAGGCCAGCCTCAGGACCCTCATCAAGGCC<br>CACATCCCCCTGTGGGAAACGGAGGCACTCCAGTGCTGAAGTTTGTTATGCTGGGTCCCTTTCACCCCATCACTCACCTG<br>TTCCTTCACCTGGTCACTCCCCCAGGGGAAGTGTGACAGAAGATACGTGGCTCAATGCTTCTGTCCATGGTGGGTCAGGC<br>CTTGGCCCTGCAGTTTTTCCATTTCAGTACTGTGTAGAGACTGACATCCCTCTCAAAACAAGGAAAACTTCTGAAGATCA<br>AGCTGCCATACTACCAGGAAAATTAGAGCTGTGTTCAGATGACCAAGGGAGTTTATCACCAGCCCGGGAGACTTCAATAG<br>ATGATGGCCTTGGATCTCAGTATCCTTTAAAGAAAGATTCATGTGGTGATCAGTTTCTTTCAGTTCCTTCACCCTTTACC<br>TGGAGCAAACCAAAGCCTGGCCACACCCCTATATTTCGCACATCTTCATTACCTCCACTAGACTGGCCTTTACCAGCTCA<br>TTTTGGACAATGTGAACTGAAAATAGAAGTGCAACCTAAAACTCATCATCGAGCCCATTATGAAACTGAAGGTAGCCGAG<br>GGGCAGTAAAAGCATCTACTGGGGGACATCCTGTTGTGAAGCTCCTGGGCTATAACGAAAAGCCAATAAATCTACAAATG<br>TTTATTGGGACAGCAGATGATCGATATTTACGACCTCATGCATTTTACCAGGTGCATCGAATCACTGGGAAGACAGTCGC<br>TACTGCAAGCCAAGAGATAATAATTGCCAGTACAAAAGTTCTGGAAATTCCACTTCTTCCTGAAAATAATATGTCAGCCA<br>GTATTGATTGTGCAGGTATTTTGAAACTCCGCAATTCAGATATAGAACTTCGAAAAGGAGAAACTGATATTGGCAGAAAG<br>AATACTAGAGTACGACTTGTGTTTCGTGTACACATCCCACAGCCCAGTGGAAAAGTCCTTTCTCTGCAGATAGCCTCTAT<br>ACCCGTTGAGTGCTCCCAGCGGTCTGCTCAAGAACTTCCTCATATTGAGAAGTACAGTATCAACAGTTGTTCTGTAAATG<br>GAGGTCATGAAATGGTTGTGACTGGATCTAATTTTCTTCCAGAATCCAAAATCATTTTTCTTGAAAAAGGACAAGATGGA<br>CGACCTCAGTGGGAGGTAGAAGGGAAGATAATCAGGGAAAAATGTCAAGGGGCTCACATTGTCCTTGAAGTTCCTCCATA<br>TCATAACCCAGCAGTTACAGCTGCAGTGCAGGTGCACTTTTATCTTTGCAATGGCAAGAGGAAAAAAAGCCAGTCTCAAC<br>GTTTTACTTATACACCAGTTTTGATGAAGCAAGAACACAGAGAAGAGATTGATTTGTCTTCAGTTCCATCTTTGCCTGTG<br>CCTCATCCTGCTCAGACCCAGAGGCCTTCCTCTGATTCAGGGTGTTCACATGACAGTGTACTGTCAGGACAGAGAAGTTT |
| SOX15 | ORFeome | JBP017_SOX15_F | JBP045_attL2_pDONR 221_201 | 58 | ura | ATGGCGCTACCAGGCTCCTCACAGGACCAGGCCTGGAGCCTGGAGCCTCCGGCTGCCACGGCTGCTGCCTCCTCATCTTC<br>GGGACCCCAGGAGCGGGAGGGCGCTGGGAGCCCCGCGGCCCCCGGGACGCTGCCCCTGGAGAAGGTGAAGCGGCCGATGA<br>ACGCGTTCATGGTGTGGAGCTCCGCTCAGCGCCGCCAGATGGCGCAGCAGAACCCCAAGATGCACAACTCCGAGATCTCC<br>AAGCGCCTGGGCGCGCAGTGGAAGCTGCTGGACGAGGACGAGAAGCGGCCCTTCGTGGAGGAGGCCAAGCGGCTCCGCGC<br>CCGACACCTGCGCGACTACCCCGACTACAAGTACCGGCCTCGGCGCAAGGCCAAGAGCTCGGGCGCCGGACCTTCCCGCT<br>GCGGACAGGGAAGAGGCAACCTGGCCAGCGGCGGCCCGCTCTGGGGGCCGGGGTACGCGACCACCCAACCGAGCAGAGGC<br>TTTGGGTACAGACCCCCCAGCTACTCGACAGCCTACCTGCCTGGCAGCTATGGCTCTTCCCACTGCAAACTGGAAGCCCC<br>CTCACCGTGCTCCCTCCCTCAGAGTGACCCTAGGCTCCAGGGGGAACTGCTGCCCACCTATACCCACTACCTGCCCCCTG<br>GCTCTCCCACTCCATACAACCCTCCCCTTGCTGGTGCCCCCATGCCCCTAACCCACCTCTTG |
| SOX17 | ORFeome | JBP018_SOX17_F | JBP043_attL2_pENTR 223.1 | 58 | ura | ATGAGCAGCCCGGATGCGGGATACGCCAGTGACGACCAGAGCCAGACCCAGAGCGCGCTGCCCGCGGTGATGGCCGGGCT<br>GGGCCCCTGCCCCTGGGCCGAGTCGCTGAGCCCCATCGGGGACATGAAGGTGAAGGGCGAGGTGCCGGCGAACAGCGGAG<br>CACCGGCCGGGGCCGCGGGCCGAGCCAAGGGCGAGTCCCGTATCCGGCGGCCGATGAACGCTTTCATGGTGTGGGCTAAG<br>GACGAGCGCAAGCGGCTGGCGCAGCAGAATCCAGACCTGCACAACGCCGAGTTGAGCAAGATGCTGGGCAAGTCGTGGAA<br>GGCGCTGACGCTGGCGGAGAAGCGGCCCTTCGTGGAGGAGGCAGAGCGGCTGCGCGTGCAGCACATGCAGGACCACCCCA<br>ACTACAAGTACCGGCCGCGGCGGCGCAAGCAGGTGAAGCGGCTGAAGCGGGTGGAGGGCGGCTTCCTGCACGGCCTGGCT<br>GAGCCGCAGGCGGCCGCGCTGGGCCCCGAGGGCGGCCGCGTGGCCATGGACGGCCTGGGCCTCCAGTTCCCCGAGCAGGG<br>CTTCCCCGCCGGCCCGCCGCTGCTGCCTCCGCACATGGGCGGCCACTACCGCGACTGCCAGAGTCTGGGCGCGCCTCCGC<br>TCGACGGCTACCCGTTGCCCACGCCCGACACGTCCCCGCTGGACGGCGTGGACCCCGACCCGGCTTTCTTCGCCGCCCCG<br>ATGCCCGGGGACTGCCCGGCGGCCGGCACCTACAGCTACGCGCAGGTCTCGGACTACGCTGGCCCCCCGGAGCCTCCCGC<br>CGGTCCCATGCACCCCCGACTCGGCCCAGAGCCCGCGGGTCCCTCGATTCCGGGCCTCCTGGCGCCACCCAGCGCCCTTC<br>ACGTGTACTACGGCGCGATGGGCTCGCCCGGGGCGGGCGGCGGGCGCGGCTTCCAGATGCAGCCGCAACACCAGCACCAG<br>CACCAGCACCAGCACCACCCCCCGGGCCCCGGACAGCCGTCGCCCCCTCCGGAGGCACTGCCCTGCCGGGACGGCACGGA<br>CCCCAGTCAGCCCGCCGAGCTCCTCGGGGAGGTGGACCGCACGGAATTTGAACAGTATCTGCACTTCGTGTGCAAGCCTG<br>AGATGGGCCTCCCCTACCAGGGGCATGACTCCGGTGTGAATCTCCCCGACAGCCACGGGGCCATTTCCTCGGTGGTGTCC<br>GACGCCAGCTCCGCGGTATATTACTGCAACTATCCTGACGTG |

| | | | | | | |
|---|---|---|---|---|---|---|
| SOX7 | pDEST-AD | JBP005 | JBP006 | 58 | ura | ATGGCTTCGCTGCTGGGAGCCTACCCTTGGCCCGAGGGTCTCGAGTGCCCGGCCCTGGACGCCGAGCTGTCGGATGGACA<br>ATCGCCGCCGGCCGTCCCCCGGCCCCCGGGGGACAAGGGCTCCGAGAGCCGTATCCGGCGGCCCATGAACGCCTTCATGG<br>TTTGGGCCAAGGACGAGAGGAAACGGCTGGCAGTGCAGAACCCGGACCTGCACAACGCCGAGCTCAGCAAGATGCTGGGA<br>AAGTCGTGGAAGGCGCTGACGCTGTCCCAGAAGAGGCCGTACGTGGACGAGGCGGAGCGGCTGCGCCTGCAGCACATGCA<br>GGACTACCCCAACTACAAGTACCGGCCGCGCAGGAAGAAGCAGGCCAAGCGGCTGTGCAAGCGCGTGGACCCGGGCTTCC<br>TTCTGAGCTCCCTCTCCCGGGACCAGAACGCCCTGCCGGAGAAGAGAAGCGGCAGCCGGGGGGCGCTGGGGGAGAAGGAG<br>GACAGGGGTGAGTACTCCCCCGGCACTGCCCTGCCCAGCCTCCGGGGCTGCTACCACGAGGGGCCGGCTGGTGGTGGCGG<br>CGGCGGCACCCCGAGCAGTGTGGACACGTACCCGTACGGGCTGCCCACACCTCCTGAAATGTCTCCCCTGGACGTGCTGG<br>AGCCGGAGCAGACCTTCTTCTCCTCCCCCTGCCAGGAGGAGCATGGCCATCCCCGCCGCATCCCCCACCTGCCAGGGCAC<br>CCGTACTCACCGGAGTACGCCCCAAGCCCTCTCCACTGTAGCCACCCCCTGGGCTCCCTGGCCCTTGGCCAGTCCCCCGG<br>CGTCTCCATGATGTCCCCTGTACCCGGCTGTCCCCCATCTCCTGCCTATTACTCCCCGGCCACCTACCACCCACTCCACT<br>CCAACCTCCAAGCCCACCTGGGCCAGCTTTCCCCGCCTCCTGAGCACCCTGGCTTCGACGCCCTGGATCAACTGAGCCAG<br>GTGGAACTCCTGGGGGACATGGATCGCAATGAATTCGACCAGTATTTGAACACTCCTGGCCACCCAGACTCCGCCACAGG<br>GGCCATGGCCCTCAGTGGGCATGTTCCGGTCTCCCAGGTGACACCAACGGGTCCCACAGAGACCAGCCTCATCTCCGTCC<br>TGGCTGATGCCACGGCCACGTACTACAACAGCTACAGTGTGTCATAG |
| SOX11 | ORFeome | JBP038_SOX11_F | JBP045_attL2_pDONR221_201 | 58 | ura | ATGGTGCAGCAGGCGGAGAGCTTGGAAGCGGAGAGCAACCTGCCCCGGGAGGCGCTGGACACGGAGGAGGGCGAATTCAT<br>GGCTTGCAGCCCGGTGGCCCTGGACGAGAGCGACCCAGACTGGTGCAAGACGGCGTCGGGCCACATCAAGCGGCCGATGA<br>ACGCGTTCATGGTATGGTCCAAGATCGAACGCAGGAAGATCATGGAGCAGTCTCCGGACATGCACAACGCCGAGATCTCC<br>AAGAGGCTGGGCAAGCGCTGGAAAATGCTGAAGGACAGCGAGAAGATCCCGTTCATCCGGGAGGCGGAGCGGCTGCGGCT<br>CAAGCACATGGCCGACTACCCCGACTACAAGTACCGGCCCCGGAAAAAGCCCAAAATGGACCCCTCGGCCAAGCCCAGCG<br>CCAGCCAGAGCCCAGAGAAGAGCGCGGCCGGCGGCGGCGGCGGGAGCGCGGGCGGAGGCGCGGGCGGTGCCAAGACCTCC<br>AAGGGCTCCAGCAAGAAATGCGGCAAGCTCAAGGCCCCCGCGGCCGCGGGCGCCAAGGCGGGCGCGGGCAAGGCGGCCCA<br>GTCCGGGGACTACGGGGGCGCGGGCGACGACTACGTGCTGGGCAGCCTGCGCGTGAGCGGCTCGGGCGGCGGCGGCGCGG<br>GCAAGACGGTCAAGTGCGTGTTTCTGGATGAGGACGACGACGACGACGACGACGACGAGCTGCAGCTGCAGATCAAA<br>CAGGAGCCGGACGAGGAGGACGAGGAACCACCGCACCAGCAGCTCCTGCAGCCGCCGGGGCAGCAGCCGTCGCAGCTGCT<br>GAGACGCTACAACGTCGCCAAAGTGCCCGCCAGCCCTACGCTGAGCAGCTCGGCGGAGTCCCCCGAGGGAGCGAGCCTCT<br>ACGACGAGGTGCGGGCCGGCGCGACCTCGGGCGCCGGGGGCGGCAGCCGCCTCTACTACAGCTTCAAGAACATCACCAAG<br>CAGCACCCGCCGCCGCTCGCGCAGCCCGCGCTGTCGCCCGCGTCCTCGCGCTCGGTGTCCACCTCCTCGTCCAGCAGCAG<br>CGGCAGCAGCAGCGGCAGCAGCGGCGAGGACGCCGACGACCTGATGTTCGACCTGAGCTTGAATTTCTCTCAAAGCGCGC<br>ACAGCGCCAGCGAGCAGCAGCTGGGGGGCGGCGCGGCGGCCGGGAACCTGTCCCTGTCGCTGGTGGATAAGGATTTGGAT<br>TCGTTCAGCGAGGGCAGCCTGGGCTCCCACTTCGAGTTCCCCGACTACTGCACGCCGGAGCTGAGCGAGATGATCGCGGG<br>GGACTGGCTGGAGGCGAACTTCTCCGACCTGGTGTTCACATAT |
| SOX4 | ORFeome | JBP039_SOX4_F | JBP045_attL2_pDONR221_201 | 58 | ura | ATGGTGCAGCAAACCAACAATGCCGAAAACACGGAGGCACTGCTCGCAGGTGAAAGCTCCGATAGCGGTGCAGGCCTTGA<br>GTTGGGGATCGCTTCTAGTCCCACCCCTGGAAGTACCGCCTCAACAGGCGGGAAGGCTGACGATCCATCCTGGTGCAAAA<br>CACCGTCTGGGCACATTAAGCGCCCCATGAACGCCTTCATGGTGTGGAGCCAGATAGAACGGAGAAAAATCATGGAGCAA<br>TCTCCTGACATGCATAATGCCGAGATCAGCAAGCGGCTGGGCAAGAGATGGAAACTGCTGAAGGACAGCGATAAAATCCC<br>CTTCATCCGTGAAGCAGAGCGGCTGCGGCTGAAGCACATGGCTGATTACCCCGACTACAAATATCGACCTCGGAAGAAGG<br>TTAAGTCTGGCAACGCCAATTCATCGAGCTCCGCCGCTGCATCTAGTAAGCCTGGAGAGAAAGGCGATAAGGTCGGCGGT<br>TCCGGTGGTGGAGGTCATGGTGGCGGGGGTGGTGGTGGATCATCGAACGCCGGCGGTGGTGGAGGTGGTGCTAGTGGGGG<br>AGGTGCAAATTCCAAACCAGCCCAGAAGAAAAGCTGTGGTTCCAAGGTGGCTGGTGGTGCAGGAGGTGGCGTCTCCAAAC<br>CACACGCTAAGCTGATCCTGGCAGGCGGCGGTGGTGGTGGAAAAGCAGCAGCTGCAGCAGCAGCTTCTTTCGCAGCAGAA<br>CAGGCAGGTGCAGCCGCCCTGCTACCTCTGGGTGCAGCAGCTGACCACCATAGCCTCTACAAGGCCAGAACTCCATCTGC<br>AAGCGCCTCAGCAAGCTCCGCAGCAAGCGCCAGCGCCGCCCTGGCAGCGCCTGGGAAACACCTGGCTGAGAAGAAAGTGA<br>AGCGGGTGTACCTGTTTGGTGGCCTTGGAACATCTAGTTCACCAGTTGGTGGAGTAGGTGCAGGAGCTGATCCAAGCGAC<br>CCTCTTGGCTTGTATGAGGAAGAGGGTGCAGGATGCTCCCCAGATGCACCATCTCTCAGTGGCCGGAGCAGCGCCGCTTC<br>CTCTCCAGCAGCAGGAAGGAGCCCAGCTGACCATCGTGGTTATGCATCACTGCGAGCAGCTAGCCCAGCACCAAGTTCAG<br>CTCCATCCCACGCATCGAGCTCCGCCTCTAGTCATTCATCGAGCTCCTCTAGTTCAGGAAGCTCCTCTAGTGACGATGAA<br>TTCGAGGACGATCTGCTCGATCTGAACCCCTCAAGCAATTTCGAATCAATGAGCCTGGGCTCCTTTTCCTCTAGTAGCGC<br>CCTGGACAGAGATCTGGACTTCAACTTTGAACCTGGCTCCGGGTCTCACTTCGAGTTTCCCGATTACTGTACTCCTGAGG<br>TGAGCGAGATGATTAGTGGCGACTGGCTCGAGAGCTCCATAAGCAACCTGGTGTTCACCTACCTGTTG |
| SOX6 | pDEST-AD | JBP005 | JBP006 | 58 | ura | ATGAAAGGTCATGGTGAGCTTCAAGGACATGAAAGAAGAATGTCTTCCAAGCAAGCCACCTCTCCATTTGCCTGTGCAGC<br>TGATGGAGAGGATGCAATGACCCAGGATTTAACCTCAAGGGAAAAGGAAGAGGGCAGTGATCAACATGTGGCCTCCCATC<br>TGCCTCTGCACCCCATAATGCACAACAAACCTCACTCTGAGGAGCTACCAACACTTGTCAGTACCATTCAACAAGATGCT<br>GACTGGGACAGCGTTCTGTCATCTCAGCAAAGAATGGAATCAGAGAATAATAAGTTATGTTCCCTATATTCCTTCCGAAA<br>TACCTCTACCTCACCACATAAGCCTGACGAAGGGAGTCGGGACCGTGAGATAATGACCAGTGTTACTTTTGGAACCCCAG<br>AGCGCCGCAAAGGGAGTCTTGCCGATGTGGTGGACACACTGAAACAGAAGAAGCTTGAGGAAATGACTCGGACTGAACAA<br>GAGGATTCCTCCTGCATGGAAAAACTACTTTCAAAAGATTGGAAGGAAAAAATGGAAAGACTAAATACCAGTGAACTTCT<br>TGGAGAAATTAAAGGTACACCTGAGAGCCTGGCAGAAAAAGAACGGCAGCTCTCCACCATGATTACCCAGCTGATCAGTT<br>TACGGGAGCAGCTACTGGCAGCGCATGATGAACAGAAAAAACTGGCAGCGTCACAAATTGAGAAACAACGGCAGCAAATG<br>GACCTTGCTCGCCAACAGCAAGAACAGATTGCGAGACAACAGCAGCAACTTCTGCAACAGCAGCACAAAATTAATCTCCT<br>GCAGCAACAGATCCAGGTTCAGGGTCACATGCCTCCGCTCATGATCCCAATTTTTCCACATGACCAGCGGACCCTGGCAG<br>CAGCTGCTGCTGCCCAACAGGGATTCCTCTTCCCCCCTGGAATAACATACAAACCAGGTGATAACTACCCCGTACAGTTC<br>ATTCCATCAACAATGGCAGCTGCTGCTGCTTCTGGACTCAGCCCTTTACAGCTCCAGAAGGGTCATGTCTCCCACCCACA<br>AATTAACCAAAGGCTAAAGGGCCTAAGTGACCGTTTTGGCAGGAATTTGGACACCTTTGAACATGGTGGTGGCCACTCTT<br>ACAACCACAAACAGATTGAGCAGCTCTATGCCGCTCAGCTGGCCAGCATGCAGGTGTCACCTGGAGCAAAGATGCCATCA<br>ACTCCACAGCCACCAAACACAGCAGGGACGGTCTCACCTACTGGGATAAAAAATGAAAAGAGAGGGACCAGCCCTGTAAC<br>TCAAGTTAAGGATGAAGCAGCAGCACAGCCTCTGAATCTCTCATCCCGACCCAAGACAGCAGAGCCTGTAAAGTCCCCAA<br>CGTCTCCCACCCAGAACCTCTTCCCAGCCAGCAAAACCAGCCCTGTCAATCTGCCAAACAAAAGCAGCATCCCTAGCCCC<br>ATTGGAGGAAGCCTGGGAAGAGGATCCTCTTTAGATATCCTATCTAGTCTCAACTCCCCTGCCCTTTTTGGGGATCAGGA<br>TACAGTGATGAAAGCCATTCAGGAGGCGCGGAAGATGCGAGAGCAGATCCAGCGGGAGCAACAGCAGCAACAGCCACATG<br>GTGTTGACGGGAAACTGTCCTCCATAAATAATATGGGGCTGAACAGCTGCAGGAATGAAAAGGAAAGAACGCGCTTTGAG<br>AATTTGGGGCCCCAGTTAACGGGAAAGTCAAATGAAGATGGAAAACTGGGCCCAGGTGTCATCGACCTTACTCGGCCAGA<br>AGATGCAGAGGGAAGTAAAGCAATGAATGGCTCTGCAGCTAAACTACAGCAGTATTATTGTTGGCCAACAGGAGGTGCCA<br>CTGTGGCTGAAGCACGAGTCTACAGGGACGCCCGCGGCCGTGCCAGCAGCGAGCCACACATTAAGCGACCAATGAATGCA<br>TTCATGGTTTGGGCAAAGGATGAGAGGAGAAAAATCCTTCAGGCCTTCCCCGACATGCATAACTCCAACATTAGCAAAAT<br>CTTAGGATCTCGCTGGAAATCAATGTCCAACCAGGAGAAGCAACCTTATTATGAAGAGCAGGCCCGGCTAAGCAAGATCC<br>ACTTAGAGAAGTACCCAAACTATAAATACAAACCCCGACCGAAACGCACCTGCATTGTTGATGGCAAAAAGCTTCGGATT<br>GGGGAGTATAAGCAACTGATGAGGTCTCGGAGACAGGAGATGAGGCAGTTCTTTACTGTGGGGCAACAGCCTCAGATTCC |
| SOX30 | pDEST-AD | JBP005 | JBP006 | 58 | ura | ATGGAGAGAGCCAGACCCGAGCCGCCGCCTCAGCCGCGCCCGTTGCGTCCCGCTCCGCCCCCGCTGCCGGTCGAGGGCAC<br>CTCCTTTTGGGCAGCAGCCATGGAGCCCCCTCCGTCGTCTCCCACACTGAGCGCGGCAGCCAGTGCGACCTTGGCCTCGT<br>CGTGCGGGGAGGCAGTGGCGTCCGGCTTACAGCCCGCGGTGCGGCGGCTGCTGCAGGTGAAGCCAGAGCAGGTGTTGCTG<br>CTACCACAGCCTCAGGCCCAGAACGAGGAAGCCGCTGCCTCGTCCGCGCAGGCGCGGCTGTTGCAGTTCAGGCCCGACCT<br>GCGGCTCCTGCAGCCGCCGACAGCGTCAGACGGCGCCACCTCCAGGCCCGAGTTGCACCCGGTGCAGCCCCTGGCGCTGC<br>ATGTCAAGGCCAAGAAGCAGAAGCTGGGGCCCAGCCTGGATCAGTCAGTGGGGCCTCGAGGGGCCGTCGAAACCGGTCCT<br>AGAGCCTCCAGGGTGGTCAAGTTGGAAGGCCCCGGGCCGGCCCTCGGCTACTTCCGAGGGGACGAGAAGGGCAAGCTGGA<br>GGCGGAGGAGGTCATGAGAGACTCGATGCAAGGCGGGGCAGGCAAAAGCCCGGCAGCCATCCGAGAAGGTGTGATCAAAA<br>CGGAGGAACCCGAGAGACTCCTCGAGGACTGCAGGCTCGGCGCGGAGCCCGCGTCCAATGGCCTGGTTCATGGCAGCGCG<br>GAGGTCATCTTGGCCCCAACGTCCGGTGCCTTTGGGCCGCACCAGCAAGACCTTAGGATCCCTTTGACGCTCCACACGGT<br>CCCCCCTGGGGCCCGGATCCAGTTTCAGGGAGCTCCGCCTTCAGAGCTGATAAGATTGACCAAGGTCCCCCTGACACCAG<br>TGCCTACTAAAATGCAGTCCCTACTGGAGCCTTCTGTAAAAATTGAAACCAAAGATGTCCCGCTCACCGTGTTGCCCTCA<br>GATGCAGGCATACCAGATACTCCCTTCAGTAAGGACAGAAATGGTCATGTGAAGCGACCCATGAACGCATTTATGGTTTG<br>GGCAAGGATCCACCGACCAGCACTAGCCAAAGCTAACCCAGCAGCCAACAATGCAGAAATCAGTGTCCAGCTTGGGTTAG<br>AGTGGAACAAACTTAGTGAAGAACAAAAGAAACCCTATTACGATGAAGCACAAAAGATTAAGGAAAAGCACAGAGAGGAA<br>TTTCCTGGTTGGGTTTATCAGCCTCGTCCAGGGAAGCGAAAACGATTCCCTCTAAGTGTTTCCAATGTATTTTCTGGTAC<br>CACACAGAATATCATCTCTACAAATCCTACAACAGTTTATCCTTACCGCTCACCTACGTACTCTGTGGTAATTCCCAGCC<br>TACAGAATCCCATCACTCATCCAGTTGTACATGCCCTTACAGTCGGCCTCCCTTTGGCTATGGAAATTTTCCGAGTTCAA<br>TGCCAGAATGCCTTAGTTATTATGAAGACAGGTACCCAAAACATGAGGGTATCTTTTCAACTTTAG |
| SOX9 | pDEST-AD | JBP005 | JBP006 | 58 | ura | ATGAATCTCCTGGACCCCTTCATGAAGATGACCGACGAGCAGGAGAAGGGCCTGTCCGGCGCCCCCAGCCCCACCATGTC<br>CGAGGACTCCGCGGGCTCGCCCTGCCCGTCGGGCTCCGGCTCGGACACCGAGAACACGCGGCCCCAGGAGAACACGTTCC<br>CCAAGGGCGAGCCCGATCTGAAGAAGGAGAGCGAGGAGGACAAGTTCCCCGTGTGCATCCGCGAGGCGGTCAGCCAGGTG<br>CTCAAAGGCTACGACTGGACGCTGGTGCCCATGCCGGTGCGCGTCAACGGCTCCAGCAAGAACAAGCCGCACGTCAAGCG<br>GCCCATGAACGCCTTCATGGTGTGGGCGCAGGCGGCGCGCAGGAAGCTCGCGGACCAGTACCCGCACTTGCACAACGCCG<br>AGCTCAGCAAGACGCTGGGCAAGCTCTGGAGACTTCTGAACGAGAGCGAGAAGCGGCCCTTCGTGGAGGAGGCGGAGCGG<br>CTGCGCGTGCAGCACAAGAAGGACCACCCGGATTACAAGTACCAGCCGCGGCGGAGGAAGTCGGTGAAGAACGGGCAGGC<br>GGAGGCAGAGGAGGCCACGGAGCAGACGCACATCTCCCCCAACGCCATCTTCAAGGCGCTGCAGGCCGACTCGCCACACT<br>CCTCCTCCGGCATGAGCGAGGTGCACTCCCCCGGCGAGCACTCGGGGCAATCCCAGGGCCCACCGACCCCACCCACCACC<br>CCCAAAACCGACGTGCAGCCGGGCAAGGCTGACCTGAAGCGAGAGGGGCGCCCCTTGCCAGAGGGGGGCAGACAGCCCCC<br>TATCGACTTCCGCGACGTGGACATCGGCGAGCTGAGCAGCGACGTCATCTCCAACATCGAGACCTTCGATGTCAACGAGT<br>TTGACCAGTACCTGCCGCCCAACGGCCACCCGGGGGTGCCGGCCACGCACGGCCAGGTCACCTACACGGGCAGCTACGGC<br>ATCAGCAGCACCGCGGCCACCCCGGCGAGCGCGGGCCACGTGTGGATGTCCAAGCAGCAGGCGCCGCCGCCACCCCCGCA<br>GCAGCCCCCACAGGCCCCGCCGGCCCCGCAGGCGCCCCCGCAGCCGCAGGCGGCGCCCCCACAGCAGCCGGCGGCACCCC<br>CGCAGCAGCCACAGGCGCACACGCTGACCACGCTGAGCAGCGAGCCGGGCCAGTCCCAGCGAACGCACATCAAGACGGAG<br>CAGCTGAGCCCCAGCCACTACAGCGAGCAGCAGCAGCACTCGCCCCAACAGATCGCCTACAGCCCCTTCAACCTCCCACA<br>CTACAGCCCCTCCTACCCGCCCATCACCCGCTCACAGTACGACTACACCGACCACCAGAACTCCAGCTCCTACTACAGCC<br>ACGCGGCAGGCCAGGGCACCGGCCTCTACTCCACCTTCACCTACATGAACCCCGCTCAGCGCCCCATGTACACCCCCATC<br>GCCGACACCTCTGGGGTCCCTTCCATCCCGCAGACCCACAGCCCCCAGCACTGGGAACAACCCGTCTACACACAGCTCAC<br>TCGACCTTAG |

| | | | | | | |
|---|---|---|---|---|---|---|
| SOX13 | ORFeome | JBP026_SOX13_F | JBP045_attL2_pDONR221_201 | 58 | ura | ATGTCCATGAGGAGCCCCATCTCTGCCCAGCTGGCCCTGGATGGCGTTGGCACCATGGTGAACTGCACCATCAAGTCAGA<br>GGAGAAGAAAGAGCCTTGCCACGAGGCCCCCAGGGCTCAGCCACTGCCGCTGAACCTCAGCCTGGAGACCCAGCCCGGG<br>CCTCCCAGGATAGTGCTGACCCCCAAGCTCCAGCCCAGGGGAATTTCAGGGGCTCCTGGGACTGTAGCTCTCCAGAGGGT<br>AATGGGTCCCCAGAACCCAAGAGACCAGGAGTGTCGGAGGCTGCCTCTGGAAGCCAGGAGAAGCTGGACTTCAACCGAAA<br>TTTGAAAGAAGTGGTGCCAGCCATAGAGAAGCTGTTGTCCAGTGACTGGAAGGAGAGGTTTCTAGGAAGGAACTCTATGG<br>AAGCCAAAGATGTCAAAGGGACCCAAGAGAGCCTAGCAGAGAAGGAGCTCCAGCTTCTGGTCATGATTCACCAGCTGTCC<br>ACCCTGCGGGACCAGCTCCTGACAGCCCACTCGGAGCAGAAGAACATGGCTGCCATGCTGTTTGAGAAGCAGCAGCAGCA<br>GATGGAGCTTGCCCGGCAGCAGCAGGAGCAGATTGCAAAGCAGCAGCAGCAGCTGATTCAGCAGCAGCATAAGATCAACC<br>TCCTTCAGCAGCAGATCCAGCAGGTTAACATGCCTTATGTCATGATCCCAGCCTTCCCCCCAAGCCACCAACCTCTGCCT<br>GTCACCCCTGACTCCCAGCTGGCCTTACCCATTCAGCCCATTCCCTGCAAACCAGTGGAGTATCCGCTGCAGCTGCTGCA<br>CAGCCCCCCTGCCCCAGTGGTGAAGAGGCCTGGGGCCATGGCCACCCACCACCCCCTGCAGGAGCCCTCCCAGCCCCTGA<br>ACCTCACAGCCAAGCCCAAGGCCCCCGAGCTGCCCAACACCTCCAGCTCCCCAAGCCTGAAGATGAGCAGCTGTGTGCCC<br>CGCCCCCCCAGCCATGGAGGCCCCACGCGGGACCTGCAGTCCAGCCCCCCGAGCCTGCCTCTGGGCTTCCTTGGTGAAGG<br>GGACGCTGTCACCAAAGCCATCCAGGATGCTCGGCAGCTGCTGCACAGCCACAGTGGGGCCTTGGATGGCTCCCCCAACA<br>CCCCCTTCCGTAAGGACCTCATCAGCCTGGACTCATCCCCAGCCAAGGAGCGGCTGGAGGACGGCTGTGTGCACCCACTG<br>GAGGAAGCCATGCTGAGCTGCGACATGGATGGCTCCCGCCACTTCCCCGAGTCCCGAAACAGCAGCCACATCAAGAGGCC<br>CATGAACGCCTTCATGGTGTGGGCCAAGGATGAGCGGAGGAAGATCCTGCAAGCCTTCCCAGACATGCACAACTCCAGCA<br>TCAGCAAGATCCTTGGATCTCGCTGGAAGTCCATGACCAACCAGGAGAAGCAGCCCTACTATGAGGAACAGGCGCGGCTG<br>AGCCGGCAGCACCTGGAGAAGTATCCTGACTACAAGTACAAGCCGCGGCCCAAGCGCACCTGCATCGTGGAGGGCAAGCG<br>GCTGCGCGTGGGAGAGTACAAGGCCCTGATGAGGACCCGGCGTCAGGATGCCCGCCAGAGCTACGTGATCCCCCCGCAGG<br>CTGGCCAGGTGCAGATGAGCTCCTCAGATGTCCTGTACCCTCGGGCAGCAGGCATGCCGCTGGCACAGCCACTGGTGGAG<br>CACTATGTCCCTCGTAGCCTGGACCCCAACATGCCTGTGATCGTCAACACCTGCAGCCTCAGAGAGGAGGGTGAGGGCAC<br>AGATGACAGGCACTCGGTGGCTGATGGCGAGATGTACCGGTACAGCGAGGACGAGGACTCGGAGGGCGAAGAGAAGAGCG<br>ATGGGGAGTTGGTGGTGCTCACAGACTTG |
| SOX5 | ORFeome | JBP040_SOX5_F | JBP045_attL2_pDONR221_201 | 58 | ura | ATGCTTACTGACCCTGATTTACCTCAGGAGTTTGAAAGGATGTCTTCCAAGCGACCAGCCTCTCCGTATGGGGAAGCAGA<br>TGGAGAGGTAGCCATGGTGACAAGCAGACAGAAAGTGGAAGAAGAGGAGAGTGACGGGCTCCCAGCCTTTCACCTTCCCT<br>TGCATGTGAGTTTTCCCAACAAGCCTCACTCTGAGGAATTTCAGCCAGTTTCTCTGCTGACGCAAGAGACTTGTGGCCAT<br>AGGACTCCCACTTCTCAGCACAATACAATGGAAGTTGATGGCAATAAAGTTATGTCTTCATTTGCCCCACACAACTCATC<br>TACCTCACCTCAGAAGGCAGAAGAAGGTGGGCGACAGAGTGGCGAGTCCTTGTCTAGTACAGCCCTGGGAACTCCTGAAC<br>GGCGCAAGGGCAGTTTAGCTGATGTTGTTGACACCTTGAAGCAGAGGAAAATGGAAGAGCTCATCAAAAACGAGCCGGAA<br>GAAACCCCCAGTATTGAAAAACTACTCTCAAAGGACTGGAAAGACAAGCTTCTTGCAATGGGATCGGGGAACTTTGGCGA<br>AATAAAAGGGACTCCCGAGAGCTTAGCTGAGAAAGAAAGGCAACTCATGGGTATGATCAACCAGCTGACCAGCCTCCGAG<br>AGCAGCTGTTGGCTGCCCACGATGAGCAGAAGAAACTAGCTGCCTCTCAGATTGAGAAACAGCGTCAGCAAATGGAGCTG<br>GCCAAGCAGCAACAAGAACAAATTGCAAGACAGCAGCAGCAGCTTCTACAGCAACAACACAAAATCAATTTGCTCCAGCA<br>ACAGATCCAGGTTCAAGGTCAGCTGCCGCCATTAATGATTCCCGTATTCCCTCCTGATCAACGGACACTGGCTGCAGCTG<br>CCCAGCAAGGATTCCTCCTCCCTCCAGGCTTCAGCTATAAGGCTGGATGTAGTGACCCTTACCCTGTTCAGCTGATCCCA<br>ACTACCATGGCAGCTGCTGCCGCAGCAACACCAGGCTTAGGCCCACTCCAACTGCAGCAGTTATATGCTGCCCAGCTAGC<br>TGCAATGCAGGTATCTCCAGGAGGGAAGCTGCCAGGCATACCCCAAGGCAACCTTGGTGCTGCTGTATCTCCTACCAGCA<br>TTCACACAGACAAGAGCACAAACAGCCCACCACCCAAAAGCAAGGATGAAGTGGCACAGCCACTGAACCTATCAGCTAAA<br>CCCAAGACCTCTGATGGCAAATCACCCACATCACCCACCTCTCCCCATATGCCAGCTCTGAGAATAAACAGTGGGGCAGG<br>CCCCCTCAAAGCCTCTGTCCCAGCAGCGTTAGCTAGTCCTTCAGCCAGAGTTAGCACAATAGGTTACTTAAATGACCATG<br>ATGCTGTCACCAAGGCAATCCAAGAAGCTCGGCAAATGAAGGAGCAACTCCGACGGGAACAACAGGTGCTTGATGGGAAG<br>GTGGCTGTTGTGAATAGTCTGGGTCTTAATAACTGCCGAACAGAAAAGGAAAAAACAACACTGGAGAGTCTGACTCAGCA<br>ACTGGCAGTTAAACAGAATGAAGAAGGAAAATTTAGCCATGCAATGATGGATTTCAATCTGAGTGGAGATTCTGATGGAA<br>GTGCTGGAGTCTCAGAGTCAAGAATTTATAGGGAATCCCGAGGGCGTGGTAGCAATGAACCCCACATAAAGCGTCCAATG<br>AATGCCTTCATGGTGTGGGCTAAAGATGAACGGAGAAAGATCCTTCAAGCCTTTCCTGACATGCACAACTCCAACATCAG<br>CAAGATATTGGGATCTCGCTGGAAAGCTATGACAAACCTAGAGAAACAGCCATATTATGAGGAGCAAGCCCGTCTCAGCA<br>AGCAGCACCTGGAGAAGTACCCTGACTATAAGTACAAGCCCAGGCCAAAGCGCACATGCCTGGTGGATGGCAAAAAGCTG<br>CGCATTGGTGAATACAAGGCAATCATGCGCAACAGGCGGCAGGAAATGCGGCAGTACTTCAATGTTGGGCAACAAGCACA<br>GATCCCCATTGCCACTGCTGGTGTTGTGTACCCTGGAGCCATCGCCATGGCTGGGATGCCCTCCCCTCACCTACCCTCGG<br>AGCACTCAAGCGTGTCTAGCAGCCCAGAGCCTGGGATGCCTGTTATCCAGAGCACTTACGGTGTGAAAGGAGAGGAGCCA<br>CATATCAAAGAAGAGATACAGGCCGAGGACATCAATGGAGAAATTTATGATGAGTACGACGAGGAAGAGGATGATCCAGA |
| TGIF2LX | pDEST-AD | JBP005 | JBP006 | 58 | ura | ATGGAGGCCGCTGCGGACGGCCCGGCTGAGACCCAAAGCCCGGTGGAAAAAGACAGCCCGGCGAAGACCCAAAGCCCAGC<br>CCAAGACACCTCAATCATGTCGAGAAATAACGCAGATACAGGCAGAGTTCTTGCCTTACCAGAGCACAAGAAGAAGCGCA<br>AGGGAAACTTGCCAGCCGAGTCCGTTAAGATCCTCCGCGACTGGATGTATAAGCATCGGTTTAAGGCCTACCCTTCAGAA<br>GAAGAGAAGCAAATGCTGTCAGAGAAGACCAATTTGTCTTTGTTGCAGATTTCTAACTGGTTTATCAATGCTCGCAGACG<br>CATTCTCCCGGATATGCTTCAACAGCGTAGAAACGACCCCATCATTGGCCACAAAACGGGCAAAGATGCCCATGCCACCC<br>ACCTGCAGAGCACCGAGGCGTCTGTGCCGGCCAAGTCAGGGCCCAGTGGTCCAGACAATGTACAAAGCCTGCCCCTGTGG<br>CCCTTGCCAAAGGGCCAGATGTCAAGAGAGAAGCAACCAGATCCGGAGTCGGCCCCTAGCCAGAAGCTCACCGGAATAGC<br>CCAGCCGAAGAAAAAGGTCAAGGTTTCTGTCACATCCCCGTCTTCTCCAGAACTTGTGTCTCCAGAGGAGCACGCCGACT<br>TCAGCAGCTTCCTGCTGCTAGTCGATGCAGCAGTACAAAGGGCTGCCGAGCTGGAGCTAGAGAAGAAGCAAGAGCCTAAT<br>CCATAG |
| TGIF2LY | pDEST-AD | JBP005 | JBP006 | 58 | ura | ATGGAGGCCGCTGCAGACGGCCCGGCTGAGACCCAAAGCCCGGTGGAAAAAGACAGCCCGGCGAAGACCCAAAGCCCAGC<br>CCAAGACACCTCAATCATGTCGAGAAATAACGCAGATACAGGCAGAGTTCTTGCCTTACCAGAGCACAAGAAGAAGCGCA<br>AGGGAAACTTGCCAGCCGAGTCCGTTAAGATCCTCCGCGACTGGATGTATAAGCATCGGTTTAAGGCCTACCCTTCAGAA<br>GAAGAGAAGCAAATGCTGTCAGAGAAGACCAATTTGTCTTTGTTGCGGATTTCTAACTGGTTTATCAATGCTCGCAGACG<br>CATTCTCCCGGATATGCTTCAACAGCGTAGAAACGACCCCATCATTGGCCACAAAACGGGCAAAGATGCCCATGCCACCC<br>ACCTGCAGAGCACCGAGGCGTCTGTGCCGGCCAAGTCAGGGCCAGTGGTCCAGACAATGTACAAAGCCTGCCCCTGTGGC<br>CCTTGCCAAAGGGCCAGATGTCAAGAGAGAAGCAACCAGATCCGGAGTCGGCCCCTAGCCAGAAGCTCACCGGAA |
| TGIF1 | pDEST-AD | JBP005 | JBP006 | 58 | ura | ATGAAAGGCAAGAAAGGTATTGTTGCAGCATCTGGCAGTGAGACTGAGGATGAGGACAGCATGGACATTCCCTTGGACCT<br>TTCTTCATCCGCTGGCTCAGGCAAGAGAAGGAGAAGGGGCAACCTACCCAAGGAGTCTGTGCAGATTCTTCGGGATTGGC<br>TGTATGAGCACCGTTACAATGCCTATCCTTCAGAGCAAGAAAAAGCGTTGCTGTCCCAGCAAACACACCTGTCTACGCTA<br>CAGGTCTGTAACTGGTTCATCAACGCCCGCCGCAGGCTCCTCCCTGACATGCTGAGAAAGGATGGCAAAGATCCAAATCA<br>GTTCACAATTTCCCGCCGTGGGGCCAAGATTTCTGAAACGAGCTCTGTGGAGTCCGTGATGGGCATCAAAAACTTCATGC<br>CAGCTCTAGAGGAGACCCCATTTCATTCCTGTACAGCTGGGCCAAACCCAACCCTAGGGAGGCCACTGTCTCCTAAGCCG<br>TCATCCCCGGGATCAGTTTTGGCTCGTCCATCAGTGATCTGCCATACCACTGTGACTGCATTGAAAGATGTCCCTTTCTC<br>TCTCTGCCAGTCGGTCGGTGTGGGACAAAACACAGATATACAGCAGATAGCGGCCAAAAACTTCACAGACACCTCTCTCA<br>TGTACCCAGAGGACACTTGTAAATCTGGACCAAGTACGAATACACAGAGTGGTCTTTTCAACACTCCTCCCCCTACTCCA<br>CCGGACCTCAACCAGGACTTCAGTGGATTTCAGCTTCTAGTGGATGTTGCACTCAAACGGGCTGCAGAGATGGAGCTTCA<br>GGCAAAACTTACAGCTTAG |
| TGIF2 | ORFeome | JBP019_TGIF2_F | JBP045_attL2_pDONR221_201 | 58 | ura | ATGTCGGACAGTGATCTAGGTGAGGACGAAGGCCTCCTCTCCCTGGCGGGCAAAAGGAAGCGCAGGGGGAACCTGCCCAA<br>GGAGTCGGTGAAGATCCTCCGGGACTGGCTGTACTTGCACCGCTACAACGCCTACCCCTCAGAGCAGGAGAAGCTGAGCC<br>TTTCTGGACAGACCAACCTGTCAGTGCTGCAAATATGTAACTGGTTCATCAATGCCCGGCGGCGGCTTCTCCCAGACATG<br>CTTCGGAAGGATGGCAAAGACCCTAATCAGTTTACCATTTCCCGCCGCGGGGGTAAGGCCTCAGATGTGGCCCTCCCCCG<br>TGGCAGCAGCCCCTCAGTGCTGGCTGTGTCTGTCCCAGCCCCCACCAATGTGCTCTCCCTGTCTGTGTGCTCCATGCCGC<br>TTCACTCAGGCCAGGGGGAAAAGCCAGCAGCCCCTTTCCCACGTGGGGAGCTGGAGTCTCCCAAGCCCCTGGTGACCCCT<br>GGTAGCACACTTACTCTGCTGACCAGGGCTGAGGCTGGAAGCCCCACAGGTGGACTCTTCAACACGCCACCACCCACACC<br>CCCAGAGCAGGACAAAGAGGACTTCAGCAGCTTCCAGCTGCTGGTGGAGGTGGCGCTACAGAGGGCTGCTGAGATGGAGC<br>TTCAGAAGCAGCAGGACCCATCACTCCCATTACTGCACACTCCCATCCCTTTAGTCTCTGAAAATCCCCAGTTG |
| POU2F3 | ORFeome | JBP020_POU2F3_F | JBP045_attL2_pDONR221_201 | 58 | ura | ATGGTGAATCTGGAGTCCATGCACACAGATATCAAGATGAGTGGGGATGTAGCCGATTCCACGGATGCTCGCAGCACTCT<br>CAGCCAGGTGGAGCCAGGAAATGATCGAAATGGCCTAGATTTCAACAGGCAGATTAAAACCGAAGATCTCAGTGACTCCC<br>TGCAGCAGACCCTCTCCCATCGGCCATGCCACCTGAGTCAAGGACCTGCCATGATGTCCGGAAACCAAATGTCTGGGCTA<br>AATGCCAGCCCATGTCAGGACATGGCTTCCCTCCATCCGCTCCAGCAGCTTGTGCTGGTTCCCGGCCACTTACAGTCTGT<br>ATCCCAGTTCCTGCTATCTCAGACCCAGCCTGGGCAGCAAGGTCTGCAGCCAAATCTCCTCCCCTTTCCACAGCAACAAA<br>GCGGTCTCCTCCTCCCACAGACTGGGCCGGGACTGGCATCCCAGGCATTTGGGCACCCTGGGCTGCCAGGATCCTCTTTA<br>GAACCCCACCTGGAAGCATCCCAGCATCTCCCAGTGCCCAAGCATCTACCCAGCTCTGGAGGGGCCGATGAGCCCAGTGA<br>CCTCGAGGAGCTGGAGAAGTTTGCCAAGACCTTCAAGCAGAGGCGCATTAAGCTGGGCTTCACACAGGGAGATGTGGGGC<br>TGGCGATGGGAAAGCTGTATGGCAACGACTTCAGCCAGACCACCATCTCACGATTTGAGGCCCTCAACCTGAGCTTCAAG<br>AACATGTGCAAGCTCAAGCCCCTGCTGGAGAAGTGGCTGAATGATGCAGAGTCCTCTCCGTCAGACCCCTCAGTGAGCAC<br>GCCCAGCTCCTACCCCAGCCTCAGTGAAGTATTTGGTAGGAAGAGAAAGAAACGGACCAGCATCGAGACCAACATCCGCC<br>TGACTCTGGAGAAGAGGTTTCAAGATAACCCAAAACCCAGCTCGGAGGAGATCTCCATGATTGCAGAGCAGTTGTCCATG<br>GAGAAGGAGGTGGTGAGGGTCTGGTTCTGCAACCGACGCCAAAAGGAGAAGCGAATCAACTGCCCTGTGGCCACACCCAT<br>CAAACCACCTGTCTACAACTCCCGGCTGGTATCTCCCTCAGGGTCTCTGGGCCCCCTCTCTGTCCCTCCTGTCCACAGTA<br>CCATGCCTGGAACAGTAACGTCATCCTGTTCCCCTGGGAACAACAGCAGGCCTTCATCTCCTGGCTCAGGACTCCACGCC<br>AGCAGCCCCACTGCATCTCAAAATAACTCCAAAGCAGCAGTGAACTCCGCCTCCAGTTTTAACTCTTCAGGATCTTGGTA<br>CCGATGGAATCATTCCACCTACCTCCACTTG |
| POU3F4 | ORFeome | JBP021_POU3F4_F | JBP043_attL2_pENTR223.1 | 58 | ura | ATGGCCACAGCTGCCTCGAATCCCTACAGCATTCTCAGTTCCACCTCCCTAGTCCATGCGGACTCTGCGGGCATGCAGCA<br>GGGGAGTCCTTTCCGCAACCCTCAGAAACTTCTCCAAAGTGATTACTTGCAGGGAGTTCCCAGCAATGGGCATCCCCTCG<br>GGCATCACTGGGTGACCAGTCTGAGCGACGGGGGCCCATGGTCCTCCACACTGGCCACCAGCCCCCTGGACCAGCAGGAC<br>GTGAAGCCCGGGCGCGAAGACCTGCAACTGGGTGCGATCATCCATCACCGCTCGCCACACGTAGCCCACCACTCACCGCA<br>CACTAACCACCCCAACGCCTGGGGGGCCAGCCCGGCACCGAACCCGTCTATCACGTCAAGCGGCCAACCCCTCAACGTGT<br>ACTCGCAGCCTGGCTTCACCGTGAGCGGCATGCTGGAACACGGGGGACTCACCCCACCTCCAGCTGCCGCCTCTGCACAG<br>AGCCTGCACCCGGTGCTCCGAGAGCCCCCGGATCACGGCGAACTGGGCTCGCACCATTGCCAGGATCACTCCGACGAGGA<br>GACGCCAACCTCTGATGAGTTGGAACAGTTCGCCAAACAATTCAAACAAAGAAGAATCAAGTTGGGCTTCACGCAGGCCG<br>ACGTGGGGTTGGCGCTGGGCACACTGTATGGTAACGTGTTCTCGCAGACCACCATCTGCAGGTTCGAAGGCTTGCAGCTG<br>AGCTTCAAAAATATGTGCAAGCTGAAGCCCCTGCTGAACAAGTGGCTGGAGGAGGCGGATTCGTCCACAGGGAGCCCGAC<br>CAGCATTGACAAGATCGCTGCACAGGGCCGCAAGCGCAAGAAGCGGACCTCCATCGAGGTGAGTGTCAAGGGCGTACTGG<br>AGACGCATTTCCTCAAGTGTCCCAAGCCTGCCGCGCAGGAGATCTCCTCGCTGGCAGACAGCCTCCAGTTGGAGAAGGAA<br>GTGGTGCGTGTCTGGTTCTGTAATCGAAGACAAAAAGAGAAAAGAATGACTCCGCCAGGGGATCAGCAGCCGCATGAGGT<br>TTATTCGCACACCGTGAAAACAGACACATCTTGCCATGATCTC |

| | | | | | | |
|---|---|---|---|---|---|---|
| POU3F1 | ORFeome | JBP022_POU3F1_F | JBP043_attL2_pENTR223.1 | 58 | ura | ATGGCCACCACCGCGCAGTACCTGCCGCGGGGCCCCGGTGGCGGAGCCGGGGGCACCGGGCCGCTTATGCACCCGGACGC<br>CGCGGCGGCGGCGGCGGCGGCGGCGGCCGCGGAGCGATTGCATGCAGGGGCCGCGTACCGCGAAGTGCAGAAGCTGATGC<br>ACCACGAGTGGCTGGGCGCGGGCGCGGGCCACCCCGTGGGCCTAGCGCACCCCCAGTGGCTACCCACGGGAGGAGGCGGC<br>GGCGGCGATTGGGCCGGCGGCCCGCACCTAGAACACGGCAAGGCAGGCGGCGGCGGCACCGGCCGAGCCGACGACGGCGG<br>CGGCGGCGGAGGTTTCCACGCGCGCCTGGTGCACCAGGGGGCGGCCCACGCGGGCGCGGCATGGGCGCAGGGCAGCACAG<br>CGCACCACTTGGGCCCGGCCATGTCGCCCTCGCCCGGGGCCAGCGGGGGCCACCAGCCCCAGCCGCTCGGGCTGTACGCG<br>CAGGCGGCCTACCCGGGGGGCGGCGGCGGCGGCCTGGCCGGGATGCTGGCGGCGGGCGGTGGCGGCGCGGGGCCGGGCCT<br>GCACCACGCGCTGCACGAGGATGGCCACGAGGCGCAGCTGGAGCCGTCGCCGCCGCCGCATCTGGGCGCCCACGGACACG<br>CACACGGACATGCACACGCGGGCGGCCTGCACGCGGCGGCGGCGCACCTGCACCCGGGCGCGGGCGGCGGCGGCTCATCG<br>GTGGGCGAGCACTCGGACGAGGATGCGCCCAGCTCGGACGACCTGGAGCAGTTCGCCAAGCAGTTCAAGCAGCGGCGCAT<br>CAAGCTGGGCTTTACGCAGGCCGACGTGGGGCTGGCGCTGGGCACGCTCTACGGTAACGTGTTCTCGCAGACCACCATCT<br>GCCGCTTCGAGGCCCTGCAGCTGAGCTTCAAGAACATGTGCAAGCTCAAGCCGCTGCTCAACAAGTGGCTGGAGGAGACC<br>GACTCGTCCAGCGGCAGCCCCACCAACCTGGACAAGATCGCGGCGCAGGGCCGCAAGCGCAAGAAGCGCACGTCCATCGA<br>GGTGGGGGTCAAAGGCGCGCTCGAGAGCCACTTTCTCAAGTGCCCCAAGCCCTCGGCGCACGAGATCACCGGCTTGGCAG<br>ACAGCCTGCAGCTGGAGAAGGAGGTGGTGCGCGTCTGGTTCTGCAACCGGCGGCAGAAGGAGAAGCGCATGACCCCTGCG<br>GCCGGCGCGGGCCACCCGCCCATGGACGATGTATACGCGCCTGGGGAGCTAGGGCCTGGCGGGGGCGGCGCATCGCCACC<br>CTCCGCGCCCCCACCGCCCCCGCCGGCGGCGCTGCACCACCACCACCACCACACACTGCCCGGCTCAGTGCAG |
| GATA3 | ORFeome | JBP024_GATA3_F | JBP045_attL2_pDONR221_201 | 58 | ura | ATGGAGGTGACGGCGGACCAGCCGCGCTGGGTGAGCCACCACCACCCCGCCGTGCTCAACGGGCAGCACCCGGACACGCA<br>CCACCCGGGCCTCAGCCACTCCTACATGGACGCGGCGCAGTACCCGCTGCCGGAGGAGGTGGATGTGCTTTTTAACATCG<br>ACGGTCAAGGCAACCACGTCCCGCCCTACTACGGAAACTCGGTCAGGGCCACGGTGCAGAGGTACCCTCCGACCCACCAC<br>GGGAGCCAGGTGTGCCGCCCGCCTCTGCTTCATGGATCCCTACCCTGGCTGGACGGCGGCAAAGCCCTGGGCAGCCACCA<br>CACCGCCTCCCCCTGGAATCTCAGCCCCTTCTCCAAGACGTCCATCCACCACGGCTCCCCGGGGCCCCTCTCCGTCTACC<br>CCCCGGCCTCGTCCTCCTCCTTGTCGGGGGGCCACGCCAGCCCGCACCTCTTCACCTTCCCGCCCACCCCGCCGAAGGAC<br>GTCTCCCCGGACCCATCGCTGTCCACCCCAGGCTCGGCCGGCTCGGCCCGGCAGGACGAGAAAGAGTGCCTCAAGTACCA<br>GGTGCCCCTGCCCGACAGCATGAAGCTGGAGTCGTCCCACTCCCGTGGCAGCATGACCGCCCTGGGTGGAGCCTCCTCGT<br>CGACCCACCACCCCATCACCACCTACCCGCCCTACGTGCCCGAGTACAGCTCCGGACTCTTCCCCCCCAGCAGCCTGCTG<br>GGCGGCTCCCCCACCGGCTTCGGATGCAAGTCCAGGCCCAAGGCCCGGTCCAGCACAGAAGGCAGGGAGTGTGTGAACTG<br>TGGGGCAACCTCGACCCCACTGTGGCGGCGAGATGGCACGGGACACTACCTGTGCAACGCCTGCGGGCTCTATCACAAAA<br>TGAACGGACAGAACCGGCCCCTCATTAAGCCCAAGCGAAGGCTGTCTGCAGCCAGGAGAGCAGGGACGTCCTGTGCGAAC<br>TGTCAGACCACCACAACCACACTCTGGAGGAGGAATGCCAATGGGGACCCTGTCTGCAATGCCTGTGGGCTCTACTACAA<br>GCTTCACAATATCAACAGACCCCTGACTATGAAGAAGGAAGGCATCCAGACCAGAAACCGAAAAATGTCTAGCAAATCCA<br>AAAAGTGCAAAAAAGTGCATGACTCACTGGAGGACTTCCCCAAGAACAGCTCGTTTAACCCGGCCGCCCTCTCCAGACAC<br>ATGTCCTCCCTGAGCCACATCTCGCCCTTCAGCCACTCCAGCCACATGCTGACCACGCCCACGCCGATGCACCCGCCATC<br>CAGCCTGTCCTTTGGACCACACCACCCCTCCAGCATGGTCACCGCCATGGGTTTG |
| GATA6 | ORFeome | JBP041_GATA6_F | JBP045_attL2_pDONR221_201 | 58 | ura | ATGGCCTTGACTGACGGCGGCTGGTGCTTACCGAAGCGATTCGGAGCCGCGGGTGCTGATGCCAGCGACTCCAGAGCCTT<br>TCCAGCTCGGGAGCCCTCTACGCCGCCTTCTCCTATCTCTTCCTCGTCTTCCTCTTGCTCCAGGGGAGGAGAGAGGGGAC<br>CTGGCGGAGCCAGTAACTGTGGAACGCCTCAGCTAGATACGGAGGCTGCCGCCGGACCCCCGGCTCGATCGCTTCTGCTC<br>AGTTCTTACGCTTCGCATCCCTTCGGAGCTCCTCACGGACCTTCGGCTCCTGGTGTCGCGGGTCCTGGGGGCAACCTGTC<br>GAGCTGGGAGGACTTGCTGCTGTTCACTGACCTCGACCAAGCCGCGACAGCCAGCAAGCTGCTGTGGTCCAGTCGAGGCG<br>CCAAGCTGAGCCCCTTCGCACCCGAACAGCCGGAGGAGATGTACCAGACCCTTGCCGCTCTTTCTAGCCAGGGTCCAGCC<br>GCTTACGATGGCGCTCCTGGAGGATTTGTTCACTCTGCGGCTGCAGCGGCAGCAGCCGCGGCCGCAGCTAGCTCCCCTGT<br>CTACGTTCCTACCACCCGCGTTGGTTCCATGCTGCCCGGACTACCTTACCATCTGCAGGGATCGGGCAGTGGACCAGCCA<br>ACCATGCTGGAGGCGCGGGAGCGCATCCCGGATGGCCTCAAGCTTCGGCAGACAGTCCTCCATATGGTAGTGGAGGCGGC<br>GCAGCTGGAGGAGGAGCTGCTGGACCTGGTGGCGCTGGATCAGCTGCTGCGCATGTCTCTGCGCGATTCCCTTACTCTCC<br>CAGCCCTCCCATGGCCAACGGAGCTGCGAGGGAGCCAGGAGGATACGCTGCGGCAGGAAGTGGGGGAGCGGGAGGAGTTA<br>GTGGCGGAGGAAGTAGCTTAGCGGCTATGGGCGGACGCGAACCCCAGTACAGCTCTCTGTCAGCCGCCCGGCCGCTGAAC<br>GGGACGTACCACCACCATCACCATCATCATCACCACCATCCGAGCCCCTACTCGCCCTACGTGGGTGCGCCACTTACGCC<br>TGCCTGGCCCGCTGGACCATTCGAAACCCCGGTGCTTCATAGCCTTCAGAGCCGTGCTGGAGCCCCGCTTCCGGTTCCCC<br>GAGGTCCCAGTGCAGACCTTCTGGAGGACCTGTCCGAGAGCCGCGAGTGCGTGAATTGCGGCTCTATCCAGACGCCGCTG<br>TGGCGACGGGACGGCACCGGACACTACCTGTGCAACGCTTGCGGGCTCTACAGCAAGATGAACGGCCTCAGCCGGCCCCT<br>CATCAAGCCCCAGAAGCGCGTGCCTTCATCACGGCGCCTTGGATTGTCCTGTGCCAACTGTCACACCACAACTACCACCT<br>TATGGCGCAGAAACGCCGAGGGTGAACCCGTGTGCAATGCTTGTGGACTGTACATGAAACTCCATGGGGTGCCCAGACCA<br>CTTGCTATGAAAAAAGAGGGAATTCAAACCAGGAAACGAAAACCTAAGAACATAAATAAATCAAAGACTTGCTCTGGTAA<br>TAGCAATAATTCCATTCCCATGACTCCAACTTCCACCTCTTCTAACTCAGATGATTGCAGCAAAAATACTTCCCCCACAA<br>CACAACCTACAGCCTCAGGGGCGGGTGCCCCGGTGATGACTGGTGCGGGAGAGAGCACCAATCCCGAGAACAGCGAGCTC<br>AAGTATTCGGGTCAAGATGGACTCTACATAGGCGTCAGTCTCGCCTCGCCGGCCGAAGTCACGTCCTCCGTGCGACCGGA<br>TTCCTGGTGCGCCCTGGCCCTGGCC |
| GATA4 | pDEST-AD | JBP005 | JBP006 | 58 | ura | ATGTATCAGAGCTTGGCCATGACCGCCAACCACGGGCCGCCCCCCGGTGCCTACGAGGCGGGCGGCCCCGGCGCCTTCAT<br>GCACGGCGCGGGCGCCGCGTCCTCGCCAGTCTACGTGCCCACACCGCGGGTGCCCTCCTCCGTGCTGGGCCTGTCCTACC<br>TCCAGGGCGGAGGCGCGGGCTCTGCGTCCGGAGGCGCCTCGGGCGGCAGCTCCGGTGGGGCCGCGTCTGGTGCGGGGCCC<br>GGGACCCAGCAGGGCAGCCCGGGATGGAGCCAGGCGGGAGCCGACGGAGCCGCTTACACCCCGCCGCCGGTGTCGCCGCG<br>CTTCTCCTTCCCGGGGACCACCGGGTCCCTGGCGGCCGCCGCCGCCGCTGCCGCGGCCCGGGAAGCTGCGGCCTACAGCA<br>GTGGCGGCGGAGCGGCGGGTGCGGGCCTGGCGGGCCGCGAGCAGTACGGGCGCGCCGGCTTCGCGGGCTCCTACTCCAGC<br>CCCTACCCGGCTTACATGGCCGACGTGGGCGCGTCCTGGGCCGCAGCCGCCGCCGCCTCCGCCGGCCCCTTCGACAGCCC<br>GGTCCTGCACAGCCTGCCCGGCCGGGCCAACCCGGCCGCCCGACACCCCAATCTCGATATGTTTGACGACTTCTCAGAAG<br>GCAGAGAGTGTGTCAACTGTGGGGCTATGTCCACCCCGCTCTGGAGGCGAGATGGGACGGGTCACTATCTGTGCAACGCC<br>TGCGGCCTCTACCACAAGATGAACGGCATCAACCGGCCGCTCATCAAGCCTCAGCGCCGGCTGTCCGCCTCCCGCCGAGT<br>GGGCCTCTCCTGTGCCAACTGCCAGACCACCACCACCACGCTGTGGCGCCGCAATGCGGAGGGCGAGCCTGTGTGCAATG<br>CCTGCGGCCTCTACATGAAGCTCCACGGGGTCCCCAGGCCTCTTGCAATGCGGAAAGAGGGGATCCAAACCAGAAAACGG<br>AAGCCCAAGAACCTGAATAAATCTAAGACACCAGCAGCTCCTTCAGGCAGTGAGAGCCTTCCTCCCGCCAGCGGTGCTTC<br>CAGCAACTCCAGCAACGCCACCACCAGCAGCAGCGAGGAGATGCGTCCCATCAAGACGGAGCCTGGCCTGTCATCTCACT<br>ACGGGCACAGCAGCTCCGTGTCCCAGACGTTCTCAGTCAGTGCGATGTCTGGCCATGGGCCCTCCATCCACCCTGTCCTC<br>TCGGCCCTGAAGCTCTCCCCACAAGGCTATGCGTCTCCCGTCAGCCAGTCTCCACAGACCAGCTCCAAGCAGGACTCTTG<br>GAACAGCCTGGTCTTGGCCGACAGTCACGGGGACATAATCACTGCG |
| GATA2 | pDEST-AD | JBP005 | JBP006 | 58 | ura | ATGGAGGTGGCGCCGGAGCAGCCGCGCTGGATGGCGCACCCGGCCGTGCTGAATGCGCAGCACCCCGACTCACACCACCC<br>GGGCCTGGCGCACAACTACATGGAACCCGCGCAGCTGCTGCCTCCAGACGAGGTGGACGTCTTCTTCAATCACCTCGACT<br>CGCAGGGCAACCCCTACTATGCCAACCCCGCTCACGCGCGGGCGCGCGTCTCCTACAGCCCCGCGCACGCCCGCCTGACC<br>GGAGGCCAGATGTGCCGCCCACACTTGTTGCACAGCCCGGGTTTGCCCTGGCTGGACGGGGGCAAAGCAGCCCTCTCTGC<br>CGCTGCGGCCCACCACCACAACCCCTGGACCGTGAGCCCCTTCTCCAAGACGCCACTGCACCCCTCAGCTGCTGGAGGCC<br>CTGGAGGCCCACTCTCTGTGTACCCAGGGGCTGGGGGTGGAGCGGGGGAGGCAGCGGGAGCTCAGTGGCCTCCCTCACC<br>CCTACAGCAGCCCACTCTGGCTCCCACCTTTTCGGCTTCCCACCCACGCCACCCAAAGAAGTGTCTCCTGACCCTAGCAC<br>CACGGGGGCTGCGTCTCCAGCCTCATCTTCCGCGGGGGGTAGTGCAGCCCGAGGAGAGGACAAGGACGGCGTCAAGTACC<br>AGGTGTCACTGACGGAGAGCATGAAGATGGAAAGTGGCAGTCCCCTGCGCCCAGGCCTAGCTACTATGGGCACCCAGCCT<br>GCTACACACCACCCCATCCCCACCTACCCCTCCTATGTGCCGGCGGCTGCCCACGACTACAGCAGCGGACTCTTCCACCC<br>CGGAGGCTTCCTGGGGGGACCGGCCTCCAGCTTCACCCCTAAGCAGCGCAGCAAGGCTCGTTCCTGTTCAGAAGGCCGGG<br>AGTGTGTCAACTGTGGGGCCACAGCCACCCCTCTCTGGCGGCGGGACGGCACCGGCCACTACCTGTGCAATGCCTGTGGC<br>CTCTACCACAAGATGAATGGGCAGAACCGACCACTCATCAAGCCCAAGCGAAGACTGTCGGCCGCCAGAAGAGCCGGCAC<br>CTGTTGTGCAAATTGTCAGACGACAACCACCACCTTATGGCGCCGAAACGCCAACGGGGACCCTGTCTGCAACGCCTGTG<br>GCCTCTACTACAAGCTGCACAATGTTAACAGGCCACTGACCATGAAGAAGGAAGGGATCCAGACTCGGAACCGGAAGATG<br>TCCAACAAGTCCAAGAAGAGCAAGAAAGGGGCGGAGTGCTTCGAGGAGCTGTCAAAGTGCATGCAGGAGAAGTCATCCCC<br>CTTCAGTGCAGCTGCCCTGGCTGGACACATGGCACCTGTGGGCCACCTCCCGCCCTTCAGCCACTCCGGACACATCCTGC<br>CCACTCCGACGCCCATCCACCCCTCCTCCAGCCTCTCCTTCGGCCACCCCCACCCGTCCAGCATGGTGACCGCCATGGGC<br>TAG |
| GATA5 | pDEST-AD | JBP005 | JBP006 | 58 | ura | ATGTACCAGAGCCTGGCGCTGGCCGCGAGCCCCCGCCAGGCCGCCTACGCCGACTCGGGCTCCTTCCTGCACGCTCCGGG<br>CGCCGGCTCTCCGATGTTTGTGCCGCCGGCGCGCGTCCCCTCGATGCTGTCCTACCTGTCCGGGTGTGAGCCGAGCCCGC<br>AGCCCCCCGAGCTCGCTGCGCGCCCCGGCTGGGCGCAGACAGCCACCGCGGATTCGTCGGCCTTCGGCCCGGGCAGTCCG<br>CACCCCCCAGCCGCGCACCCGCCCGGGGCCACCGCCTTCCCTTTCGCGCACAGCCCCTCGGGGCCCGGCAGCGGCGGCAG<br>CGCGGGGGGCCGAGACGGCAGTGCCTACCAGGGCGCGCTGTTGCCTCGAGAACAGTTCGCGGCCCCGCTTGGGCGGCCGG<br>TGGGGACCTCGTACTCCGCCACCTACCCGGCCTACGTGAGCCCCGACGTGGCCCAGTCCTGGACTGCCGGGCCCTTCGAT<br>GGCAGCGTCCTGCACGGCCTCCCAGGCCGCAGGCCCACCTTCGTGTCCGACTTCTTGGAGGAGTTCCCGGGTGAGGGTCG<br>TGAGTGTGTCAACTGCGGGGCCCTGTCCACACCGCTGTGGCGCCGAGATGGCACCGGCCACTACCTGTGCAATGCCTGCG<br>GCCTCTACCACAAGATGAATGGCGTCAACCGGCCGCTCGTTCGGCCTCAGAAGCGCCTGTCCTCGTCCCGCCGCGCCGGC<br>CTCTGCTGCACCAACTGCCACACGACCAACACCACGCTGTGGCGGCGGAACTCGGAGGGGGAGCCCGTGTGCAATGCCTG<br>CGGCCTCTACATGAAGCTGCACGGGGTGCCGCGGCCTCTGGCTATGAAGAAAGAAAGCATCCAGACACGGAAGCGGAAGC<br>CAAAGACCATCGCCAAGGCCAGGGGCTCCTCAGGATCCACAAGGAATGCCTCGGCCTCCCCATCTGCTGTCGCCAGCACT<br>GACAGCTCAGCAGCCACTTCCAAAGCCAAGCCCAGCCTGGCGTCCCCAGTGTGCCCTGGGCCCAGCATGGCCCCCCAGGC<br>CTCTGGCCAGGAGGATGACTCTCTTGCCCCCGGCCACTTGGAGTTCAAGTTCGAGCCTGAGGACTTTGCCTTCCCCTCCA<br>CGGCCCCGAGCCCCCAGGCTGGCCTCAGGGGGGCTCTGCGCCAAGAGGCCTGGTGTGCGCTGGCCTTGGCC |

| | | | | | | |
|---|---|---|---|---|---|---|
| FOXL2_DBD_FOXO3 | FOXL2 strain | JBP115_FOXL2_DBD_FOXO3_IDR_F | JBP116_FOXL2_DBD_FOXO3_IDR_R | 58 | 47-48 | ATGGCAGAGGCACCGGCTTCCCCGGCCCCGCTCTCTCCGCTCGAAGTGGAGCTGGACCCGGAGTTCGAGCCCCAGAGCCGTCCGCGATCCTGTACGTGGCCCCTGCAAAGGCCGGAGCTCCAAGCGAGCCCTGCCAAGCCCTCGGGGGAGACGGCCGCCGACTCCATGATCCCCGAGGAGGAGGACGATGAAGACGACGAGGACGGCGGGGGACGGGCCGGCTCGGCCATGGCGATCGGCGGCGGCGGCGGGAGCGGCACGCTGGGCTCCGGGCTGCTCCTTGAGGACTCGGCCCGGGTGCTGGCACCCGGAGGGCAAGACCCCGGGTCTGGGCCAGCCACCGCGGCGGGCGGGCTGAGCGGGGGTACACAGGCGCTGCTGCAGCCTCAGCAACCGCTGCCACCACCGCAGCCGGGGGCGGCTGGGGGCTCCGGGCAGCCGAGGAAATGTTCGTCGCGGCGGAACGCCTGGGGAAACAAGCCCCCGTACTCGTACGTGGCGCTCATCGCCATGGCGATCCGCGAGAGCGCGGAGAAGAGGCTCACGCTGTCCGGCATCTACCAGTACATCATCGCGAAGTTCCCGTTCTACGAGAAGAATAAGAAGGGCTGGCAAAATAGCATCCGCCACAACCTCAGCCTCAACGAGTGCTTCATCAAGGTGCCGCGCGAGGGCGGCGGCGAGCGCAAGGGCAACTACTGGACGCTGGACCCGGCCTGCGAAGACATGTTCGAGAAGAAGAGCGGAAAAGCCCCCCGGCGGCGGGCTGTCTCCATGGACAATAGCAACAAGTATACCAAGAGCCGTGGCCGCGCAGCCAAGAAGAAGGCAGCCCTGCAGACAGCCCCCGAATCAGCTGACGACAGTCCCTCCCAGCTCTCCAAGTGGCCTGGCAGCCCCACGTCACGCAGCAGTGATGAGCTGGATGCGTGGACGGACTTCCGTTCACGCACCAATTCTAACGCCAGCACAGTCAGTGGCCGCCTGTCGCCCATCATGGCAAGCACAGAGTTGGATGAAGTCCAGGACGATGATGCGCCTCTCTCGCCCATGCTCTACAGCAGCTCAGCCAGCCTGTCACCTTCAGTAAGCAAGCCGTGCACGGTGGAACTGCCACGGCTGACTGATATGGCAGGCACCATGAATCTGAATGATGGGCTGACTGAAAACCTCATGGACGACCTGCTGGATAACATCACGCTCCCGCCATCCCAGCCATCGCCCACTGGGGGACTCATGCAGCGGAGCTCTAGCTTCCCGTATACCACCAAGGGCTCGGGCCTGGGCTCCCCAACCAGCTCCTTTAACAGCACGGTGTTCGGACCTTCATCTCTGAACTCCCTACGCCAGTCTCCCATGCAGACCATCCAAGAGAACAAGCCAGCTACCTTCTCTTCCATGTCACACTATGGTAACCAGACACTCCAGGACCTGCTCACTTCGGACTCACTTAGCCACAGCGATGTCATGATGACACAGTCGGACCCCTTGATGTCTCAGGCCAGCACCGCTGTGTCTGCCCAGAATTCCCGCCGGAACGTGATGCTTCGCAATGATCCGATGATGTCCTTTGCTGCCCAGCCTAACCAGGGAAGTTTGGTCAATCAGAACTTGCTCCACCACCAGCACCAAACCCAGGGCGCTCTTGGTGGCAGCCGTGCCTTGTCGAATTCTGTCAGCAACATGGGCTTGAGTGAGTCCAGCAGCCTTGGGTCAGCCAAACACCAGCAGCAGTCTCCTGTCAGCCAGTCTATGCAAACCCTCTCGGACTCTCTCTCAGGCTCCTCCTTGTACTCAACTAGTGCAAACCTGCCCGTCATGGGCCATGAGAAGTTCCCCAGCGACTTGGACCTGGACATGTTCAATGGGAGCTTGGAATGTGACATGGAGTCCATTATCCGTAGTGAACTCATGGATGCTGATGGGTTGGATTTTAACTTTGATTCCCTCATCTCCACACAGAATGTTGTTGGTTTGAACGTGGGGAACTTCACTGGTGCTAAGCAGGCCTCATCTCAGAGCTGGGTGCCAGGC |
| FOXL2_DBD_FOXA2 | FOXL2 strain | JBP117_FOXL2_DBD_FOXA2_IDR_F | JBP118_FOXL2_DBD_FOXA2_IDR_R | 58 | 49-50 | ATGCTGGGAGCGGTGAAGATGGAAGGGCACGAGCCGTCCGACTGGAGCAGCTACTATGCAGAGCCCGAGGGCTACTCCTCCGTGAGCAACATGAACGCCGGCCTGGGGATGAACGGCATGAACACGTACATGAGCATGTCGGCGGCCGCCATGGGCAGCGGCTCGGGCAACATGAGCGCGGGCTCCATGAACATGTCGTCGTACGTGGGCGCTGGCATGAGCCCGTCCCTGGCGGGGATGTCCCCCGGCGCGGGCGCCATGGCGGGCATGGGCGGCTCGGCCGGGGCGGCTGGCGTGGCGGGCATGGGGCCGCACTTGAGTCCCAGCCTGAGCCCGCTCGGGGGGCAGGCGGCCGGGGCCATGGGCGGCCTGGCCCCCTACGCCAACATGAACTCCATGAGCCCCATGTACGGGCAGGCGGGCCTGAGCCGCGCCCGCGACCCCAAGACCTACAGGCGCAGCTACACGCACGCAAAGAAGCCCCCGTACTCGTACGTGGCGCTCATCGCCATGGCGATCCGCGAGAGCGCGGAGAAGAGGCTCACGCTGTCCGGCATCTACCAGTACATCATCGCGAAGTTCCCGTTCTACGAGAAGAATAAGAAGGGCTGGCAAAATAGCATCCGCCACAACCTCAGCCTCAACGAGTGCTTCATCAAGGTGCCGCGCGAGGGCGGCGGCGAGCGCAAGGGCAACTACTGGACGCTGGACCCGGCCTGCGAAGACATGTTCGAGAAGGGCTGCTACCTGCGCCGCCAGAAGCGCTTCAAGTGCGAGAAGCAGCTGGCGCTGAAGGAGGCCGCAGGCGCCGCCGGCAGCGGCAAGAAGGCGGCCGCCGGGGCCCAGGCCTCACAGGCTCAACTCGGGGAGGCCGCCGGGCCGGCCTCCGAGACTCCGGCGGGCACCGAGTCGCCTCACTCGAGCGCCTCCCCGTGCCAGGAGCACAAGCGAGGGGGCCTGGGAGAGCTGAAGGGGACGCCGGCTGCGGCGCTGAGCCCCCCAGAGCCGGCGCCCTCTCCCGGGCAGCAGCAGGCCGCGGCCCACCTGCTGGGCCCGCCCCACCACCCGGGCCTGCCGCCTGAGGCCCACCTGAAGCCGGAACACCACTACGCCTTCAACCACCCGTTCTCCATCAACAACCTCATGTCCTCGGAGCAGCAGCACCACCACAGCCACCACCACCACCAGCCCCACAAAATGGACCTCAAGGCCTACGAACAGGTGATGCACTACCCCGGCTACGGTTCCCCCATGCCTGGCAGCTTGGCCATGGGCCCGGTCACGAACAAAACGGGCCTGGACGCCTCGCCCCTGGCCGCAGATACCTCCTACTACCAGGGGGTGTACTCCCGGCCCATTATGAACTCCTCTTTG |
| FOXL2_DBD_FOXF1 | FOXL2 strain | JBP121_FOXL2_DBD_FOXF1_IDR_F | JBP122_FOXL2_DBD_FOXF1_IDR_R | 58 | 55-56 | ATGGACCCCGCGTCGTCCGGCCCGTCCAAGGCCAAGAAGACCAACGCCGGCATCCGGCGCCCGGAGAAGCCCCCGTACTCGTACGTGGCGCTCATCGCCATGGCGATCCGCGAGAGCGCGGAGAAGAGGCTCACGCTGTCCGGCATCTACCAGTACATCATCGCGAAGTTCCCGTTCTACGAGAAGAATAAGAAGGGCTGGCAAAATAGCATCCGCCACAACCTCAGCCTCAACGAGTGCTTCATCAAGGTGCCGCGCGAGGGCGGCGGCGAGCGCAAGGGCAACTACTGGACGCTGGACCCGGCCTGCGAAGACATGTTCGAGAAGGAGGGCTCCTTTCGGCGGCGGCCGCGCGGCTTCCGAAGGAAATGCCAGGCGCTCAAGCCCATGTACAGCATGATGAACGGGCTCGGCTTCAACCACCTCCCGGACACCTACGGCTTCCAGGGCTCGGCCGGCGGCCTCTCGTGCCCGCCCAACAGCCTGGCGCTGGAGGGCGGCCTGGGCATGATGAACGGCCACTTGCCGGGCAACGTGGACGGCATGGCCCTGCCCAGCCACTCGGTGCCCCACCTGCCTTCCAACGGCGGCCACTCGTACATGGGCGGCTGCGGCGGCGCGGCGGCCGGCGAGTACCCGCACCACGACAGCTCGGTGCCCGCCTCCCCGCTGCTGCCCACCGGCGCCGGTGGGGTCATGGAGCCGCACGCCGTCTACTCGGGCTCGGCGGCGGCCTGGCCGCCCTCGGCGTCCGCGGCGCTCAACAGCGGCGCCTCTTATATCAAGCAGCAGCCCCTGTCCCCCTGTAACCCCGCGGCCAACCCCCTGTCCGGCAGCCTCTCCACGCACTCCCTGGAGCAGCCGTATCTGCACCAGAACAGCCACAACGCCCCAGCCGAGCTGCAAGGCATCCCGCGGTATCACTCGCAGTCGCCCAGCATGTGTGACCGAAAGGAGTTTGTCTTCTCTTTCAACGCCATGGCGTCCTCTTCCATGCACTCGGCCGGCGGGGGCTCCTACTACCACCAGCAGGTCACCTACCAAGACATCAAGCCTTGCGTGATGTGGATCCAC |
| FOXL2_DBD_FOXP3 | FOXL2 strain | JBP123_FOXL2_DBD_FOXP3_IDR_F | JBP124_FOXL2_DBD_FOXP3_IDR_R | 58 | 57-58 | ATGCCCAACCCCAGGCCTGGCAAGCCCTCGGCCCCTTCCTTGGCCCTTGGCCCATCCCCAGGAGCCTCGCCCAGCTGGAGGGCTGCACCCAAAGCCTCAGACCTGCTGGGGGCCCGGGGCCCAGGGGGAACCTTCCAGGGCCGAGATCTTCGAGGCGGGGCCCATGCCTCCTCTTCTTCCTTGAACCCCATGCCACCATCGCAGCTGCAGCTGCCCACACTGCCCCTAGTCATGGTGGCACCCTCCGGGGCACGGCTGGGCCCCTTGCCCCACTTACAGGCACTCCTCCAGGACAGGCCACATTTCATGCACCAGCTCTCAACGGTGGATGCCCACGCCCGGACCCCTGTGCTGCAGGTGCACCCCCTGGAGAGCCCAGCCATGATCAGCCTCACACCACCCACCACCGCCACTGGGGTCTTCTCCCTCAAGGCCCGGCCTGGCCTCCCACCTGGGATCAACGTGGCCAGCCTGGAATGGGTGTCCAGGGAGCCGGCACTGCTCTGCACCTTCCCAAATCCCAGTGCACCCAGGAAGGACAGCACCCTTTCGGCTGTGCCCCAGAGCTCCTACCCACTGCTGGCAAATGGTGTCTGCAAGTGGCCCGGATGTGAGAAGGTCTTCGAAGAGCCAGAGGACTTCCTCAAGCACTGCCAGGCGGACCATCTTCTGGATGAGAAGGGCAGGGCACAATGTCTCCTCCAGAGAGAGATGGTACAGTCTCTGGAGCAGCAGCTGGTGCTGGAGAAGGAGAAGCTGAGTGCCATGCAGGCCCACCTGGCTGGGAAAATGGCACTGACCAAGGCTTCATCTGTGGCATCATCCGACAAGGGCTCCTGCTGCATCGTAGCTGCTGGCAGCCAAGGCCCTGTCGTCCCAGCCTGGTCTGGCCCCCGGGAGGCCCCTGACAGCCTGTTTGCTGTCCGGAGGCACCTGTGGGGTAGCCATGGAAACAGCACATTCCCAGAGTTCCTCCACAACATGGACTACTTCAAGTTCCACAACATGCGAAAGCCCCCGTACTCGTACGTGGCGCTCATCGCCATGGCGATCCGCGAGAGCGCGGAGAAGAGGCTCACGCTGTCCGGCATCTACCAGTACATCATCGCGAAGTTCCCGTTCTACGAGAAGAATAAGAAGGGCTGGCAAAATAGCATCCGCCACAACCTCAGCCTCAACGAGTGCTTCATCAAGGTGCCGCGCGAGGGCGGCGGCGAGCGCAAGGGCAACTACTGGACGCTGGACCCGGCCTGCGAAGACATGTTCGAGAAGAAACGGAGCCAGAGGCCCAGCAGGTGTTCCAACCCTACACCTGGCCCC |
| FOXL2_DBD_FOXP1 | FOXL2 strain | JBP125_FOXL2_DBD_FOXP1_IDR_F | JBP126_FOXL2_DBD_FOXP1_IDR_R | 58 | 59-60 | ATGATGCAAGAATCTGGGACTGAGACAAAAAGTAACGGTTCAGCCATCCAGAATGGGTCGGGCGGCAGCAACCACTTACTAGAGTGCGGCGGTCTTCGGGAGGGGCGGTCCAACGGAGAGACGCCGGCCGTGGACATCGGGGCAGCTGACCTCGCCCACGCCCAGCAGCAGCAGCAACAGGCACTTCAGGTGGCAAGACAGCTCCTTCTTCAGCAGCAACAGCAGCAGCAAGTTAGTGGATTAAAATCTCCCAAGAGGAATGACAAACAACCAGCTCTTCAGGTTCCCGTGTCAGTGGCTATGATGACACCTCAAGTTATCACTCCCCAGCAAATGCAGCAGATCCTCCAGCAACAAGTGCTGAGCCCTCAGCAGCTCCAGGTTCTCCTCCAGCAGCAGCAGGCCCTCATGCTTCAACAGCAGCAGCTTCAAGAGTTTTATAAAAAACAACAGGAACAGTTGCAGCTTCAACTTTTACAACAACAACATGCTGGAAAACAGCCTAAAGAGCAACAGCAGGTGGCTACCCAGCAGTTGGCTTTTCAGCAGCAGCTTTTACAGATGCAGCAGTTACAGCAGCAGCACCTCCTGTCTTTGCAGCGCCAAGGCCTTCTGACAATTCAGCCCGGGCAGCCTGCCCTTCCCCTTCAACCTCTTGCTCAAGGCATGATTCCAACAGAACTGCAGCAGCTCTGGAAAGAAGTGACAAGTGCTCATACTGCAGAAGAAACCACAGGCAACAATCACAGCAGTTTGGATCTGACCACGACATGTGTCTCCTCCTCTGCACCTTCCAAGACCTCCTTAATAATGAACCCACATGCCTCTACCAATGGACAGCTCTCAGTCCACACTCCCAAAAGGGAAAGTTTGTCCCATGAGGAGCACCCCCATAGCCATCCTCTCTATGGACATGGTGTATGCAAGTGGCCAGGCTGTGAAGCAGTGTGCGAAGATTTCCAATCATTTCTAAAACATCTCAACAGTGAGCATGCGCTGGACGATAGAAGTACAGCCCAATGTAGAGTACAAATGCAGGTTGTACAGCAGTTAGAGCTACAGCTTGCAAAAGACAAAGAACGCCTGCAAGCCATGATGACCCACCTGCATGTGAAGTCTACAGAACCCAAAGCCGCCCCTCAGCCCTTGAATCTGGTATCAAGTGTCACTCTCTCCAAGTCCGCATCGGAGGCTTCTCCACAGAGCTTACCTCATACTCCAACGACCCCAACCGCCCCCCTGACTCCCGTCACCCAAGGCCCCTCTGTCATCACAACCACCAGCATGCACACGGTGGGACCCATCCGCAGGCGGTACTCAGACAAATACAACGTGCCCATTTCGTCAGCAGATATTGCGCAGAACCAAGAATTTTATAAGAACGCAGAAGTTAGAAAGCCCCCGTACTCGTACGTGGCGCTCATCGCCATGGCGATCCGCGAGAGCGCGGAGAAGAGGCTCACGCTGTCCGGCATCTACCAGTACATCATCGCGAAGTTCCCGTTCTACGAGAAGAATAAGAAGGGCTGGCAAAATAGCATCCGCCACAACCTCAGCCTCAACGAGTGCTTCATCAAGGTGCCGCGCGAGGGCGGCGGCGAGCGCAAGGGCAACTACTGGACGCTGGACCCGGCCTGCGAAGACATGTTCGAGAAGAAACGAAGGCCACAAAAGATCAGTGGTAACCCTTCCCTTATTAAAAACATGCAGAGCAGCCACGCCTACTGCACACCTCTCAATGCAGCTTTACAGGCTTCAATGGCTGAGAATAGTATACCTCTATACACTACCGCTTCCATGGGAAATCCCACTCTGGGCAACTTAGCCAGCGCAATACGGGAAGAGCTGAACGGGGCAATGGAGCATACCAACAGCAACGAGAGTGACAGCAGTCCAGGCAGATCTCCTATGCAAGCCGTGCATCCTGTACACGTCAAAGAAGAGCCCCTCGATCCAGAGGAAGCTGAAGGGCCCCTGTCCTTAGTGACAACAGCCAACCACAGTCCAGATTTTGACCATGACAGAGATTACGAAGATGAACCAGTAAACGAGGACATGGAGTCAGGCCTCATGGGC |
| FOXL2_DBD_FOXP2 | FOXL2 strain | JBP127_FOXL2_DBD_FOXP2_IDR_F | JBP190_FOXL2_DBD_FOXP2_IDR_R2 | 58 | 61-62 | ATGATGCAGGAATCTGCGACAGAGACAATAAGCAACAGTTCAATGAATCAAAATGGAATGAGCACTCTAAGCAGCCAATTAGATGCTGGCAGCAGAGATGGAAGATCAAGTGGTGACACCAGCTCTGAAGTAAGCACAGTAGAACTGCTGCATCTGCAACAACAGCAGGCTCTCCAGGCAGCAAGACAACTTCTTTTACAGCAGCAAACAAGTGGATTGAAATCTCCTAAGAGCAGTGATAAACAGAGACCACTGCAGGAATTGCTTCCAGAAACAAAATTATGTATCTGTGGCCACTCTTCTGGTGATGGGCATCCTCACAACACATTTGCAGTGCCTGTGTCAGTGGCCATGATGACTCCCCAGGTGATCACCCCTCAGCAAATGCAGCAGATCCTTCAGCAACAAGTCCTGTCTCCTCAGCAGCTACAAGCCCTTCTCCAACAACAGCAGGCTGTCATGCTGCAGCAGCAACAACTACAAGAGTTTTACAAGAAACAGCAAGAGCAGTTACATCTTCAGCTTTTGCAGCAGCAGCAGCAACAGCAGCAGCAGCAACAACAGCAGCAACAACAGCAGCAGCAACAACAACAACAACAGCAGCAACAACAGCAGCAGCAGCAGCAACAGCAGCAGCAGCAGCAACAGCATCCTGGAAAGCAAGCGAAAGAGCAGCAGCAGCAGCAGCAGCAACAGCAATTGGCAGCCCAGCAGCTTGTCTTCCAGCAGCAGCTTCTCCAGATGCAACAACTCCAGCAGCAGCAGCATCTGCTCAGCCTTCAGCGTCAGGGACTCATCTCCATTCCACCTGGCCAGGCAGCACTTCCTGTCCAATCGCTGCCTCAAGCTGGCTTAAGTCCTGCTGAGATTCAGCAGTTATGGAAAGAAGTGACTGGAGTTCACAGTATGGAAGACAATGGCATTAAACATGGAGGGCTAGACCTCACTACTAACAATTCCTCCTCGACTACCTCCTCCAACACTTCCAAAGCATCACCACCAATAACTCATCATTCCATAGTGAATGGACAGTCTTCAGTTCTAAGTGCAAGACGAGACAGCTCGTCACATGAGGAGACTGGGGCCTCTCACACTCTCTATGGCCATGGAGTTTGCAAATGGCCAGGCTGTGAAAGCATTTGTGAAGATTTTGGACAGTTTTTAAAGCACCTTAACAATGAACACGCATTGGATGACCGAAGCACTGCTCAGTGTCGAGTGCAAATGCAGGTGGTGCAACAGTTAGAAATACAGCTTTCTAAAGAACGCGAACGTCTTCAAGCAATGATGACCCACTTGCACATGCGACCCTCAGAGCCCAAACCATCTCCCAAACCTCTAAATCTGGTGTCTAGTGTCACCATGTCGAAGAATATGTTGGAGACATCCCCACAGAGCTTACCTCAAACCCCTACCACACCAACGGCCCCAGTCACCCCGATTACCCAGGGACCCTCAGTAATCACCCCAGCCAGTGTGCCCAATGTGGGAGCCATACGAAGGCGACATTCAGACAAATACAACATTCCCATGTCATCAGAAATTGCCCCAAACTATGAATTTTATAAAAATGCAGATGTCAGAAAGCCCCCGTACTCGTACGTGGCGCTCATCGCCATGGCGATCCGCGAGAGCGCGGAGAAGAGGCTCACGCTGTCCGGCATCTACCAGTACATCATCGCGAAGTTCCCGTTCTACGAGAAGAATAAGAAGGGCTGGCAAAATAGCATCCGCCACAACCTCAGCCTCAACGAGTGCTTCATCAAGGTGCCGCGCGAGGGCGGCGGCGAGCGCAAGGGCAACTACTGGACGCTGGACCCGGCCTGCGAAGACATGTTCGAGAAGAGGTCACAAAAGATAACAGGAAGTCCAACCTTAGTAAAAAATATACCTACCAGTTTAGGCTATGGAGCAGCTCTTAATGCCAGTTTGCAGGCTGCCTTGGCAGAGAGCAGTTTACCTTTGCTAAGTAATCCTGGACTGATAAATAATGCATCCAGTGGCCTACTGCAGGCCGTCCACGAAGACCTCAATGGTTCTCTGGATCACATTGACAGCAATGGAAACAGTAGTCCGGGCTGCTCACCTCAGCCGCACATACATTCAATCCACGTCAAGGAAGAGCCAGTGATTGCAGAGGATGAAGACTGCCCAATGTCCTTAGTGACAACAGCTAATCACAGTCCAGAATTAGAAGACGACAGAGAGATTGAAGAAGAGCCTTTATCTGAAGATCT |

| | | | | | | |
|---|---|---|---|---|---|---|
| ERG_DBD_ELK4_IDR | ERG strain | JBP151_ERG_DBD_ELK4_IDR_F | JBP152_ERG_DBD_ELK4_IDR_R | 58 | 65-66 | ATGGACAGTGCTATCACCCTTTGGCAGTTCCTCCTGGAGCTCCTGTCGGACAGCTCCAACTCCAGCTGCATCACCTGGGA<br>AGGCACCAACGGGGAGTTCAAGATGACGGATCCCGACGAGGTGGCCCGGCGCTGGGGAGAGCGGAAGAGCAAACCCAACA<br>TGAACTACGATAAGCTCAGCCGCGCCCTCCGTTACTACTATGACAAGAACATCATGACCAAGGTCCATGGGAAGCGCTAC<br>GCCTACAAGTTCGACTATCCAGAGATTTTGAACATGGATCCAATGACAGTGGGCAGGATTGAGGGTGACTGTGAAAGTTT<br>AAACTTCAGTGAAGTCAGCAGCAGTTCCAAAGATGTGGAGAATGGAGGGAAAGATAAACCACCTCAGCCTGGTGCCAAGA<br>CCTCTAGCCGCAATGACTACATACACTCTGGCTTATATTCTTCATTTACTCTCAACTCTTTGAACTCCTCCAATGTAAAG<br>CTTTTCAAATTGATAAAGACTGAGAATCCAGCCGAGAAACTGGCAGAGAAAAAATCTCCTCAGGAGCCCACACCATCTGT<br>CATCAAATTTGTCACGACACCTTCCAAAAAGCCACCGGTTGAACCTGTTGCTGCCACCATTTCAATTGGCCCAAGTATTT<br>CTCCATCTTCAGAAGAAACTATCCAAGCTTTGGAGACATTGGTTTCCCCAAAACTGCCTTCCCTGGAAGCCCCAACCTCT<br>GCCTCTAACGTAATGACTGCTTTTGCCACCACACCACCCATTTCGTCCATACCCCCTTTGCAGGAACCTCCCAGAACACC<br>TTCACCACCACTGAGTTCTCACCCAGACATCGACACAGACATTGATTCAGTGGCTTCTCAGCCAATGGAACTTCCAGAGA<br>ATTTGTCACTGGAGCCTAAAGACCAGGATTCAGTCTTGCTAGAAAAGGACAAAGTAAATAATTCATCAAGATCCAAGAAA<br>CCCAAAGGGTTAGAACTGGCACCCACCCTTGTGATCACGAGCAGTGATCCAAGCCCACTGGGAATACTGAGCCCATCTCT<br>CCCTACAGCTTCTCTTACACCAGCATTTTTTTCACAGGTAGCTTGCTCGCTCTTTATGGTGTCACCATTGCTTTCATTTA<br>TTTGCCCTTTTAAGCAAATCCAGAATTTATACACTCAAGTTTGCTTTCTGTTACTTAGGTTTGTCTTAGAAAGGTTATGT<br>GTGACTGTCATGTAG |
| ERG_DBD_ELK1_IDR | ERG strain | JBP153_ERG_DBD_ELK1_IDR_F | JBP154_ERG_DBD_ELK1_IDR_R | 58 | 67-68 | ATGGACCCATCTGTGACGCTTTGGCAGTTCCTCCTGGAGCTCCTGTCGGACAGCTCCAACTCCAGCTGCATCACCTGGGA<br>AGGCACCAACGGGGAGTTCAAGATGACGGATCCCGACGAGGTGGCCCGGCGCTGGGGAGAGCGGAAGAGCAAACCCAACA<br>TGAACTACGATAAGCTCAGCCGCGCCCTCCGTTACTACTATGACAAGAACATCATGACCAAGGTCCATGGGAAGCGCTAC<br>GCCTACAAGTTCGACTACCCTGAGGTCGCAGGGTGCTCCACTGAGGACTGCCCGCCCCAGCCAGAGGTGTCTGTTACCTC<br>CACCATGCCAAATGTGGCCCCTGCTGCTATACATGCCGCCCCAGGGGACACTGTCTCTGGAAAGCCAGGCACACCCAAGG<br>GTGCAGGAATGGCAGGCCCAGGCGGTTTGGCACGCAGCAGCCGGAACGAGTACATGCGCTCGGGCCTCTATTCCACCTTC<br>ACCATCCAGTCTCTGCAGCCGCAGCCACCCCCTCATCCTCGGCCTGCTGTGGTGCTCCCCAGTGCAGCTCCTGCAGGGGC<br>AGCAGCGCCCCCCTCGGGGAGCAGGAGCACCAGTCCAAGCCCCTTGGAGGCCTGTCTGGAGGCTGAAGAGGCCGGCTTGC<br>CTCTGCAGGTCATCCTGACCCCGCCCGAGGCCCCAAACCTGAAATCGGAAGAGCTTAATGTGGAGCCGGGTTTGGCCGG<br>GCTTTGCCCCCAGAAGTGAAAGTAGAAGGGCCCAAGGAAGAGTTGGAAGTTGCGGGGGAGAGAGGGTTTGTGCCAGAAAC<br>CACCAAGGCCGAGCCAGAAGTCCCTCCACAGGAGGGCGTGCCAGCCCGGCTGCCCGCGGTTGTTATGGACACCGCAGGGC<br>AGGCGGGCGGCCATGCGGCTTCCAGCCCTGAGATCTCCCAGCCGCAGAAGGGCCGGAAGCCCCGGGACCTAGAGCTTCCA<br>CTCAGCCCGAGCCTGCTAGGTGGGCCGGGACCCGAACGGACCCCAGGATCGGGAAGTGGCTCCGGCCTCCAGGCTCCGGG<br>GCCGGCGCTGACCCCATCCCTGCTTCCTACGCATACATTGACCCCGGTGCTGCTGACACCCAGCTCGCTGCCTCCTAGCA<br>TTCACTTCTGGAGCACCCTGAGTCCCATTGCGCCCCGTAGCCCGGCCAAGCTCTCCTTCCAGTTTCCATCCAGTGGCAGC<br>GCCCAGGTGCACATCCCTTCTATCAGCGTGGATGGCCTCTCGACCCCCGTGGTGCTCTCCCCAGGGCCCCAGAAGCCATA<br>G |
| ERG_DBD_FLI1_IDR | ERG strain | JBP155_ERG_DBD_FLI1_IDR_F | JBP156_ERG_DBD_FLI1_IDR_R | N/A | 71-72 | ATGGACGGGACTATTAAGGAGGCTCTGTCGGTGGTGAGCGACGACCAGTCCCTCTTTGACTCAGCGTACGGAGCGGCAGC<br>CCATCTCCCCAAGGCCGACATGACTGCCTCGGGGAGTCCTGACTACGGGCAGCCCCACAAGATCAACCCCCTCCCACCAC<br>AGCAGGAGTGGATCAATCAGCCAGTGAGGGTCAACGTCAAGCGGGAGTATGACCACATGAATGGATCCAGGGAGTCTCCG<br>GTGGACTGCAGCGTTAGCAAATGCAGCAAGCTGGTGGGCGGAGGCGAGTCCAACCCCATGAACTACAACAGCTATATGGA<br>CGAGAAGAATGGCCCCCCTCCTCCCAACATGACCACCAACGAGAGGAGAGTCATCGTCCCCGCAGACCCCACACTGTGGA<br>CACAGGAGCATGTGAGGCAATGGCTGGAGTGGGCCATAAAGGAGTACAGCTTGATGGAGATCGACACATCCTTTTTCCAG<br>AACATGGATGGCAAGGAACTGTGTAAAATGAACAAGGAGGACTTCCTCCGCGCCACCACCCTCTACAACACGGAAGTGCT<br>GTTGTCACACCTCAGTTACCTCAGGGAAAGTTCACTGCTGGCCTATAATACAACCTCCCACACCGACCAATCCTCACGAT<br>TGAGTGTCAAAGAAGACCCTTCTTATGACTCAGTCAGAAGAGGAGCTTGGGGCAATAACATGAATTCTGGCCTCAACAAA<br>AGTCCTCCCCTTGGAGGGGCACAAACGATCAGTAAGAATACAGAGCAACGGCCCCAGCCAGATCCGTATCAGATCCTGGG<br>CCCGACCAGCAGTCGCCTAGCCAACCCTGGAAGCGGGCAGATCCAGCTTTGGCAGTTCCTCCTGGAGCTCCTGTCGGACA<br>GCTCCAACTCCAGCTGCATCACCTGGGAAGGCACCAACGGGGAGTTCAAGATGACGGATCCCGACGAGGTGGCCCGGCGC<br>TGGGGAGAGCGGAAGAGCAAACCCAACATGAACTACGATAAGCTCAGCCGCGCCCTCCGTTACTACTATGACAAGAACAT<br>CATGACCAAGGTCCATGGGAAGCGCTACGCCTACAAGTTCGACTTCCACGGCATTGCCCAGGCTCTGCAGCCACATCCGA<br>CCGAGTCGTCCATGTACAAGTACCCTTCTGACATCTCCTACATGCCTTCCTACCATGCCCACCAGCAGAAGGTGAACTTT<br>GTCCCTCCCCATCCATCCTCCATGCCTGTCACTTCCTCCAGCTTCTTTGGAGCCGCATCACAATACTGGACCTCCCCCAC<br>GGGGGGAATCTACCCCAACCCCAACGTCCCCCGCCATCCTAACACCCACGTGCCTTCACACTTAGGCAGCTACTACTTG |
| ERG_DBD_ERF_IDR | ERG strain | JBP157_ERG_DBD_ERF_IDR_F | JBP158_ERG_DBD_ERF_IDR_R | 58 | 73-74 | ATGAAGACCCCGGCGGACACAGGGTTTGCCTTCCCGGATTGGGCCTACAAGCCAGAGTCGTCCCCTGGCTCAAGGCAGAT<br>CCAGCTTTGGCAGTTCCTCCTGGAGCTCCTGTCGGACAGCTCCAACTCCAGCTGCATCACCTGGGAAGGCACCAACGGGG<br>AGTTCAAGATGACGGATCCCGACGAGGTGGCCCGGCGCTGGGGAGAGCGGAAGAGCAAACCCAACATGAACTACGATAAG<br>CTCAGCCGCGCCCTCCGTTACTACTATGACAAGAACATCATGACCAAGGTCCATGGGAAGCGCTACGCCTACAAGTTCAA<br>TTTCAACAAACTGGTGCTGGTCAATTACCCATTCATTGATGTGGGGTTGGCTGGGGGTGCAGTGCCCCAGAGTGCCCCGC<br>CAGTGCCGTCGGGTGGTAGCCACTTCCGCTTCCCTCCCTCAACGCCCTCCGAGGTGCTGTCCCCCACCGAGGACCCCCGC<br>TCACCACCAGCCTGCTCTTCATCTTCATCTTCCCTCTTCTCGGCTGTGGTGGCCCGCCGCCTGGGCCGAGGCTCAGTCAG<br>TGACTGTAGTGATGGCACGTCAGAGCTGGAGGAACCGCTGGGAGAGGATCCCCGCGCCCGACCACCCGGCCCTCCGGATC<br>TGGGTGCCTTCCGAGGGCCCCCGCTGGCCCGCCTGCCCCATGACCCTGGTGTCTTCCGAGTCTATCCCCGGCCTCGGGGT<br>GGCCCTGAACCCCTCAGCCCCTTCCCTGTGTCGCCTCTGGCCGGTCCTGGATCCCTGCTGCCCCCTCAGCTCTCCCCGGC<br>TCTGCCCATGACGCCCACCCACCTGGCCTACACTCCCTCGCCCACGCTGAGCCCGATGTACCCCAGTGGTGGCGGGGGGC<br>CCAGCGGCTCAGGGGGAGGCTCCCACTTCTCCTTCAGCCCTGAGGACATGAAACGGTACCTGCAGGCCCACACCCAAAGC<br>GTCTACAACTACCACCTCAGCCCCCGCGCCTTCCTGCACTACCCTGGGCTGGTGGTGCCCCAGCCCCAGCGCCCTGACAA<br>GTGCCCGCTGCCGCCCATGGCACCCGAGACCCCACCGGTCCCCTCCTCGGCCTCGTCATCCTCTTCTTCTTCTTCCTCCC<br>CATTCAAGTTTAAGCTCCAGCCGCCCCCACTCGGACGCCGGCAGCGGGCAGCTGGGGAGAAGGCCGTAGCCGGTGCTGAC<br>AAGAGCGGTGGCAGTGCAGGCGGGCTGGCTGAGGGGGCAGGGGCGCTAGCCCCACCGCCCCCGCCACCACAGATCAAGGT<br>GGAGCCCATCTCGGAAGGCGAGTCGGAGGAGGTAGAGGTGACTGACATCAGTGATGAGGATGAGGAAGACGGGGAGGTGT<br>TCAAGACGCCCCGTGCCCCACCTGCACCCCCTAAGCCTGAGCCCGGCGAGGCACCCGGGGCATCCCAGTGCATGCCCCTC<br>AAGCTACGCTTTAAGCGGCGCTGGAGTGAAGACTGTCGCCTCGAAGGGGGTGGGGGCCCCGCTGGGGGCTTTGAGGATGA<br>GGGTGAGGACAAGAAGGTGCGTGGGGAGGGGCCTGGGGAGGCTGGGGGGCCCCTCACCCCAAGGCGGGTGAGCTCTGACC<br>TCCAGCATGCCACGGCCCAGCTCTCCCTGGAGCACCGAGACTCCTAG |
| ERG_DBD_ELF1_IDR | ERG strain | JBP159_ERG_DBD_ELF1_IDR_F | JBP160_ERG_DBD_ELF1_IDR_R | 58 | 75-76 | ATGGCTGCTGTTGTCCAACAGAACGACCTAGTATTTGAATTTGCTAGTAACGTCATGGAGGATGAACGACAGCTTGGTGA<br>TCCAGCTATTTTTCCTGCCGTAATTGTGGAACATGTTCCTGGTGCTGATATTCTCAATAGTTATGCCGGTCTAGCCTGTG<br>TGGAAGAGCCCAATGACATGATTACTGAGAGTTCACTGGATGTTGCTGAAGAAGAAATCATAGACGATGATGATGATGAC<br>ATCACCCTTACAGTTGAAGCTTCTTGTCATGACGGGGATGAAACAATTGAAACTATTGAGGCTGCTGAGGCACTCCTCAA<br>TATGGATTCCCCTGGCCCTATGCTGGATGAAAAACGAATAAATAATAATATATTTAGTTCACCTGAAGATGACATGGTTG<br>TTGCCCCAGTCACCCATGTGTCCGTCACATTAGATGGGATTCCTGAAGTGATGGAAACACAGCAGGTGCAAGAAAAATAT<br>GCAGACTCACCGGGAGCCTCATCACCAGAACAGCCTAAGAGGAAAAAAGGAAGAAAAACTAAACCACCACGACCAGATTC<br>CCCAGCCACTACGCCAAATATATCTGTGAAGAAGAAAAACAAAGATGGAAAGGGAAACACAATTTATCTTCTTTGGCAGT<br>TCCTCCTGGAGCTCCTGTCGGACAGCTCCAACTCCAGCTGCATCACCTGGGAAGGCACCAACGGGGAGTTCAAGATGACG<br>GATCCCGACGAGGTGGCCCGGCGCTGGGGAGAGCGGAAGAGCAAACCCAACATGAACTACGATAAGCTCAGCCGCGCCCT<br>CCGTTACTACTATGACAAGAACATCATGACCAAGGTCCATGGGAAGCGCTACGCCTACAAGTTCGACAAAGAAATGCCAA<br>AAGATCTTATATATATAAATGATGAGGATCCAAGTTCCAGCATAGAGTCTTCAGATCCATCACTATCTTCATCAGCCACT<br>TCAAATAGGAATCAAACCAGCCGGTCGAGAGTATCTTCAAGTCCAGGGGTAAAAGGAGGAGCCACTACAGTTCTAAAACC<br>AGGGAATTCTAAAGCTGCAAAACCCAAAGATCCTGTGGAAGTTGCACAACCATCAGAAGTTTTGAGGACAGTGCAGCCCA<br>CGCAGTCTCCATATCCTACCCAGCTCTTCCGGACTGTTCATGTAGTACAGCCAGTACAGGCTGTCCCAGAGGGAGAAGCA<br>GCTAGAACCAGTACCATGCAGGATGAAACATTAAATTCTTCCGTTCAGAGTATTAGGACTATACAGGCTCCAACCCAAGT<br>TCCAGTGGTTGTGTCTCCTAGGAATCAGCAGTTGCATACAGTAACACTCCAAACAGTGCCACTCACAACAGTTATAGCCA<br>GCACAGATCCATCAGCAGGTACTGGATCTCAGAAGTTTATTTTACAAGCCATTCCATCATCACAGCCCATGACAGTACTG<br>AAAGAAAATGTCATGCTGCAGTCACAAAAGGCGGGCTCTCCTCCTTCAATTGTCTTGGGCCCTGCCCAGGTTCAGCAGGT<br>CCTTACTAGCAATGTTCAGACCATTTGCAATGGAACCGTCAGTGTGGCTTCCTCTCCATCCTTCAGTGCTACTGCACCTG<br>TGGTGACCTTTTCTCCTCGCAGTTCACAGCTGGTTGCTCACCCACCTGGCACTGTAATCACTTCAGTTATCAAAACTCAA<br>GAAACAAAAACTCTTACACAGGAAGTAGAGAAAAAGGAATCTGAAGATCATTTGAAAGAGAACACTGAGAAAACGGAGCA<br>GCAGCCACAGCCTTATGTGATGGTAGTGTCCAGTTCCAATGGATTTACTTCTCAGGTAGCTATGAAACAAAACGAACTGC<br>TGGAACCCAACTCTTTTTAG |
| ERG_DBD_ELF2_IDR | ERG strain | JBP161_ERG_DBD_ELF2_IDR_F | JBP162_ERG_DBD_ELF2_IDR_R | 58 | 77-78 | ATGACATCAGCAGTGGTTGACAGTGGAGGTACTATTTTGGAGCTTTCCAGCAATGGAGTAGAAAATCAAGAGGAAAGTGA<br>AAAGGTTTCTGAATATCCAGCAGTGATTGTGGAGCCAGTTCCAAGTGCCAGATTAGAGCAGGGCTATGCAGCCCAGGTTC<br>TGGTTTATGATGATGAGACTTATATGATGCAAGATGTGGCAGAAGAACAAGAAGTTGAGACAGAGAATGTGGAAACAGTG<br>GAAGCATCAGTTCACAGCAGTAATGCACACTGTACAGATAAGACAATTGAAGCTGCTGAAGCCCTGCTTCATATGGAATC<br>TCCTACCTGCTTGAGGGATTCAAGAAGTCCTGAATTCATCCATGCTGCTATGAGGCCAGATGTCATTACAGAAACTGTAG<br>TGGAGGTGTCAACTGAAGAGTCTGAACCCATGGATACCTCTCCTATTCCAACATCACCAGATAGCCATGAACCAATGAAA<br>AAGAAAAAAGTTGGCCGTAAACCAAAGACCCAGCAATCACCAATTTCCAATGGCTCTCCTGAGTTAGGTATAAAGAAGAA<br>ACCAAGAGAAGGAAAAGGAAACACAACCTATTTGCTTTGGCAGTTCCTCCTGGAGCTCCTGTCGGACAGCTCCAACTCCA<br>GCTGCATCACCTGGGAAGGCACCAACGGGGAGTTCAAGATGACGGATCCCGACGAGGTGGCCCGGCGCTGGGGAGAGCGG<br>AAGAGCAAACCCAACATGAACTACGATAAGCTCAGCCGCGCCCTCCGTTACTACTATGACAAGAACATCATGACCAAGGT<br>CCATGGGAAGCGCTACGCCTACAAGTTCGACGATATGCCGAAAAACATAGTGGTCATAGATGATGACAAAAGTGAAACCT<br>GTAATGAAGATTTAGCAGGAACTACTGATGAAAAATCATTAGAACGAGTGTCACTGTCTGCAGAAAGTCTCCTGAAAGCA<br>GCATCCTCTGTTCGCAGTGGAAAAAATTCATCCCCTATAAACTGCTCCAGAGCAGAGAAGGGTGTAGCTAGAGTTGTGAA<br>TATCACTTCCCCTGGGCACGATGCTTCATCCAGATCTCCTACTACCACTGCATCTGTGTCAGCAACAGCAGCTCCAAGGA<br>CAGTTCGTGTGGCAATGCAGGTACCTGTTGTAATGACATCATTGGGTCAGAAAATTTCAACTGTGGCAGTTCAGTCAGTT<br>AATGCAGGCGCACCATTAATAACCAGCACTAGTCCAACAACAGCGACCTCTCCAAAGGTAGTCATTCAGACAATCCCTAC<br>TGTGATGCCAGCTTCTACTGAAAATGGAGACAAAATCACCATGCAGCCTGCCAAAATTATTACCATCCCAGCTACACAGC<br>TTGCACAGTGTCAACTGCAGACAAAGTCAAATCTGACTGGATCAGGAAGCATTAACATTGTTGGAACCCCATTGGCTGTG<br>AGAGCACTTACCCCTGTTTCAATAGCCCATGGTACACCTGTAATGAGACTATCAATGCCTACTCAGCAGGCATCTGGCCA<br>GACTCCTCCTCGAGTTATCAGTGCAGTCATAAAGGGGCCAGAGGTTAAATCGGAAGCAGTGGCAAAAAAGCAAGAACATG<br>ATGTGAAAACTTTGCAGCTAGTAGAAGAAAAACCAGCAGATGGAAATAAGACAGTGACCCACGTAGTGGTTGTCAGTGCG<br>CCTTCAGCTATTGCCCTTCCTGTAACTATGAAAACAGAAGGACTAGTGACATGTGAGAAA |
| ELF1_DBD_ELK4_IDR | ELF1 strain | JBP165_ELF1_DBD_ELK4_IDR_F | JBP166_ELF1_DBD_ELK4_IDR_R | N/A | 65-66 | ATGGACAGTGCTATCACCTGGGAGTTTTTACTGGCACTGCTCCAGGACAAGGCTACTTGTCCTAAATACATCAAGTGGAC<br>CCAGCGAGAGAAAGGCATTTTTAAATTGGTGGATTCTAAAGCAGTGTCCAGGTTGTGGGGGAAGCACAAAAACAAACCTG<br>ATATGAATTATGAGACCATGGGAAGAGCACTCAGGTACTATTACCAAAGGGGTATTCTGGCAAAAGTGGAAGGTCAGCGC<br>TTGGTGTATCAGTT------TTATCCAGAGATTTTGAACATGGATCCAATGACAGTGGGCAGGATTGAGGGTGACTGTGA<br>AAGTTTAAACTTCAGTGAAGTCAGCAGCAGTTCCAAAGATGTGGAGAATGGAGGGAAAGATAAACCACCTCAGCCTGGTG<br>CCAAGACCTCTAGCCGCAATGACTACATACACTCTGGCTTATATTCTTCATTTACTCTCAACTCTTTGAACTCCTCCAAT<br>GTAAAGCTTTTCAAATTGATAAAGACTGAGAATCCAGCCGAGAAACTGGCAGAGAAAAAATCTCCTCAGGAGCCCACACC<br>ATCTGTCATCAAATTTGTCACGACACCTTCCAAAAAGCCACCGGTTGAACCTGTTGCTGCCACCATTTCAATTGGCCCAA<br>GTATTTCTCCATCTTCAGAAGAAACTATCCAAGCTTTGGAGACATTGGTTTCCCCAAAACTGCCTTCCCTGGAAGCCCCA<br>ACCTCTGCCTCTAACGTAATGACTGCTTTTGCCACCACACCACCCATTTCGTCCATACCCCCTTTGCAGGAACCTCCCAG<br>AACACCTTCACCACCACTGAGTTCTCACCCAGACATCGACACAGACATTGATTCAGTGGCTTCTCAGCCAATGGAACTTC<br>CAGAGAATTTGTCACTGGAGCCTAAAGACCAGGATTCAGTCTTGCTAGAAAAGGACAAAGTAAATAATTCATCAAGATCC<br>AAGAAACCCAAAGGGTTAGAACTGGCACCCACCCTTGTGATCACGAGCAGTGATCCAAGCCCACTGGGAATACTGAGCCC<br>ATCTCTCCCTACAGCTTCTCTTACACCAGCATTTTTTTCACAGGTAGCTTGCTCGCTCTTTATGGTGTCACCATTGCTTT<br>CATTTATTTGCCCTTTTAAGCAAATCCAGAATTTATACACTCAAGTTTGCTTTCTGTTACTTAGGTTTGTCTTAGAAAGG<br>TTATGTGTGACTGTCATGTAG |

| | | | | | | |
|---|---|---|---|---|---|---|
| ELF1_DBD_ELK1_ID | ELF1 strain | JBP167_ELF1_DBD_ELK1_IDR_F | JBP168_ELF1_DBD_ELK1_IDR_R | 58 | 67-68 | ATGGACCCATCTGTGACGTGGGAGTTTTTACTGGCACTGCTCCAGGACAAGGCTACTTGTCCTAAATACATCAAGTGGAC<br>CCAGCGAGAGAAAGGCATTTTTAAATTGGTGGATTCTAAAGCAGTGTCCAGGTTGTGGGGGAAGCACAAAAACAAACCTG<br>ATATGAATTATGAGACCATGGGAAGAGCACTCAGGTACTATTACCAAAGGGGTATTCTGGCAAAAGTGGAAGGTCAGCGC<br>TTGGTGTATCAGTTTTACCCTGAGGTCGCAGGGTGCTCCACTGAGGACTGCCCGCCCCAGCCAGAGGTGTCTGTTACCTC<br>CACCATGCCAAATGTGGCCCCTGCTGCTATACATGCCGCCCCAGGGGACACTGTCTCTGGAAAGCCAGGCACACCCAAGG<br>GTGCAGGAATGGCAGGCCCAGGCGGTTTGGCACGCAGCAGCCGGAACGAGTACATGCGCTCGGGCCTCTATTCCACCTTC<br>ACCATCCAGTCTCTGCAGCCGCAGCCACCCCCTCATCCTCGGCCTGCTGTGGTGCTCCCCAGTGCAGCTCCTGCAGGGGC<br>AGCAGCGCCCCCCTCGGGGAGCAGGAGCACCAGTCCAAGCCCCTTGGAGGCCTGTCTGGAGGCTGAAGAGGCCGGCTTGC<br>CTCTGCAGGTCATCCTGACCCCGCCCGAGGCCCCAAACCTGAAATCGGAAGAGCTTAATGTGGAGCCGGGTTTGGGCCGG<br>GCTTTGCCCCCAGAAGTGAAAGTAGAAGGGCCCAAGGAAGAGTTGGAAGTTGCGGGGGAGAGAGGGTTTGTGCCAGAAAC<br>CACCAAGGCCGAGCCAGAAGTCCCTCCACAGGAGGGCGTGCCAGCCCGGCTGCCCGCGGTTGTTATGGACACCGCAGGGC<br>AGGCGGGCGGCCATGCGGCTTCCAGCCCTGAGATCTCCCAGCCGCAGAAGGGCCGGAAGCCCCGGGACCTAGAGCTTCCA<br>CTCAGCCCGAGCCTGCTAGGTGGGCCGGGACCCGAACGGACCCCAGGATCGGGAAGTGGCTCCGGCCTCCAGGCTCCGGG<br>GCCGGCGCTGACCCCATCCCTGCTTCCTACGCATACATTGACCCCGGTGCTGCTGACACCCAGCTCGCTGCCTCCTAGCA<br>TTCACTTCTGGAGCACCCTGAGTCCCATTGCGCCCCGTAGCCCGGCCAAGCTCTCCTTCCAGTTTCCATCCAGTGGCAGC<br>GCCCAGGTGCACATCCCTTCTATCAGCGTGGATGGCCTCTCGACCCCCGTGGTGCTCTCCCCAGGGCCCCAGAAGCCATA<br>G |
| ELF1_DBD_FLI1_IDR | ELF1 strain | JBP169_ELF1_DBD_FLI1_IDR_F | JBP170_ELF1_DBD_FLI1_IDR_R | 58 | 71-72 | ATGGACGGGACTATTAAGGAGGCTCTGTCGGTGGTGAGCGACGACCAGTCCCTCTTTGACTCAGCGTACGGAGCGGCAGC<br>CCATCTCCCCAAGGCCGACATGACTGCCTCGGGGAGTCCTGACTACGGGCAGCCCCACAAGATCAACCCCCTCCCACCAC<br>AGCAGGAGTGGATCAATCAGCCAGTGAGGGTCAACGTCAAGCGGGAGTATGACCACATGAATGGATCCAGGGAGTCTCCG<br>GTGGACTGCAGCGTTAGCAAATGCAGCAAGCTGGTGGGCGGAGGCGAGTCCAACCCCATGAACTACAACAGCTATATGGA<br>CGAGAAGAATGGCCCCCCTCCTCCCAACATGACCACCAACGAGAGGAGAGTCATCGTCCCCGCAGACCCCACACTGTGGA<br>CACAGGAGCATGTGAGGCAATGGCTGGAGTGGGCCATAAAGGAGTACAGCTTGATGGAGATCGACACATCCTTTTTCCAG<br>AACATGGATGGCAAGGAACTGTGTAAAATGAACAAGGAGGACTTCCTCCGCGCCACCACCCTCTACAACACGGAAGTGCT<br>GTTGTCACACCTCAGTTACCTCAGGGAAAGTTCACTGCTGGCCTATAATACAACCTCCCACACCGACCAATCCTCACGAT<br>TGAGTGTCAAAGAAGACCCTTCTTATGACTCAGTCAGAAGAGGAGCTTGGGGCAATAACATGAATTCTGGCCTCAACAAA<br>AGTCCTCCCCTTGGAGGGGCACAAACGATCAGTAAGAATACAGAGCAACGGCCCCAGCCAGATCCGTATCAGATCCTGGG<br>CCCGACCAGCAGTCGCCTAGCCAACCCTGGAAGCGGGCAGATCCAGTGGGAGTTTTTACTGGCACTGCTCCAGGACAAGG<br>CTACTTGTCCTAAATACATCAAGTGGACCCAGCGAGAGAAAGGCATTTTTAAATTGGTGGATTCTAAAGCAGTGTCCAGG<br>TTGTGGGGGAAGCACAAAAACAAACCTGATATGAATTATGAGACCATGGGAAGAGCACTCAGGTACTATTACCAAAGGGG<br>TATTCTGGCAAAAGTGGAAGGTCAGCGCTTGGTGTATCAGTTTTTCCACGGCATTGCCCAGGCTCTGCAGCCACATCCGA<br>CCGAGTCGTCCATGTACAAGTACCCTTCTGACATCTCCTACATGCCTTCCTACCATGCCCACCAGCAGAAGGTGAACTTT<br>GTCCCTCCCCATCCATCCTCCATGCCTGTCACTTCCTCCAGCTTCTTTGGAGCCGCATCACAATACTGGACCTCCCCCAC<br>GGGGGGAATCTACCCCAACCCCAACGTCCCCCGCCATCCTAACACCCACGTGCCTTCACACTTAGGCAGCTACTACTTG |
| ELF1_DBD_ERF_IDR | ELF1 strain | JBP171_ELF1_DBD_ERF_IDR_F | JBP172_ELF1_DBD_ERF_IDR_R | 58 | 73-74 | ATGAAGACCCCGGCGGACACAGGGTTTGCCTTCCCGGATTGGGCCTACAAGCCAGAGTCGTCCCCTGGCTCAAGGCAGAT<br>CCAGTGGGAGTTTTTACTGGCACTGCTCCAGGACAAGGCTACTTGTCCTAAATACATCAAGTGGACCCAGCGAGAGAAAG<br>GCATTTTTAAATTGGTGGATTCTAAAGCAGTGTCCAGGTTGTGGGGGAAGCACAAAAACAAACCTGATATGAATTATGAG<br>ACCATGGGAAGAGCACTCAGGTACTATTACCAAAGGGGTATTCTGGCAAAAGTGGAAGGTCAGCGCTTGGTGTATCAGTT<br>TTTCAACAAACTGGTGCTGGTCAATTACCCATTCATTGATGTGGGGTTGGCTGGGGGTGCAGTGCCCCAGAGTGCCCCGC<br>CAGTGCCGTCGGGTGGTAGCCACTTCCGCTTCCCTCCCTCAACGCCCTCCGAGGTGCTGTCCCCCACCGAGGACCCCCGC<br>TCACCACCAGCCTGCTCTTCATCTTCATCTTCCCTCTTCTCGGCTGTGGTGGCCCGCCGCCTGGGCCGAGGCTCAGTCAG<br>TGACTGTAGTGATGGCACGTCAGAGCTGGAGGAACCGCTGGGAGAGGATCCCCGCGCCCGACCACCCGGCCCTCCGGATC<br>TGGGTGCCTTCCGAGGGCCCCCGCTGGCCCGCCTGCCCCATGACCCTGGTGTCTTCCGAGTCTATCCCCGGCCTCGGGGT<br>GGCCCTGAACCCCTCAGCCCCTTCCCTGTGTCGCCTCTGGCCGGTCCTGGATCCCTGCTGCCCCCTCAGCTCTCCCCGGC<br>TCTGCCCATGACGCCCACCCACCTGGCCTACACTCCCTCGCCCACGCTGAGCCCGATGTACCCCAGTGGTGGCGGGGGGC<br>CCAGCGGCTCAGGGGGAGGCTCCCACTTCTCCTTCAGCCCTGAGGACATGAAACGGTACCTGCAGGCCCACACCCAAAGC<br>GTCTACAACTACCACCTCAGCCCCCGCGCCTTCCTGCACTACCCTGGGCTGGTGGTGCCCCAGCCCCAGCGCCCTGACAA<br>GTGCCCGCTGCCGCCCATGGCACCCGAGACCCCACCGGTCCCCTCCTCGGCCTCGTCATCCTCTTCTTCTTCTTCCTCCC<br>CATTCAAGTTTAAGCTCCAGCCGCCCCCACTCGGACGCCGGCAGCGGGCAGCTGGGGAGAAGGCCGTAGCCGGTGCTGAC<br>AAGAGCGGTGGCAGTGCAGGCGGGCTGGCTGAGGGGGCAGGGGCGCTAGCCCCACCGCCCCCGCCACCACAGATCAAGGT<br>GGAGCCCATCTCGGAAGGCGAGTCGGAGGAGGTAGAGGTGACTGACATCAGTGATGAGGATGAGGAAGACGGGGAGGTGT<br>TCAAGACGCCCCGTGCCCCACCTGCACCCCCTAAGCCTGAGCCCGGCGAGGCACCCGGGGCATCCCAGTGCATGCCCCTC<br>AAGCTACGCTTTAAGCGGCGCTGGAGTGAAGACTGTCGCCTCGAAGGGGGTGGGGGCCCCGCTGGGGGCTTTGAGGATGA<br>GGGTGAGGACAAGAAGGTGCGTGGGGAGGGGCCTGGGGAGGCTGGGGGGCCCCTCACCCCAAGGCGGGTGAGCTCTGACC<br>TCCAGCATGCCACGGCCCAGCTCTCCCTGGAGCACCGAGACTCCTAG |
| ELF1_DBD_ELF2_ID | ELF1 strain | JBP173_ELF1_DBD_ELF2_IDR_F | JBP174_ELF1_DBD_ELF2_IDR_R | 58 | 77-78 | ATGACATCAGCAGTGGTTGACAGTGGAGGTACTATTTTGGAGCTTTCCAGCAATGGAGTAGAAAATCAAGAGGAAAGTGA<br>AAAGGTTTCTGAATATCCAGCAGTGATTGTGGAGCCAGTTCCAAGTGCCAGATTAGAGCAGGGCTATGCAGCCCAGGTTC<br>TGGTTTATGATGATGAGACTTATATGATGCAAGATGTGGCAGAAGAACAAGAAGTTGAGACAGAGAATGTGGAAACAGTG<br>GAAGCATCAGTTCACAGCAGTAATGCACACTGTACAGATAAGACAATTGAAGCTGCTGAAGCCCTGCTTCATATGGAATC<br>TCCTACCTGCTTGAGGGATTCAAGAAGTCCTGAATTCATCCATGCTGCTATGAGGCCAGATGTCATTACAGAAACTGTAG<br>TGGAGGTGTCAACTGAAGAGTCTGAACCCATGGATACCTCTCCTATTCCAACATCACCAGATAGCCATGAACCAATGAAA<br>AAGAAAAAAGTTGGCCGTAAACCAAAGACCCAGCAATCACCAATTTCCAATGGCTCTCCTGAGTTAGGTATAAAGAAGAA<br>ACCAAGAGAAGGAAAAGGAAACACAACCTATTTGTGGGAGTTTTTACTGGCACTGCTCCAGGACAAGGCTACTTGTCCTA<br>AATACATCAAGTGGACCCAGCGAGAGAAAGGCATTTTTAAATTGGTGGATTCTAAAGCAGTGTCCAGGTTGTGGGGGAAG<br>CACAAAAACAAACCTGATATGAATTATGAGACCATGGGAAGAGCACTCAGGTACTATTACCAAAGGGGTATTCTGGCAAA<br>AGTGGAAGGTCAGCGCTTGGTGTATCAGTTTGATATGCCGAAAAACATAGTGGTCATAGATGATGACAAAAGTGAAACCT<br>GTAATGAAGATTTAGCAGGAACTACTGATGAAAAATCATTAGAACGAGTGTCACTGTCTGCAGAAAGTCTCCTGAAAGCA<br>GCATCCTCTGTTCGCAGTGGAAAAAATTCATCCCCTATAAACTGCTCCAGAGCAGAGAAGGGTGTAGCTAGAGTTGTGAA<br>TATCACTTCCCCTGGGCACGATGCTTCATCCAGATCTCCTACTACCACTGCATCTGTGTCAGCAACAGCAGCTCCAAGGA<br>CAGTTCGTGTGGCAATGCAGGTACCTGTTGTAATGACATCATTGGGTCAGAAAATTTCAACTGTGGCAGTTCAGTCAGTT<br>AATGCAGGCGCACCATTAATAACCAGCACTAGTCCAACAACAGCGACCTCTCCAAAGGTAGTCATTCAGACAATCCCTAC<br>TGTGATGCCAGCTTCTACTGAAAATGGAGACAAAATCACCATGCAGCCTGCCAAAATTATTACCATCCCAGCTACACAGC<br>TTGCACAGTGTCAACTGCAGACAAAGTCAAATCTGACTGGATCAGGAAGCATTAACATTGTTGGAACCCCATTGGCTGTG<br>AGAGCACTTACCCCTGTTTCAATAGCCCATGGTACACCTGTAATGAGACTATCAATGCCTACTCAGCAGGCATCTGGCCA<br>GACTCCTCCTCGAGTTATCAGTGCAGTCATAAAGGGGCCAGAGGTTAAATCGGAAGCAGTGGCAAAAAAGCAAGAACATG<br>ATGTGAAAACTTTGCAGCTAGTAGAAGAAAAACCAGCAGATGGAAATAAGACAGTGACCCACGTAGTGGTTGTCAGTGCG<br>CCTTCAGCTATTGCCCTTCCTGTAACTATGAAAACAGAAGGACTAGTGACATGTGAGAAA |
| ELF1_DBD_ERG_IDR | ELF1 strain | JBP175_ELF1_DBD_ERG_IDR_F | JBP176_ELF1_DBD_ERG_IDR_R | 65 | 69-70 | ATGGTCAGCACTATTAAGGAAGCCTTATCAGTTGTGAGTGAGGACCAGTCGTTGTTTGAGTGTGCCTACGGAACGCCACA<br>CCTGGCTAAGACAGAGATGACCGCGTCCTCCTCCAGCGACTATGGACAGACTTCCAAGATGAGCCCACGCGTCCCTCAGC<br>AGGATTGGCTGTCTCAACCCCCAGCCAGGGTCACCATCAAAATGGAATGTAACCCTAGCCAGGTGAATGGCTCAAGGAAC<br>TCTCCTGATGAATGCAGTGTGGCCAAAGGCGGGAAGATGGTGGGCAGCCCAGACACCGTTGGGATGAACTACGGCAGCTA<br>CATGGAGGAGAAGCACATGCCACCCCCAAACATGACCACGAACGAGCGCAGAGTTATCGTGCCAGCAGATCCTACGCTAT<br>GGAGTACAGACCATGTGCGGCAGTGGCTGGAGTGGGCGGTGAAAGAATATGGCCTTCCAGACGTCAACATCTTGTTATTC<br>CAGAACATCGATGGGAAGGAACTGTGCAAGATGACCAAGGACGACTTCCAGAGGCTCACCCCCAGCTACAATGCCGACAT<br>CCTTCTCTCACATCTCCACTACCTCAGAGAGACTCCTCTTCCACATTTGACTTCAGATGATGTTGATAAAGCCTTACAAA<br>ACTCTCCACGGTTAATGCATGCTAGAAACACAGGGGGTGCAGCTTTTATTTTCCCAAATACTTCAGTATATCCTGAAGCT<br>ACGCAAAGAATTACAACTAGGCCAGATTTACCATATGAGCCCCCCAGGAGATCAGCCTGGACCGGTCACGGCCACCCCAC<br>GCCCCAGTCGAAAGCTGCTCAACCATCTCCTTCCACAGTGCCCAAAACTGAAGACCAGCGTCCTCAGTTAGATCCTTATC<br>AGATTCTTGGACCAACAAGTAGCCGCCTTGCAAATCCAGGCAGTGGCCAGATCCAGTGGGAGTTTTTACTGGCACTGCTC<br>CAGGACAAGGCTACTTGTCCTAAATACATCAAGTGGACCCAGCGAGAGAAAGGCATTTTTAAATTGGTGGATTCTAAAGC<br>AGTGTCCAGGTTGTGGGGGAAGCACAAAAACAAACCTGATATGAATTATGAGACCATGGGAAGAGCACTCAGGTACTATT<br>ACCAAAGGGGTATTCTGGCAAAAGTGGAAGGTCAGCGCTTGGTGTATCAGTTTTTCCACGGGATCGCCCAGGCCCTCCAG<br>CCCCACCCCCCGGAGTCATCTCTGTACAAGTACCCCTCAGACCTCCCGTACATGGGCTCCTATCACGCCCACCCACAGAA<br>GATGAACTTTGTGGCGCCCCACCCTCCAGCCCTCCCCGTGACATCTTCCAGTTTTTTGCTGCCCCAAACCCATACTGGA<br>ATTCACCAACTGGGGGTATATACCCCAACACTAGGCTCCCCACCAGCCATATGCCTTCTCATCTGGGCACTTACTAC |
| SOX5_IDR | JBP093_SOX5_IDR_repair | N/A | N/A | N/A | 79-80 | ATGCTTACTGACCCTGATTTACCTCAGGAGTTTGAAAGGATGTCTTCCAAGCGACCAGCCTCTCCGTATGGGGAAGCAGA<br>TGGAGAGGTAGCCATGGTGACAAGCAGACAGAAAGTGGAAGAAGAGGAGAGTGACGGGCTCCCAGCCTTTCACCTTCCCT<br>TGCATGTGAGTTTTCCCAACAAGCCTCACTCTGAGGAATTTCAGCCAGTTTCTCTGCTGACGCAAGAGACTTGTGGCCAT<br>AGGACTCCCACTTCTCAGCACAATACAATGGAAGTTGATGGCAATAAAGTTATGTCTTCATTTGCCCCACACAACTCATC<br>TACCTCACCTCAGAAGGCAGAAGAAGGTGGGCGACAGAGTGGCGAGTCCTTGTCTAGTACAGCCCTGGGAACTCCTGAAC<br>GGCGCAAGGGCAGTTTAGCTGATGTTGTTGACACCTTGAAGCAGAGGAAAATGGAAGAGCTCATCAAAAACGAGCCGGAA<br>GAAACCCCCAGTATTGAAAAACTACTCTCAAAGGACTGGAAAGACAAGCTTCTTGCAATGGGATCGGGGAACTTTGGCGA<br>AATAAAAGGGACTCCCGAGAGCTTAGCTGAGAAAGAAAGGCAACTCATGGGTATGATCAACCAGCTGACCAGCCTCCGAG<br>AGCAGCTGTTGGCTGCCCACGATGAGCAGAAGAAACTAGCTGCCTCTCAGATTGAGAAACAGCGTCAGCAAATGGAGCTG<br>GCCAAGCAGCAACAAGAACAAATTGCAAGACAGCAGCAGCAGCTTCTACAGCAACAACACAAAATCAATTTGCTCCAGCA<br>ACAGATCCAGGTTCAAGGTCAGCTGCCGCCATTAATGATTCCCGTATTCCCTCCTGATCAACGGACACTGGCTGCAGCTG<br>CCCAGCAAGGATTCCTCCTCCCTCCAGGCTTCAGCTATAAGGCTGGATGTAGTGACCCTTACCCTGTTCAGCTGATCCCA<br>ACTACCATGGCAGCTGCTGCCGCAGCAACACCAGGCTTAGGCCCACTCCAACTGCAGCAGTTATATGCTGCCCAGCTAGC<br>TGCAATGCAGGTATCTCCAGGAGGGAAGCTGCCAGGCATACCCCAAGGCAACCTTGGTGCTGCTGTATCTCCTACCAGCA<br>TTCACACAGACAAGAGCACAAACAGCCCACCACCCAAAAGCAAGGATGAAGTGGCACAGCCACTGAACCTATCAGCTAAA<br>CCCAAGACCTCTGATGGCAAATCACCCACATCACCCACCTCTCCCCATATGCCAGCTCTGAGAATAAACAGTGGGGCAGG<br>CCCCCTCAAAGCCTCTGTCCCAGCAGCGTTAGCTAGTCCTTCAGCCAGAGTTAGCACAATAGGTTACTTAAATGACCATG<br>ATGCTGTCACCAAGGCAATCCAAGAAGCTCGGCAAATGAAGGAGCAACTCCGACGGGAACAACAGGTGCTTGATGGGAAG<br>GTGGCTGTTGTGAATAGTCTGGGTCTTAATAACTGCCGAACAGAAAAGGAAAAAACAACACTGGAGAGTCTGACTCAGCA<br>ACTGGCAGTTAAACAGAATGAAGAAGGAAAATTTAGCCATGCAATGATGGATTTCAATCTGAGTGGAGATTCTGATGGAA<br>GTGCTGGAGTCTCAGAGTCAAGAATTTATAGGGAATCCCGAGGGCGTGGTAGCAATGAACCCCACATAAAGCCCAGGCCA<br>AAGCGCACATGCCTGGTGGATGGCAAAAAGCTGCGCATTGGTGAATACAAGGCAATCATGCGCAACAGGCGGCAGGAAAT<br>GCGGCAGTACTTCAATGTTGGGCAACAAGCACAGATCCCCATTGCCACTGCTGGTGTTGTGTACCCTGGAGCCATCGCCA<br>TGGCTGGGATGCCCTCCCCTCACCTACCCTCGGAGCACTCAAGCGTGTCTAGCAGCCCAGAGCCTGGGATGCCTGTTATC<br>CAGAGCACTTACGGTGTGAAAGGAGAGGAGCCACATATCAAAGAAGAGATACAGGCCGAGGACATCAATGGAGAAATTTA<br>TGATGAGTACGACGAGGAAGAGGATGATCCAGATGTAGATTATGGGAGTGACAGTGAAAACCATATTGCAGGACAAGCCA<br>AC |

| | | | | | | |
|---|---|---|---|---|---|---|
| SOX6_IDR | JBP094_SOX6_IDR_repair | N/A | N/A | N/A | 81-82 | ATGAAAGGTCATGGTGAGCTTCAAGGACATGAAAGAAGAATGTCTTCCAAGCAAGCCACCTCTCCATTTGCCTGTGCAGC<br>TGATGGAGAGGATGCAATGACCCAGGATTTAACCTCAAGGGAAAAGGAAGAGGGCAGTGATCAACATGTGGCCTCCCATC<br>TGCCTCTGCACCCCATAATGCACAACAAACCTCACTCTGAGGAGCTACCAACACTTGTCAGTACCATTCAACAAGATGCT<br>GACTGGGACAGCGTTCTGTCATCTCAGCAAAGAATGGAATCAGAGAATAATAAGTTATGTTCCCTATATTCCTTCCGAAA<br>TACCTCTACCTCACCACATAAGCCTGACGAAGGGAGTCGGGACCGTGAGATAATGACCAGTGTTACTTTTGGAACCCCAG<br>AGCGCCGCAAAGGGAGTCTTGCCGATGTGGTGGACACACTGAAACAGAAGAAGCTTGAGGAAATGACTCGGACTGAACAA<br>GAGGATTCCTCCTGCATGGAAAAACTACTTTCAAAAGATTGGAAGGAAAAAATGGAAAGACTAAATACCAGTGAACTTCT<br>TGGAGAAATTAAAGGTACACCTGAGAGCCTGGCAGAAAAAGAACGGCAGCTCTCCACCATGATTACCCAGCTGATCAGTT<br>TACGGGAGCAGCTACTGGCAGCGCATGATGAACAGAAAAAACTGGCAGCGTCACAAATTGAGAAACAACGGCAGCAAATG<br>GACCTTGCTCGCCAACAGCAAGAACAGATTGCGAGACAACAGCAGCAACTTCTGCAACAGCAGCACAAAATTAATCTCCT<br>GCAGCAACAGATCCAGGTTCAGGGTCACATGCCTCCGCTCATGATCCCAATTTTTCCACATGACCAGCGGACCCTGGCAG<br>CAGCTGCTGCTGCCCAACAGGGATTCCTCTTCCCCCCTGGAATAACATACAAACCAGGTGATAACTACCCCGTACAGTTC<br>ATTCCATCAACAATGGCAGCTGCTGCTGCTTCTGGACTCAGCCCTTTACAGCTCCAGAAGGGTCATGTCTCCCACCCACA<br>AATTAACCAAAGGCTAAAGGGCCTAAGTGACCGTTTTGGCAGGAATTTGGACACCTTTGAACATGGTGGTGGCCACTCTT<br>ACAACCACAAACAGATTGAGCAGCTCTATGCCGCTCAGCTGGCCAGCATGCAGGTGTCACCTGGAGCAAAGATGCCATCA<br>ACTCCACAGCCACCAAACACAGCAGGGACGGTCTCACCTACTGGGATAAAAAATGAAAAGAGAGGGACCAGCCCTGTAAC<br>TCAAGTTAAGGATGAAGCAGCAGCACAGCCTCTGAATCTCTCATCCCGACCCAAGACAGCAGAGCCTGTAAAGTCCCCAA<br>CGTCTCCCACCCAGAACCTCTTCCCAGCCAGCAAAACCAGCCCTGTCAATCTGCCAAACAAAAGCAGCATCCCTAGCCCC<br>ATTGGAGGAAGCCTGGGAAGAGGATCCTCTTTAGATATCCTATCTAGTCTCAACTCCCCTGCCCTTTTTGGGGATCAGGA<br>TACAGTGATGAAAGCCATTCAGGAGGCGCGGAAGATGCGAGAGCAGATCCAGCGGGAGCAACAGCAGCAACAGCCACATG<br>GTGTTGACGGGAAACTGTCCTCCATAAATAATATGGGGCTGAACAGCTGCAGGAATGAAAAGGAAAGAACGCGCTTTGAG<br>AATTTGGGGCCCCAGTTAACGGGAAAGTCAAATGAAGATGGAAAACTGGGCCCAGGTGTCATCGACCTTACTCGGCCAGA<br>AGATGCAGAGGGAAGTAAAGCAATGAATGGCTCTGCAGCTAAACTACAGCAGTATTATTGTTGGCCAACAGGAGGTGCCA<br>CTGTGGCTGAAGCACGAGTCTACAGGGACGCCCGCGGCCGTGCCAGCAGCGAGCCACACATTAAACCCCGACCGAAACGC<br>ACCTGCATTGTTGATGGCAAAAAGCTTCGGATTGGGGAGTATAAGCAACTGATGAGGTCTCGGAGACAGGAGATGAGGCA<br>GTTCTTTACTGTGGGGCAACAGCCTCAGATTCCAATCACCACAGGAACAGGTGTTGTGTATCCTGGTGCTATCACTATGG<br>CAACTACCACACCATCGCCTCAGATGACATCTGACTGCTCTAGCACCTCGGCCAGCCCGGAGCCCAGCCTCCCGGTCATC<br>CAGAGCACTTATGGTATGAAGACAGATGGCGGAAGCCTAGCTGGAAATGAAATGATCAATGGAGAGGATGAAATGGAAAT |
| SOX7_IDR | JBP095_SOX7_IDR_repair | N/A | N/A | N/A | 83-84 | ATGGCTTCGCTGCTGGGAGCCTACCCTTGGCCCGAGGGTCTCGAGTGCCCGGCCCTGGACGCCGAGCTGTCGGATGGACA<br>ATCGCCGCCGGCCGTCCCCCGGCCCCCGGGGGACAAGGGCTCCGAGAGCCGTATCCGGCGGCCGCGCAGGAAGAAGCAGG<br>CCAAGCGGCTGTGCAAGCGCGTGGACCCGGGCTTCCTTCTGAGCTCCCTCTCCCGGGACCAGAACGCCCTGCCGGAGAAG<br>AGAAGCGGCAGCCGGGGGGCGCTGGGGGAGAAGGAGGACAGGGGTGAGTACTCCCCCGGCACTGCCCTGCCCAGCCTCCG<br>GGGCTGCTACCACGAGGGGCCGGCTGGTGGTGGCGGCGGCGGCACCCCGAGCAGTGTGGACACGTACCCGTACGGGCTGC<br>CCACACCTCCTGAAATGTCTCCCCTGGACGTGCTGGAGCCGGAGCAGACCTTCTTCTCCTCCCCCTGCCAGGAGGAGCAT<br>GGCCATCCCCGCCGCATCCCCCACCTGCCAGGGCACCCGTACTCACCGGAGTACGCCCCAAGCCCTCTCCACTGTAGCCA<br>CCCCCTGGGCTCCCTGGCCCTTGGCCAGTCCCCCGGCGTCTCCATGATGTCCCCTGTACCCGGCTGTCCCCCATCTCCTG<br>CCTATTACTCCCCGGCCACCTACCACCCACTCCACTCCAACCTCCAAGCCCACCTGGGCCAGCTTTCCCCGCCTCCTGAG<br>CACCCTGGCTTCGACGCCCTGGATCAACTGAGCCAGGTGGAACTCCTGGGGGACATGGATCGCAATGAATTCGACCAGTA<br>TTTGAACACTCCTGGCCACCCAGACTCCGCCACAGGGGCCATGGCCCTCAGTGGGCATGTTCCGGTCTCCCAGGTGACAC<br>CAACGGGTCCCACAGAGACCAGCCTCATCTCCGTCCTGGCTGATGCCACGGCCACGTACTACAACAGCTACAGTGTGTCA<br>TAG |
| SOX9_IDR | JBP096_SOX9_IDR_repair | N/A | N/A | N/A | 85-86 | ATGAATCTCCTGGACCCCTTCATGAAGATGACCGACGAGCAGGAGAAGGGCCTGTCCGGCGCCCCCAGCCCCACCATGTC<br>CGAGGACTCCGCGGGCTCGCCCTGCCCGTCGGGCTCCGGCTCGGACACCGAGAACACGCGGCCCCAGGAGAACACGTTCC<br>CCAAGGGCGAGCCCGATCTGAAGAAGGAGAGCGAGGAGGACAAGTTCCCCGTGTGCATCCGCGAGGCGGTCAGCCAGGTG<br>CTCAAAGGCTACGACTGGACGCTGGTGCCCATGCCGGTGCGCGTCAACGGCTCCAGCAAGAACAAGCCGCACGTCCAGCC<br>GCGGCGGAGGAAGTCGGTGAAGAACGGGCAGGCGGAGGCAGAGGAGGCCACGGAGCAGACGCACATCTCCCCCAACGCCA<br>TCTTCAAGGCGCTGCAGGCCGACTCGCCACACTCCTCCTCCGGCATGAGCGAGGTGCACTCCCCCGGCGAGCACTCGGGG<br>CAATCCCAGGGCCCACCGACCCCACCCACCACCCCCAAAACCGACGTGCAGCCGGGCAAGGCTGACCTGAAGCGAGAGGG<br>GCGCCCCTTGCCAGAGGGGGGCAGACAGCCCCCTATCGACTTCCGCGACGTGGACATCGGCGAGCTGAGCAGCGACGTCA<br>TCTCCAACATCGAGACCTTCGATGTCAACGAGTTTGACCAGTACCTGCCGCCCAACGGCCACCCGGGGGTGCCGGCCACG<br>CACGGCCAGGTCACCTACACGGGCAGCTACGGCATCAGCAGCACCGCGGCCACCCCGGCGAGCGCGGGCCACGTGTGGAT<br>GTCCAAGCAGCAGGCGCCGCTGCCACCCCCGCAGCAGCCCCCACAGGCCCCGCCGGCCCCGCAGGCGCCCCCGCAGCCGC<br>AGGCGGCGCCCCCACAGCAGCCGGCGGCACCCCCGCAGCAGCCACAGGCGCACACGCTGACCACGCTGAGCAGCGAGCCG<br>GGCCAGTCCCAGCGAACGCACATCAAGACGGAGCAGCTGAGCCCCAGCCACTACAGCGAGCAGCAGCAGCACTCGCCCCA<br>ACAGATCGCCTACAGCCCCTTCAACCTCCCACACTACAGCCCCTCCTACCCGCCCATCACCCGCTCACAGTACGACTACA<br>CCGACCACCAGAACTCCAGCTCCTACTACAGCCACGCGGCAGGCCAGGGCACCGGCCTCTACTCCACCTTCACCTACATG<br>AACCCCGCTCAGCGCCCCATGTACACCCCCATCGCCGACACCTCTGGGGTCCCTTCCATCCCGCAGACCCACAGCCCCCA<br>GCACTGGGAACAACCCGTCTACACACAGCTCACTCGACCTTAG |
| SOX13_IDR | JBP097_SOX13_IDR_repair | N/A | N/A | N/A | 87-88 | ATGTCCATGAGGAGCCCCATCTCTGCCCAGCTGGCCCTGGATGGCGTTGGCACCATGGTGAACTGCACCATCAAGTCAGA<br>GGAGAAGAAAGAGCCTTGCCACGAGGCCCCCCAGGGCTCAGCCACTGCCGCTGAACCTCAGCCTGGAGACCCAGCCCGGG<br>CCTCCCAGGATAGTGCTGACCCCCAAGCTCCAGCCCAGGGGAATTTCAGGGGCTCCTGGGACTGTAGCTCTCCAGAGGGT<br>AATGGGTCCCCAGAACCCAAGAGACCAGGAGTGTCGGAGGCTGCCTCTGGAAGCCAGGAGAAGCTGGACTTCAACCGAAA<br>TTTGAAAGAAGTGGTGCCAGCCATAGAGAAGCTGTTGTCCAGTGACTGGAAGGAGAGGTTTCTAGGAAGGAACTCTATGG<br>AAGCCAAAGATGTCAAAGGGACCCAAGAGAGCCTAGCAGAGAAGGAGCTCCAGCTTCTGGTCATGATTCACCAGCTGTCC<br>ACCCTGCGGGACCAGCTCCTGACAGCCCACTCGGAGCAGAAGAACATGGCTGCCATGCTGTTTGAGAAGCAGCAGCAGCA<br>GATGGAGCTTGCCCGGCAGCAGCAGGAGCAGATTGCAAAGCAGCAGCAGCAGCTGATTCAGCAGCAGCATAAGATCAACC<br>TCCTTCAGCAGCAGATCCAGCAGGTTAACATGCCTTATGTCATGATCCCAGCCTTCCCCCCAAGCCACCAACCTCTGCCT<br>GTCACCCCTGACTCCCAGCTGGCCTTACCCATTCAGCCCATTCCCTGCAAACCAGTGGAGTATCCGCTGCAGCTGCTGCA<br>CAGCCCCCCTGCCCCAGTGGTGAAGAGGCCTGGGGCCATGGCCACCCACCACCCCCTGCAGGAGCCCTCCCAGCCCCTGA<br>ACCTCACAGCCAAGCCCAAGGCCCCCGAGCTGCCCAACACCTCCAGCTCCCCAAGCCTGAAGATGAGCAGCTGTGTGCCC<br>CGCCCCCCCAGCCATGGAGGCCCCACGCGGGACCTGCAGTCCAGCCCCCCGAGCCTGCCTCTGGGCTTCCTTGGTGAAGG<br>GGACGCTGTCACCAAAGCCATCCAGGATGCTCGGCAGCTGCTGCACAGCCACAGTGGGGCCTTGGATGGCTCCCCCAACA<br>CCCCCTTCCGTAAGGACCTCATCAGCCTGGACTCATCCCCAGCCAAGGAGCGGCTGGAGGACGGCTGTGTGCACCCACTG<br>GAGGAAGCCATGCTGAGCTGCGACATGGATGGCTCCCGCCACTTCCCCGAGTCCCGAAACAGCAGCCACATCAAGCCGCG<br>GCCCAAGCGCACCTGCATCGTGGAGGGCAAGCGGCTGCGCGTGGGAGAGTACAAGGCCCTGATGAGGACCCGGCGTCAGG<br>ATGCCCGCCAGAGCTACGTGATCCCCCCGCAGGCTGGCCAGGTGCAGATGAGCTCCTCAGATGTCCTGTACCCTCGGGCA<br>GCAGGCATGCCGCTGGCACAGCCACTGGTGGAGCACTATGTCCCTCGTAGCCTGGACCCCAACATGCCTGTGATCGTCAA<br>CACCTGCAGCCTCAGAGAGGAGGGTGAGGGCACAGATGACAGGCACTCGGTGGCTGATGGCGAGATGTACCGGTACAGCG<br>AGGACGAGGACTCGGAGGGCGAAGAGAAGAGCGATGGGGAGTTGGTGGTGCTCACAGACTTG |
| SOX17_IDR | JBP098_SOX17_IDR_repair | N/A | N/A | N/A | 89-90 | ATGAGCAGCCCGGATGCGGGATACGCCAGTGACGACCAGAGCCAGACCCAGAGCGCGCTGCCCGCGGTGATGGCCGGGCT<br>GGGCCCCTGCCCCTGGGCCGAGTCGCTGAGCCCCATCGGGGACATGAAGGTGAAGGGCGAGGCGCCGGCGAACAGCGGAG<br>CACCGGCCGGGGCCGCGGGCCGAGCCAAGGGCGAGTCCCGTATCCGGCGGCCGCGGCGGCGCAAGCAGGTGAAGCGGCTG<br>AAGCGGGTGGAGGGCGGCTTCCTGCACGGCCTGGCTGAGCCGCAGGCGGCCGCGCTGGGCCCCGAGGGCGGCCGCGTGGC<br>CATGGACGGCCTGGGCCTCCAGTTCCCCGAGCAGGGCTTCCCCGCCGGCCCGCCGCTGCTGCCTCCGCACATGGGCGGCC<br>ACTACCGCGACTGCCAGAGTCTGGGCGCGCCTCCGCTCGACGGCTACCCGTTGCCCACGCCCGACACGTCCCCGCTGGAC<br>GGCGTGGACCCCGACCCGGCTTTCTTCGCCGCCCCGATGCCCGGGGACTGCCCGGCGGCCGGCACCTACAGCTACGCGCA<br>GGTCTCGGACTACGCTGGCCCCCCGGAGCCTCCCGCCGGTCCCATGCACCCCCGACTCGGCCCAGAGCCCGCGGTCCCT<br>CGATTCCGGGCCTCCTGGCGCCACCCAGCGCCCTTCACGTGTACTACGGCGCGATGGGCTCGCCCGGGGCGGGCGGCGGG<br>CGCGGCTTCCAGATGCAGCCGCAACACCAGCACCAGCACCAGCACCAGCACCACCCCCCGGGCCCCGGACAGCCGTCGCC<br>CCCTCCGGAGGCACTGCCCTGCCGGGACGGCACGGACCCCAGTCAGCCCGCCGAGCTCCTCGGGGAGGTGGACCGCACGG<br>AATTTGAACAGTATCTGCACTTCGTGTGCAAGCCTGAGATGGGCCTCCCCTACCAGGGGCATGACTCCGGTGTGAATCTC<br>CCCGACAGCCACGGGGCCATTTCCTCGGTGGTGTCCGACGCCAGCTCCGCGGTATATTACTGCAACTATCCTGACGTG |
| SOX30_IDR | JBP099_SOX30_IDR_repair | N/A | N/A | N/A | 91-92 | ATGGAGAGAGCCAGACCCGAGCCGCCGCCTCAGCCGCGCCCGTTGCGTCCCGCTCCGCCCCCGCTGCCGGTCGAGGGCAC<br>CTCCTTTTGGGCAGCAGCCATGGAGCCCCCTCCGTCGTCTCCCACACTGAGCGCGGCAGCCAGTGCGACCTTGGCCTCGT<br>CGTGCGGGGAGGCAGTGGCGTCCGGCTTACAGCCCGCGGTGCGGCGGCTGCTGCAGGTGAAGCCAGAGCAGGTGTTGCTG<br>CTACCACAGCCTCAGGCCCAGAACGAGGAAGCCGCTGCCTCGTCCGCGCAGGCGCGGCTGTTGCAGTTCAGGCCCGACCT<br>GCGGCTCCTGCAGCCGCCGACAGCGTCAGACGGCGCCACCTCCAGGCCCGAGTTGCACCCGGTGCAGCCCCTGGCGCTGC<br>ATGTCAAGGCCAAGAAGCAGAAGCTGGGGCCCAGCCTGGATCAGTCAGTGGGGCCTCGAGGGGCCGTCGAAACCGGTCCT<br>AGAGCCTCCAGGGTGGTCAAGTTGGAAGGCCCCGGGCCGGCCCTCGGCTACTTCCGAGGGGACGAGAAGGGCAAGCTGGA<br>GGCGGAGGAGGTCATGAGAGACTCGATGCAAGGCGGGGCAGGCAAAAGCCCGGCAGCCATCCGAGAAGGTGTGATCAAAA<br>CGGAGGAACCCGAGAGACTCCTCGAGGACTGCAGGCTCGGCGCGGAGCCCGCGTCCAATGGCCTGGTTCATGGCAGCGCG<br>GAGGTCATCTTGGCCCCAACGTCCGGTGCCTTTGGGCCGCACCAGCAAGACCTTAGGATCCCTTTGACGCTCCACACGGT<br>CCCCCCTGGGGCCCGGATCCAGTTTCAGGGAGCTCCGCCTTCAGAGCTGATAAGATTGACCAAGGTCCCCCTGACACCAG<br>TGCCTACTAAAATGCAGTCCCTACTGGAGCCTTCTGTAAAAATTGAAACCAAAGATGTCCCGCTCACCGTGTTGCCCTCA<br>GATGCAGGCATACCAGATACTCCCTTCAGTAAGGACAGAAATGGTCATGTGTATCAGCCTCGTCCAGGGAAGCGAAAACG<br>ATTCCCTCTAAGTGTTTCCAATGTATTTTCTGGTACCACACAGAATATCATCTCTACAAATCCTACAACAGTTTATCCTT<br>ACCGCTCACCTACGTACTCTGTGGTAATTCCCAGCCTACAGAATCCCATCACTCATCCAGTTGTACATGCCCTTACAGTC<br>GGCCTCCCTTTGGCTATGGAAATTTTCCGAGTTCAATGCCAGAATGCCTTAGTTATTATGAAGACAGGTACCCAAAACAT<br>GAGGGTATCTTTTCAACTTTAG |
| SOX5_DBD | SOX5 strain | JBP100_SOX5_DBD_F | JBP101_SOX5_DBD_R | 58 | ura | AAGCGTCCAATGAATGCCTTCATGGTGTGGGCTAAAGATGAACGGAGAAAGATCCTTCAAGCCTTTCCTGACATGCACAA<br>CTCCAACATCAGCAAGATATTGGGATCTCGCTGGAAAGCTATGACAAACCTAGAGAAACAGCCATATTATGAGGAGCAAG<br>CCCGTCTCAGCAAGCAGCACCTGGAGAAGTACCCTGACTATAAGTAC |
| SOX6_DBD | SOX6 strain | JBP102_SOX6_DBD_F | JBP103_SOX6_DBD_R | 65 | ura | AAGCGACCAATGAATGCATTCATGGTTTGGGCAAAGGATGAGAGGAGAAAAATCCTTCAGGCCTTCCCCGACATGCATAA<br>CTCCAACATTAGCAAAATCTTAGGATCTCGCTGGAAATCAATGTCCAACCAGGAGAAGCAACCTTATTATGAAGAGCAGG<br>CCCGGCTAAGCAAGATCCACTTAGAGAAGTACCCAAACTATAAATAC |
| SOX7_DBD | SOX7 strain | JBP105_SOX7_DBD_F | JBP106_SOX7_DBD_R | 65 | ura | CGGCCCATGAACGCCTTCATGGTTTGGGCCAAGGACGAGAGGAAACGGCTGGCAGTGCAGAACCCGGACCTGCACAACGC<br>CGAGCTCAGCAAGATGCTGGGAAAGTCGTGGAAGGCGCTGACGCTGTCCCAGAAGAGGCCGTACGTGGACGAGGCGGAGC<br>GGCTGCGCCTGCAGCACATGCAGGACTACCCCAACTACAAGTAC |
| SOX9_DBD | SOX9 strain | JBP107_SOX9_DBD_F | JBP108_SOX9_DBD_R | 58 | ura | AAGCGGCCCATGAACGCCTTCATGGTGTGGGCGCAGGCGGCGCGCAGGAAGCTCGCGGACCAGTACCCGCACTTGCACAA<br>CGCCGAGCTCAGCAAGACGCTGGGCAAGCTCTGGAGACTTCTGAACGAGAGCGAGAAGCGGCCCTTCGTGGAGGAGGCGG<br>AGCGGCTGCGCGTGCAGCACAAGAAGGACCACCCGGATTACAAGTAC |
| SOX13_DBD | SOX13 strain | JBP109_SOX13_DBD_F | JBP110_SOX13_DBD_R | 58 | ura | AAGAGGCCCATGAACGCCTTCATGGTGTGGGCCAAGGATGAGCGGAGGAAGATCCTGCAAGCCTTCCCAGACATGCACAA<br>CTCCAGCATCAGCAAGATCCTTGGATCTCGCTGGAAGTCCATGACCAACCAGGAGAAGCAGCCCTACTATGAGGAACAGG<br>CGCGGCTGAGCCGGCAGCACCTGGAGAAGTATCCTGACTACAAGTAC |
| SOX17_DBD | SOX17 strain | JBP111_SOX17_DBD_F | JBP112_SOX17_DBD_R | 58 | ura | CGGCCGATGAACGCTTTCATGGTGTGGGCTAAGGACGAGCGCAAGCGGCTGGCGCAGCAGAATCCAGACCTGCACAACGC<br>CGAGTTGAGCAAGATGCTGGGCAAGTCGTGGAAGGCGCTGACGCTGGCGGAGAAGCGGCCCTTCGTGGAGGAGGCAGAGC<br>GGCTGCGCGTGCAGCACATGCAGGACCACCCCAACTACAAGTAC |
| SOX30_DBD | SOX30 strain | JBP113_SOX30_DBD_F | JBP114_SOX30_DBD_R | 62 | ura | AAGCGACCCATGAACGCATTTATGGTTTGGGCAAGGATCCACCGACCAGCACTAGCCAAAGCTAACCCAGCAGCCAACAA<br>TGCAGAAATCAGTGTCCAGCTTGGGTTAGAGTGGAACAAACTTAGTGAAGAACAAAAGAAACCCTATTACGATGAAGCAC<br>AAAAGATTAAGGAAAAGCACAGAGAGGAATTTCCTGGTTGGGTT |

| Oligo type | Name | Description | Sequence |
|---|---|---|---|
| primer | JBP004 | Sits on MNase of Josh_gBlock (F) | CGATAAGTATGGACGCGGTCTGG |
| primer | ADO10 | HO locus primer (R) | AAACAAATCAGTGCCGGTAACGC |
| primer | JBP005 | pDEST_AD TF plasmid lift (F) with homology to Josh_Gblock | GGATCACGATATCGATTACAAGGATGATGATGATAAGgcggccacaagtttgtacaaaaaagcaggcacc |
| primer | JBP006 | pDEST_AD TF plasmid lift (R) with homology to Josh_gBlock | GCTTATTTAGAAGTGGCGCGCCTCTAagaagtggcgcgccctacatcaaaccactttgtacaagaaagctgg |
| primer | JBP007_ATF4_F | ATF4 (F) with homology to Josh_Gblock | ggatcacgatatcgattacaaggatgatgatgataaggcggccATGACCGAAATGAGCTTCCTGAGC |
| primer | JBP008_FOS_F | FOS (F) with homology to Josh_Gblock | ggatcacgatatcgattacaaggatgatgatgataaggcggccATGATGTTCTCGGGCTTCAACGC |
| primer | JBP009_FOXP3_F | FOXP3 (F) with homology to Josh_Gblock | ggatcacgatatcgattacaaggatgatgatgataaggcggccATGCCCAACCCCAGGCCTG |
| primer | JBP010_FOXA2_F | FOXA2 (F) with homology to Josh_Gblock | ggatcacgatatcgattacaaggatgatgatgataaggcggccATGCTGGGAGCGGTGAAGATG |
| primer | JBP011_FLI1_F | FLI1 (F) with homology to Josh_Gblock | ggatcacgatatcgattacaaggatgatgatgataaggcggccATGGACGGGACTATTAAGGAGGCTC |
| primer | JBP012_FOXP1_F | FOXP1 (F) with homology to Josh_Gblock | ggatcacgatatcgattacaaggatgatgatgataaggcggccATGATGCAAGAATCTGGGACTGAGAC |
| primer | JBP013_HOXA11_F | HOXA11 (F) with homology to Josh_Gblock | ggatcacgatatcgattacaaggatgatgatgataaggcggccATGGATTTTGATGAGCGTGGTCCC |
| primer | JBP014_HOXA10_F | HOXA10 (F) with homology to Josh_Gblock | ggatcacgatatcgattacaaggatgatgatgataaggcggccATGTCATGCTCGGAGAGCCCC |
| primer | JBP015_HOXC9_F | HOXC9 (F) with homology to Josh_Gblock | ggatcacgatatcgattacaaggatgatgatgataaggcggccATGTCGGCGACGGGGC |
| primer | JBP017_SOX15_F | SOX15 (F) with homology to Josh_Gblock | ggatcacgatatcgattacaaggatgatgatgataaggcggccATGGCGCTACCAGGCTCCTC |
| primer | JBP018_SOX17_F | SOX17 (F) with homology to Josh_Gblock | ggatcacgatatcgattacaaggatgatgatgataaggcggccATGAGCAGCCCGGATGCG |
| primer | JBP019_TGIF2_F | TGIF2 (F) with homology to Josh_Gblock | ggatcacgatatcgattacaaggatgatgatgataaggcggccATGTCGGACAGTGATCTAGGTGAGG |
| primer | JBP020_POU2F3_F | POU2F3 (F) with homology to Josh_Gblock | ggatcacgatatcgattacaaggatgatgatgataaggcggccATGGTGAATCTGGAGTCCATGCAC |
| primer | JBP021_POU3F4_F | POU3F4 (F) with homology to Josh_Gblock | ggatcacgatatcgattacaaggatgatgatgataaggcggccATGGCCACAGCTGCCTCG |
| primer | JBP022_POU3F1_F | POU3F1 (F) with homology to Josh_Gblock | ggatcacgatatcgattacaaggatgatgatgataaggcggccATGGCCACCACCGCGC |
| primer | JBP024_GATA3_F | GATA3 (F) with homology to Josh_Gblock | ggatcacgatatcgattacaaggatgatgatgataaggcggccATGGAGGTGACGGCGGACC |
| primer | JBP025_FOXF1_F | FOXF1 (F) with homology to Josh_Gblock | ggatcacgatatcgattacaaggatgatgatgataaggcggccATGGACCCCGCGTCGTC |
| primer | JBP026_SOX13_F | SOX13 (F) with homology to Josh_Gblock | ggatcacgatatcgattacaaggatgatgatgataaggcggccATGTCCATGAGGAGCCCCATCTC |
| primer | JBP028_FOXL1_F | FOXL1 (F) with homology to Josh_Gblock | ggatcacgatatcgattacaaggatgatgatgataaggcggccATGAGTCACCTCTTCGATCCCCG |
| primer | JBP029_FOXL2_F | FOXL2 (F) with homology to Josh_Gblock | ggatcacgatatcgattacaaggatgatgatgataaggcggccATGATGGCCAGCTACCCCGAG |
| primer | JBP030_FOXP2_F | FOXP2 (F) with homology to Josh_Gblock | ggatcacgatatcgattacaaggatgatgatgataaggcggccATGATGCAGGAATCTGCGACAGAG |
| primer | JBP031_ELF2_F | ELF2 (F) with homology to Josh_Gblock | ggatcacgatatcgattacaaggatgatgatgataaggcggccATGACATCAGCAGTGGTTGACAGTG |
| primer | JBP032_MNT_F | MNT (F) with homology to Josh_Gblock | ggatcacgatatcgattacaaggatgatgatgataaggcggccATGAGCATAGAGACGCTACTGGAGG |
| primer | JBP033_MLXIPL_F | MLXIPL (F) with homology to Josh_Gblock | ggatcacgatatcgattacaaggatgatgatgataaggcggccATGGCTGGCGCGCTGG |
| primer | JBP034_MLX_F | MLX (F) with homology to Josh_Gblock | ggatcacgatatcgattacaaggatgatgatgataaggcggccATGACGGAGCCTGGAGCTTCTC |
| primer | JBP035_MXD4_F | MXD4 (F) with homology to Josh_Gblock | ggatcacgatatcgattacaaggatgatgatgataaggcggccATGGAGCTGAACTCCCTGCTGATC |
| primer | JBP036_HOXB9_F | HOXB9 (F) with homology to Josh_Gblock | ggatcacgatatcgattacaaggatgatgatgataaggcggccATGTCCATTTCTGGGACGCTTAGC |
| primer | JBP037_HOXC13_F | HOXC13 (F) with homology to Josh_Gblock | ggatcacgatatcgattacaaggatgatgatgataaggcggccATGACGACTTCGCTGCTCCTG |
| primer | JBP038_SOX11_F | SOX11 (F) with homology to Josh_Gblock | ggatcacgatatcgattacaaggatgatgatgataaggcggccATGGTGCAGCAGGCGGAG |
| primer | JBP039_SOX4_F | SOX4 (F) with homology to Josh_Gblock | ggatcacgatatcgattacaaggatgatgatgataaggcggccATGGTGCAGCAAACCAACAATGC |
| primer | JBP040_SOX5_F | SOX5 (F) with homology to Josh_Gblock | ggatcacgatatcgattacaaggatgatgatgataaggcggccATGCTTACTGACCCTGATTTACCTCAGG |
| primer | JBP041_GATA6_F | GATA6 (F) with homology to Josh_Gblock | ggatcacgatatcgattacaaggatgatgatgataaggcggccATGGCCTTGACTGACGGCG |
| primer | JBP043_attL2_pENTR223.1 | attL2 site of pENTR223.1 (R) with homology to Josh_Gblock | GCTTATTTAGAAGTGGCGCGCCTCTAagaagtggcgcgccctaCTATCTATCTACCAACTTTGTACAAGAAAGCTGGGC |
| primer | JBP044_attL2_pENTR223 | attL2 site of pENTR223 (R) with homology to Josh_Gblock | GCTTATTTAGAAGTGGCGCGCCTCTAagaagtggcgcgccctaCTATCTATCTATGCCAACTTTGTACAAGAAAGTTGGGT |
| primer | JBP045_attL2_pDONR221_201 | attL2 site of pDONR201/221 (R) with homology to Josh_Gblock | GCTTATTTAGAAGTGGCGCGCCTCTAagaagtggcgcgccctaCTATCTATCTAGCCAACTTTGTACAAGAAAGCTGGGT |
| primer | JBP100_SOX5_DBD_F | SOX DBD primer with homology to Josh G-block | ggatcacgatatcgattacaaggatgatgatgataaggcggccAAGCGTCCAATGAATGCCTTCATGG |
| primer | JBP101_SOX5_DBD_R | SOX DBD primer with homology to Josh G-block | GCTTATTTAGAAGTGGCGCGCCTCTAagaagtggcgcgccctaGTACTTATAGTCAGGGTACTTCTCCAGGTG |
| primer | JBP102_SOX6_DBD_F | SOX DBD primer with homology to Josh G-block | ggatcacgatatcgattacaaggatgatgatgataaggcggccAAGCGACCAATGAATGCATTCATGG |
| primer | JBP103_SOX6_DBD_R | SOX DBD primer with homology to Josh G-block | CTTATTTAGAAGTGGCGCGCCTCTAagaagtggcgcgccctaGTATTTATAGTTTGGGTACTTCTCTAAGTGGATCTTGC |
| primer | JBP105_SOX7_DBD_F | SOX DBD primer with homology to Josh G-block | ggatcacgatatcgattacaaggatgatgatgataaggcggccCGGCCCATGAACGCCTTCATG |
| primer | JBP106_SOX7_DBD_R | SOX DBD primer with homology to Josh G-block | GCTTATTTAGAAGTGGCGCGCCTCTAagaagtggcgcgccctaGTACTTGTAGTTGGGGTAGTCCTGCATG |
| primer | JBP107_SOX9_DBD_F | SOX DBD primer with homology to Josh G-block | ggatcacgatatcgattacaaggatgatgatgataaggcggccAAGCGGCCCATGAACGCC |
| primer | JBP108_SOX9_DBD_R | SOX DBD primer with homology to Josh G-block | GCTTATTTAGAAGTGGCGCGCCTCTAagaagtggcgcgccctaGTACTTGTAATCCGGGTGGTCCTTCTTG |
| primer | JBP109_SOX13_DBD_F | SOX DBD primer with homology to Josh G-block | ggatcacgatatcgattacaaggatgatgatgataaggcggccAAGAGGCCCATGAACGCCTTC |
| primer | JBP110_SOX13_DBD_R | SOX DBD primer with homology to Josh G-block | GCTTATTTAGAAGTGGCGCGCCTCTAagaagtggcgcgccctaGTACTTGTAGTCAGGATACTTCTCCAGGTG |
| primer | JBP111_SOX17_DBD_F | SOX DBD primer with homology to Josh G-block | ggatcacgatatcgattacaaggatgatgatgataaggcggccCGGCCGATGAACGCTTTCATG |
| primer | JBP112_SOX17_DBD_R | SOX DBD primer with homology to Josh G-block | GCTTATTTAGAAGTGGCGCGCCTCTAagaagtggcgcgccctaGTACTTGTAGTTGGGGTGGTCCTG |
| primer | JBP113_SOX30_DBD_F | SOX DBD primer with homology to Josh G-block | ggatcacgatatcgattacaaggatgatgatgataaggcggccAAGCGACCCATGAACGCATTTATGG |
| primer | JBP114_SOX30_DBD_R | SOX DBD primer with homology to Josh G-block | GCTTATTTAGAAGTGGCGCGCCTCTAagaagtggcgcgccctaAACCCAACCAGGAAATTCCTCTCTGTG |
| primer | JBP115_FOXL2_DBD_FOXO3_IDR_F | FOXL2 DBD primer with homology to other FOX IDR | GCCGAGGAAATGTTCGTCGCGGCGGAACGCCTGGGGAAACAAGCCCCCGTACTCGTACGTG |
| primer | JBP116_FOXL2_DBD_FOXO3_IDR_R | FOXL2 DBD primer with homology to other FOX IDR | CCATGGAGACAGCCCGCCGCCGGGGGGCTTTTCCGCTCTTCTTCTCGAACATGTCTTCGCAGGC |
| primer | JBP117_FOXL2_DBD_FOXA2_IDR_F | FOXL2 DBD primer with homology to other FOX IDR | CGACCCCAAGACCTACAGGCGCAGCTACACGCACGCAAAGAAGCCCCCGTACTCGTACGTG |
| primer | JBP118_FOXL2_DBD_FOXA2_IDR_R | FOXL2 DBD primer with homology to other FOX IDR | TCTCGCACTTGAAGCGCTTCTGGCGGCGCAGGTAGCAGCCCTTCTCGAACATGTCTTCGCAGGC |
| primer | JBP121_FOXL2_DBD_FOXF1_IDR_F | FOXL2 DBD primer with homology to other FOX IDR | CAAGGCCAAGAAGACCAACGCCGGCATCCGGCGCCCGGAGAAGCCCCCGTACTCGTACGTG |
| primer | JBP122_FOXL2_DBD_FOXF1_IDR_R | FOXL2 DBD primer with homology to other FOX IDR | TCCTTCGGAAGCCGCGCGGCCGCCGCCGAAAGGAGCCCTCCTTCTCGAACATGTCTTCGCAGGC |
| primer | JBP123_FOXL2_DBD_FOXP3_IDR_F | FOXL2 DBD primer with homology to other FOX IDR | CCTCCACAACATGGACTACTTCAAGTTCCACAACATGCGAAAGCCCCCGTACTCGTACGTG |

| | | | |
|---|---|---|---|
| primer | JBP124_FOXL2_DBD_FOXP3_IDR_R | FOXL2 DBD primer with homology to other FOX IDR | GTGTAGGGTTGGAACACCTGCTGGGCCTCTGGCTCCGTTTCTTCTCGAACATGTCTTCGCAGGC |
| primer | JBP125_FOXL2_DBD_FOXP1_IDR_F | FOXL2 DBD primer with homology to other FOX IDR | TGCGCAGAACCAAGAATTTTATAAGAACGCAGAAGTTAGAAAGCCCCCGTACTCGTACGTG |
| primer | JBP126_FOXL2_DBD_FOXP1_IDR_R | FOXL2 DBD primer with homology to other FOX IDR | TAAGGGAAGGGTTACCACTGATCTTTTGTGGCCTTCGTTTCTTCTCGAACATGTCTTCGCAGGC |
| primer | JBP127_FOXL2_DBD_FOXP2_IDR_F | FOXL2 DBD primer with homology to other FOX IDR | TGCCCCAAACTATGAATTTTATAAAAATGCAGATGTCAGAAAGCCCCCGTACTCGTACGTG |
| primer | JBP190_FOXL2_DBD_FOXP2_IDR_R2 | FOXL2 DBD primer with homology to other FOX IDR | TTTTTACTAAGGTTGGACTTCCTGTTATCTTTTGTGACCTCTTCTCGAACATGTCTTCGCAGGC |
| primer | JBP151_ERG_DBD_ELK4_F | ERG DBD primer with homology to other GABPA (Ets) IDR | tttgtacaaaaaagcaggcaccATGGACAGTGCTATCACCCTTTGGCAGTTCCTCCTGGAGC |
| primer | JBP152_ERG_DBD_ELK4_R | ERG DBD primer with homology to other GABPA (Ets) IDR | TGCCCACTGTCATTGGATCCATGTTCAAAATCTCTGGATAGTCGAACTTGTAGGCGTAGCGC |
| primer | JBP153_ERG_DBD_ELK1_F | ERG DBD primer with homology to other GABPA (Ets) IDR | tttgtacaaaaaagcaggcaccATGGACCCATCTGTGACGCTTTGGCAGTTCCTCCTGGAGC |
| primer | JBP154_ERG_DBD_ELK1_R | ERG DBD primer with homology to other GABPA (Ets) IDR | GCGGGCAGTCCTCAGTGGAGCACCCTGCGACCTCAGGGTAGTCGAACTTGTAGGCGTAGCGC |
| primer | JBP155_ERG_DBD_FLI1_F | ERG DBD primer with homology to other GABPA (Ets) IDR | CAGCAGTCGCCTAGCCAACCCTGGAAGCGGGCAGATCCAGCTTTGGCAGTTCCTCCTGGAGC |
| primer | JBP156_ERG_DBD_FLI1_R | ERG DBD primer with homology to other GABPA (Ets) IDR | CGGTCGGATGTGGCTGCAGAGCCTGGGCAATGCCGTGGAAGTCGAACTTGTAGGCGTAGCGC |
| primer | JBP157_ERG_DBD_ERF_F | ERG DBD primer with homology to other GABPA (Ets) IDR | CTACAAGCCAGAGTCGTCCCCTGGCTCAAGGCAGATCCAGCTTTGGCAGTTCCTCCTGGAGC |
| primer | JBP158_ERG_DBD_ERF_R | ERG DBD primer with homology to other GABPA (Ets) IDR | CATCAATGAATGGGTAATTGACCAGCACCAGTTTGTTGAAGTCGAACTTGTAGGCGTAGCGC |
| primer | JBP159_ERG_DBD_ELF1_F | ERG DBD primer with homology to other GABPA (Ets) IDR | GAAGAAAAACAAAGATGGAAAGGGAAACACAATTTATCTTCTTTGGCAGTTCCTCCTGGAGC |
| primer | JBP160_ERG_DBD_ELF1_R | ERG DBD primer with homology to other GABPA (Ets) IDR | CCTCATCATTTATATATATAAGATCTTTTGGCATTTCTTTGTCGAACTTGTAGGCGTAGCGC |
| primer | JBP161_ERG_DBD_ELF2_F | ERG DBD primer with homology to other GABPA (Ets) IDR | GAAGAAACCAAGAGAAGGAAAAGGAAACACAACCTATTTGCTTTGGCAGTTCCTCCTGGAGC |
| primer | JBP162_ERG_DBD_ELF2_R | ERG DBD primer with homology to other GABPA (Ets) IDR | TTTTGTCATCATCTATGACCACTATGTTTTTCGGCATATCGTCGAACTTGTAGGCGTAGCGC |
| primer | JBP165_ELF1_DBD_ELK4_F | ELF1 DBD primer with homology to other GABPA (Ets) IDR | tttgtacaaaaaagcaggcaccATGGACAGTGCTATCACCTGGGAGTTTTTACTGGCACTGCTC |
| primer | JBP166_ELF1_DBD_ELK4_R | ELF1 DBD primer with homology to other GABPA (Ets) IDR | TGCCCACTGTCATTGGATCCATGTTCAAAATCTCTGGATAAAACTGATACACCAAGCGCTGACC |
| primer | JBP167_ELF1_DBD_ELK1_F | ELF1 DBD primer with homology to other GABPA (Ets) IDR | tttgtacaaaaaagcaggcaccATGGACCCATCTGTGACGTGGGAGTTTTTACTGGCACTGCTC |
| primer | JBP168_ELF1_DBD_ELK1_R | ELF1 DBD primer with homology to other GABPA (Ets) IDR | GCGGGCAGTCCTCAGTGGAGCACCCTGCGACCTCAGGGTAAAACTGATACACCAAGCGCTGACC |
| primer | JBP169_ELF1_DBD_FLI1_F | ELF1 DBD primer with homology to other GABPA (Ets) IDR | CAGCAGTCGCCTAGCCAACCCTGGAAGCGGGCAGATCCAGTGGGAGTTTTTACTGGCACTGCTC |
| primer | JBP170_ELF1_DBD_FLI1_R | ELF1 DBD primer with homology to other GABPA (Ets) IDR | CGGTCGGATGTGGCTGCAGAGCCTGGGCAATGCCGTGGAAAAACTGATACACCAAGCGCTGACC |
| primer | JBP171_ELF1_DBD_ERF_F | ELF1 DBD primer with homology to other GABPA (Ets) IDR | CTACAAGCCAGAGTCGTCCCCTGGCTCAAGGCAGATCCAGTGGGAGTTTTTACTGGCACTGCTC |
| primer | JBP172_ELF1_DBD_ERF_R | ELF1 DBD primer with homology to other GABPA (Ets) IDR | CATCAATGAATGGGTAATTGACCAGCACCAGTTTGTTGAAAAACTGATACACCAAGCGCTGACC |
| primer | JBP173_ELF1_DBD_ELF2_F | ELF1 DBD primer with homology to other GABPA (Ets) IDR | GAAGAAACCAAGAGAAGGAAAAGGAAACACAACCTATTTGTGGGAGTTTTTACTGGCACTGCTC |
| primer | JBP174_ELF1_DBD_ELF2_R | ELF1 DBD primer with homology to other GABPA (Ets) IDR | TTTTGTCATCATCTATGACCACTATGTTTTTCGGCATATCAAACTGATACACCAAGCGCTGACC |
| primer | JBP175_ELF1_DBD_ERG_F | ELF1 DBD primer with homology to other GABPA (Ets) IDR | AAGTAGCCGCCTTGCAAATCCAGGCAGTGGCCAGATCCAGTGGGAGTTTTTACTGGCACTGCTC |
| primer | JBP176_ELF1_DBD_ERG_R | ELF1 DBD primer with homology to other GABPA (Ets) IDR | CCGGGGGGTGGGGCTGGAGGGCCTGGGCGATCCCGTGGAAAAACTGATACACCAAGCGCTGACC |
| gRNA | FOXO3_sense | sgRNA guide for cutting the DBD with GTTTT/GATCA sequence | TATGCAGTGACAGGTTGTGCGTTTT |
| gRNA | FOXO3_antisense | sgRNA guide for cutting the DBD with GTTTT/GATCA sequence | GCACAACCTGTCACTGCATAGATCA |
| gRNA | FOXA2_sense | sgRNA guide for cutting the DBD with GTTTT/GATCA sequence | AGCGAGATGTACGAGTAGGGGTTTT |
| gRNA | FOXA2_antisense | sgRNA guide for cutting the DBD with GTTTT/GATCA sequence | CCCTACTCGTACATCTCGCTGATCA |
| gRNA | FOXJ2_sense | sgRNA guide for cutting the DBD with GTTTT/GATCA sequence | GATGGCATAGGTGATGAGAGGTTTT |
| gRNA | FOXJ2_antisense | sgRNA guide for cutting the DBD with GTTTT/GATCA sequence | CTCTCATCACCTATGCCATCGATCA |
| gRNA | FOXF1_sense | sgRNA guide for cutting the DBD with GTTTT/GATCA sequence | AGCGCGATGTAGGAATAGGGGTTTT |
| gRNA | FOXF1_antisense | sgRNA guide for cutting the DBD with GTTTT/GATCA sequence | CCCTATTCCTACATCGCGCTGATCA |
| gRNA | FOXP3_sense | sgRNA guide for cutting the DBD with GTTTT/GATCA sequence | TGTGCAGACTCAGGTTGTGGGTTTT |
| gRNA | FOXP3_antisense | sgRNA guide for cutting the DBD with GTTTT/GATCA sequence | CCACAACCTGAGTCTGCACAGATCA |
| gRNA | FOXP1_sense | sgRNA guide for cutting the DBD with GTTTT/GATCA sequence | ACGCACTGCATTCTTCCACGGTTTT |
| gRNA | FOXP1_antisense | sgRNA guide for cutting the DBD with GTTTT/GATCA sequence | CGTGGAAGAATGCAGTGCGTGATCA |
| gRNA | FOXP2_sense | sgRNA guide for cutting the DBD with GTTTT/GATCA sequence | TCAGGCGTAATGCAGCAACTGTTTT |
| gRNA | FOXP2_antisense | sgRNA guide for cutting the DBD with GTTTT/GATCA sequence | AGTTGCTGCATTACGCCTGAGATCA |
| gRNA | ELK4_sense | sgRNA guide for cutting the DBD with GTTTT/GATCA sequence | AAGAGGTGGCTCGTCTCTGGGTTTT |
| gRNA | ELK4_antisense | sgRNA guide for cutting the DBD with GTTTT/GATCA sequence | CCAGAGACGAGCCACCTCTTGATCA |
| gRNA | ELK1_sense | sgRNA guide for cutting the DBD with GTTTT/GATCA sequence | GTGAAGTCCAGGAGATGATGGTTTT |
| gRNA | ELK1_antisense | sgRNA guide for cutting the DBD with GTTTT/GATCA sequence | CATCATCTCCTGGACTTCACGATCA |
| gRNA | ERG_sense | sgRNA guide for cutting the DBD with GTTTT/GATCA sequence | TGTCCGACAGGAGCTCCAGGGTTTT |
| gRNA | ERG_antisense | sgRNA guide for cutting the DBD with GTTTT/GATCA sequence | CCTGGAGCTCCTGTCGGACAGATCA |
| gRNA | FLI1_sense | sgRNA guide for cutting the DBD with GTTTT/GATCA sequence | TGTCGGAGAGCAGCTCCAGGGTTTT |
| gRNA | FLI1_antisense | sgRNA guide for cutting the DBD with GTTTT/GATCA sequence | CCTGGAGCTGCTCTCCGACAGATCA |
| gRNA | ERF_sense | sgRNA guide for cutting the DBD with GTTTT/GATCA sequence | GCATTCTGCACAAGACCAAGGTTTT |
| gRNA | ERF_antisense | sgRNA guide for cutting the DBD with GTTTT/GATCA sequence | CTTGGTCTTGTGCAGAATGCGATCA |
| gRNA | ELF1_sense | sgRNA guide for cutting the DBD with GTTTT/GATCA sequence | CTCAGGTACTATTACCAAAGGTTTT |
| gRNA | ELF1_antisense | sgRNA guide for cutting the DBD with GTTTT/GATCA sequence | CTTTGGTAATAGTACCTGAGGATCA |
| gRNA | ELF2_sense | sgRNA guide for cutting the DBD with GTTTT/GATCA sequence | GACATGAACTATGAAACCATGTTTT |
| gRNA | ELF2_antisense | sgRNA guide for cutting the DBD with GTTTT/GATCA sequence | ATGGTTTCATAGTTCATGTCGATCA |
| gRNA | SOX5_sense | sgRNA guide for cutting the DBD with GTTTT/GATCA sequence | AATGAATGCCTTCATGGTGTGTTTT |
| gRNA | SOX5_antisense | sgRNA guide for cutting the DBD with GTTTT/GATCA sequence | ACACCATGAAGGCATTCATTGATCA |
| gRNA | SOX6_sense | sgRNA guide for cutting the DBD with GTTTT/GATCA sequence | TGTTGGAGTTATGCATGTCGGTTTT |
| gRNA | SOX6_antisense | sgRNA guide for cutting the DBD with GTTTT/GATCA sequence | CGACATGCATAACTCCAACAGATCA |
| gRNA | SOX7_sense | sgRNA guide for cutting the DBD with GTTTT/GATCA sequence | GTCCCAGAAGAGGCCGTACGGTTTT |
| gRNA | SOX7_antisense | sgRNA guide for cutting the DBD with GTTTT/GATCA sequence | CGTACGGCCTCTTCTGGGACGATCA |
| gRNA | SOX9_sense | sgRNA guide for cutting the DBD with GTTTT/GATCA sequence | CCTTCGTGGAGGAGGCGGAGGTTTT |
| gRNA | SOX9_antisense | sgRNA guide for cutting the DBD with GTTTT/GATCA sequence | CTCCGCCTCCTCCACGAAGGGATCA |
| gRNA | SOX13_sense | sgRNA guide for cutting the DBD with GTTTT/GATCA sequence | CCTACTATGAGGAACAGGCGGTTTT |
| gRNA | SOX13_antisense | sgRNA guide for cutting the DBD with GTTTT/GATCA sequence | CGCCTGTTCCTCATAGTAGGGATCA |
| gRNA | SOX17_sense | sgRNA guide for cutting the DBD with GTTTT/GATCA sequence | GCCGAGTTGAGCAAGATGCTGTTTT |
| gRNA | SOX17_antisense | sgRNA guide for cutting the DBD with GTTTT/GATCA sequence | AGCATCTTGCTCAACTCGGCGATCA |
| gRNA | SOX30_sense | sgRNA guide for cutting the DBD with GTTTT/GATCA sequence | GATTAAGGAAAAGCACAGAGGTTTT |
| gRNA | SOX30_antisense | sgRNA guide for cutting the DBD with GTTTT/GATCA sequence | CTCTGTGCTTTTCCTTAATCGATCA |
| gRNA | ura_sense | sgRNA guide for cutting the ura sequence with GTTTT/GATCA sequence | GGTGTGGGTTTAGATGACAAGTTTT |
| gRNA | ura_antisense | sgRNA guide for cutting the ura sequence with GTTTT/GATCA sequence | TTGTCATCTAAACCCACACCGATCA |
| ss oligo | JBP093_SOX5_IDR_repair | Repair DNA for CRISPR to create SOX IDR-only strain | TAGGGAATCCCGAGGGCGTGGTAGCAATGAACCCCACATAAAGCCCAGGCCAAAGCGCACATGCCTGGTGGATGGCAAAA |
| ss oligo | JBP094_SOX6_IDR_repair | Repair DNA for CRISPR to create SOX IDR-only strain | CAGGGACGCCCGCGGCCGTGCCAGCAGCGAGCCACACATTAAACCCCGACCGAAACGCACCTGCATTGTTGATGGCAAAA |
| ss oligo | JBP095_SOX7_IDR_repair | Repair DNA for CRISPR to create SOX IDR-only strain | CCGGCCCCCGGGGGACAAGGGCTCCGAGAGCCGTATCCGGCGGCCGCGCAGGAAGAAGCAGGCCAAGCGGCTGTGCAAGC |
| ss oligo | JBP096_SOX9_IDR_repair | Repair DNA for CRISPR to create SOX IDR-only strain | GGTGCGCGTCAACGGCTCCAGCAAGAACAAGCCGCACGTCCAGCCGCGGCGGAGGAAGTCGGTGAAGAACGGGCAGGCGG |
| ss oligo | JBP097_SOX13_IDR_repair | Repair DNA for CRISPR to create SOX IDR-only strain | CTCCCGCCACTTCCCCGAGTCCCGAAACAGCAGCCACATCAAGCCGCGGCCCAAGCGCACCTGCATCGTGGAGGGCAAGC |
| ss oligo | JBP098_SOX17_IDR_repair | Repair DNA for CRISPR to create SOX IDR-only strain | CGGGGCCGCGGGCCGAGCCAAGGGCGAGTCCCGTATCCGGCGGCCGCGGCGGCGCAAGCAGGTGAAGCGGCTGAAGCGGG |
| ss oligo | JBP099_SOX30_IDR_repair | Repair DNA for CRISPR to create SOX IDR-only strain | ACCAGATACTCCCTTCAGTAAGGACAGAAATGGTCATGTGTATCAGCCTCGTCCAGGGAAGCGAAAACGATTCCCTCTAA |
